# Tales of 1,008 Small Molecules: Phenomic Profiling through Live-cell Imaging in a Panel of Reporter Cell Lines

**DOI:** 10.1101/2020.03.13.990093

**Authors:** Michael J. Cox, Steffen Jaensch, Jelle Van de Waeter, Laure Cougnaud, Daan Seynaeve, Soulaiman Benalla, Seong Joo Koo, Ilse Van Den Wyngaert, Jean-Marc Neefs, Dmitry Malkov, Mart Bittremieux, Margino Steemans, Pieter J. Peeters, Jörg Kurt Wegner, Hugo Ceulemans, Emmanuel Gustin, Yolanda T. Chong, Hinrich W.H. Göhlmann

## Abstract

Phenomic profiles are high-dimensional sets of readouts that can comprehensively capture the biological impact of chemical and genetic perturbations in cellular assay systems. Phenomic profiling of compound libraries can be used for compound target identification or mechanism of action (MoA) prediction and other applications in drug discovery. To devise an economical set of phenomic profiling assays, we assembled a library of 1,008 approved drugs and well-characterized tool compounds manually annotated to 218 unique MoAs, and we profiled each compound at four concentrations in live-cell, high-content imaging screens against a panel of 15 reporter cell lines, which expressed a diverse set of fluorescent organelle and pathway markers in three distinct cell lineages. For 41 of 83 testable MoAs, phenomic profiles accurately ranked the reference compounds (AUC-ROC ≥0.9). MoAs could be better resolved by screening compounds at multiple concentrations than by including replicates at a single concentration. Screening additional cell lineages and fluorescent markers increased the number of distinguishable MoAs but this effect quickly plateaued. There remains a substantial number of MoAs that were hard to distinguish from others under the current study’s conditions. We discuss ways to close this gap, which will inform the design of future phenomic profiling efforts.

## Introduction

In the last two decades, technological and analytical innovations in high-throughput microscopy and transcriptomics have enabled the large-scale phenomic profiling of small-molecule libraries [1–8]. These profiling approaches quantify the phenotypic response of cells to compound treatment by simultaneously measuring changes in hundreds or thousands of features, be they transcript levels assessed with gene expression technologies or the morphological characteristics of cells in a microscopy image. The resulting multivariate readouts capture the impact of specific chemistry across many biological processes at play in the context of a given cell population, which makes them more informative than traditional uni- or low-dimensional assays. Phenomic profiling has proven its value in drug discovery for target / mechanism of action (MoA) identification, hit extension, hit diversification, and hit triaging efforts [9,10].

The most straightforward strategy for characterizing compound MoA via a phenomic profiling assay is to include a set of reference compounds that display high selectivity for targets of interest, and to base MoA predictions for uncharacterized compounds on their proximity to these reference compounds in the phenotypic space. This nearest-reference approach is data efficient, however there is no attempt to learn MoA-specific feature weighting. This is of particular importance as poly-pharmacology, i.e., the simultaneous engagement of multiple targets, is the rule rather than the exception for small molecules [11], and a nearest-reference approach will either assign a molecule exhibiting poly-pharmacology to the phenomically most dominant activity only, or will not assign the molecule at all if reference compounds with similar phenomic profiles are not present in the experiment.

For poly-pharmacological compounds, morphological and transcriptional readouts reflect the actions on multiple, functionally unrelated targets that are convolved into a complex phenotype, and their activity on specific and relevant targets must therefore be deconvolved from these phenomic profiles, which is far from trivial. Deconvolution can be achieved using supervised machine learning models to predict target activity [7]. This approach requires that many of the phenomically profiled compounds have also been tested in the target- or MoA-specific assays it aims to cover. These compounds are used as training examples to identify the most informative phenomic features to predict activities in each assay of interest. Consequently, the data scale required for supervised deconvolution can be prohibitive.

A continuing challenge in drug discovery is the design of an affordable set of phenomic profiling assays with maximal coverage of therapeutic targets. Ideally, this assay set should be able to document a phenotypic response towards at least the vast majority of the 549 human proteins targeted by 999 FDA-approved small-molecule drugs for human disease [12]. Until recently, most studies that use phenomic profiling have been restricted in scope, typically using fewer than 100 reference compounds annotated to a limited number of targets [1,2,4,13–15] and are not sufficiently broadly representative to inform design principles for generic sets of phenomic profiling assays.

In the current study, we used a nearest-reference approach to explore how various screening parameters affect the ability to distinguish MoAs from each other. We used gene-editing to create a panel of 15 reporter cell lines by introducing different combinations of 12 blue fluorescent protein (BFP), green fluorescent protein (GFP), or red fluorescent protein (RFP)/fusionRed signalling pathway and organelle markers into the A549, HepG2, and WPMY1 cell backgrounds, and profiled these reporter cell lines in live-cell, high-content imaging screens against a library of 1,008 small molecules, manually annotated with 218 unique MoA descriptors, at four concentrations.

## Results

### Library of 1,008 Reference Compounds and 169 Natural Products

We assembled a set of 1,008 well-characterized reference compounds, composed of FDA-approved drugs and commercially available tool compounds, and manually annotated each compound with one or more MoA descriptors (Supplementary Table S1). In total, 218 unique MoA descriptors were assigned to the reference compound set. Of the 1,008 reference compounds, 829 (82%) were labelled with only a single MoA descriptor. Of the 218 MoAs, 132 (61%) were assigned to ≥3 co-annotated compounds and 92 (42%) were assigned to ≥5 co-annotated compounds (Supplementary Fig. S1). To increase coverage of the phenotypic space, 169 natural products were added to this chemical library. As an additional source of annotation, the gene names of the reference compound targets were downloaded from the ChEMBL and IUPHAR databases [16,17]. In total, 690 compounds could be mapped to the gene names of 692 unique targets (median number of targets per compound = 2) (Supplementary Table S1; Supplementary Fig. S1).

### Panel of 15 Reporter Cell Lines

Monitoring the morphologies of multiple subcellular organelles and the activation status of different intracellular signalling pathways improves the predictive power of phenomic profiling [4,13], so we constructed a panel of reporter cell lines for use in live-cell screens. We selected three cell types – each from a different lineage – for gene-editing to express a diverse set of fluorescent markers: the A549 lung adenocarcinoma, the HepG2 hepatocellular carcinoma, and the WPMY1 prostatic stromal myofibroblast lines. Each reporter cell line carried a BFP segmentation cassette that enabled detection of the nuclear and cell boundaries. In addition, each reporter cell line expressed a GFP- and RFP/fusionRed-tagged organelle or pathway marker from its endogenous chromosomal locus. Five different combinations of organelle and pathway markers were introduced into each cell type. Tagging of these proteins did not significantly perturb their cellular localization or molecular function (Supplementary Fig. S2). These cell lines are available for purchase from MilliporeSigma.

### Experimental and Analytical Pipeline

Our experimental and analytical pipeline is outlined in Figure 1. Live cells were imaged 24 hours after being treated with compound. Following segmentation with automated image analysis software, features were extracted from each cell and data from live cells was aggregated at the well-level. A modified version of the minimum redundancy maximum relevance (mRMR) algorithm [18] was used to select an informative subset of features for each reporter cell line; between 22 and 58 features were sufficient to capture the phenotypic diversity present in each of the 15 reporter cell line screens. We refer to the retained normalized features as “imaging signature” representing the phenotype induced by a compound treatment. Active reference compound treatments were ranked and the area under the receiver operating characteristic curve (AUC-ROC) was computed. MoAs with high AUC-ROC values (≥0.9) were considered distinguishable from other MoAs.

**Figure 1:**
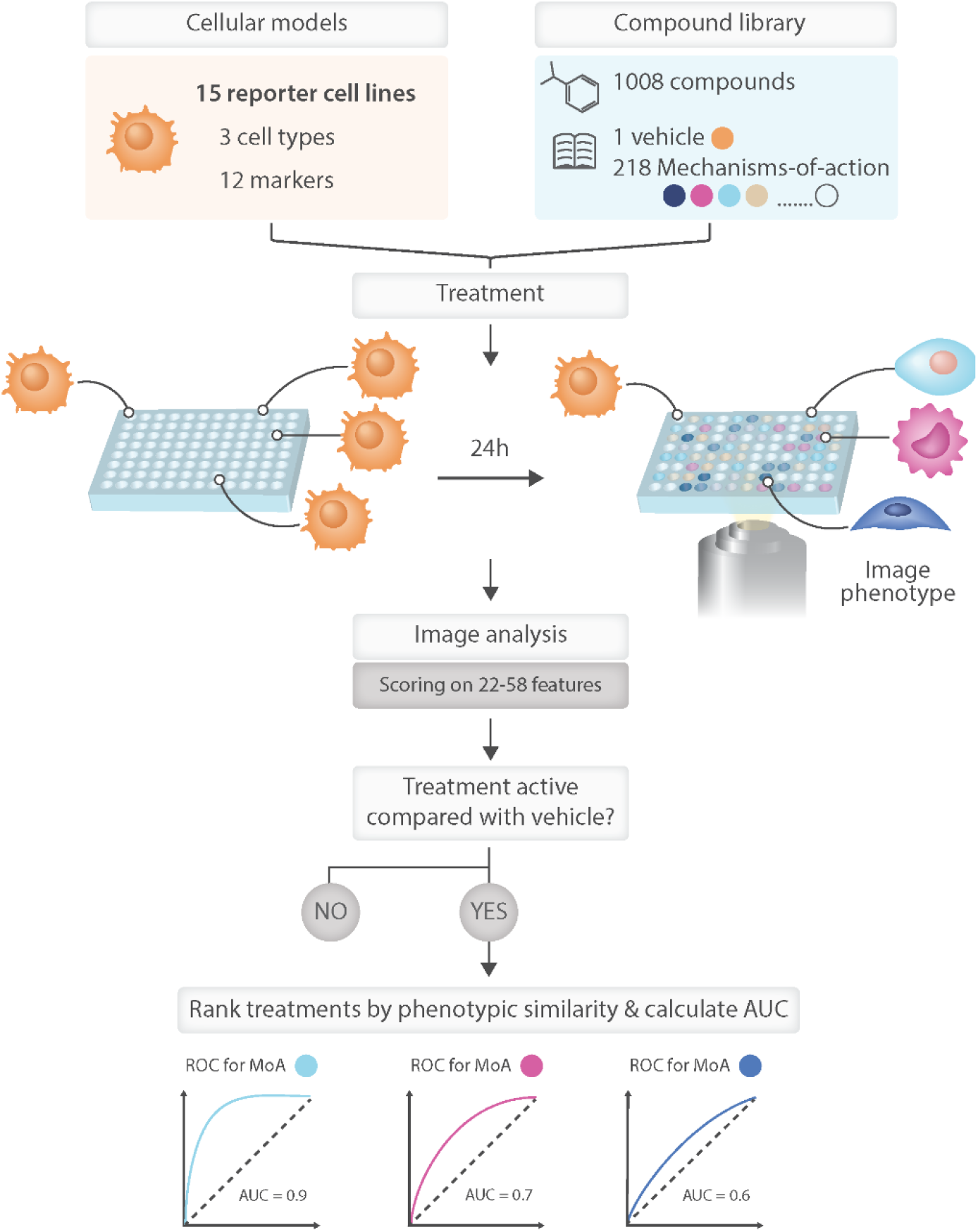
Overview of experimental and analytical steps in phenotypic screen. Fifteen reporter cell lines (three cell types × five marker combinations) were screened against a chemical library that contained 1,008 reference compounds which had been manually annotated with MoA information. Compounds were screened at four concentrations (0.3, 1, 3, and 9 μM) and live cells were imaged 24 hours after treatment. Following segmentation, ~500 features were computed per cell, aggregated at well-level and z-score normalized against vehicle (DMSO control). The mRMR algorithm was then used to select an informative subset of non-redundant features that exhibited high reproducibility across replicate compound treatments for each reporter cell line, referred to as imaging signature, which describes the phenotypic response to each compound treatment. A compound treatment was deemed to be phenotypically active if its imaging signature was significantly different from that of DMSO control and highly similar across replicates. To quantify the extent to which each MoA with ≥3 active members in a single reporter cell line could be distinguished from other MoAs, we computed AUC-ROC values based on ranking all active treatments by the Pearson correlations between their imaging signatures.

### Compound Activity

The compound library contained 1,177 unique chemicals, 628 (53%) of which were phenotypically active in ≥1 cell line. Most of these active compounds (Supplementary Table S2) induced significant and reproducible phenotypic responses in multiple reporter cell lines (323-455 active compounds per cell line). Indeed, using just three reporter cell lines, we could capture 92% of the active compounds in the library (Supplementary Fig. S3a). Reference compounds were more likely to be phenotypically active than natural products (57% versus 33% active, respectively) and they also tended to exhibit this activity at lower concentrations, probably because they were designed to be potent and to have drug-like physiochemical properties (Supplementary Fig. S3b). Strikingly, reference compounds co-annotated to the same MoA almost always display similar activity in each of the reporter cell lines, indicating that the MoA annotation is largely consistent with whether or not a compound exhibits phenotypic activity (Fig. 2a and Supplementary Table S3).

**Figure 2:**
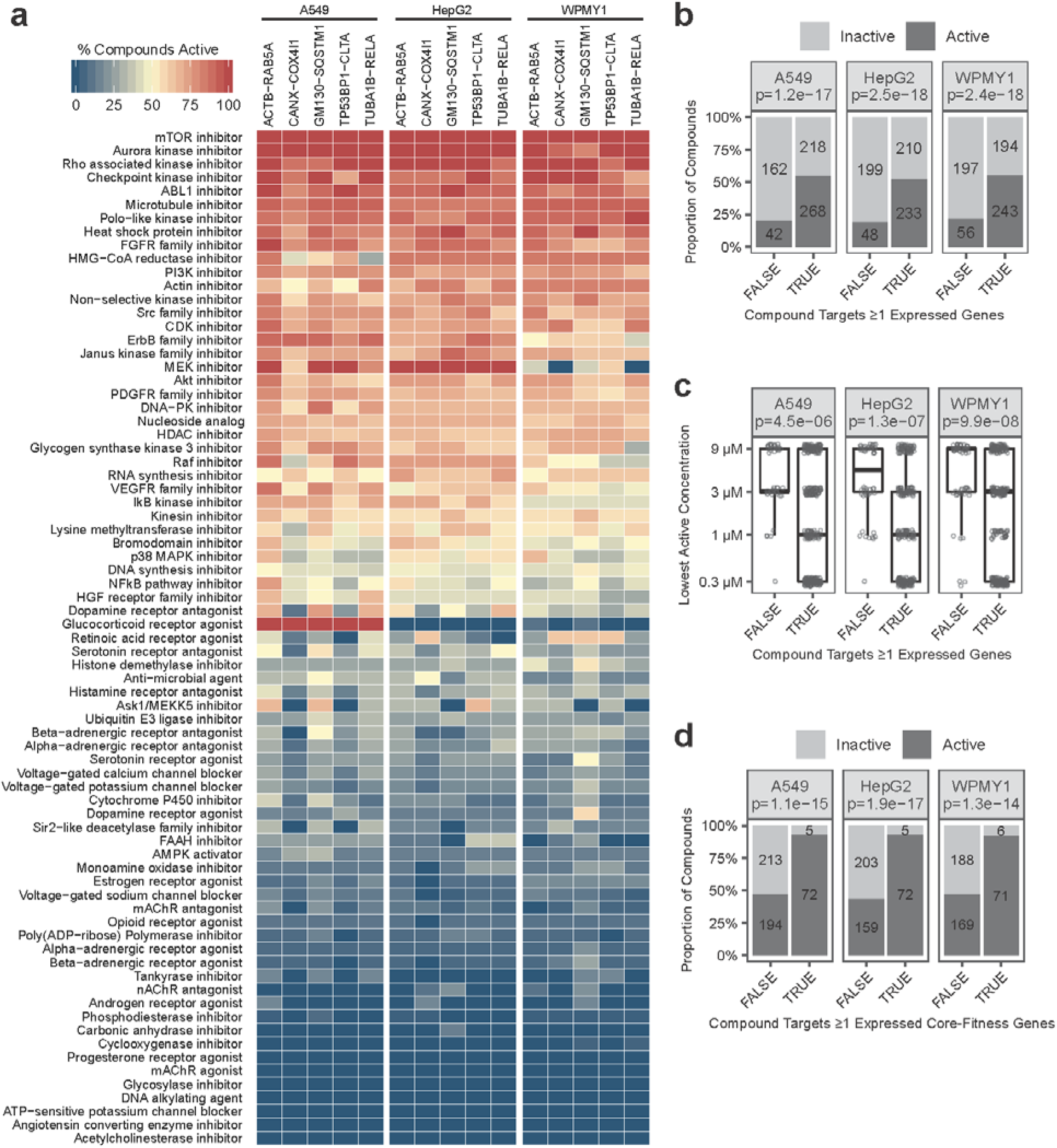
Compounds annotated to expressed targets and core-fitness genes are more likely to have phenotypic activity. (a) Heat map showing the percentage of compounds that induced significant phenotypic activity for MoAs with ≥6 compounds tested (complete data in Supplementary Table S2 and S3). (b) Compounds annotated to expressed targets were more likely to be active than compounds annotated to targets that are not expressed. Target expression was determined using microarray data. Each dark-grey bar shows the number of compounds that are active in ≥1 of the five cell lines within each cell type; each light-grey bar shows the number of compounds that are not active in any of the five cell lines within each cell type. Fisher’s exact test p-values are indicated below the cell type names in the header. (c) Active compounds annotated to expressed targets tended to be active at lower concentrations than active compounds annotated to targets that are not expressed. The vertical axis shows the lowest active concentration across the five cell lines within each cell type for each active compound. Wilcoxon rank sum test p-values are indicated below the cell type names in the header. (d) Compounds that target core-fitness genes were more likely to be active than compounds that target non-core-fitness genes. Only compounds annotated to ≥1 expressed target were considered. As in (a), target expression was determined using microarray data. Each dark-grey bar shows the number of compounds that are active in ≥1 of the five cell lines within each cell type, and each light-grey bar shows the number of compounds that are not active in any of the five cell lines within each cell type. Fisher’s exact test p-values are indicated below the cell type names in the header.

Our chemical library contained 690 reference compounds and natural products that had been annotated to 692 unique gene targets. Using microarray experiments we determined that each of the A549, HepG2, and WPMY1 cell types expressed ~45% of the annotated target genes (Supplementary Fig. S4a and S4b). Compounds mapped to ≥1 expressed target were much more likely to display phenotypic activity than compounds that targeted genes that were not expressed (~53% *versus* ~22% active; Fisher’s exact test p≤1.2×10^−17^) (Fig. 2b). That compounds mapped to non-expressed targets primarily show off-target activity is suggested by the fact that they tend to exhibit their initial phenotypic activity at much higher concentrations than compounds whose targets are expressed (Fig. 2c).

The glucocorticoid receptor (GR) agonists provide the most striking example of a group of co-annotated compounds that displayed differential phenotypic activity between cell lines. Although the NR3C1 gene that encodes GR is expressed in all of our reporter lines, the GR agonists only exhibited significant phenotypic activity in the A549 cell background (Fig. 2a and Supplementary Fig. S5). This differential activity is mirrored by a much stronger transcriptional response to GR agonists in A549 cells than in HepG2 and WPMY1 cells (Supplementary Fig. S5a), including stronger changes in A549 cells in gene sets annotated to GR-related gene ontology terms and, interestingly, to cytoplasm organization gene ontology terms (Supplementary Fig. S5b). We suspect that one or more post-transcriptional regulatory mechanisms, including differential splicing of the NR3C1 transcript and the use of alternate translational start sites, regulation of GR nuclear localization, and the cell-type-dependent association with different co-repressors and co-activators (reviewed in [19]), explains the differential activity of GR agonists observed in our reporter cell lines.

Approximately 50% of the compounds annotated to targets expressed in our reporter cell lines were not phenotypically active (Fig. 2b). We believe that a major reason for this might be the ability of cells to adapt to the perturbation of a target because of the inherent redundancy built into their genetic networks and signalling pathways. Genome-wide CRISPR-Cas9 screens have identified a group of core-fitness genes that are required by multiple, genetically diverse cell lines for their optimal proliferation and viability in culture [20,21], and whose function cannot be completely buffered by redundant elements in the cell. We found that >90% of compounds targeting core-fitness genes expressed in our cell lines were phenotypically active (Fig. 2d). In contrast, <50% of the compounds annotated to expressed genes that were not part of the core-fitness group were active. This observation argues that when panels of different cell lines are being selected for MoA prediction screens, a combination of lineages that maximizes the number of drug targets that are required for optimal fitness in ≥1 cellular context would increase the number of active compounds observed.

### Using Phenotypes to Infer Mechanism of Action

For a phenotype to be interpretable, it must be linked to a group of reference compounds acting on the same target or on targets that function in the same biological pathway. The diversity of the chemically-induced phenotypes observed in a given reporter cell line can be represented graphically by projecting the imaging signatures of active compound treatments onto a two-dimensional t-SNE map [22] (Fig. 3 and Supplementary Fig. S6). These maps show that some co-annotated compounds form coherent clusters (e.g., kinesin inhibitors) in phenotypic space whereas others do not (e.g., vascular endothelial growth factor receptor [VEGFR] inhibitors and Abelson murine leukemia viral oncogene homolog 1 [ABL1] inhibitors).

**Figure 3:**
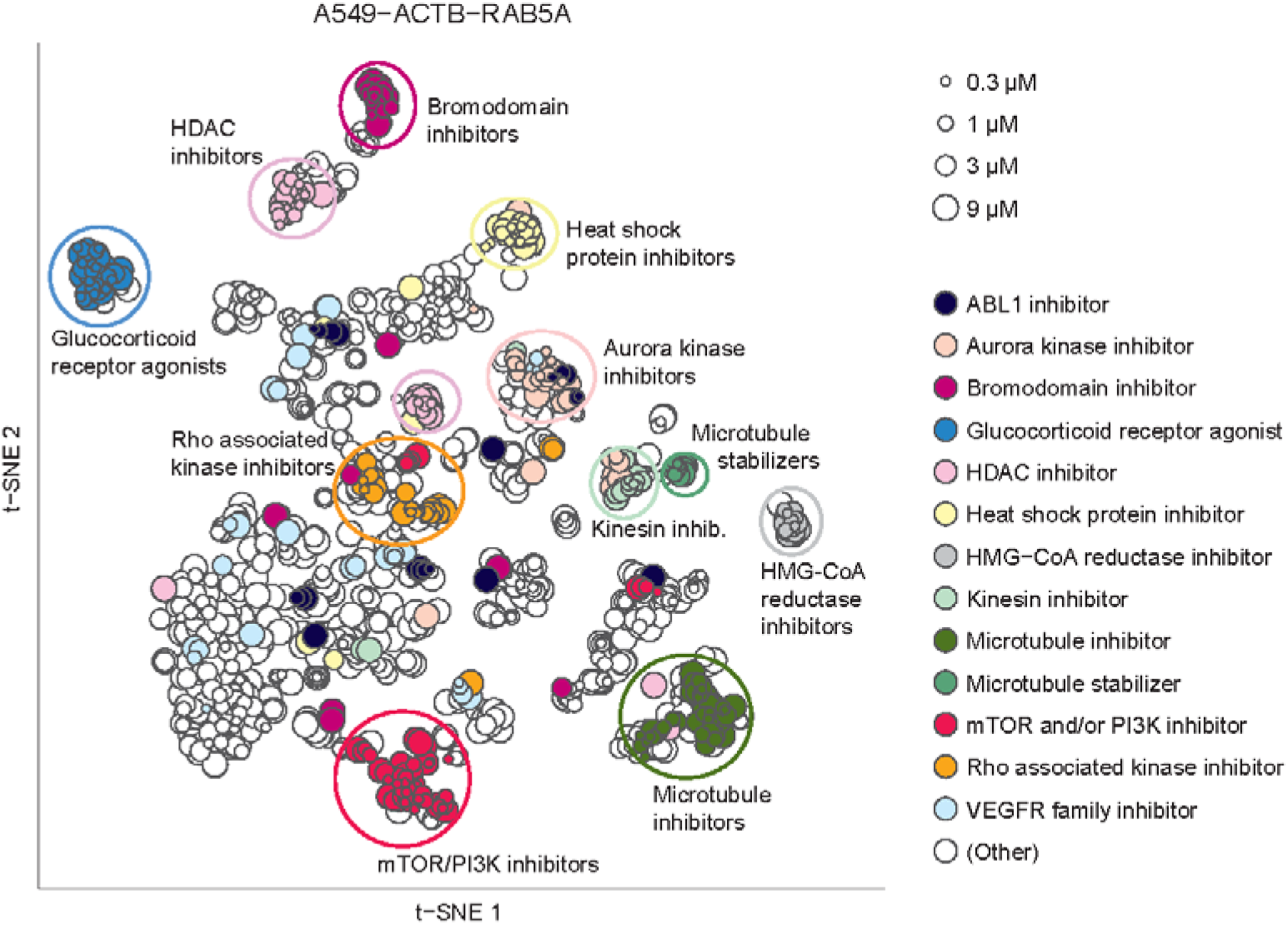
t-SNE map of all active treatments in the A549-ACTB-RAB5A cell line based on the Pearson correlation between their imaging signatures. Each point corresponds to a unique treatment. Selected MoAs are indicated by colours, 11 of which form distinct phenotypic clusters, while ABL1 inhibitors and VEGFR family inhibitors are examples of MoAs that showed diverse phenotypes and hence poor clustering. Circles indicating pheno-clusters were drawn manually.

The coherence of phenomic clustering of a MoA with ≥3 active members can be formally evaluated by ranking all signatures in a single reporter cell line based on their Pearson correlations and computing the AUC-ROC value for the categorization as MoA member versus non-member [5,13] (Fig. 4a, Supplementary Fig. S7, Supplementary Fig. S8, and Supplementary Table S3). Using AUC-ROC ≥0.9 as a cut-off, we found that 41 of the 83 MoAs were distinguishable from other MoAs in one or more reporter cell lines. Of the 48 MoAs with ≥5 active compounds, we found 17 MoAs with an AUC-ROC value ≥0.9 in one or more reporter cell lines, 11 of which exceeded this AUC-ROC threshold in >50% of the reporter cell lines.

**Figure 4:**
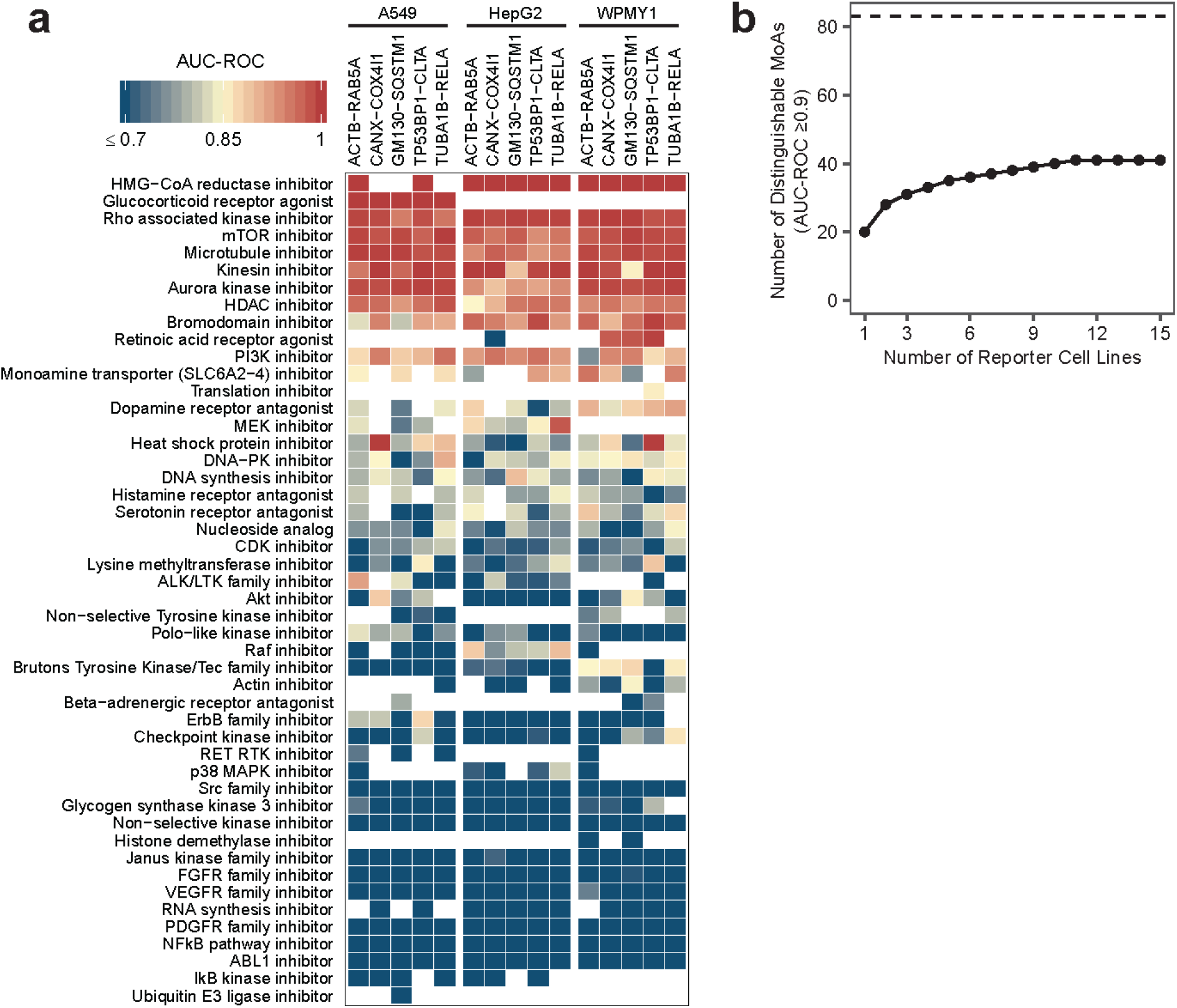
Summary how well each MoA can be distinguished from other MoAs based on compound ranking by phenotypic similarity. (a) Heat map showing AUC-ROC values for all MoAs with ≥5 active compounds in the respective cell line (complete data in Supplementary Fig. S9 and Supplementary Table S3); empty entries (white) correspond to MoAs with <5 active compounds. (b) The number of distinguishable MoAs (AUC-ROC ≥0.9) increased with additional reporter cell lines, but this effect quickly plateaued. We considered all 83 MoAs (indicated by the dashed line) with ≥3 active compounds and ranked the cell lines by how many (additional) distinguishable MoAs they contribute.

Overall, the AUC-ROC values for most of the 83 testable MoAs were similar across most reporter cell lines, indicating that no particular combination of fluorescent markers and cell lineage dramatically out-performed all others (Supplementary Fig. S8). One of the few groups for which we observed substantial cell-lineage dependency is the mitochondrial toxicants. As detection of mitochondrial toxicity is of broad interest in drug discovery, we further investigated this group which includes oxidative phosphorylation uncouplers (CCCP, FCCP, malonoben, BAM-15), oxidative phosphorylation inhibitors (oligomycin A), and respiratory chain inhibitors (rotenone, antimycin A, phenformin, 3-nitropropionic acid). Taken together as one MoA, the mitochondrial toxicants reached an AUC-ROC value ≥0.9 in all five A549 cell lines and markedly lower AUC-ROC values in the HepG2 and WPMY1 cell lines (Supplementary Fig. S9a). In particular, for the A549-CANX-COX4I1 cell line, we observed a tight cluster on the t-SNE map that includes all active concentrations of the mitochondrial toxicants, with a concentration-dependent exception for rotenone, which clusters with the mitochondrial toxicants at 0.3 μM but with microtubule inhibitors at higher concentrations (Supplementary Fig. S9b). The latter observation is consistent with a reported off-target effect of rotenone on microtubules [23]. Additionally, we found a number of compounds annotated for MoAs not related to mitochondrial toxicity to consistently cluster with the known mitochondrial toxicants in all five A549 cell lines, including the ErbB family inhibitor TAK 165, the fatty acid amide hydrolase (FAAH) inhibitor PF 3845, and the mitogen-activated protein kinase kinase (MEK) and IκB kinase inhibitor arctigenin. Indeed, mitochondrial toxicity of these three compounds was confirmed in the Glu/Gal assay (Supplementary Table S4), and literature evidence exists that PF 3845 and arctigenin affect mitochondrial activity [24,25].

### Exploring Characteristics of Distinguishable MoAs

We wanted to better understand why some active co-annotated compounds induced coherent phenotypic clusters with high AUC-ROC values whereas others did not. In general, compounds sharing the same MoA had low chemical similarity (mean Tanimoto index <0.5 for all but four MoAs), and we observed no correlation between the chemical similarity among co-annotated compounds and the AUC-ROC values (r^2^ = 0.004) (Supplementary Fig. S10a and S10b). We theorized that compounds annotated to distinguishable MoAs might induce the strongest overall phenotypic responses in our reporter cell lines or that they might exhibit the strongest perturbation of a single fluorescent marker or phenotypic feature. Neither hypothesis was true. The AUC-ROC calculated for a given MoA did not correlate with the Euclidean distance of their imaging signatures from that of DMSO-treated controls, nor did the absolute z-score of the single most perturbed feature in each imaging signature (Supplementary Fig. S10c and S10d). We did find that active compounds that target the products of core-fitness genes were on average 2.4 times more likely to be linked to distinguishable MoAs than compounds only annotated to expressed targets not in this gene set (Supplementary Fig. S11), but this difference was statistically significant (Fisher’s exact test p<0.05) in only six of the 15 reporter cell lines. Further research is needed to confirm a link between core-fitness genes and distinguishability of MoAs.

The number of distinguishable MoAs increased with each additional cell line that was screened (Fig. 4b), but this effect quickly plateaued. From the 83 MoAs having ≥3 active compounds, maximally 41 could be distinguished, and >90% of the distinguishable MoAs were already covered with seven cell lines. Also increasing the number of cell types or the number of marker pairs increased the number of distinguishable MoAs, but again, this effect soon reached a plateau. As might be expected, the number of phenotypically active compounds per cell line increased with increasing compound concentration: from 109 active MoA-annotated compounds per cell line on average at 0.3 μM to 343 at 9 μM. Overall, MoAs could be better distinguished if compounds were screened at multiple concentrations (Fig. 5a) and this increase was on average 2.7 times larger compared to screening compounds at a single concentration but with multiple replicates (Supplementary Fig. S12). This probably reflects the fact that the reference compounds in our library have a range of potencies for their annotated targets and no single concentration is optimal for maximizing on-target effects and minimizing off-target effects for all compounds.

**Figure 5:**
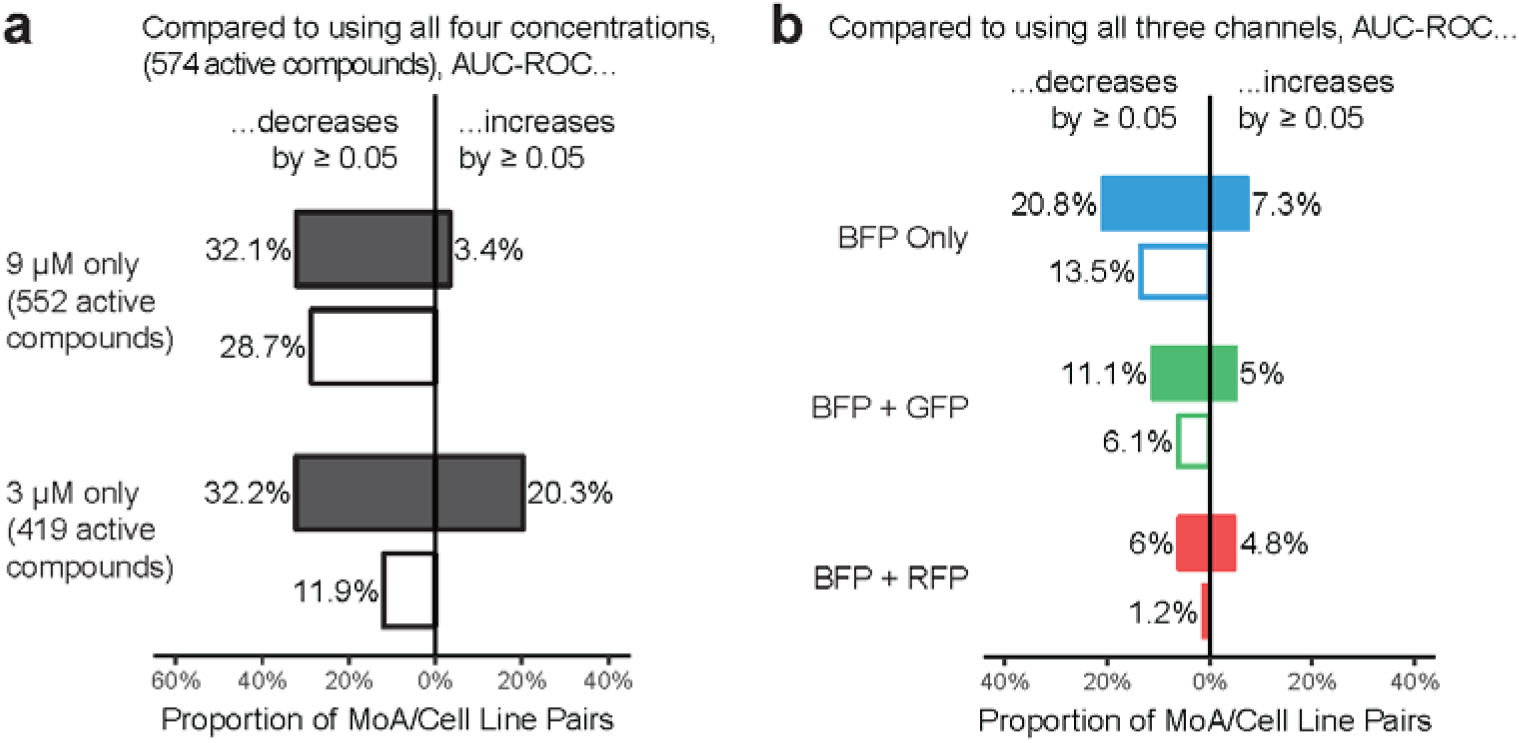
Summary of the degree to which different experimental factors affected how well MoAs can be distinguished. (a) When using only the 9 μM concentration compared to all four concentrations, the AUC-ROC decreased by ≥0.05 for 32.1% (113) of the MoA/cell line pairs, whereas the AUC-ROC increased by ≥0.05 for only 3.4% (12) of the MoA/cell line pairs, resulting in a net AUC-ROC decrease of ≥0.05 for 28.7% (101) of the MoA/cell line pairs. Using only the 3 μM concentration resulted in a similar trend but compared to the “9 μM only”-case more MoA/cell line pairs benefited from the exclusion of the other concentrations. In each case all active treatments at the respective concentration(s) were considered for the AUC-ROC computation; for the “3 μM only”-case this means substantially fewer compounds than for the “9 μM only”- and “all concentrations”-cases. (b) When using only the information from the BFP channel compared to using the information from all three fluorescent channels, we observed a net AUC-ROC decrease of ≥0.05 for only 13.5% of the MoA/cell line pairs, indicating that most of the information is contained in the BFP channel alone. Adding the GFP channel reduces the net loss to 6.1%; adding the RFP channel reduces the net loss to 1.2%. We have not observed strong patterns as to which kind of markers (structural versus signalling / bright versus dim) are most beneficial for MoA distinction, or for which MoAs distinction is particularly sensitive across cell types to the exclusion of particular markers. In all cases, the AUC-ROC is based on the exact same set of active treatments (active calling based on the “all three channels”-signature). For the AUC-ROC analysis we repeated the mRMR feature selection for each considered combination of channels (rather than just dropping features related to the excluded channel(s) from the “all three channels”-signature) for an unbiased comparison between the different combinations of channels.

We used a similar analysis to understand the contributions made by the different fluorescent markers in each reporter cell line to the AUC-ROC values calculated for the different MoAs. The same BFP markers were expressed in every cell line and were used to segment the cell and nucleus. Using the intensity, texture, and shape features derived from BFP fluorescence alone did not dramatically (>0.05) reduce the AUC-ROC values for almost 80% of MoAs (Fig. 5b), indicating that the phenotypic information provided by the BFP marker was significant. Using additional information from the GFP and RFP/fusionRed markers improved overall predictive performance, but it is notable that we did not see any one individual reporter cell line dramatically outperform all others.

## Discussion

We developed a live-cell image-based profiling assay and used it to catalogue the morphological phenotypes induced by a set of >1,000 well-annotated reference compounds in 15 reporter cell lines in an effort to better understand how to efficiently design such image-based phenomic profiling screens that can simultaneously distinguish a broad range of compound MoAs. To our knowledge, this study is the most extensive experimental design exercise conducted in image-based phenomic profiling published thus far.

Our panel of cell lines expressed 12 different fluorescently-tagged proteins from their endogenous chromosomal loci, enabling us to monitor the morphology of different organelles and the activity of various signalling pathways in three distinct cell types. We observed that 57% of reference compounds were phenotypically active in ≥1 reporter line, which is comparable to previous image-based profiling screens [26]. There was a clear trend for compounds co-annotated to the same MoA to be either all active or all inactive in any given reporter line. Furthermore, we observed that compounds annotated to targets expressed in the assayed cell type were far more likely to be phenotypically active than compounds annotated to non-expressed targets. With the notable exception of GR agonists, the activity profiles of the different MoA groups were similar across all 15 reporter cell lines. This may be explained by the fact that there is a significant overlap between the gene sets expressed in A549, HepG2, and WPMY1 cell lines. Only 13% of all the ~12,000 expressed genes are unique to a single cell background, and nearly 70% of the expressed targets annotated to compounds in our chemical library are transcribed in all of our reporter cell types (Supplementary Fig. S4). These substantially overlapping expression patterns may explain why we recovered more than 90% of the phenotypically active compounds using just three reporter lines. The inclusion of additional cell types would only be justified if they significantly increased the coverage of ‘active biology’; their selection could be guided by differentiation status or constitutive or stimulus-induced expression patterns of targets. For example, according to the Human Protein Atlas [27], the THP-1 monocytic cell line expresses 1,114 genes not expressed by A549 and HepG2 cells. Notably, our initial attempts to increase the diversity of cell lineages by constructing monocyte and fibroblast reporter cell lines were frustrated by the difficulty of gene-editing these lineages; screening these cell types or primary cells using fixation and multiplexed dyes to measure cellular phenotypes, such as in the cell painting assay, may be the best way forward [28].

We were able to use the phenotypes induced by a broad range of pharmacologically distinct compounds to distinguish their MoA. Some MoAs, such as GR agonists and inhibitors of aurora kinase, 3-hydroxy-3-methyl-glutaryl-coenzyme A (HMG-CoA) reductase, kinesin, microtubules, and Rho-associated kinase, were very clearly distinguished (i.e., AUC-ROC ≥0.99), indicating that our imaging platform could be used directly to reliably annotate uncharacterized compounds for such MoAs. For many other MoAs we achieved slightly lower AUC-ROC values ≥0.9 and such models are useful for MoA hypothesis generation, limiting the number of target-specific orthogonal assays that need to be performed. Nevertheless, only 41 of the 83 testable MoAs could be distinguished accurately (ROC-AUC ≥0.9). This failure to distinguish certain MoAs may stem from several causes. For instance, for inhibitors of DNA synthesis, MEK, and heat shock protein, we observed highly MoA-specific phenotypes in all cell lines for subsets of these reference compounds, but the presence of outliers in these annotation groups resulted in relatively low performance estimates. For other MoAs such as ABL1 inhibitors and platelet-derived growth factor receptor (PDGFR) family inhibitors, we observed highly diverse phenotypes for the majority of compounds in these groups, indicating that the phenotypic activity in our reporter cell line panel is primarily driven by modulation of off-targets.

The lack of a consistent (or any) phenotypic activity may reflect the absence of key mechanistic targets in the imaging assay – implying a need for a different cell system – or differences in compound potency, response time, or poly-pharmacology. The impact of potency differences can be mitigated by the inclusion of multiple concentrations. Indeed, we observed higher performance gains when including additional compound concentrations compared to additional replicates of a single concentration. While not explored in this study, multiple time points could be added, and they may prove useful in unmasking effects. Nonetheless, the nearest-reference approach used in this study is inherently vulnerable to poly-pharmacology, because it assigns a query compound to the nearest reference(s) based on the same unweighted imaging features for all MoAs. Compounds can therefore be assigned to only one of multiple MoAs, salvaging some but not all information. For instance, several compounds annotated to different MoAs instead clustered with known mitochondrial toxicants. Hence, phenomic profiling combined with nearest-reference missed the known MoAs, but not without merit as it flagged certain compound liabilities. Other compounds, however, may end up drawn in limbo amid multiple references without any assignment at all. Recent large-scale MoA-annotation efforts [12,29] may enable growing the size of highly selective reference libraries to some extent, but neither do they address poly-pharmacology of uncharacterized compounds (such as hits from a screen for which one wants to make MoA predictions) nor do they overcome the relative lack of highly selective drugs or chemical probes for many MoAs.

The supervised deconvolution approach in Simm et al., 2018 [7], may be better suited to capture poly-pharmacology than the nearest-reference approach. For each assay (target/MoA) it models, it leverages a large volume of imaged compounds and their activity labels, which enables it to tune model-specific schemes to identify the most predictive imaging features for each assay. That supervised deconvolution approach yielded models that reliably (AUC-ROC ≥0.9) predict compound activity for 10 targets mapped to MoAs that were not phenotypically distinctive in any of the reporter cell lines in our study. Simm et al. imaged a H4 neuroglioma cell line, which may have contributed to the increased coverage of MoAs. However, we could compare our results to an unpublished in-house study similar in design and scale to Simm et al. that was based on a HepG2 cell line. That large-scale study yielded predictive models for five targets mapped to MoAs that were not phenotypically distinctive in any of the five HepG2 cell lines in our study. Even though the HepG2 assays in both studies were not identical, these findings suggest that augmenting phenomic profiling at scale with assay activity data and supervised machine learning can expand the set of predictable targets/MoAs. Amenable datasets already exist in the industry, and the creation of public counterparts is underway.

We observed that the genetic background of the reporter cell line did not dramatically affect the overall AUC-ROC values calculated for the different compound MoAs, which is consistent with the results of large-scale transcriptomic profiling experiments where the prediction of multiple MoAs was similarly effective in several cell types [30]. However, there were a few clear exceptions, like the GR agonists and mitochondrial toxicants that induced more predictive phenotypes in A549 cells than in HepG2 and WPMY1 cells. We also observed that the number of distinguishable MoAs benefited similarly from the inclusion of additional markers as from inclusion of additional cell types, albeit only modestly. Comparison of AUC-ROC values demonstrated that, overall, no particular combination of GFP and RFP/fusionRed markers outperformed all others. One reason for this observation may be the fact that all reporter cell lines expressed BFP nuclear and cytoplasmic segmentation markers, and in many cases these on their own were sufficient to distinguish MoAs with reasonable accuracy. The predictive power of nuclear and cytoplasmic markers has recently been noted in both live- and fixed-cell phenotypic screens using small sets of annotated reference compounds [4,31] and our results extend this finding.

Overall, our findings suggest a strategy to improve target/MoA predictive performance of image-based phenomic profiling. Firstly, the addition of complementary cellular models and stimuli by a transcriptomics-driven selection could improve target coverage, possibly also aiming to specifically increase the coverage of fitness genes. Secondly, a data-efficient nearest-reference approach like that employed in the current study could be used to evaluate the target/MoA predictive performance of a large panel of cellular models and treatment conditions, also including additional concentrations and time points. Additional markers could also be evaluated, however based on our results it seems unlikely to have a dramatic impact. Finally, for a panel of top-performing imaging assays, images could be generated for large numbers of compounds combined with activity labels from orthogonal assays and supervised deconvolution approaches to address poly-pharmacology. While this strategy calls for significant investments to be made, we believe it will advance the field of phenomic profiling and contribute to the discovery of better therapeutics.

## Methods

### Reagents

The following reagents were used: fetal bovine serum (FBS; BioWest, Nuaillé, France; S1810); minimal essential medium (MEM; Gibco, Thermo Fisher Scientific, Waltham, MA; 21090); MEM, no phenol red (Gibco 51200); Dulbecco’s modified Eagle’s medium (DMEM; Gibco 11960); FluoroBrite™ DMEM (Gibco A18967); RPMI-1640 (Sigma, St. Louis, MO; R0883); RPMI-1640, no phenol red (Sigma R7509); 200 mM L-glutamine (Sigma G7513); 10,000 U/ml penicillin-streptomycin (Pen-Strep) (Gibco 15140); 100 mM sodium pyruvate (Sigma S8636); 384-well poly-D-lysine-coated microClear plates (Greiner, Kremsmünster, Austria; 781946 custom-barcoded).

### Cell Lines

A549, HepG2, and WPMY1 cells were custom gene-edited by MilliporeSigma Cell Design Studio™. Five different variants of each cell type were constructed. Each variant contained a segmentation cassette that expressed both a TagBFP protein (Evrogen, Moscow, Russia) and a nuclear-localized TagBFP protein, used to identify the cytoplasmic and nuclear regions of cells during image analysis, respectively. Alternative codons were used for the two TagBFP sequences in the segmentation cassette to prevent loss of nuclear-localized BFP signal due to spontaneous homologous recombination. The segmentation cassette also contained two translational (2A) skip sites, one located between the BFP sequences and the other at the 3’-end of the cassette. The segmentation cassette was integrated into the genome by homologous recombination at the first protein-coding exon of either beta-actin (ACTB) or vimentin (VIM), placing the expression of the segmentation markers under the control of the ACTB/VIM promoter without disrupting the expression of the host gene.

Following integration of the segmentation cassette, each cell line was further gene-edited to express one GFP-tagged organelle or pathway marker protein from its endogenous promoter and an additional RFP- or fusionRed-tagged organelle/pathway marker from its chromosomal locus. Five different variants of each cell type were created by introducing different pairs of green and red markers: (i) GFP-ACTB (actin cytoskeleton) and RFP-RAB5A (early endosome), (ii) CANX-GFP (endoplasmic reticulum) and COX4I1-RFP (mitochondria), (iii) GFP-GM130 (Golgi) and fusionRed-SQSTM1 (autophagosome), (iv) GFP-TUBA1B (microtubules) and fusionRed-RELA (NF-κB signalling), and (v) GFP-TP53BP1 (non-homologous end joining-mediated repair of DNA double strand breaks) and CLTA-fusionRed (clathrin-mediated endocytosis). The fluorescent protein tags used were TagGFP2, TagRFP, and fusionRed (Evrogen).

### Cell Culture

Cells were maintained using standard laboratory techniques at 37°C and 5% CO_2_. During cell line maintenance and propagation, cells were cultured in the presence of phenol red; but, this was omitted during imaging as it obscures the signal from RFP/fusionRed-tagged proteins. HepG2 cells were cultured in MEM, 10% FBS, 2 mM L-glutamine, 100 U/ml Pen-Strep. A549 cells were cultured in RPMI-1640, 10% FBS, 2 mM L-glutamine, 100 U/ml Pen-Strep. WPMY1 cells were cultured in DMEM, 5% FBS, 4 mM L-glutamine, 1 mM sodium pyruvate, 100 U/ml Pen-Strep.

### Microscopy Screen

Cells were seeded in medium containing no phenol red on to 384-well poly-D-lysine-coated microClear plates at the appropriate density and grown for 20 to 40 hours before treatment with compound at 0.3, 1, 3, or 9 μM. To control for experimental variation, two technical replicates were included for each compound treatment in each imaging batch and two batches were prepared for each reporter cell line. Cells were grown for an additional 24 hours and imaged on a Cell Voyager 7000 (Yokogawa, Tokyo, Japan) high-throughput microscope equipped with a climate control chamber (37°C, 5% CO_2_). Images of BFP (ex: 405 nm, em: 445/45), GFP (ex: 488 nm, em: 525/50), and RFP/fusionRed (ex:561 nm, em: 593/46) were acquired at two to three sites per well using a 20× objective lens (Olympus, Tokyo, Japan; NA 0.75). All liquid dispensation, compound treatment, and imaging steps were conducted on an automated platform designed specifically for live-cell microscopy screens. Typically, twenty 384-well plates were imaged per day over a 20-hour period. As cells were actively growing during this time (doubling time 18-24 hours), the last 10 plates in a batch were seeded at a lower cell density than the first 10 plates to compensate for the additional growth period they would encounter before imaging. The exact seeding densities were optimized for each cell line so that cells in control wells were 50% to 80% confluent at the time of imaging.

### Exclusion of Fluorescent Compounds

Fluorescent compounds may mask the signal from one or more of the BFP/GFP/RFP/fusionRed-tagged proteins. We identified such compounds by screening them in a HepG2 cell line not expressing any fluorescently-tagged proteins and examining the resulting images by eye. The 117 of 1,294 initially screened compounds found to be fluorescent were excluded from our study, leaving 1,008 reference compounds and 169 natural products which we screened against the 15 reporter cell lines.

### Compound Annotation

Each of our 1,008 unique reference compounds was manually annotated with a MoA descriptor (e.g., “LRRK2 inhibitor”, “DNA alkylating agent”, “Serotonin receptor antagonist”, etc.), containing molecular target names rather than assigning compounds to cellular pathways or phenotypic responses as the latter descriptions of compound activity lack specificity and can manifest differently in different cell types. Compounds that acted on multiple functionally distinct targets, notably kinase inhibitors, were assigned multiple MoA descriptors. Guidance was provided by two recent compound annotation efforts that sought to manually annotate FDA-approved drugs and some preclinical compounds [12,29]; 280 compounds not annotated in either of these sources were manually annotated based on vendor catalogue information and the curated target annotations in the ChEMBL and IUPHAR chemical databases [16,17].

In addition, the gene names of individual targets were extracted from the manually curated target information in the ChEMBL and IUPHAR databases for each of the 1,177 unique chemicals (1,008 reference compounds and 169 natural products) by searching for matching chemical structures. In total, the gene names of 692 targets were mapped to 690 compounds in our chemical library.

### Microarray Analysis

A549, HepG2, and WPMY1 parental cell lines were seeded in 6-well plates and grown for 24 hours to a confluency of 70% to 90% before RNA was extracted using an RNeasy RNA extraction kit (Qiagen, Venlo, the Netherlands). Samples were prepared in triplicate, processed, and hybridized to HGU-219 arrays (Affymetrix, Santa Clara, CA) as previously described [32].

The data were pre-processed with the Robust Multi-array Average (RMA) methodology [33], summarized at the gene level (ENTREZID). A gene was considered expressed in a cell line if its log_2_-normalized intensity was higher than a threshold for all three replicates and all probe sets of the gene (gene symbol). The threshold for each replicate was computed as the 95th percentile of the first mode of a Gaussian mixture with two components fitted to the sample probe set distribution of that replicate. The nine obtained thresholds (three replicates × three cell lines) were in the range between 4.187 and 4.265 and resulted in 58% genes considered expressed in A549 cells, 57% in HepG2 cells, and 57% in WPMY1 cells.

### Cell Identification and Feature Extraction

PerkinElmer (Waltham, MA) Acapella/Columbus image analysis software was used to identify the boundaries of individual nuclei and surrounding cytoplasm based on the BFP channel. Approximately 500 features quantifying intensity, texture, shape, and spatial relationships in all three channels were calculated for each cell. The extracted features together with compressed versions of the original images and masks of the nuclei and cell outlines were imported into Phaedra [34] for visual quality control.

### Removal of Dead and Mis-segmented Cells

We trained linear support vector machine classifiers separately for each cell type to recognize live, dead, and mis-segmented cells based on manually labelled training sets comprising ~12,000 cells per cell type sampled from all batches. Five-fold cross-validation accuracy was 98.3% for A549 cells, 98.3% for HepG2 cells, and 96.7% for WMPY1. This resulted in a total of 217.3 million cells classified as live, 11.9 million cells as dead, and 12.6 million cells as mis-segmented. This clean-up step resulted on average in 11.6 (3.0%) more compounds per cell line considered active with a reproducible phenotype compared to not performing this clean-up step.

### Well-level Feature Aggregation

The mean and the median for each feature as well as cell count was calculated for all live cells in a well, resulting in a ~1,000-dimensional descriptor for each well.

### Normalization

Well-level features were normalized on a plate-by-plate and batch-by-batch basis. Each full plate contained 28 control wells (dimethyl sulfoxide [DMSO]-treated) and 280 sample wells (compound-treated). Each feature was normalized by subtracting the mean of the control wells plate-wise, followed by dividing by the standard deviation of the control wells batch-wise.

### Feature Selection

Some imaging features are highly correlated to each other and/or show poor reproducibility across replicate wells. To identify a set of representative features in an unsupervised fashion (in the sense that it does not use MoA annotations) we used a modified version of the mRMR algorithm [18] with the treatment IDs as classes. We reasoned that a feature set that is able to discriminate different treatments from each other would also be useful to discriminate different MoAs.

First, samples (i.e., wells) highly similar to DMSO controls were removed based on the Euclidean distance between the sample feature profile and the mean DMSO control profile using a 95th percentile cut-off on the null-distribution of Euclidean distances between individual DMSO control profiles and the mean DMSO control profile. Second, wells with <100 cells were removed and 1% of each feature vector of the remaining wells was trimmed at either end of the distribution to reduce the influence of outliers.

The relevance of each feature with respect to the classification variable (i.e., treatment ID) was computed as an F-statistic, and scaled *versus* the maximum obtained relevance across features. An iterative procedure, starting with the most relevant feature, selected the next feature with the best trade-off between relevance and redundancy. The redundancy of each not yet selected feature was updated in each iteration and was defined as the maximum correlation between the feature and any of the already selected features. The feature with the maximum trade-off score between relevance and redundancy, computed as relevance – (relevance × redundancy), was selected in each iteration. This procedure resulted in a ranked list of features.

To select an optimal number of features, for feature sets of increasing size the ability to discriminate pairs of replicates from pairs of non-replicates based on the Pearson correlation was measured as area under the curve (AUC). A maximum of 10,000 pairs of replicates and non-replicates were randomly selected, and the process was repeated 10 times to derive mean and standard deviation of the AUC for each number of features tested. The selected number of optimal features is the minimal number of features leading to a mean AUC within the maximum mean AUC minus one standard deviation.

The feature selection was done separately for each cell line. For the different cell lines, we obtained between 22 and 58 features. For each well, the selected feature set summarized the phenotypic state of the cell population, referred to as imaging signature.

### Active Calling

We classified each treatment (i.e., compound at a specific concentration) in each reporter cell line as phenotypically active or not (Supplementary Table S2) using two criteria: (i) (strictly) >50% of the replicates show a significant deviation in their imaging signature compared to that of DMSO control, and (ii) the replicates show highly reproducible imaging signatures.

For the first criterion, we flagged each well as active or inactive by computing the Euclidean distance between the imaging signature of that well and the mean imaging signature of all control wells (which is a 0-vector due to the z-score normalization against the controls) and comparing that distance against a threshold tau. Tau was set to the 95th percentile of the null-distribution of the Euclidean distances between the imaging signature of each individual control well and the mean imaging signature of all control wells. Thus, by definition, we considered 5% of all individual control wells as (Euclidean-)active; and we considered a treatment as (Euclidean-)active if ≥3 of four of its replicates were flagged as (Euclidean-)active.

For the second criterion, we computed the median Pearson correlation between all six pairs of replicate imaging signatures of each treatment. We constructed a null-distribution of non-replicate correlations using all wells of (Euclidean-)active treatments (between 7 and 13 million pairs, depending on the cell line), and used the 95th percentile of this null distribution as a threshold. Here, non-replicates were defined as pairs of wells with different compounds (i.e., excluding pairs of the same compound but different concentration). We considered a treatment as reproducible if its median replicate correlation was higher than that threshold.

t-SNE projections and AUC-ROC analyses were done only for the phenotypically active treatments. A compound was considered phenotypically active if ≥1 of its concentrations was classified as phenotypically active. Our data were highly reproducible as imaging signatures for active compound treatment replicates showed similarly high correlation within and across batches (Supplementary Fig. S13a and S13c).

### Treatment-level Aggregation

To obtain a single imaging signature per treatment, we calculated the median imaging signature of all four replicate wells.

### AUC-ROC Analysis

For each MoA and cell line with ≥3 active compounds, we computed the area under the receiver-operating characteristic curve (AUC-ROC), quantifying how well compounds with that MoA can be distinguished from compounds with a different MoA, following a similar approach as in Loo et al., 2007 [13] and Subramanian et al., 2017 [5] (Supplementary Fig. S7). We used each active treatment once as a query and ranked all other active treatments (except other concentrations of the query compound) based on the Pearson correlation between their imaging signatures. To avoid heavily penalizing a compound that shows high phenotypic similarity to other compounds with the same MoA at only a subset of its concentrations, we removed all but the highest ranked concentration of each compound. The correlation similarity to the query treatment together with a binary flag for each ranked treatment whether or not it has the currently considered MoA of the query treatment (compounds may be annotated with multiple MoAs) enables computing the AUC-ROC. This results in one AUC-ROC value for each active treatment and MoA it is annotated with. We then aggregate these AUC-ROC values on the MoA level by first taking for each compound the maximum AUC-ROC value over its concentrations, followed by taking the median of the per-compound AUC-ROC values within each MoA. We note that the choice of maximum and median leads to an ‘optimistic’ AUC-ROC in the sense that if all compounds of a MoA show high similarity to each other at ≥1 concentration the resulting AUC-ROC value will be high. However, if one or more compounds are phenotypically different at all concentrations from other compounds of that MoA, this will result in a lower AUC-ROC value for each individual query treatment of that MoA and hence also a lower maximum/median aggregated AUC-ROC value for that MoA. The entire AUC-ROC analysis was done separately for each cell line. If a compound belongs to multiple MoAs, it was considered independently in the AUC-ROC computation of each of its MoAs.

To assess the statistical significance of the AUC-ROC values, we computed empirical p-values by permuting the MoA annotations and deriving a null distribution of AUC-ROC values in 10,000 iterations. We adjusted p-values to account for multiple testing by controlling the false-discovery rate (FDR) at alpha = 0.05, using the Benjamini-Hochberg method [35]. Statistical significance was achieved at AUC-ROC values above 0.8137 to 0.8577, depending on the cell line. AUC-ROC values and associated FDR-adjusted p-values for all 83 testable MoAs are listed in Supplementary Table S3.

Our data were highly reproducible as the AUC-ROC values computed for the different MoAs did not exhibit sizeable batch effects (Supplementary Fig. S13b and S13c).

### Chemical Similarity

We calculated chemical similarity between all pairs of compounds using atom pair fingerprints [36] and Tanimoto distance between them, using the sdf2ap() and desc2fp() functions in the ChemmineR R-package.

### Core-fitness Genes

We combined the CRISPR results from Hart et al., 2015 [20] and Wang et al., 2015 [21] for 16,993 genes that were queried for fitness defects in nine cell lines. Using a similar criterion as in Hart et al. we defined core-fitness genes as those that caused fitness defects in at least two thirds of the tested cell lines (i.e., six or more out of nine cell lines). In total, 1,228 core-fitness genes were identified using these datasets.

## Supporting information

All supplementary figures except S6 and S8

## Acknowledgements

We would like to thank Luc Geeraert for excellent medical writing assistance as well as Victoria Eastham and Reinoud De Groot for critical reading and helpful feedback.

## Author Contributions

Y.T.C., E.G., and S.J. conceived the research. Y.T.C. and H.W.H.G. supervised the research. D.M. oversaw and participated in the creation of the reporter cell lines. M.J.C. and J.V.d.W. developed and conducted the imaging screen assay. S.J. performed the image analysis and in collaboration with L.C. and D.S. analysed all data from the imaging screen. P.J.P. contributed to the conception of the study and the manuscript. H.C. contributed analysis concepts and result interpretation. M.J.C., J.-M.N., and J.K.W. annotated the compound set used in this study. S.B., I.V.D.W., and S.J.K. conducted the glucocorticoid receptor agonist microarray experiments. L.C. analysed all microarray data. M.B. and M.S. performed the Glu/Gal assay. S.J., M.J.C., H.C., E.G., and H.W.H.G wrote and edited the manuscript.

## Competing Interests

S.J., S.J.K., I.V.D.W., J.-M.N., M.B., M.S., P.J.P., J.K.W., H.C., E.G., and H.W.H.G. are employees of Janssen Pharmaceutica N.V. The other authors declare no competing interests.

## Availability of Materials and Data

The 15 reporter cell lines are available for purchase upon request from MilliporeSigma. A data set comprising all microscopy images together with the public chemical structures of the studied compounds will be made publicly available upon publication of the manuscript.

