## Supplementary material for "Tales of 1,008 Small Molecules: Phenomic Profiling through Live-cell Imaging in a Panel of Reporter Cell Lines": All supplementary figures except S6 and S8

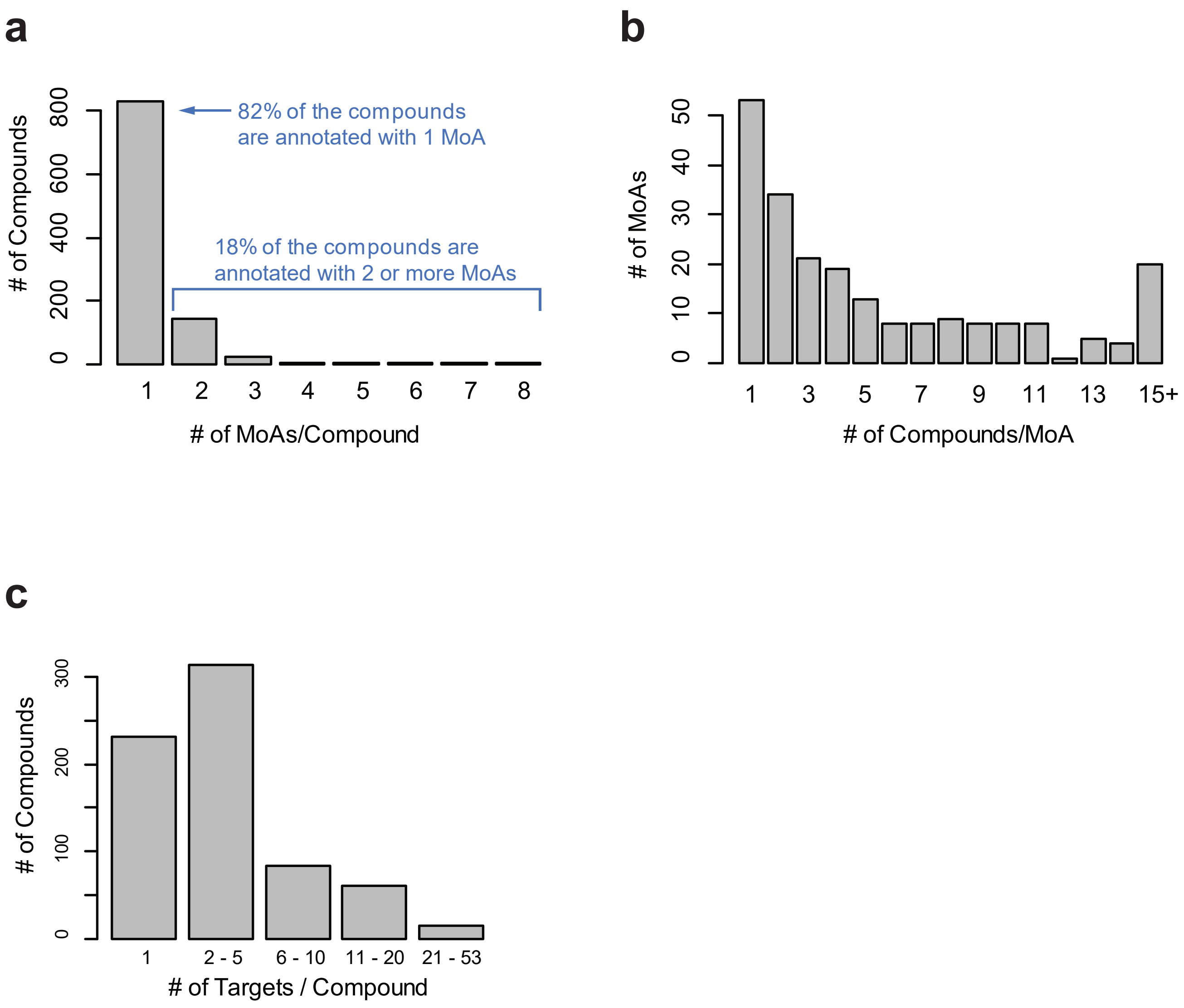


**Supplementary Figure S1**

Summary of MoA and target annotations for our chemical compound library.

(a) Number of MoAs per compound.

(b) Number of compounds per MoA.

(c) Number of targets per compound (MoA-annotated reference compounds and natural products).

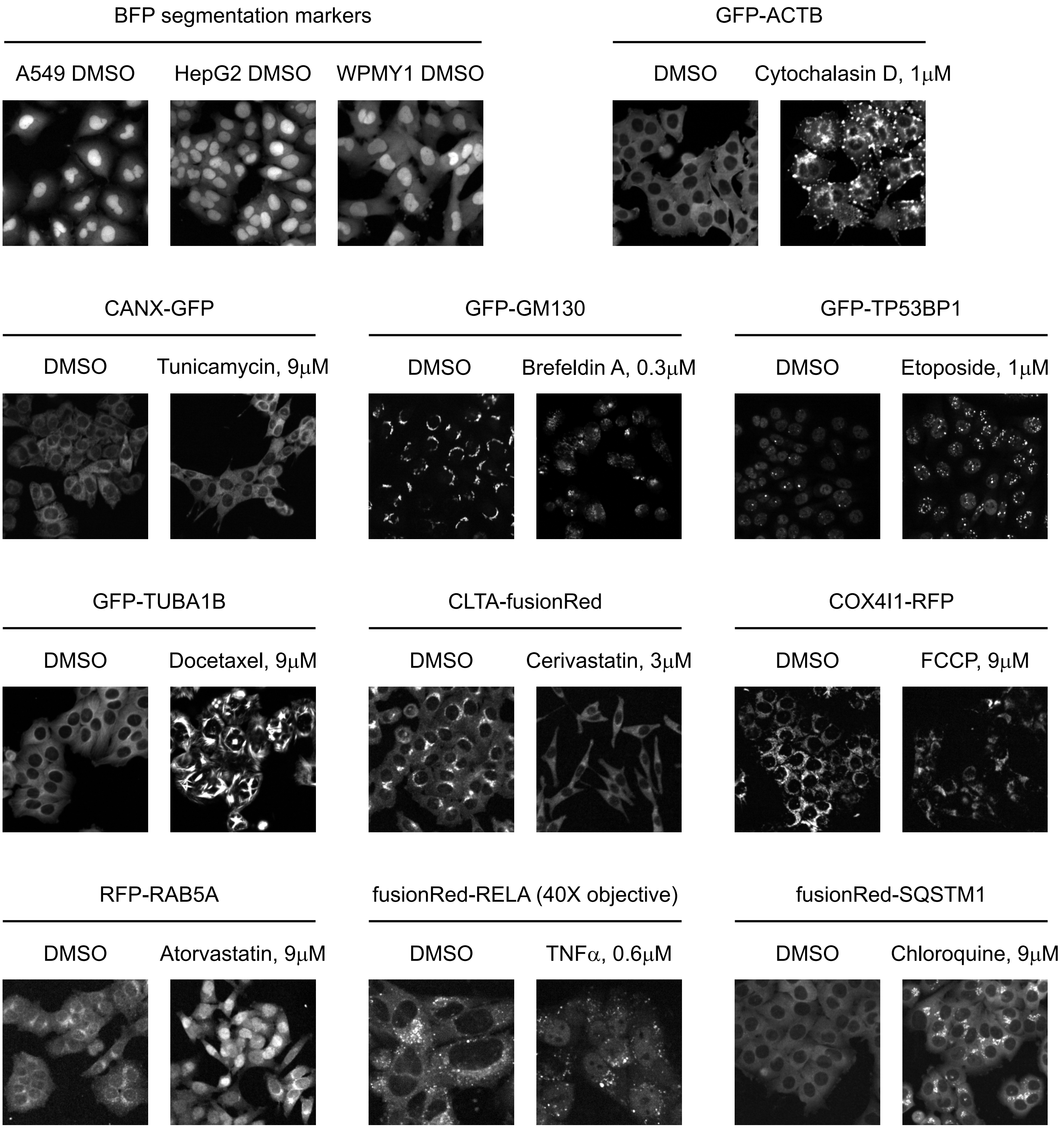


**Supplementary Figure S2**

Marker validation - microscopy images of the fluorescent markers expressed in the different reporter cell lines used in this study are shown.

The BFP nuclear and cytoplasmic segmentation markers are pictured in control (DMSO-treated) A549, HepG2, and WPMY1 cells. To ensure that the fluorescent GFP/fusionRed/RFP tags did not impact the expected subcellular localization or function of the organelle and pathway markers, cells were treated with either DMSO or a tool compound known to perturb the localization or expression of the marker being visualized. FusionRed-RELA images are from WPMY1 cells imaged at 40× magnification. All other GFP/fusionRed/RFP images were taken at 20× magnification in the HepG2 background.

a

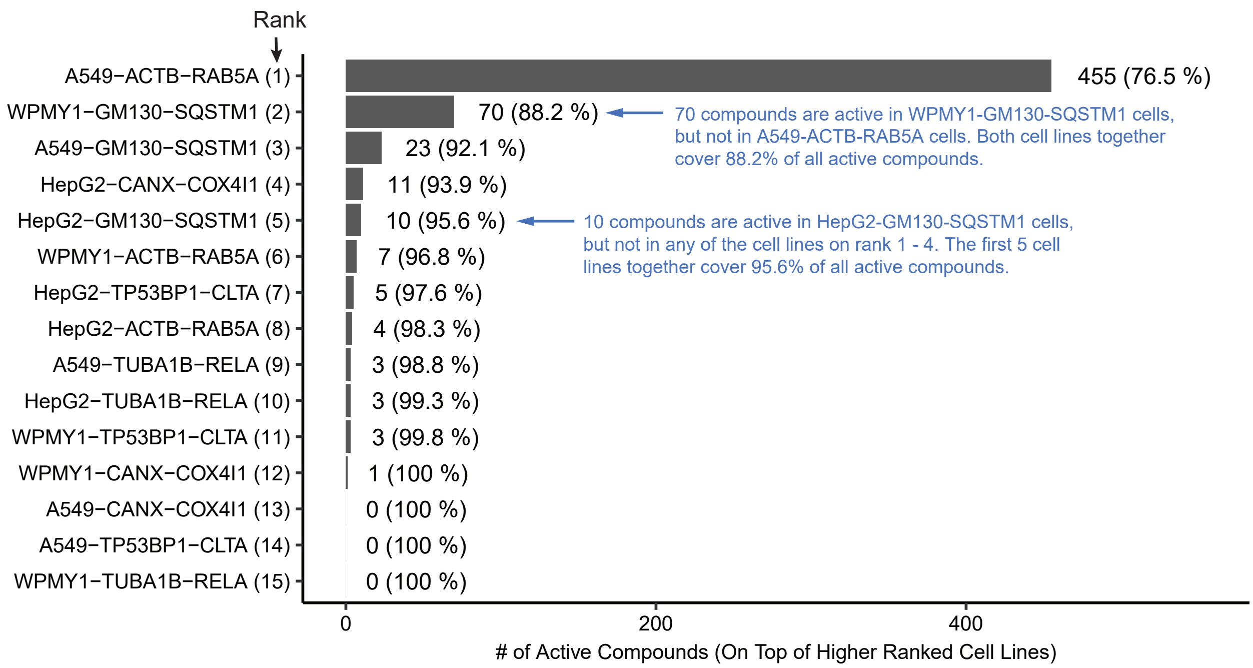


b

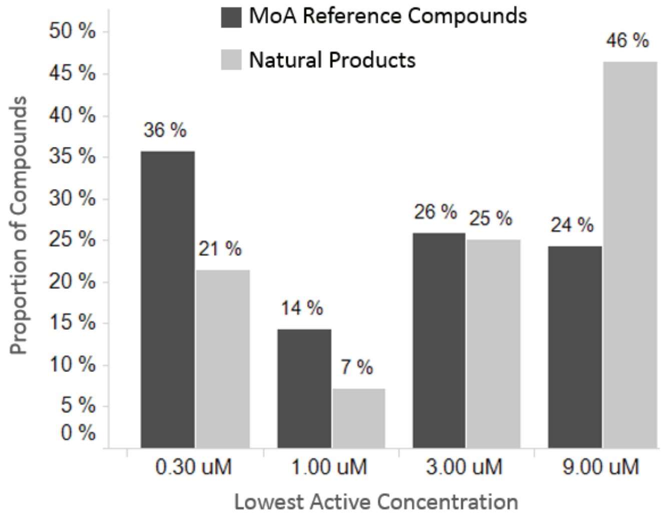


**Supplementary Figure S3**

(a) 92.1% of the active compounds were captured with just three cell lines. Cell lines were ranked by number of active compounds, starting with the A549-ACTB-RAB5A cell line which had the largest number of active compounds and then iteratively adding the cell line with the largest number of active compounds that were not active in any of the higher ranked cell lines.

(b) Natural products showed a greater tendency to be phenotypically active only at the highest tested concentration compared to the MoA reference compounds. Bar graph shows the lowest active concentration (minimum over cell lines) of compounds that were phenotypically active in ≥1 cell line.

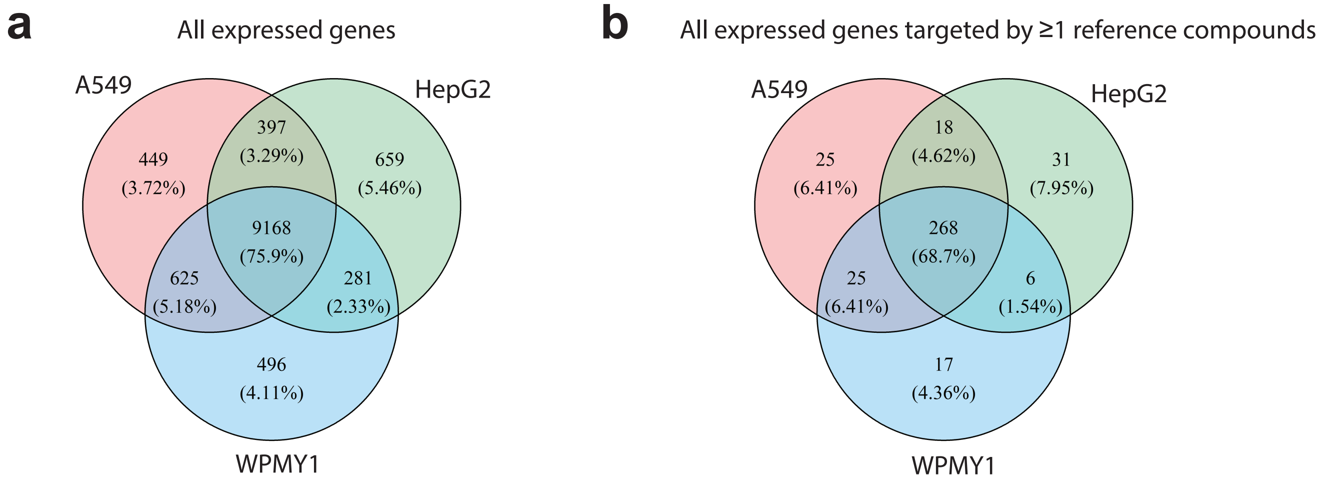


**Supplementary Figure S4**

Gene and target expression patterns in A549, HepG2, and WPMY1 cell lines.

(a) Venn diagram showing overlap of expressed genes in the three parental cell lines, based on microarray data.

(b) Same as (a) but limited to the 390 expressed genes that were targeted by ≥1 of the compounds tested (including natural products). An additional 302 gene products annotated to compounds in our chemical library were not expressed in any of the three parental cell lines.

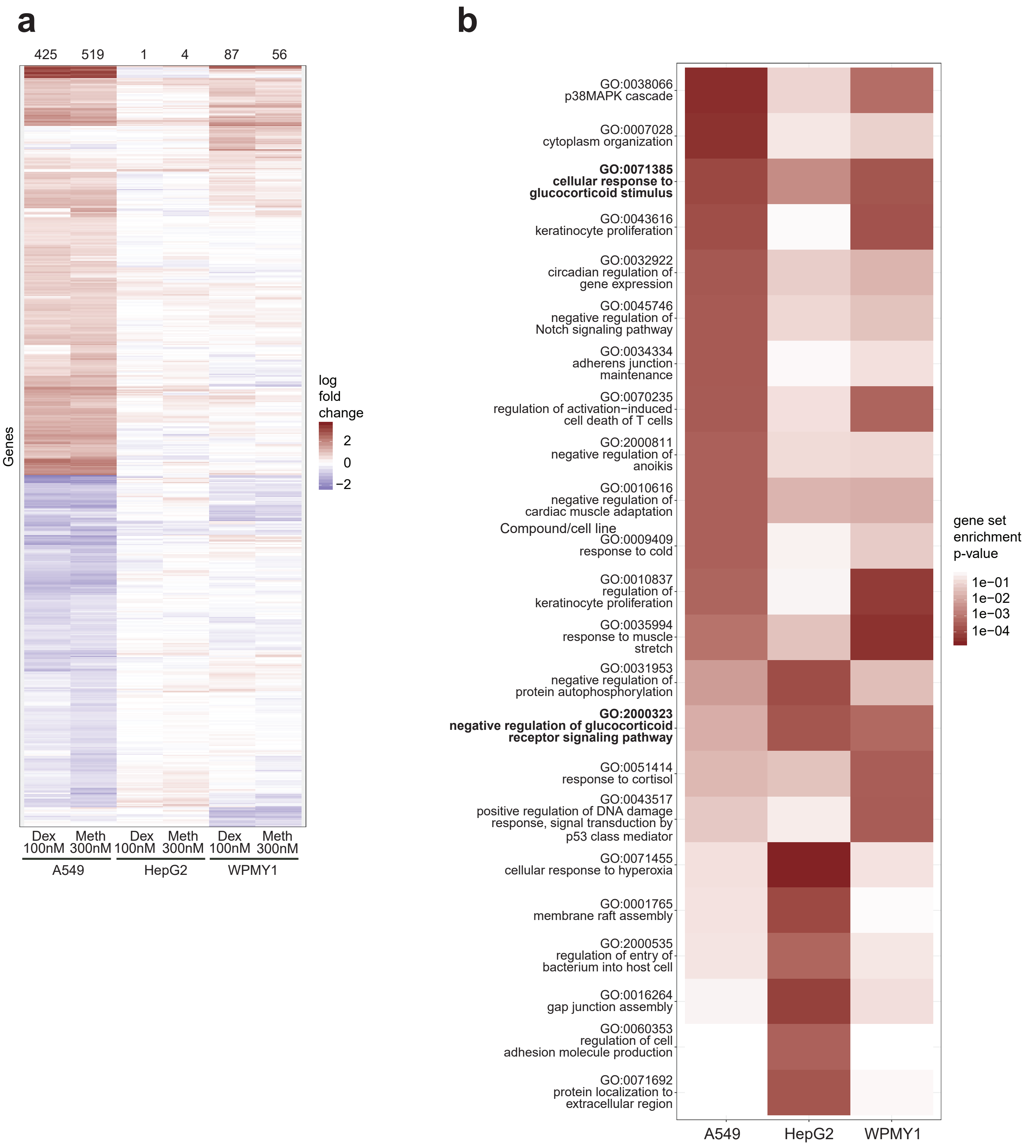


**Supplementary Figure S5**

GR agonists elicited a more robust transcriptional response in A549 cells than in HepG2 or WPMY1 cells.

(a) Hierarchically clustered heat map showing all genes that were significantly up- or down-regulated (adjusted p-value of treatment versus DMSO <0.05) in ≥1 cell line after treatment with the GR agonists dexamethasone (Dex) or methylprednisolone (Meth) for 4 hours. Numbers at the top of the heat map indicate how many genes were significantly up- or down-regulated.

(b) Heat map showing all pathways (Gene Ontology biological processes) with gene set enrichment p-value $<5\times10^{-4}$ in ≥1 cell line upon treatment with methylprednisolone at the lowest concentration (300 nM) used in the live-cell imaging screen. Pathways (rows) are sorted by increasing p-value in A549 cells.

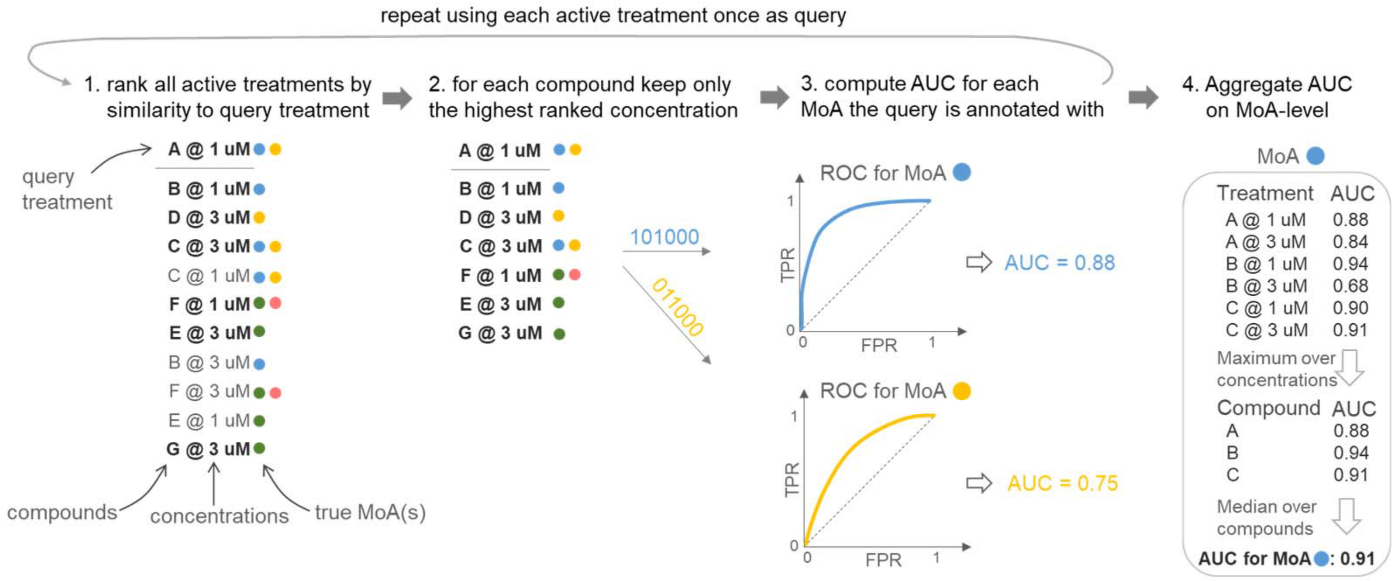


**Supplementary Figure S7**

AUC-ROC computation procedure to quantify how well each MoA can be distinguished from other MoAs, illustrated with seven compounds (named A - G) at two concentrations and annotated for four different MoAs (coloured circles). Here, compounds D and G are not active at 1 μM and therefore only occur at 3 μM in the ranking illustrated in Step 1. The sequences of 0's and 1's for the blue and the orange MoA correspond to the ranked compounds as illustrated in Step 2; they indicate whether or not the respective compound share the respective MoA with the query and define a ROC curve for each MoA. Abbreviations: FPR = false positive rate, TPR = true positive rate

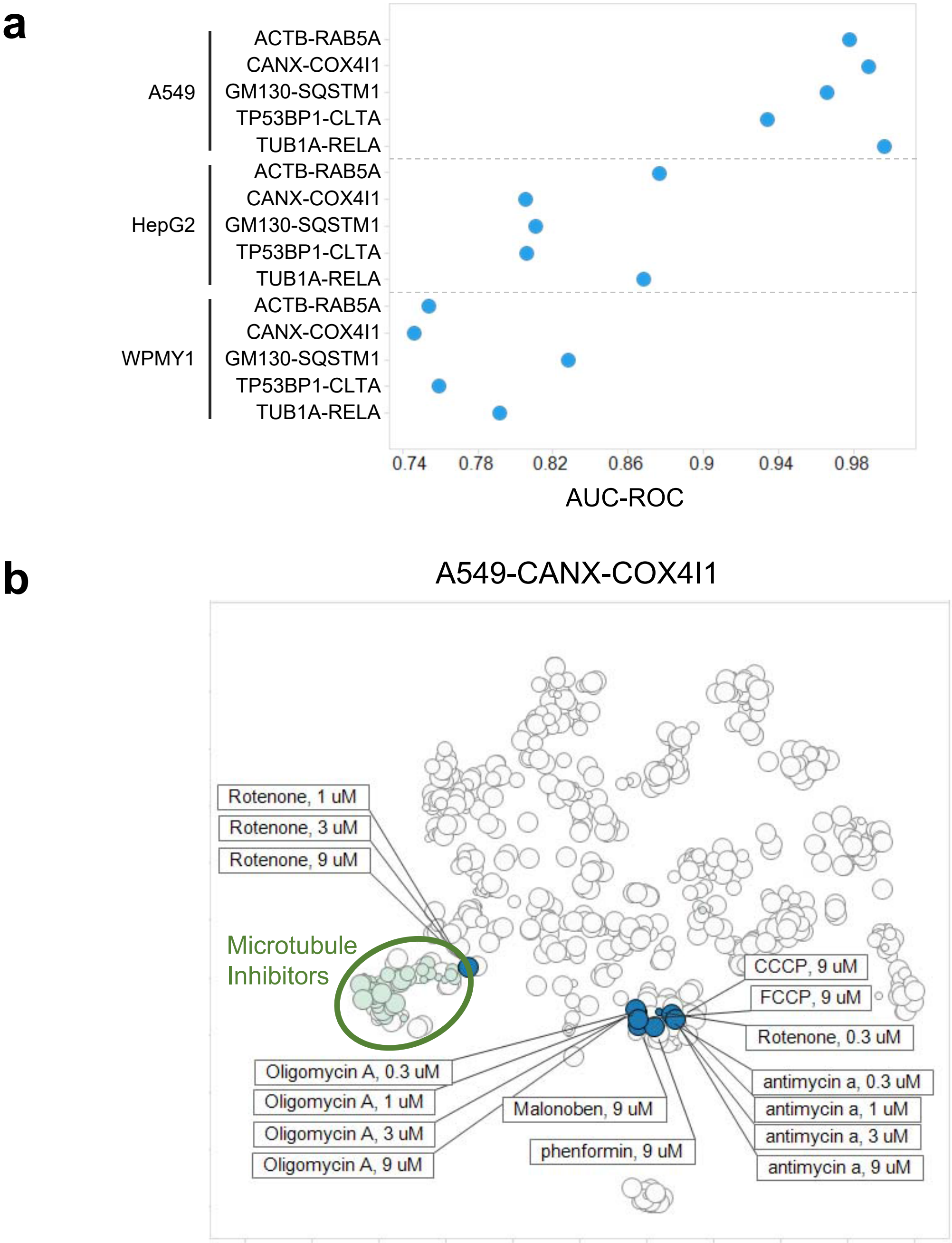


**Supplementary Figure S9**

A549 cells exhibited more functionally informative responses to mitochondrial toxicants (oligomycin A, CCCP, FCCP, rotenone, malonoben, phenformin, and antimycin A) than HepG2 and WPMY1 cells.

(a) AUC-ROC indicating how well the mitochondrial toxicants oligomycin A, CCCP, FCCP, rotenone, malonoben, phenformin, and antimycin A could be distinguished from all other MoAs in each cell line.

(b) t-SNE map showing all active treatments in the A549-CANX-COX4I1 cell line with the active concentrations of the seven mitochondrial toxicants highlighted. Rotenone clusters at high concentrations with microtubule inhibitors and this observation is consistent with a reported off-target effect of Rotenone on microtubules (Heinz, S. et al. Sci. Rep. 7, 45465 [2017]). Marker size indicates compound concentration; each marker represents a distinct treatment (median of four replicates).

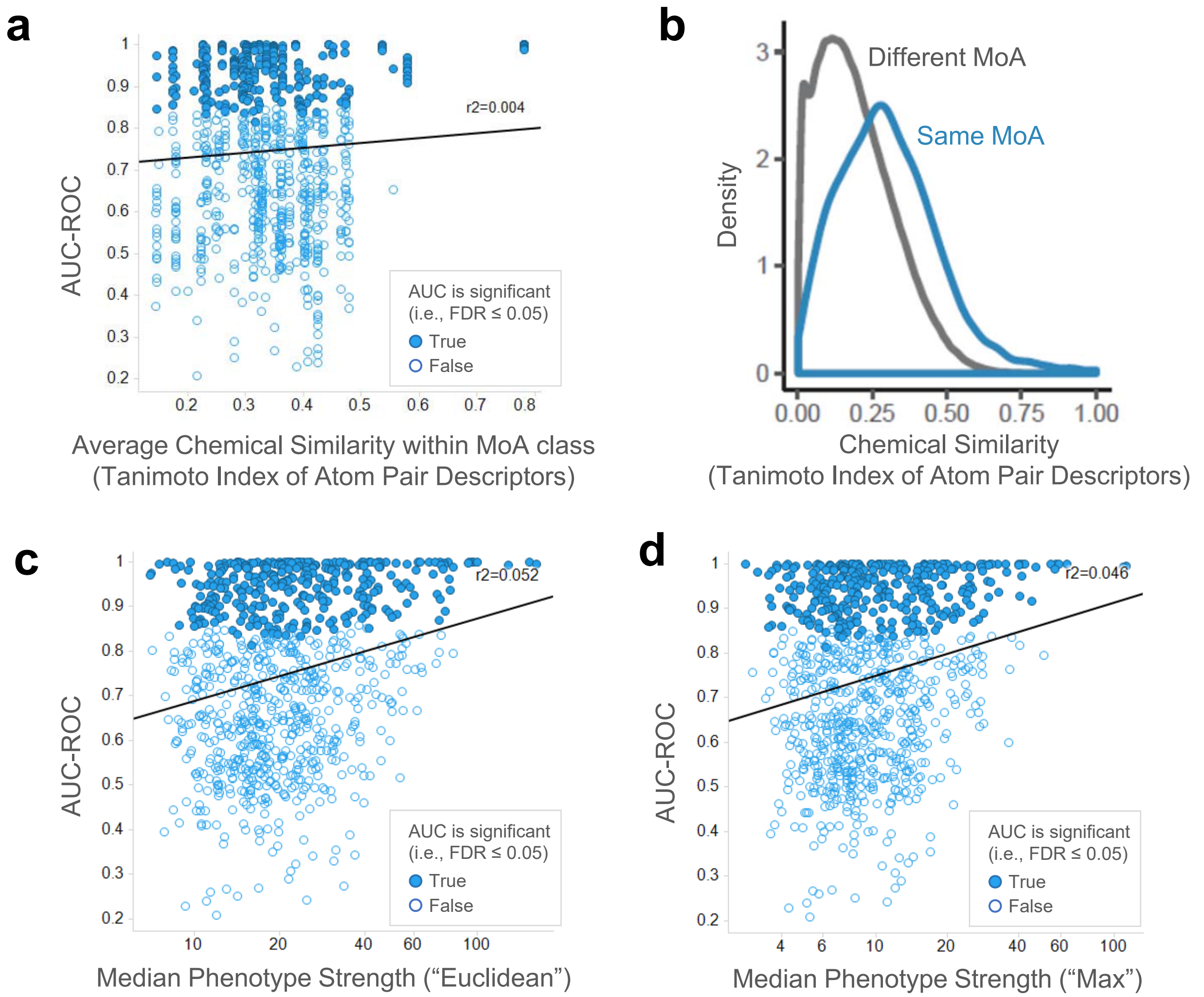


**Supplementary Figure S10**

How well MoAs can be distinguished was not explained by the chemical similarity of co-annotated compounds nor by the strength of the phenotypes they induced.

(a-b) MoA distinction performance did not correlate with chemical similarity. Each point in the scatter plot corresponds to a MoA/cell line pair. For each MoA the average chemical similarity of all compounds with that MoA active in ≥1 cell line is shown on the horizontal axis. The vertical axis shows how well the MoA could be distinguished from all other MoAs. The straight line is a least square fit of the data. (b) Density plot of chemical similarities of all tested compounds (independent of phenotypic activity) showing that most compounds with shared MoA had low chemical similarity.

(c-d) MoA distinction performance did not depend on the strength of the phenotype. (c) Each point in the scatter plot corresponds to a MoA/cell line pair. For each treatment the strength of the phenotype it induced was computed as the Euclidean distance of the treatment's signature to the mean DMSO control signature (which is a zero vector), and summarized on MoA level by taking the median phenotype strength over the same set of treatments with that MoA as considered in the AUC computation. Horizontal axis is shown on log10-scale. The straight line is a least square fit of the data. (d) Same as (c) but phenotype strength was computed as the maximum absolute feature value (i.e., z-score relative to DMSO control) in the compound's signature.

a

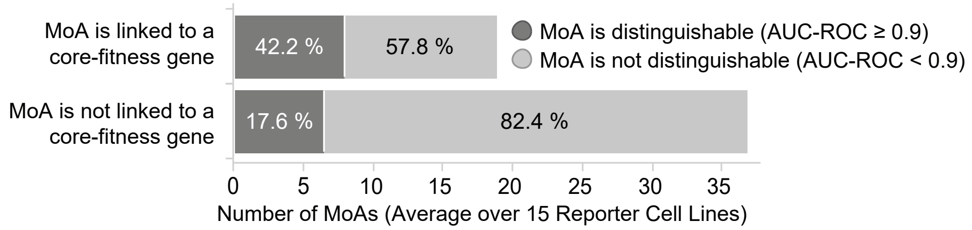


b

| Cell Line | p-value Fisher's Exact Test | Odds Ratio | # of MoAs "not distinguishable & not linked to core-fitness gene" | # of MoAs "not distinguishable & linked to core-fitness gene" | # of MoAs "distinguishable & not linked to core-fitness gene" | # of MoAs "distinguishable & linked to core-fitness gene" |
|---|---|---|---|---|---|---|
| A549-TP53BP1-CLTA | **0.00771** | 6.25 | 29 | 10 | 4 | 9 |
| HepG2-TUBA1B-RELA | **0.00826** | 5.72 | 33 | 10 | 5 | 9 |
| HepG2-CANX-COX4I1 | **0.01301** | 5.95 | 31 | 10 | 4 | 8 |
| HepG2-GM130-SQSTM1 | **0.01590** | 5.43 | 31 | 11 | 4 | 8 |
| HepG2-ACTB-RAB5A | **0.04003** | 4.08 | 33 | 11 | 5 | 7 |
| A549-CANX-COX4I1 | **0.04098** | 4.43 | 23 | 10 | 4 | 8 |
| WPMY1-TUBA1B-RELA | 0.05138 | 4.03 | 25 | 9 | 6 | 9 |
| WPMY1-GM130-SQSTM1 | 0.06032 | 3.26 | 35 | 12 | 7 | 8 |
| HepG2-TP53BP1-CLTA | 0.10819 | 3.15 | 31 | 11 | 7 | 8 |
| A549-GM130-SQSTM1 | 0.11666 | 2.70 | 33 | 12 | 7 | 7 |
| A549-ACTB-RAB5A | 0.12044 | 2.78 | 35 | 11 | 9 | 8 |
| WPMY1-CANX-COX4I1 | 0.12312 | 2.68 | 30 | 11 | 8 | 8 |
| A549-TUBA1B-RELA | 0.12624 | 2.62 | 30 | 10 | 9 | 8 |
| WPMY1-ACTB-RAB5A | 0.35559 | 1.86 | 29 | 12 | 9 | 7 |
| WPMY1-TP53BP1-CLTA | 0.54879 | 1.48 | 25 | 13 | 9 | 7 |

**Supplementary Figure S11**

Compounds that target core-fitness genes were more likely to induce phenotypes that are informative of MoA, but this result was statistically significant ($p < 0.05$) in only 6 of the 15 cell lines.

(a) Summary graph, average MoA counts over the 15 reporter cell lines shown in (b).

(b) Table showing MoA counts of the four possible categories and associated Fisher's exact test results for each of the 15 reporter cell lines individually; based on all MoAs with ≥3 active target-annotated compounds in the respective cell line.

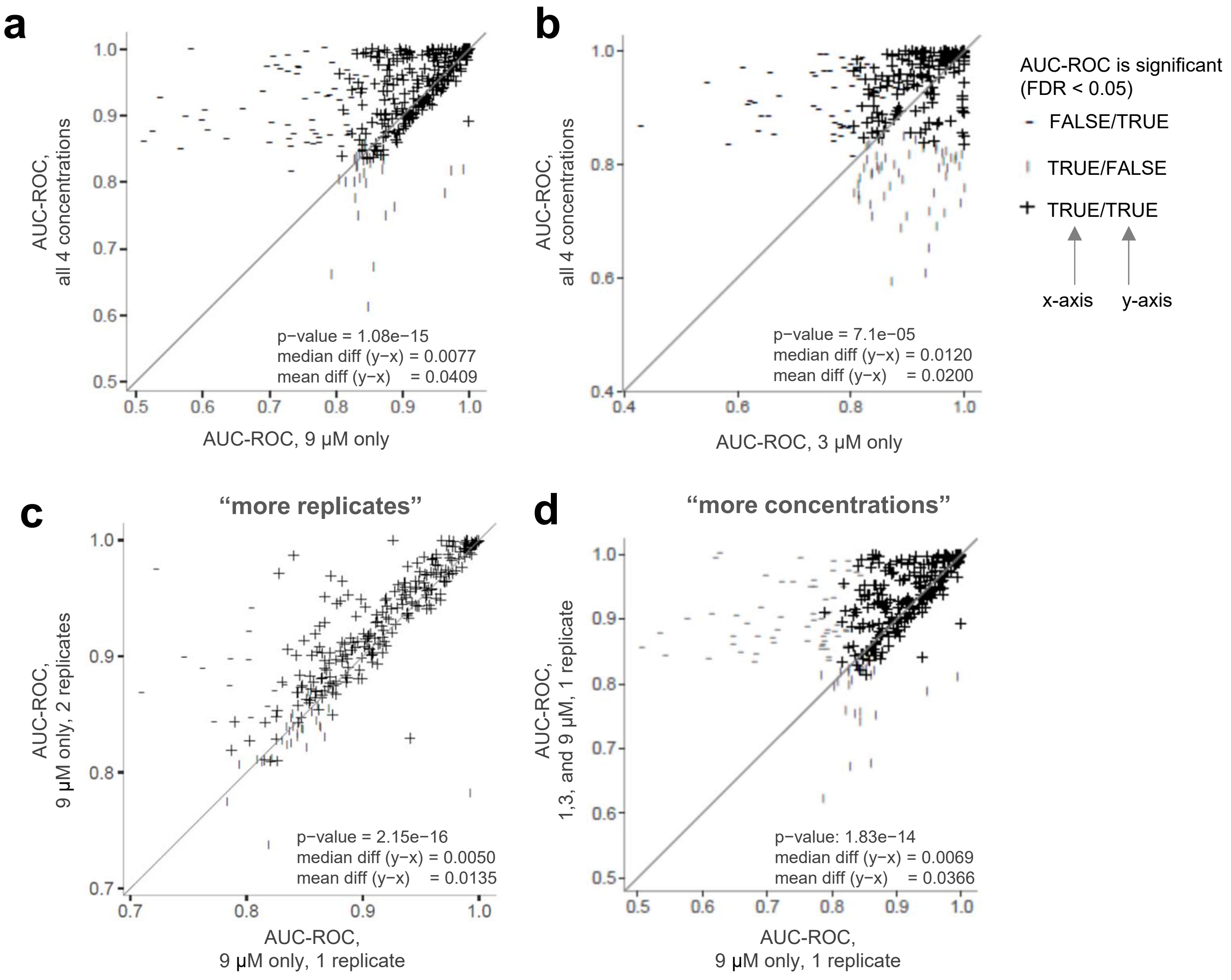


**Supplementary Figure S12**

Comparisons of MoA distinction performance in terms of AUC-ROC for certain subsets of the data.

Each point corresponds to a MoA/cell line pair. Statistical significance of differences in AUC-ROC were assessed with one-sided Wilcoxon signed rank tests with the alternative hypothesis that there is a positive shift of the AUC-ROC values on the y-axis compared to the AUC-ROC values on the x-axis.

(a-b) Distinction performance significantly improved with more concentrations. (a) AUC-ROC computed based on only the 9 µM data aggregated over both batches versus AUC-ROC computed based on data from all four concentrations aggregated over both batches (i.e., the entire available data set). (b) 3 µM versus all concentrations; analogously to (a). While the overall distinction performance significantly improved with more concentrations, for some MoAs considering only the 3 µM data yielded better performance.

(c-d) While more replicates of a single concentration also significantly improved MoA distinction performance, the improvement was 2.7 times larger on average when having more concentrations with a single replicate. This comparison was done such that for (c) and (d) approximately the same number of data points per compound was considered in the AUC-ROC computation. (c) AUC-ROC computed based on only the 9 µM data from Batch 1 (i.e., 1 data point per compound) versus AUC-ROC computed based on the 9 µM data from both batches, thereby keeping the two batches as separate data points (i.e., 2 data points per compound). (d) AUC-ROC computed based on only the 9 µM data from Batch 1 (i.e., 1 data point per compound) versus AUC-ROC computed based on the 1, 3, and 9 µM data from Batch 1 (resulting in 2.06 data points per compound on average; note that because only active treatments were considered the effective number of data points per compound was <3).

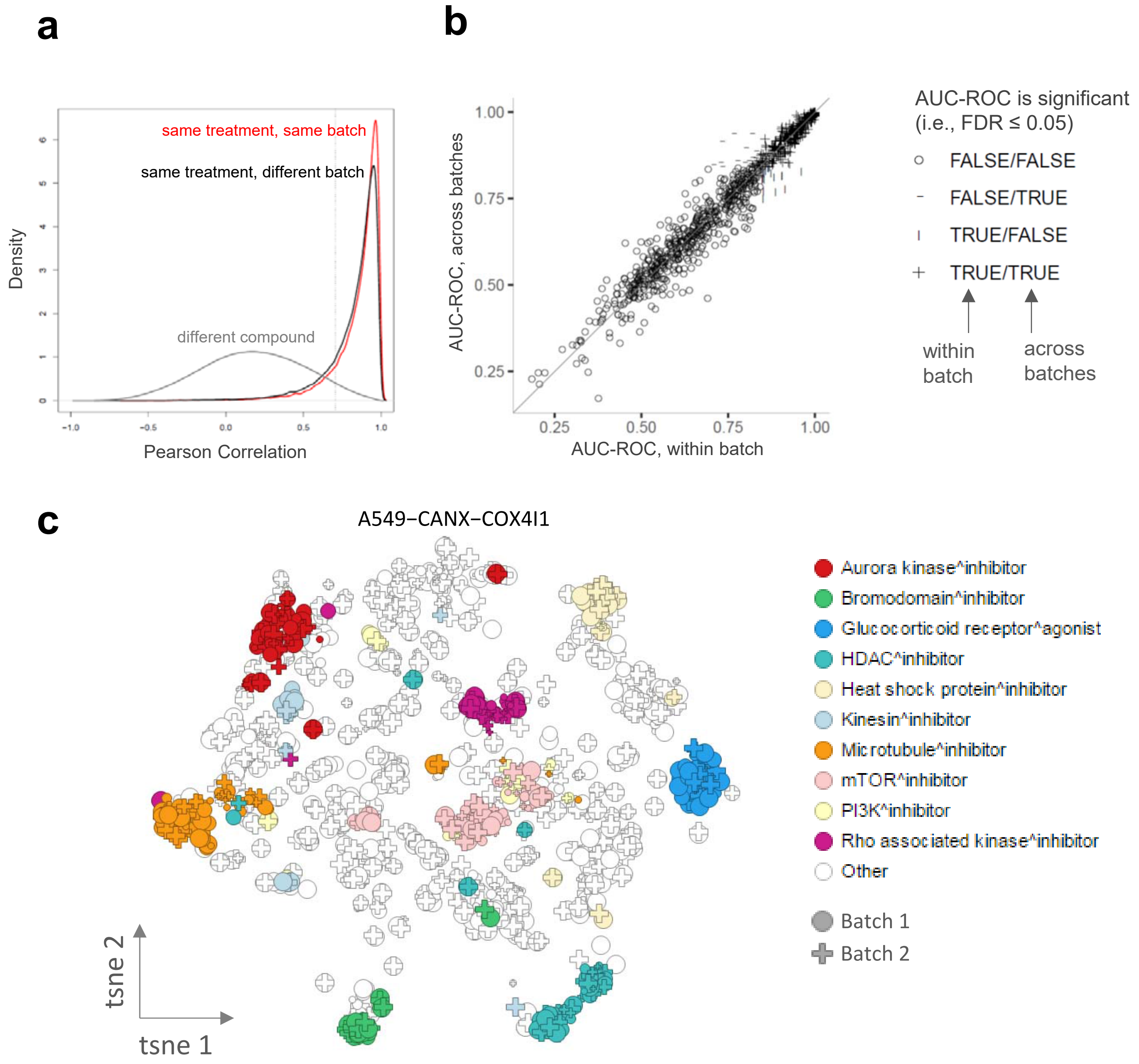


**Supplementary Figure S13**

Assessment of batch effects.

(a) Replicates of active treatments produced highly similar imaging signatures within and across batches. 87.2% of the replicate pairs within the same batch (red line) and 83.4% of the replicate pairs across batches (black line) show higher correlation than the 95th percentile (dashed vertical line) of the correlations between the signatures of different compounds (grey line). Correlations were computed separately in each cell line for all pairs of wells belonging to active treatments (here: based on Euclidean distance to DMSO controls only) and then pooled into the three distributions shown.

(b) MoA AUC-ROC did not significantly drop when comparing signatures across different batches. Each point corresponds to a MoA/cell line pair. AUC-ROC for "within batch" (horizontal axis) was computed based on the signatures from Batch 1 for all treatments; AUC-ROC value for "across batches" (vertical axis) was computed based on the signature from Batch 1 for the query treatment and the signatures from Batch 2 for all other treatments ranked against the query treatment. No significant difference between AUC-ROC "within batch" and AUC-ROC "across batch" was observed (p-value = 0.372; two-sided Wilcoxon signed rank test); median difference (AUC-ROC "across batches" – AUC-ROC "within batches") = 0; mean difference (AUC-ROC "across batches" – AUC-ROC "within batches") = -0.000298.

(c) Consistent with the results in (b) t-SNE map of active treatments in the A549-CANX-COX4I1 cell line (as illustrative example) indicates absence of sizeable batch effects. Each point corresponds to a distinct treatment and batch. Size indicates concentration (0.3 – 9 µM).
