## Supplementary Figure S6 for "Tales of 1,008 Small Molecules: Phenomic Profiling through Live-cell Imaging in a Panel of Reporter Cell Lines"

**Supplementary Figure S6**
t-SNE maps for each of the 15 cell lines, analogously to Fig. 3 in the main manuscript.

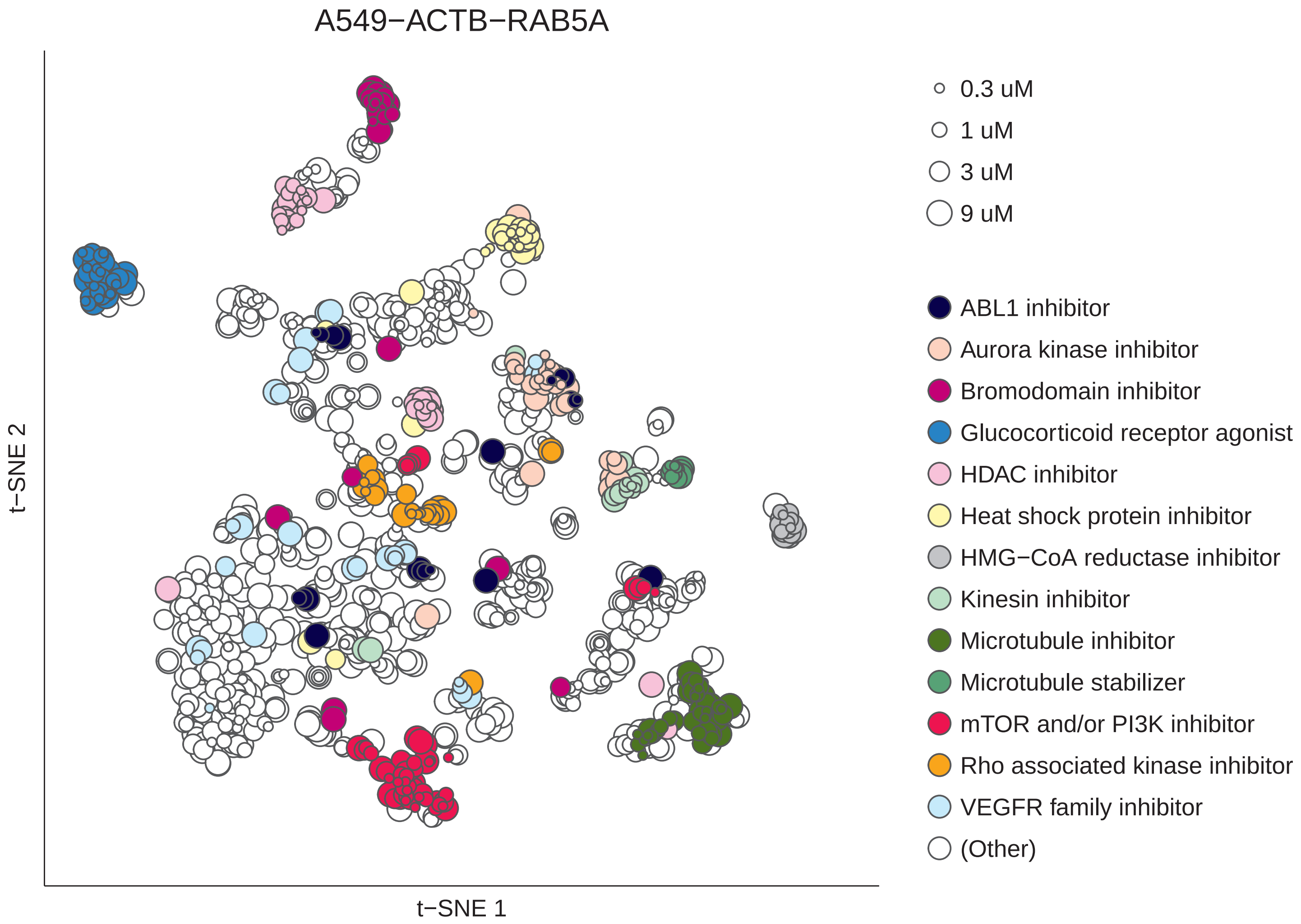
A549−ACTB−RAB5A
t−SNE 2
t−SNE 1
0.3 uM
1 uM
3 uM
9 uM
ABL1 inhibitor
Aurora kinase inhibitor
Bromodomain inhibitor
Glucocorticoid receptor agonist
HDAC inhibitor
Heat shock protein inhibitor
HMG−CoA reductase inhibitor
Kinesin inhibitor
Microtubule inhibitor
Microtubule stabilizer
mTOR and/or PI3K inhibitor
Rho associated kinase inhibitor
VEGFR family inhibitor
(Other)

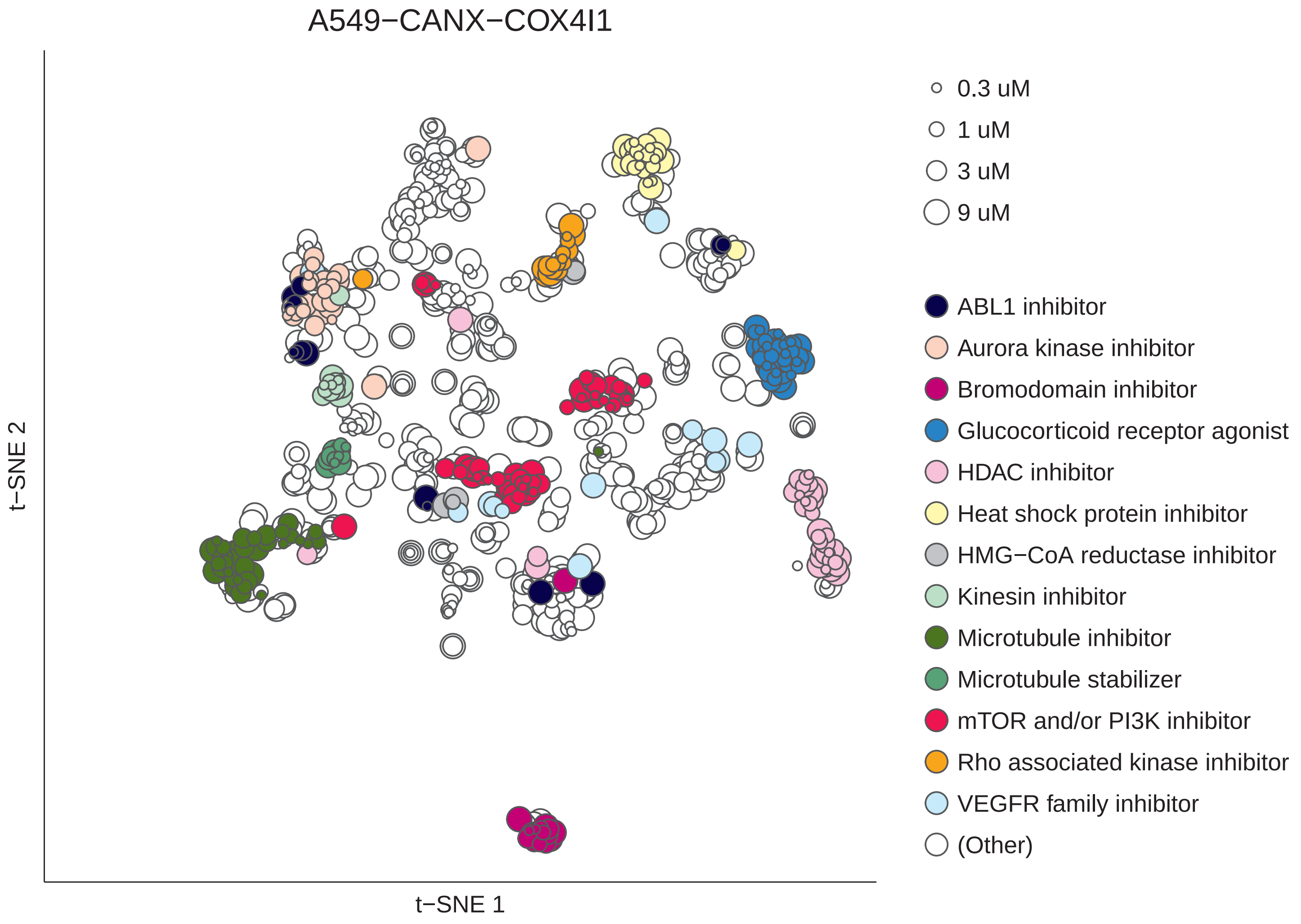
A549−CANX−COX4I1
t−SNE 2
t−SNE 1
0.3 uM
1 uM
3 uM
9 uM
ABL1 inhibitor
Aurora kinase inhibitor
Bromodomain inhibitor
Glucocorticoid receptor agonist
HDAC inhibitor
Heat shock protein inhibitor
HMG−CoA reductase inhibitor
Kinesin inhibitor
Microtubule inhibitor
Microtubule stabilizer
mTOR and/or PI3K inhibitor
Rho associated kinase inhibitor
VEGFR family inhibitor
(Other)

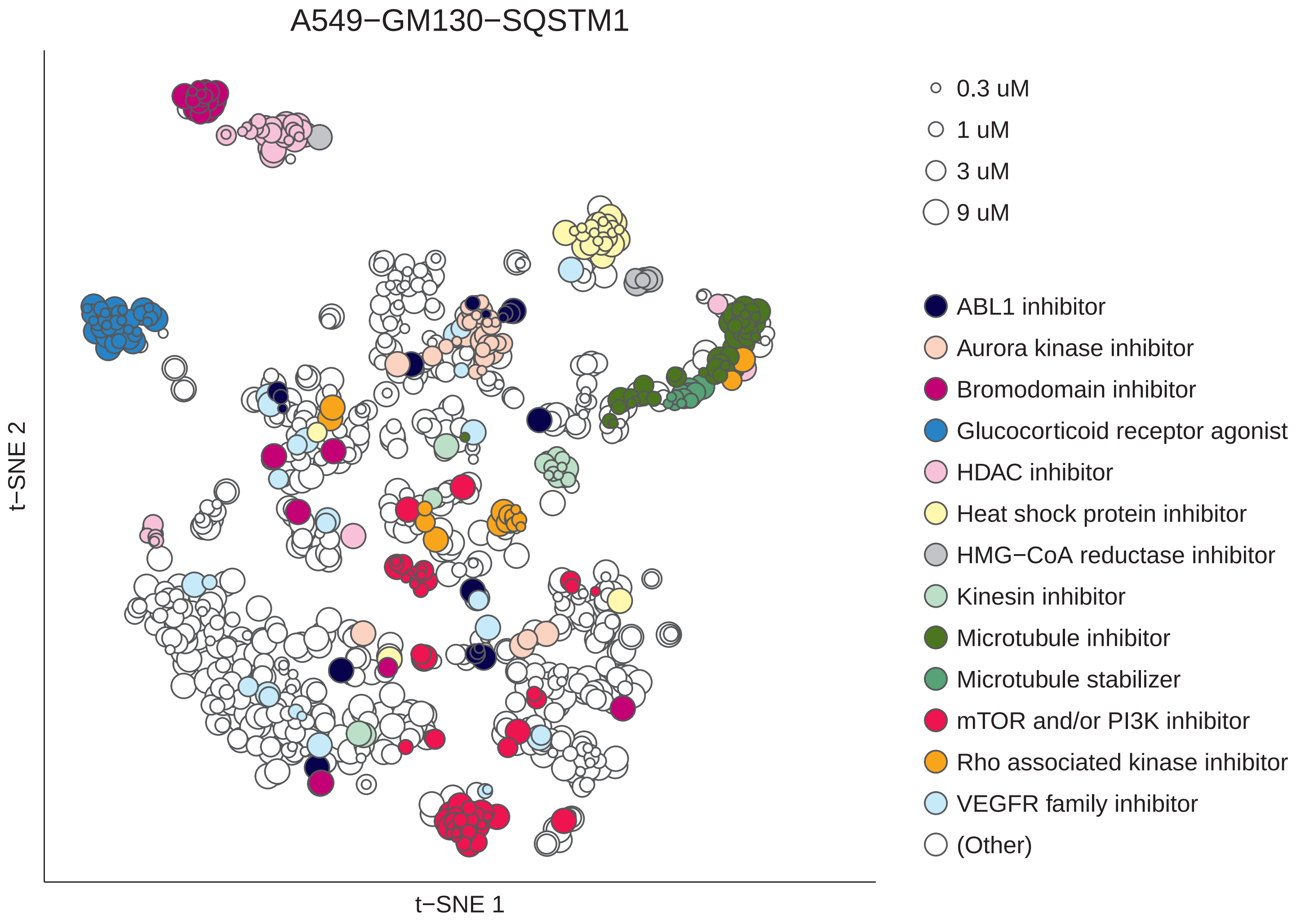

A549−GM130−SQSTM1
t−SNE 2
t−SNE 1
0.3 uM
1 uM
3 uM
9 uM
ABL1 inhibitor
Aurora kinase inhibitor
Bromodomain inhibitor
Glucocorticoid receptor agonist
HDAC inhibitor
Heat shock protein inhibitor
HMG−CoA reductase inhibitor
Kinesin inhibitor
Microtubule inhibitor
Microtubule stabilizer
mTOR and/or PI3K inhibitor
Rho associated kinase inhibitor
VEGFR family inhibitor
(Other)

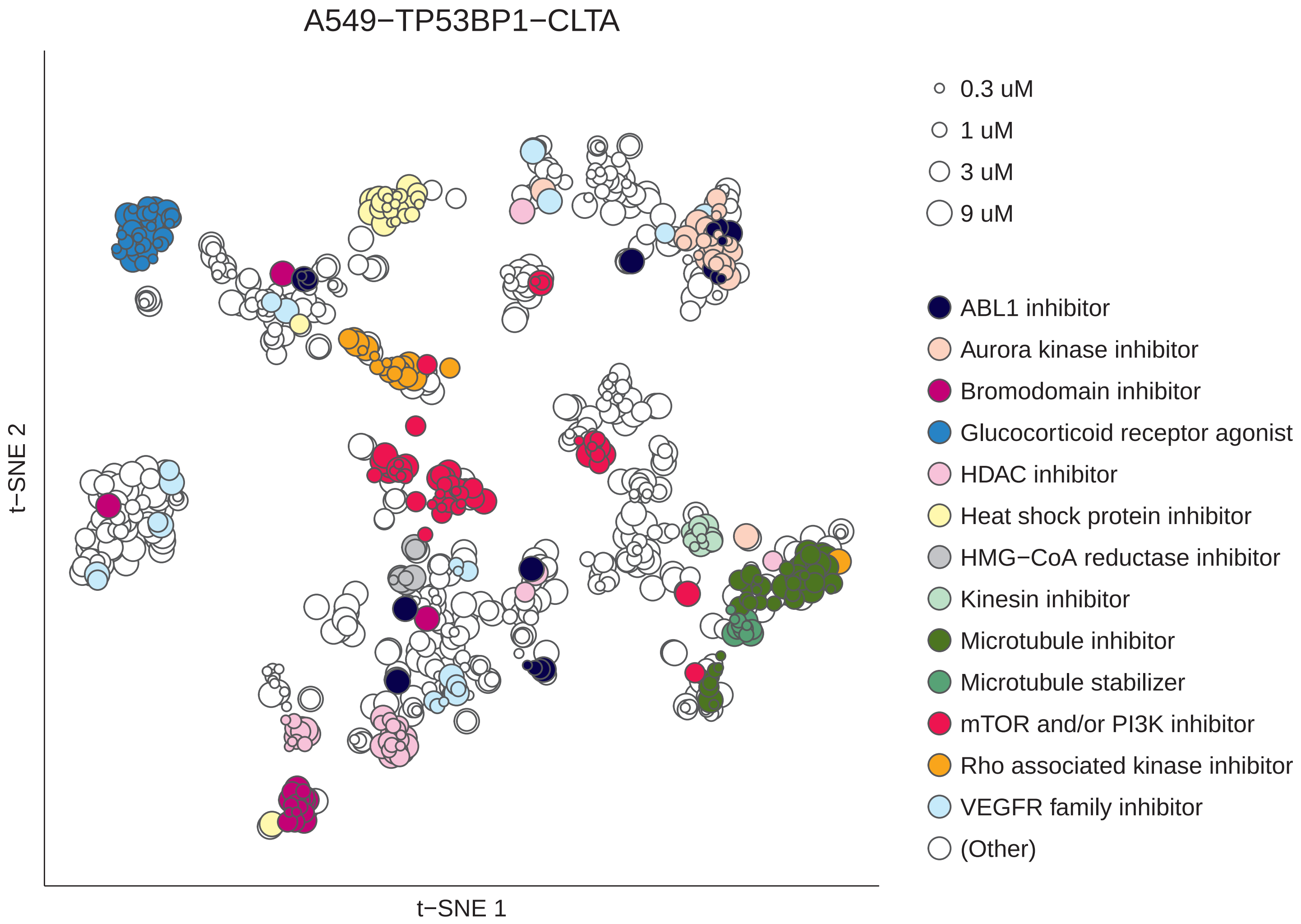
A549−TP53BP1−CLTA
t−SNE 2
t−SNE 1
0.3 uM
1 uM
3 uM
9 uM
ABL1 inhibitor
Aurora kinase inhibitor
Bromodomain inhibitor
Glucocorticoid receptor agonist
HDAC inhibitor
Heat shock protein inhibitor
HMG−CoA reductase inhibitor
Kinesin inhibitor
Microtubule inhibitor
Microtubule stabilizer
mTOR and/or PI3K inhibitor
Rho associated kinase inhibitor
VEGFR family inhibitor
(Other)

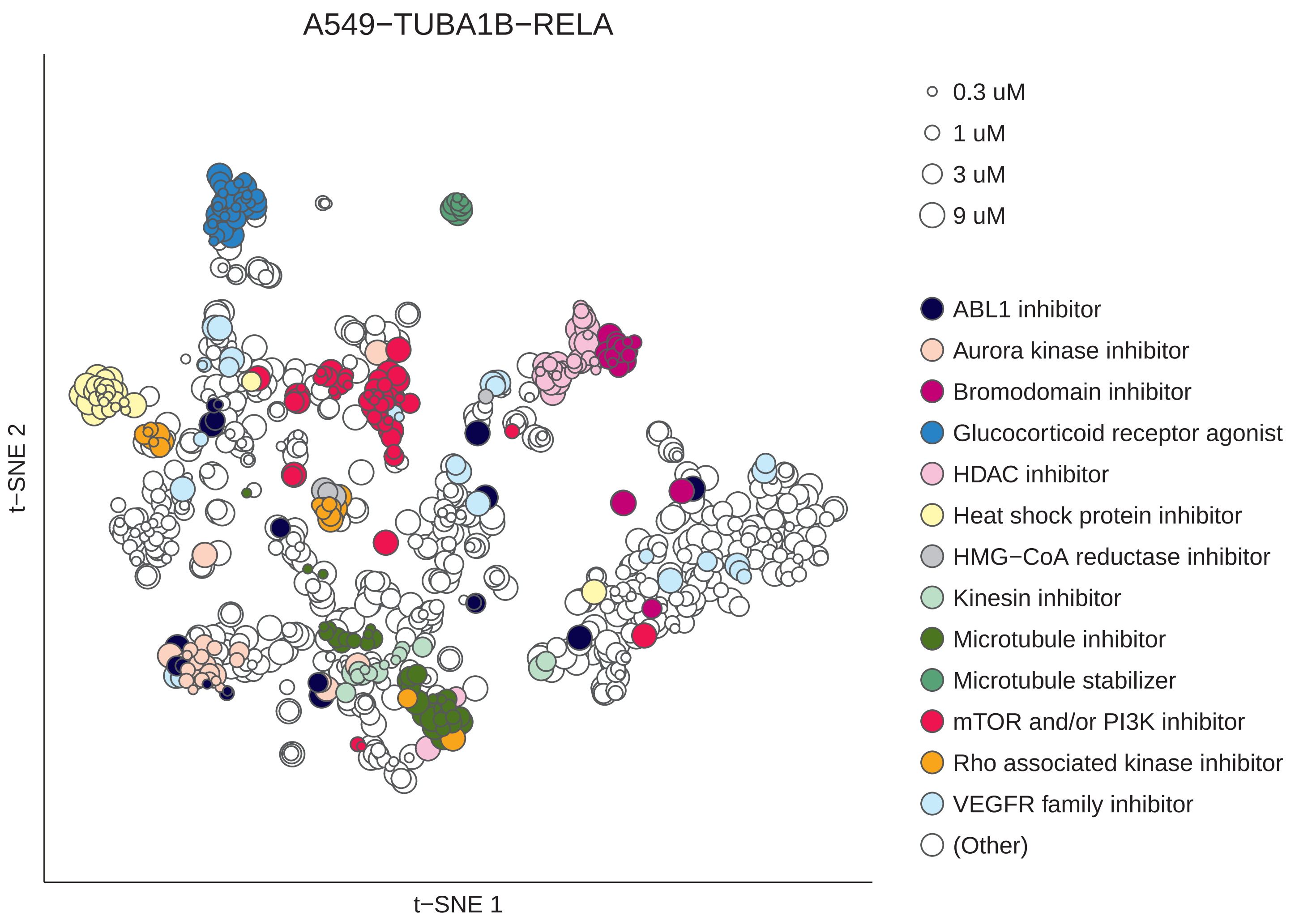
A549−TUBA1B−RELA
t−SNE 2
t−SNE 1
0.3 uM
1 uM
3 uM
9 uM
ABL1 inhibitor
Aurora kinase inhibitor
Bromodomain inhibitor
Glucocorticoid receptor agonist
HDAC inhibitor
Heat shock protein inhibitor
HMG−CoA reductase inhibitor
Kinesin inhibitor
Microtubule inhibitor
Microtubule stabilizer
mTOR and/or PI3K inhibitor
Rho associated kinase inhibitor
VEGFR family inhibitor
(Other)

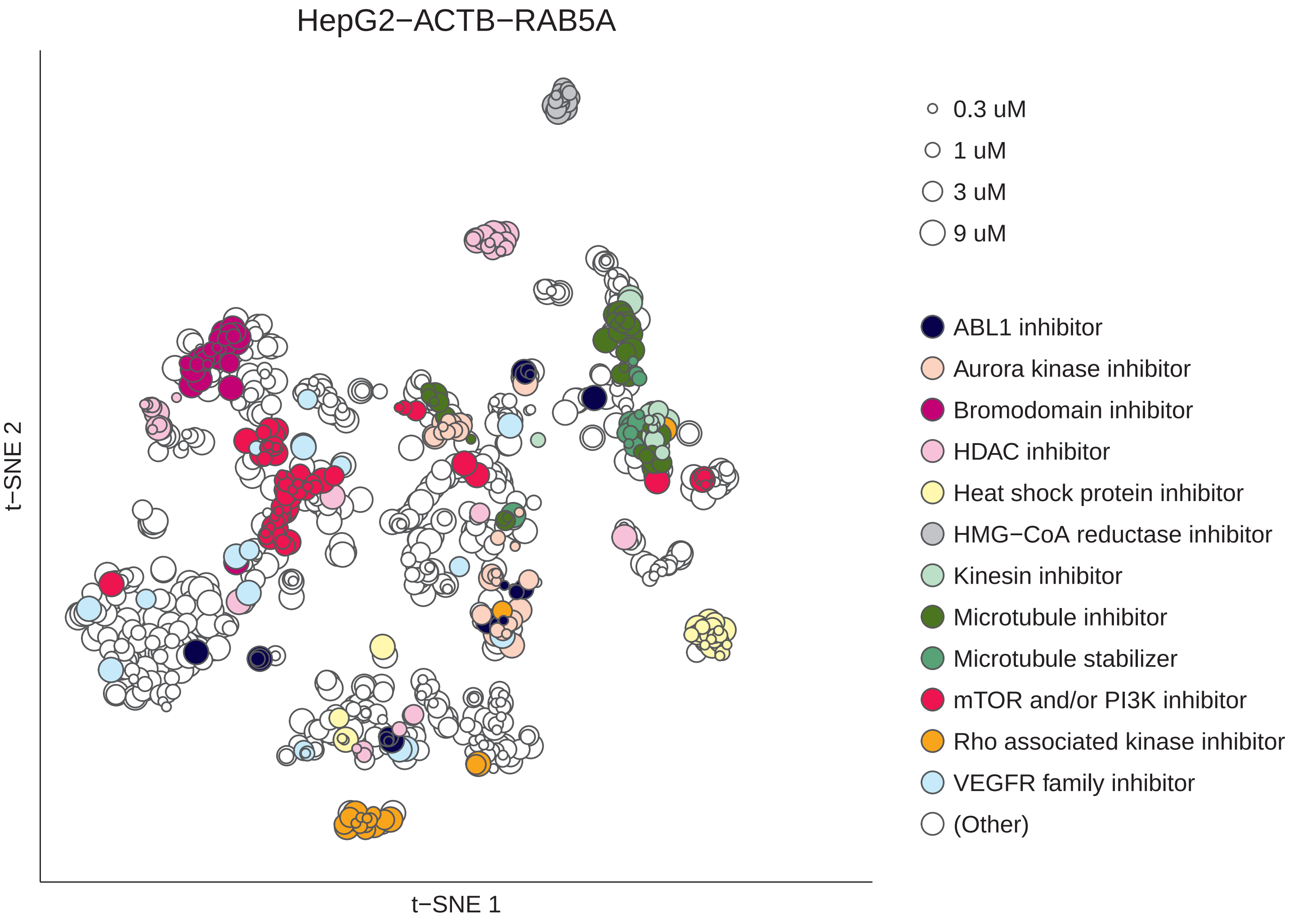
HepG2−ACTB−RAB5A
t−SNE 2
t−SNE 1
0.3 uM
1 uM
3 uM
9 uM
ABL1 inhibitor
Aurora kinase inhibitor
Bromodomain inhibitor
HDAC inhibitor
Heat shock protein inhibitor
HMG−CoA reductase inhibitor
Kinesin inhibitor
Microtubule inhibitor
Microtubule stabilizer
mTOR and/or PI3K inhibitor
Rho associated kinase inhibitor
VEGFR family inhibitor
(Other)

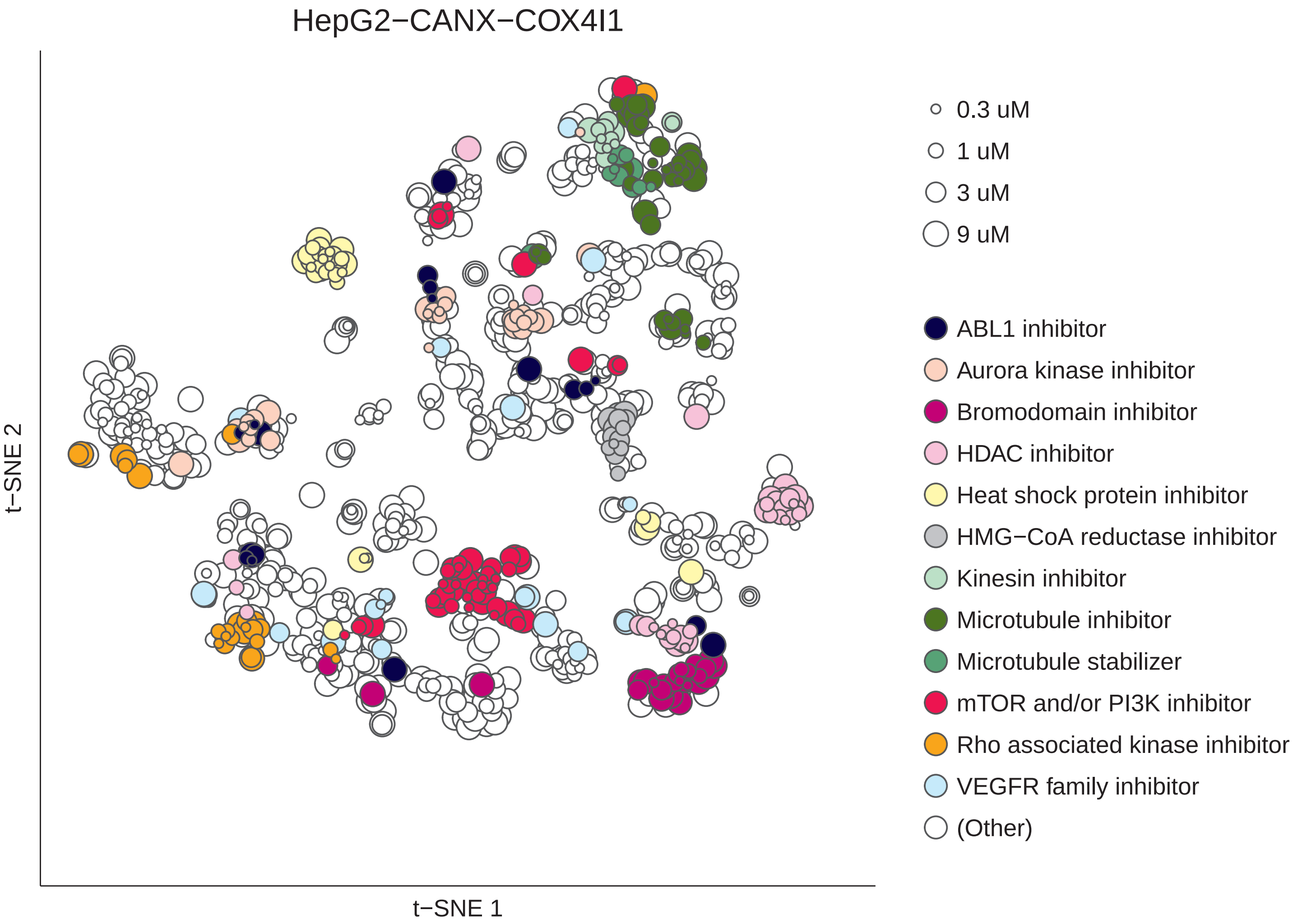
HepG2−CANX−COX4I1
t−SNE 2
t−SNE 1
0.3 uM
1 uM
3 uM
9 uM
ABL1 inhibitor
Aurora kinase inhibitor
Bromodomain inhibitor
HDAC inhibitor
Heat shock protein inhibitor
HMG−CoA reductase inhibitor
Kinesin inhibitor
Microtubule inhibitor
Microtubule stabilizer
mTOR and/or PI3K inhibitor
Rho associated kinase inhibitor
VEGFR family inhibitor
(Other)

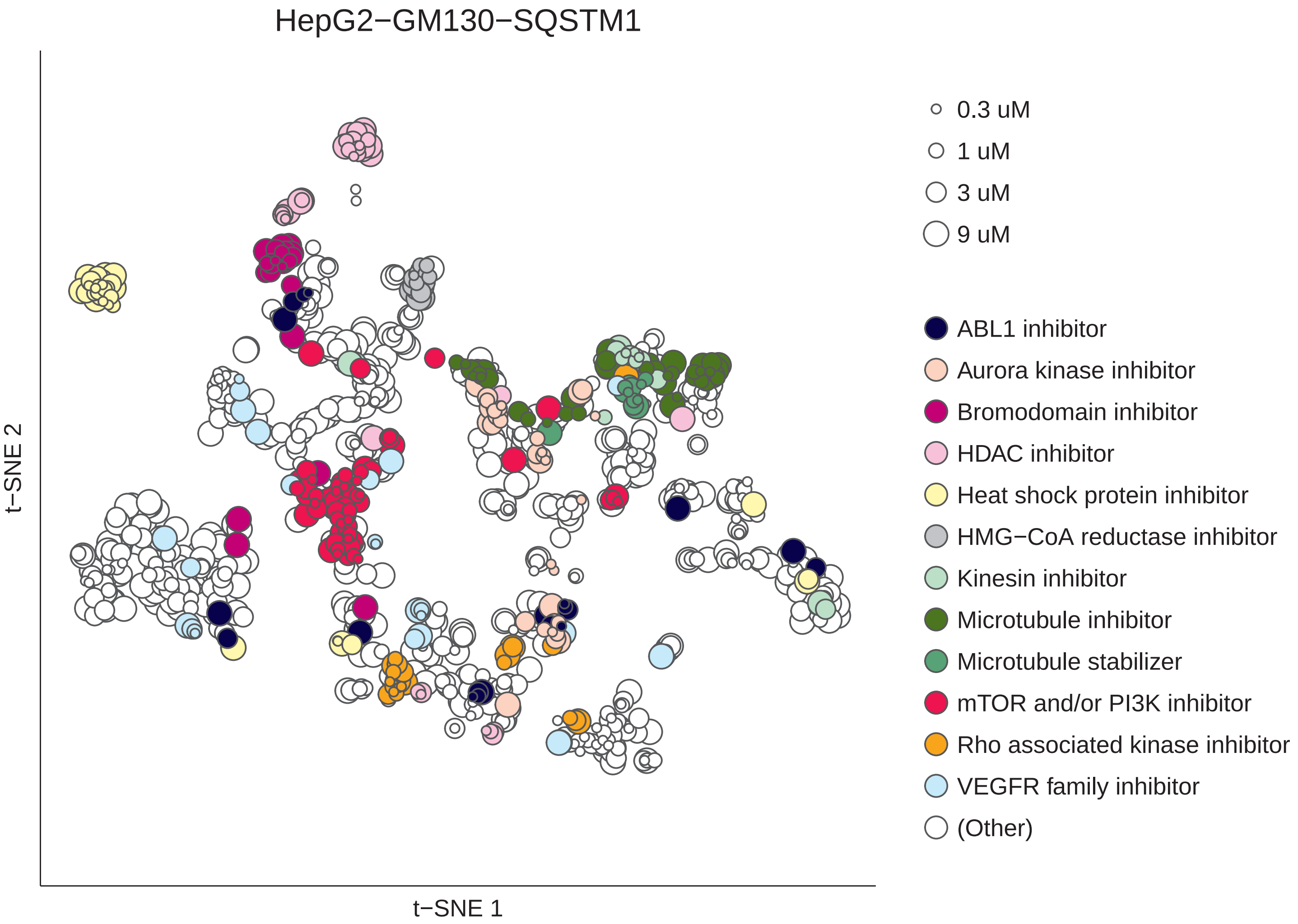
HepG2−GM130−SQSTM1
t−SNE 1
t−SNE 2
0.3 uM
1 uM
3 uM
9 uM
ABL1 inhibitor
Aurora kinase inhibitor
Bromodomain inhibitor
HDAC inhibitor
Heat shock protein inhibitor
HMG−CoA reductase inhibitor
Kinesin inhibitor
Microtubule inhibitor
Microtubule stabilizer
mTOR and/or PI3K inhibitor
Rho associated kinase inhibitor
VEGFR family inhibitor
(Other)

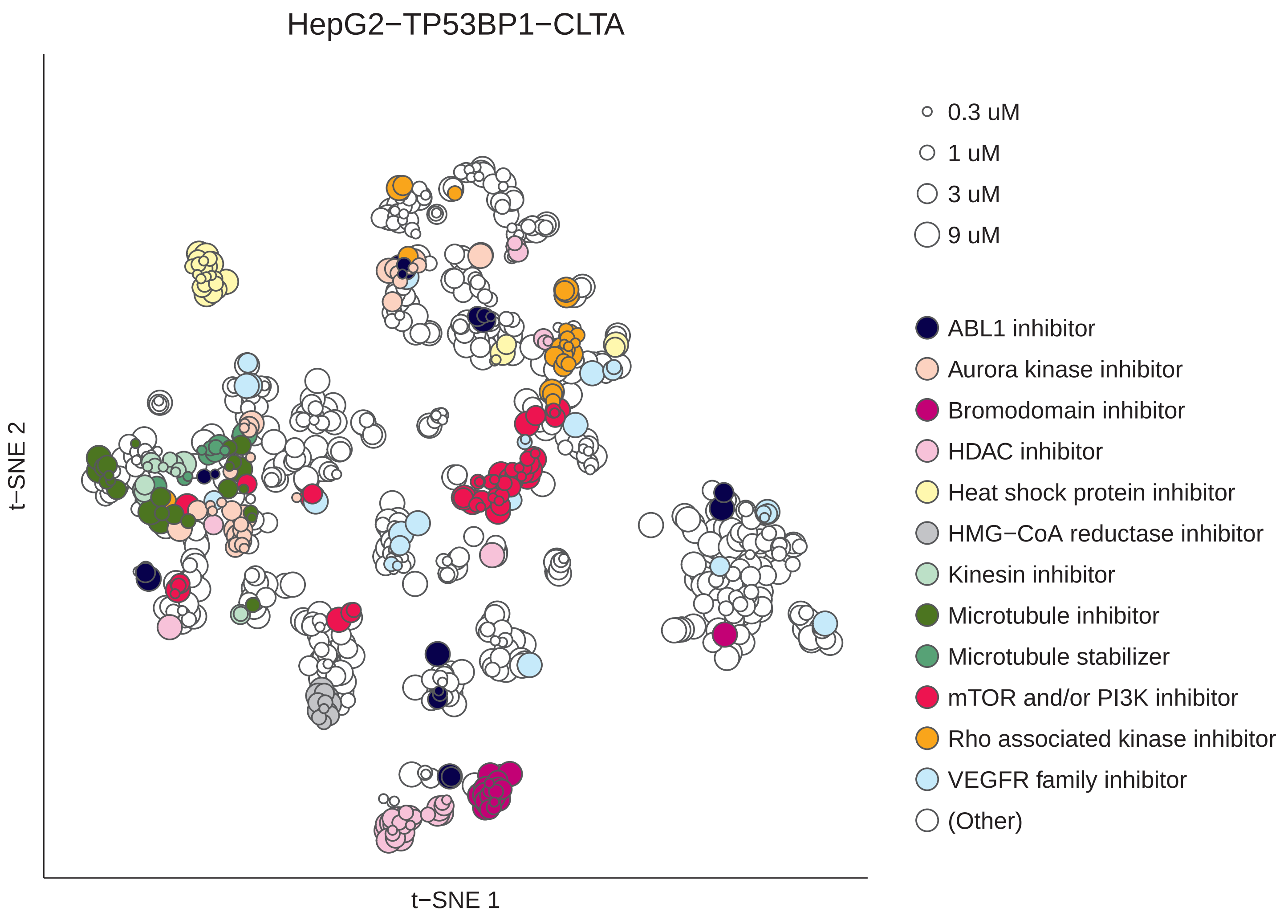
HepG2−TP53BP1−CLTA
t−SNE 2
t−SNE 1
0.3 uM
1 uM
3 uM
9 uM
ABL1 inhibitor
Aurora kinase inhibitor
Bromodomain inhibitor
HDAC inhibitor
Heat shock protein inhibitor
HMG−CoA reductase inhibitor
Kinesin inhibitor
Microtubule inhibitor
Microtubule stabilizer
mTOR and/or PI3K inhibitor
Rho associated kinase inhibitor
VEGFR family inhibitor
(Other)

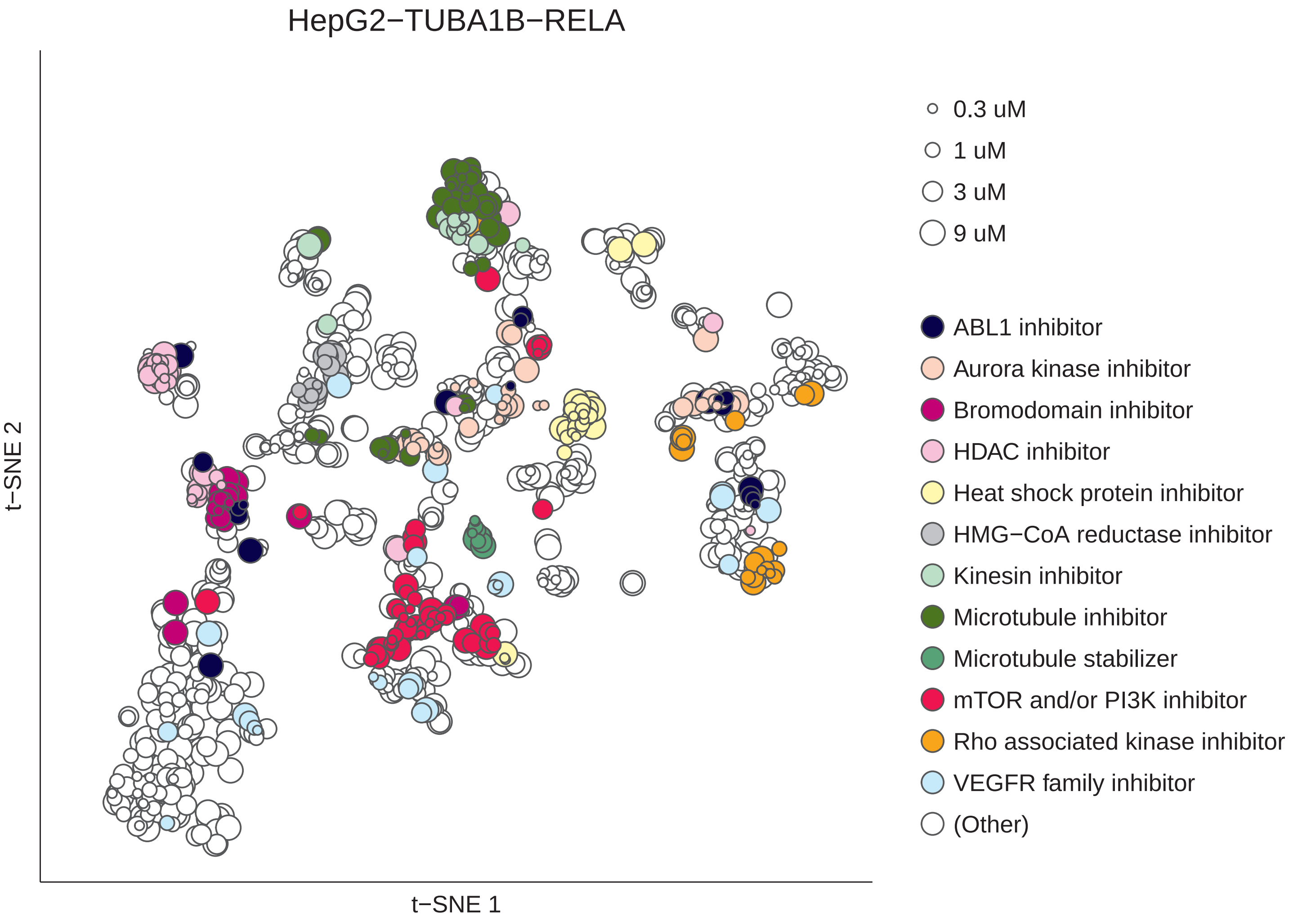
HepG2−TUBA1B−RELA
t−SNE 2
t−SNE 1
0.3 uM
1 uM
3 uM
9 uM
ABL1 inhibitor
Aurora kinase inhibitor
Bromodomain inhibitor
HDAC inhibitor
Heat shock protein inhibitor
HMG−CoA reductase inhibitor
Kinesin inhibitor
Microtubule inhibitor
Microtubule stabilizer
mTOR and/or PI3K inhibitor
Rho associated kinase inhibitor
VEGFR family inhibitor
(Other)

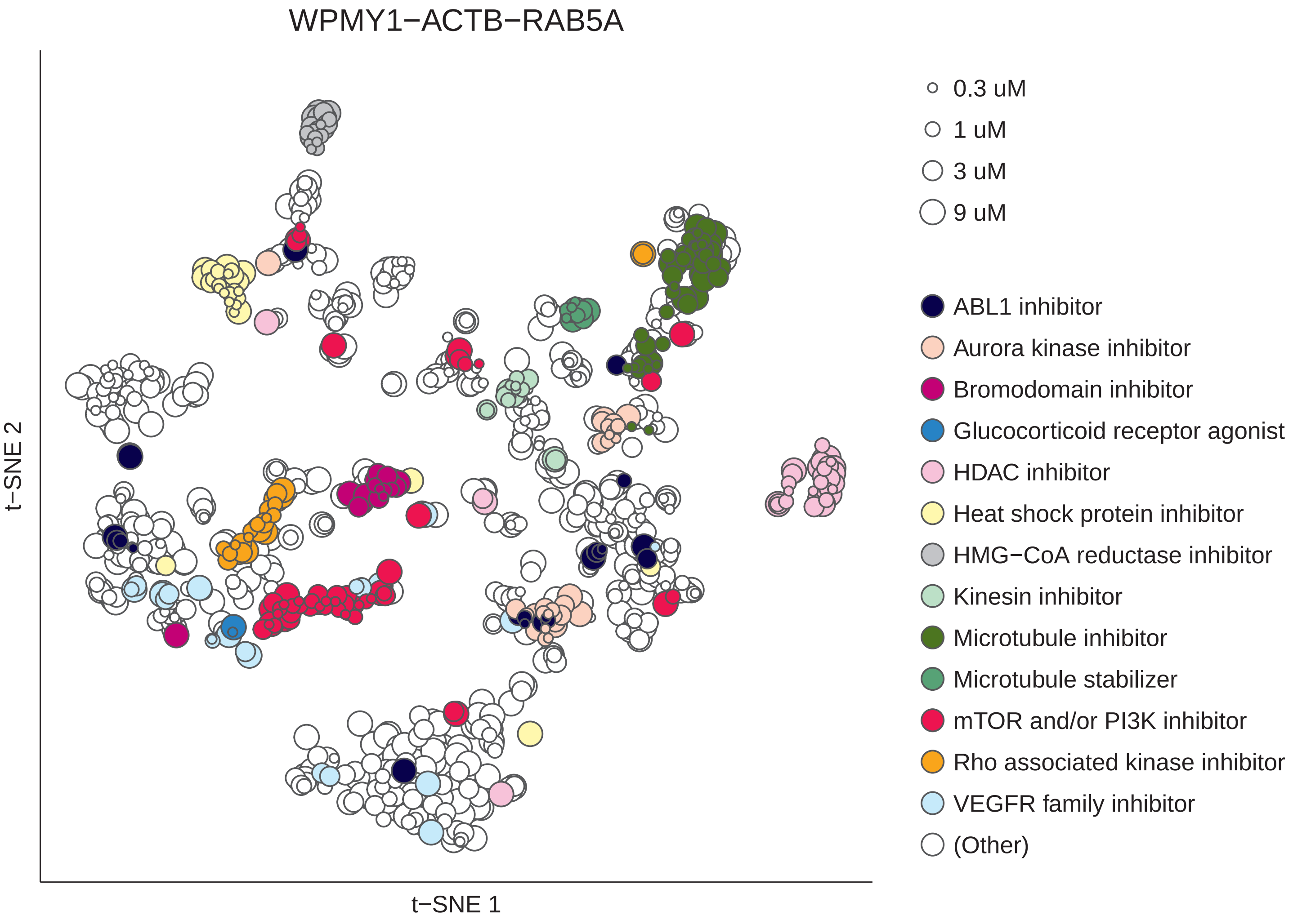
WPMY1−ACTB−RAB5A
t−SNE 2
t−SNE 1
0.3 uM
1 uM
3 uM
9 uM
ABL1 inhibitor
Aurora kinase inhibitor
Bromodomain inhibitor
Glucocorticoid receptor agonist
HDAC inhibitor
Heat shock protein inhibitor
HMG−CoA reductase inhibitor
Kinesin inhibitor
Microtubule inhibitor
Microtubule stabilizer
mTOR and/or PI3K inhibitor
Rho associated kinase inhibitor
VEGFR family inhibitor
(Other)

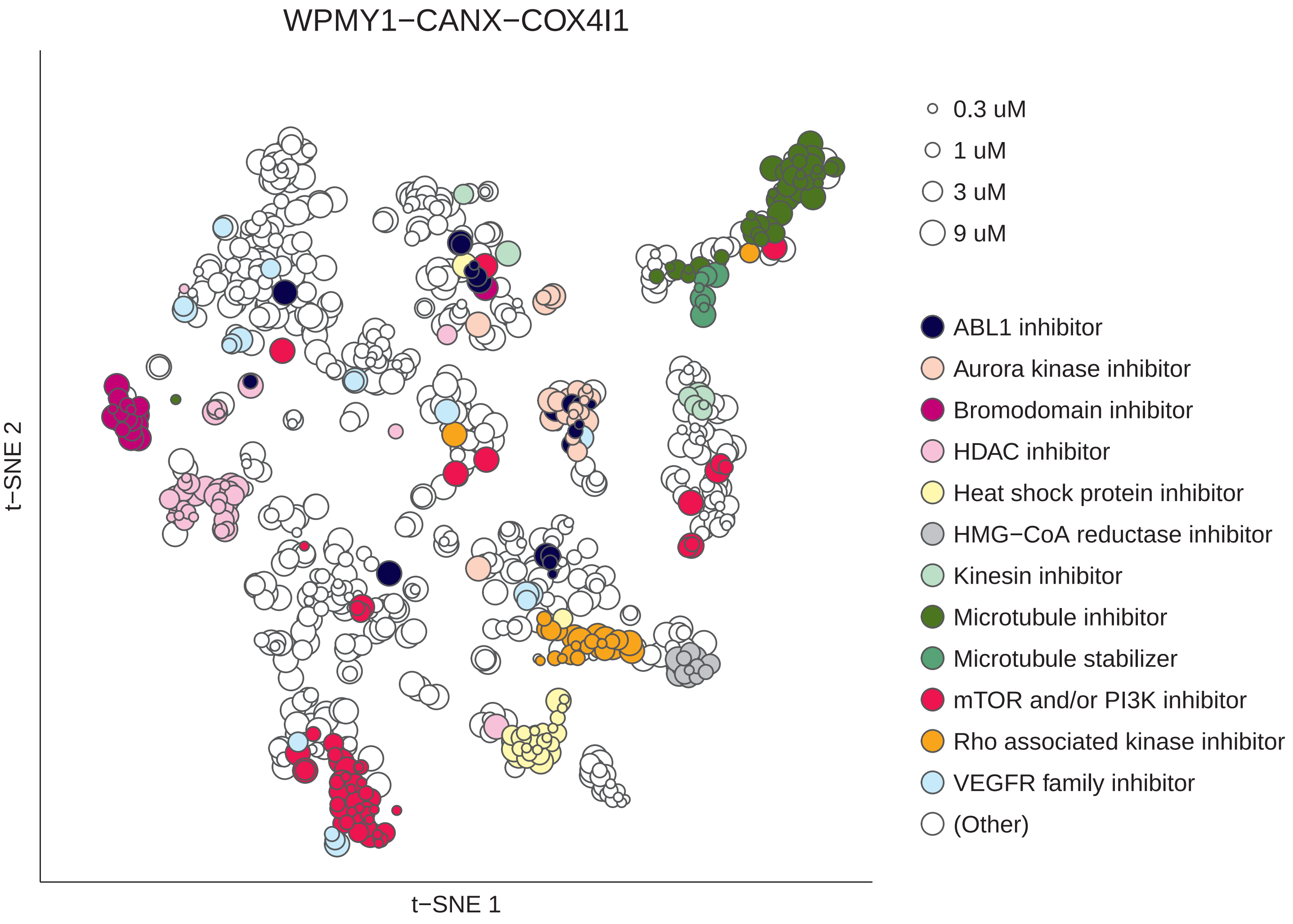

WPMY1−CANX−COX4I1
t−SNE 2
t−SNE 1
0.3 uM
1 uM
3 uM
9 uM
ABL1 inhibitor
Aurora kinase inhibitor
Bromodomain inhibitor
HDAC inhibitor
Heat shock protein inhibitor
HMG−CoA reductase inhibitor
Kinesin inhibitor
Microtubule inhibitor
Microtubule stabilizer
mTOR and/or PI3K inhibitor
Rho associated kinase inhibitor
VEGFR family inhibitor
(Other)

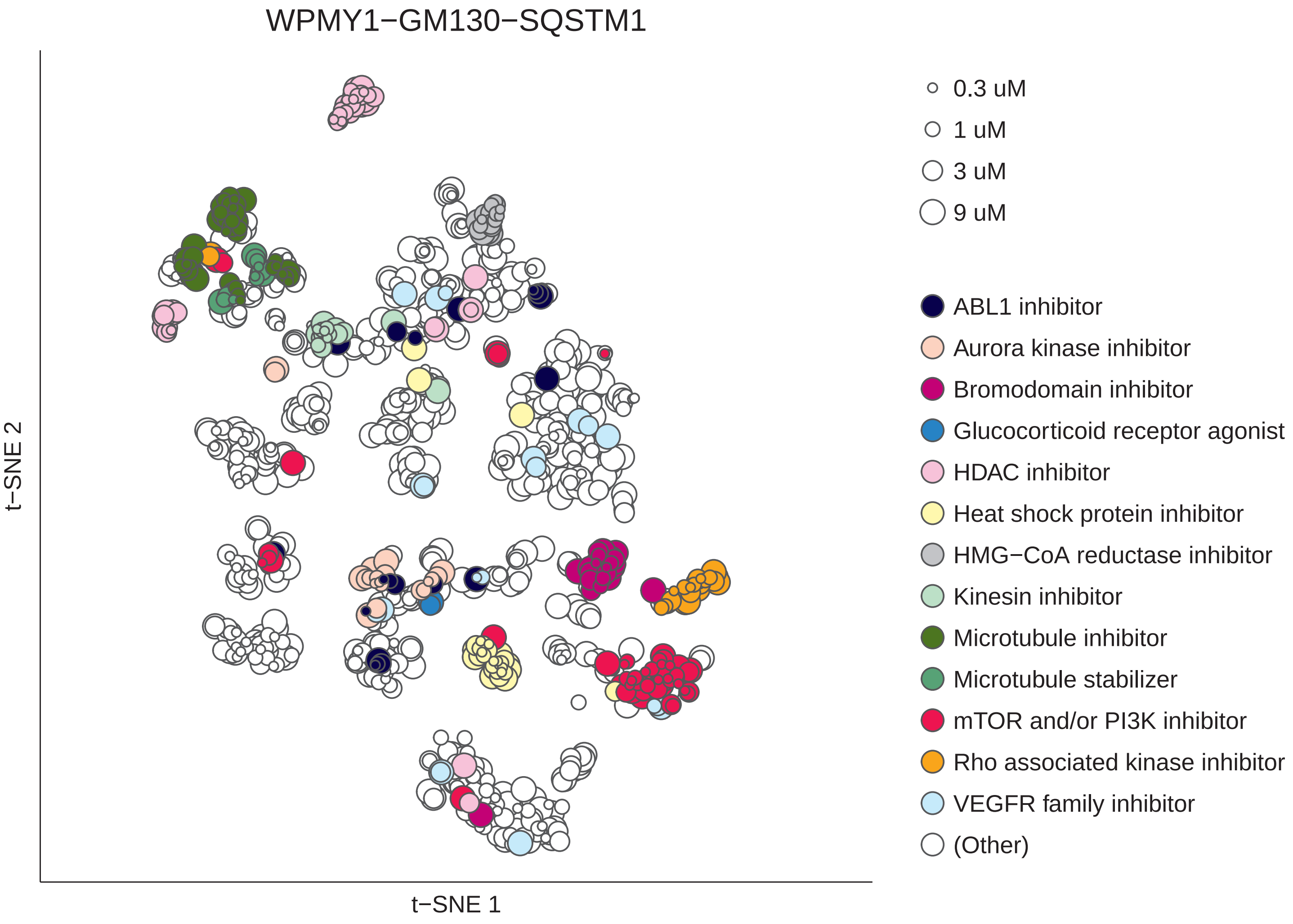
WPMY1−GM130−SQSTM1
t−SNE 2
t−SNE 1
0.3 uM
1 uM
3 uM
9 uM
ABL1 inhibitor
Aurora kinase inhibitor
Bromodomain inhibitor
Glucocorticoid receptor agonist
HDAC inhibitor
Heat shock protein inhibitor
HMG−CoA reductase inhibitor
Kinesin inhibitor
Microtubule inhibitor
Microtubule stabilizer
mTOR and/or PI3K inhibitor
Rho associated kinase inhibitor
VEGFR family inhibitor
(Other)

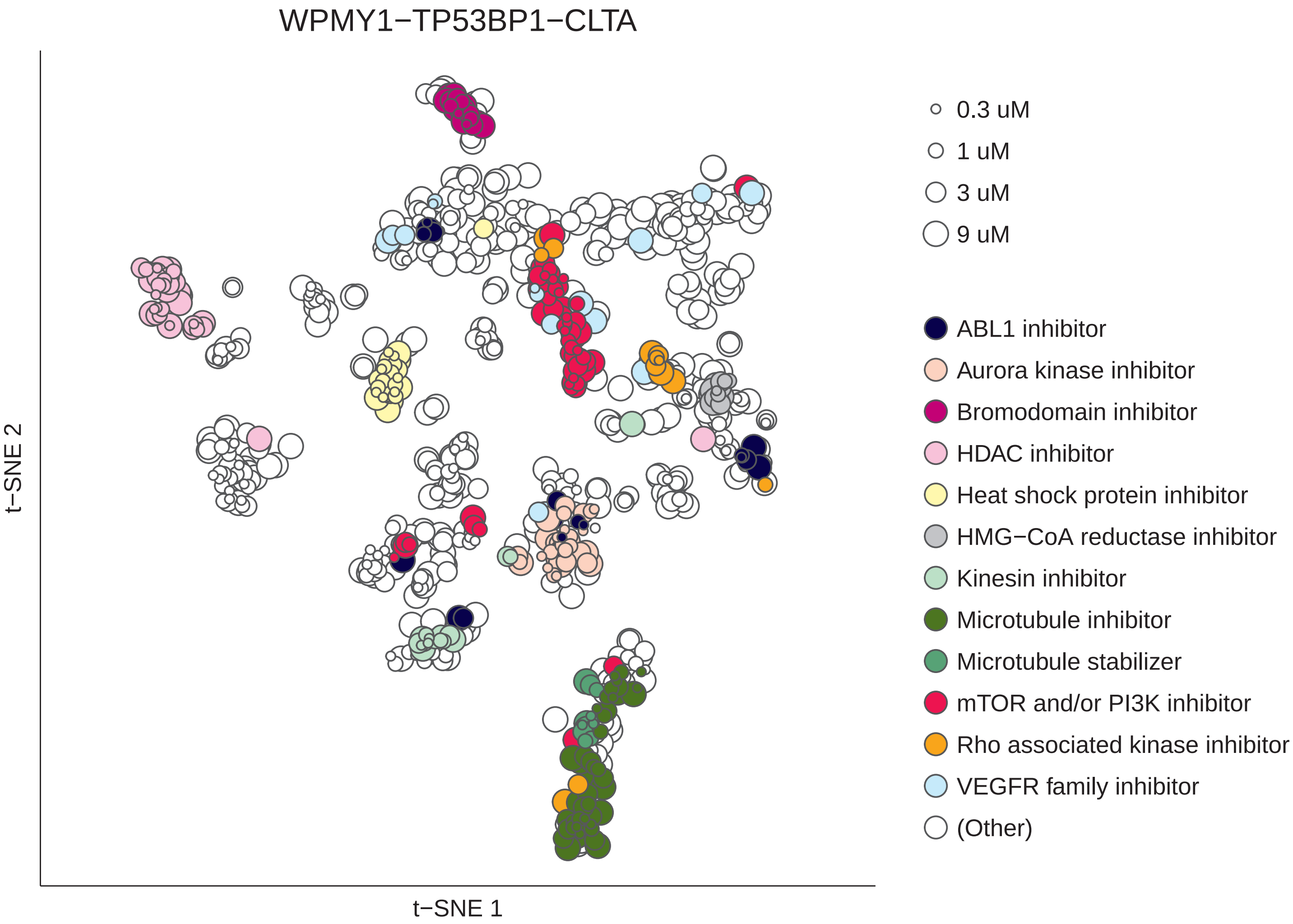

WPMY1−TP53BP1−CLTA
t−SNE 2
t−SNE 1
0.3 uM
1 uM
3 uM
9 uM
ABL1 inhibitor
Aurora kinase inhibitor
Bromodomain inhibitor
HDAC inhibitor
Heat shock protein inhibitor
HMG−CoA reductase inhibitor
Kinesin inhibitor
Microtubule inhibitor
Microtubule stabilizer
mTOR and/or PI3K inhibitor
Rho associated kinase inhibitor
VEGFR family inhibitor
(Other)

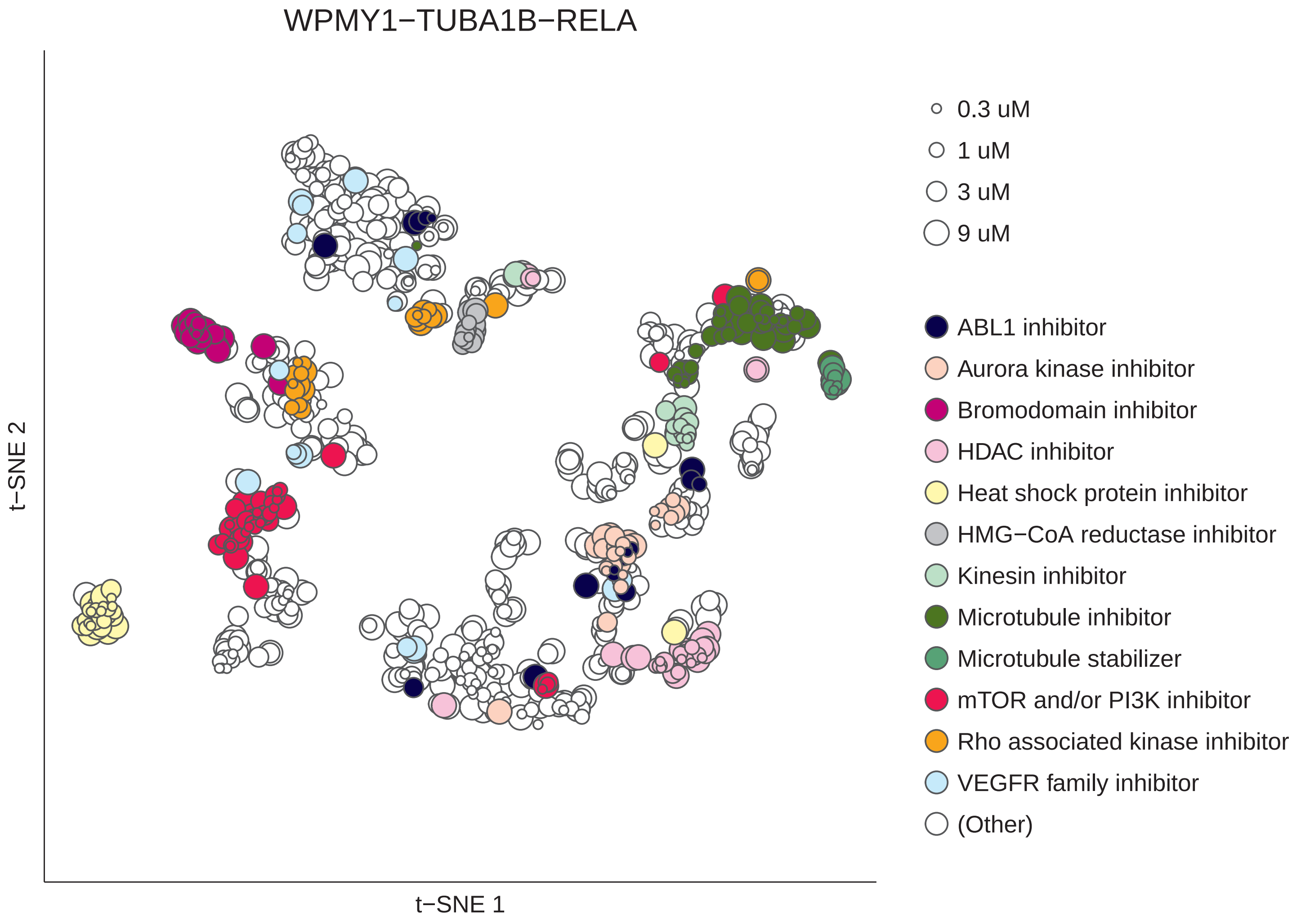
WPMY1−TUBA1B−RELA
t−SNE 2
t−SNE 1
0.3 uM
1 uM
3 uM
9 uM
ABL1 inhibitor
Aurora kinase inhibitor
Bromodomain inhibitor
HDAC inhibitor
Heat shock protein inhibitor
HMG−CoA reductase inhibitor
Kinesin inhibitor
Microtubule inhibitor
Microtubule stabilizer
mTOR and/or PI3K inhibitor
Rho associated kinase inhibitor
VEGFR family inhibitor
(Other)
