## Supplementary Figure S8 for "Tales of 1,008 Small Molecules: Phenomic Profiling through Live-cell Imaging in a Panel of Reporter Cell Lines"

**Supplementary Figure S8**
Graphical representation of the AUC-ROC values for all 83 MoAs with ≥3 active compounds, which are also listed in Supplementary Table S3.

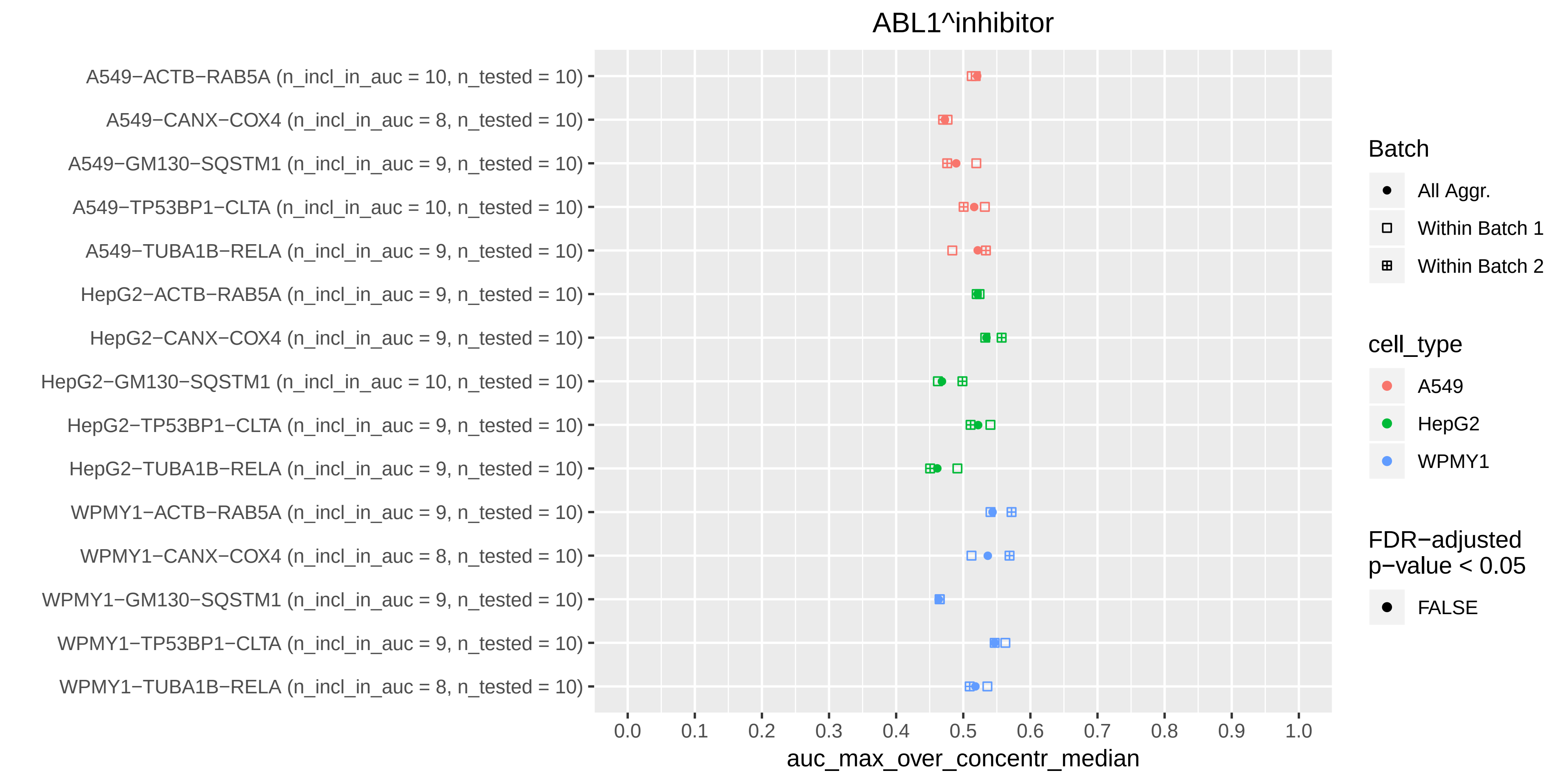
ABL1^inhibitor
A549−ACTB−RAB5A (n_incl_in_auc = 10, n_tested = 10)
A549−CANX−COX4 (n_incl_in_auc = 8, n_tested = 10)
A549−GM130−SQSTM1 (n_incl_in_auc = 9, n_tested = 10)
A549−TP53BP1−CLTA (n_incl_in_auc = 10, n_tested = 10)
A549−TUBA1B−RELA (n_incl_in_auc = 9, n_tested = 10)
HepG2−ACTB−RAB5A (n_incl_in_auc = 9, n_tested = 10)
HepG2−CANX−COX4 (n_incl_in_auc = 9, n_tested = 10)
HepG2−GM130−SQSTM1 (n_incl_in_auc = 10, n_tested = 10)
HepG2−TP53BP1−CLTA (n_incl_in_auc = 9, n_tested = 10)
HepG2−TUBA1B−RELA (n_incl_in_auc = 9, n_tested = 10)
WPMY1−ACTB−RAB5A (n_incl_in_auc = 9, n_tested = 10)
WPMY1−CANX−COX4 (n_incl_in_auc = 8, n_tested = 10)
WPMY1−GM130−SQSTM1 (n_incl_in_auc = 9, n_tested = 10)
WPMY1−TP53BP1−CLTA (n_incl_in_auc = 9, n_tested = 10)
WPMY1−TUBA1B−RELA (n_incl_in_auc = 8, n_tested = 10)
0.0
0.1
0.2
0.3
0.4
0.5
0.6
0.7
0.8
0.9
1.0
auc_max_over_concentr_median
Batch
All Aggr.
Within Batch 1
Within Batch 2
cell_type
A549
HepG2
WPMY1
FDR−adjusted p−value < 0.05
FALSE

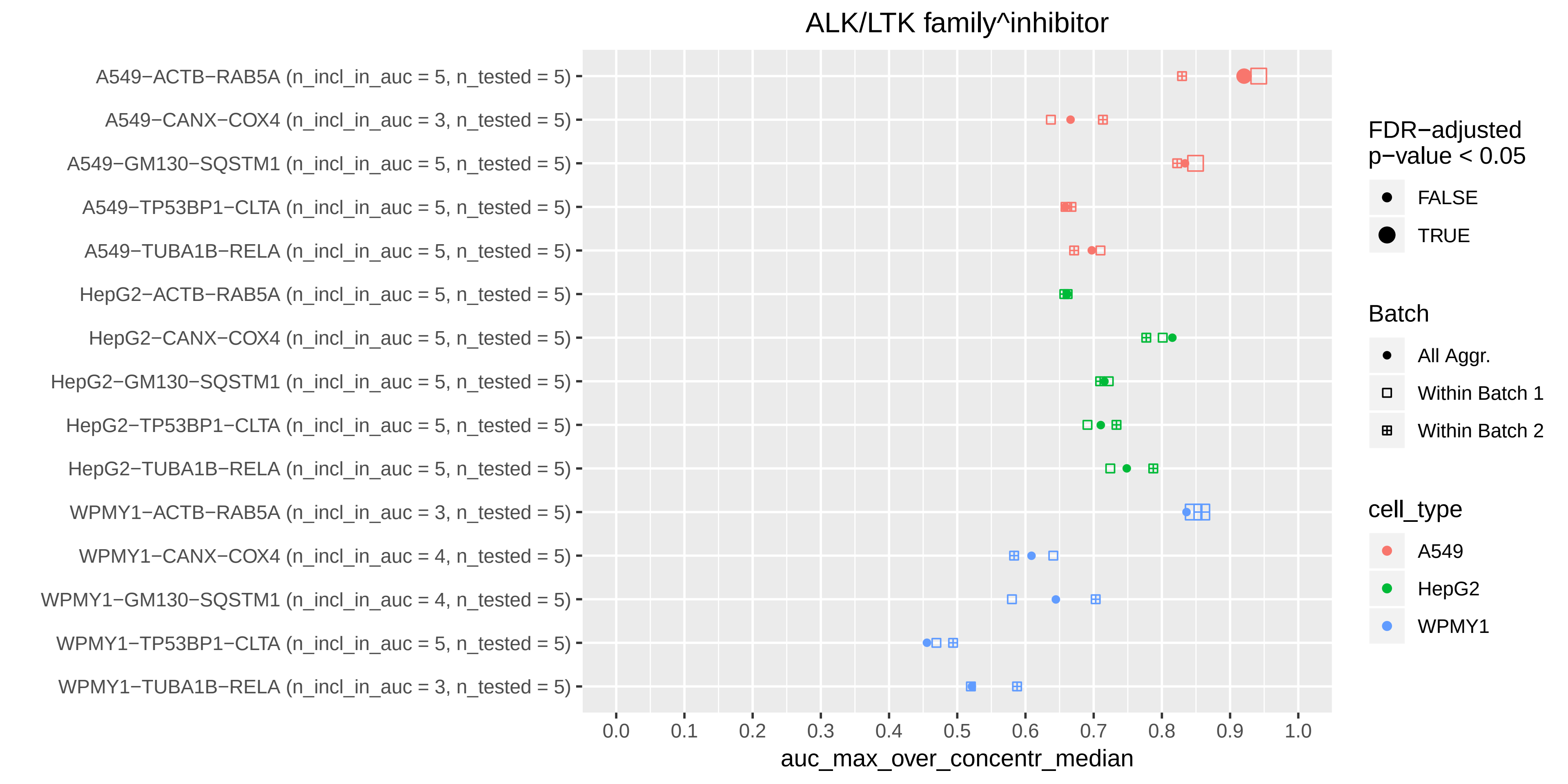
ALK/LTK family^inhibitor
A549−ACTB−RAB5A (n_incl_in_auc = 5, n_tested = 5)
A549−CANX−COX4 (n_incl_in_auc = 3, n_tested = 5)
A549−GM130−SQSTM1 (n_incl_in_auc = 5, n_tested = 5)
A549−TP53BP1−CLTA (n_incl_in_auc = 5, n_tested = 5)
A549−TUBA1B−RELA (n_incl_in_auc = 5, n_tested = 5)
HepG2−ACTB−RAB5A (n_incl_in_auc = 5, n_tested = 5)
HepG2−CANX−COX4 (n_incl_in_auc = 5, n_tested = 5)
HepG2−GM130−SQSTM1 (n_incl_in_auc = 5, n_tested = 5)
HepG2−TP53BP1−CLTA (n_incl_in_auc = 5, n_tested = 5)
HepG2−TUBA1B−RELA (n_incl_in_auc = 5, n_tested = 5)
WPMY1−ACTB−RAB5A (n_incl_in_auc = 3, n_tested = 5)
WPMY1−CANX−COX4 (n_incl_in_auc = 4, n_tested = 5)
WPMY1−GM130−SQSTM1 (n_incl_in_auc = 4, n_tested = 5)
WPMY1−TP53BP1−CLTA (n_incl_in_auc = 5, n_tested = 5)
WPMY1−TUBA1B−RELA (n_incl_in_auc = 3, n_tested = 5)
0.0
0.1
0.2
0.3
0.4
0.5
0.6
0.7
0.8
0.9
1.0
auc_max_over_concentr_median
FDR−adjusted p−value < 0.05
FALSE
TRUE
Batch
All Aggr.
Within Batch 1
Within Batch 2
cell_type
A549
HepG2
WPMY1

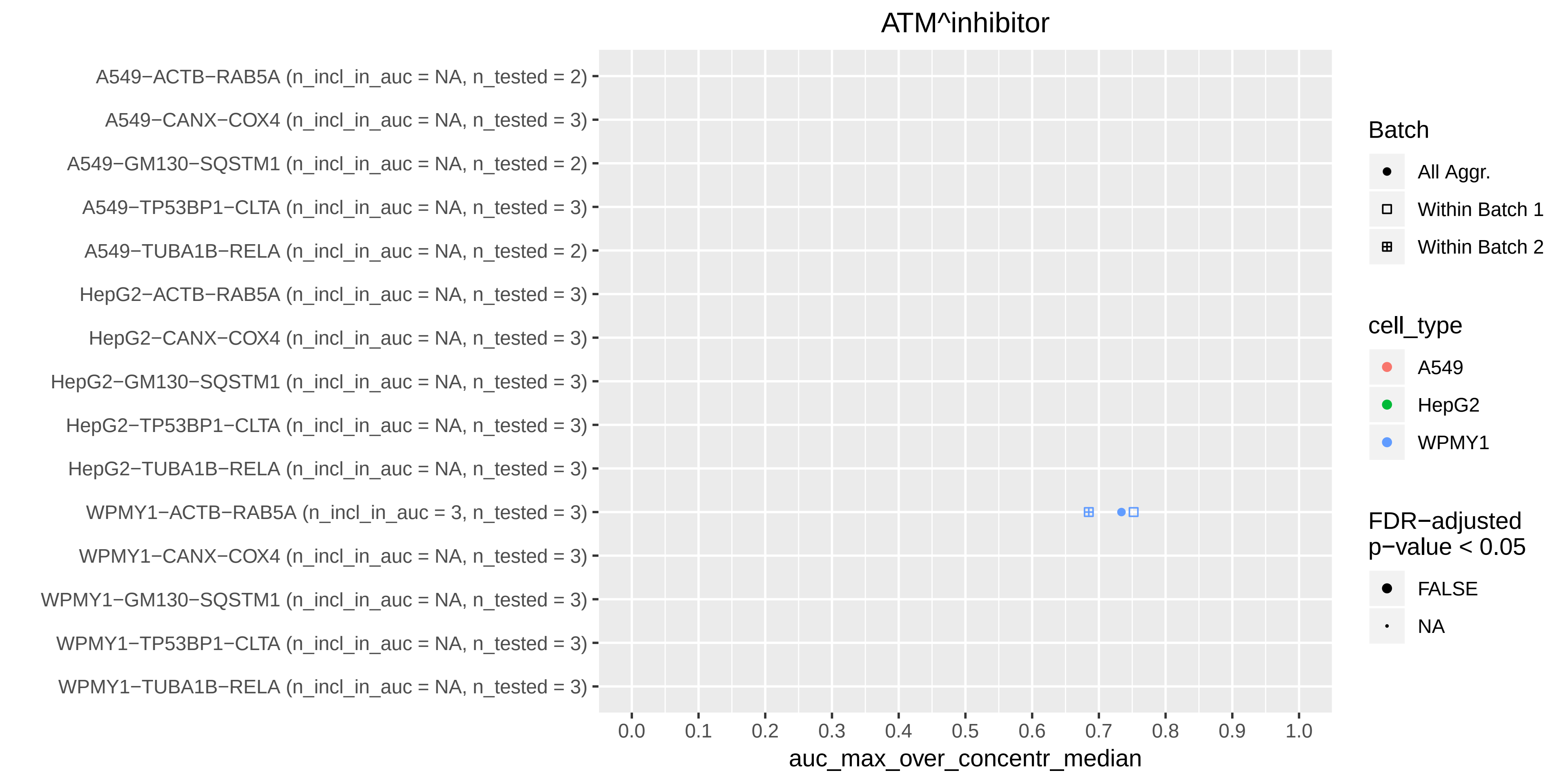

ATM^inhibitor
A549−ACTB−RAB5A (n_incl_in_auc = NA, n_tested = 2)
A549−CANX−COX4 (n_incl_in_auc = NA, n_tested = 3)
A549−GM130−SQSTM1 (n_incl_in_auc = NA, n_tested = 2)
A549−TP53BP1−CLTA (n_incl_in_auc = NA, n_tested = 3)
A549−TUBA1B−RELA (n_incl_in_auc = NA, n_tested = 2)
HepG2−ACTB−RAB5A (n_incl_in_auc = NA, n_tested = 3)
HepG2−CANX−COX4 (n_incl_in_auc = NA, n_tested = 3)
HepG2−GM130−SQSTM1 (n_incl_in_auc = NA, n_tested = 3)
HepG2−TP53BP1−CLTA (n_incl_in_auc = NA, n_tested = 3)
HepG2−TUBA1B−RELA (n_incl_in_auc = NA, n_tested = 3)
WPMY1−ACTB−RAB5A (n_incl_in_auc = 3, n_tested = 3)
WPMY1−CANX−COX4 (n_incl_in_auc = NA, n_tested = 3)
WPMY1−GM130−SQSTM1 (n_incl_in_auc = NA, n_tested = 3)
WPMY1−TP53BP1−CLTA (n_incl_in_auc = NA, n_tested = 3)
WPMY1−TUBA1B−RELA (n_incl_in_auc = NA, n_tested = 3)
0.0
0.1
0.2
0.3
0.4
0.5
0.6
0.7
0.8
0.9
1.0
auc_max_over_concentr_median
Batch
All Aggr.
Within Batch 1
Within Batch 2
cell_type
A549
HepG2
WPMY1
FDR−adjusted
p−value < 0.05
FALSE
NA

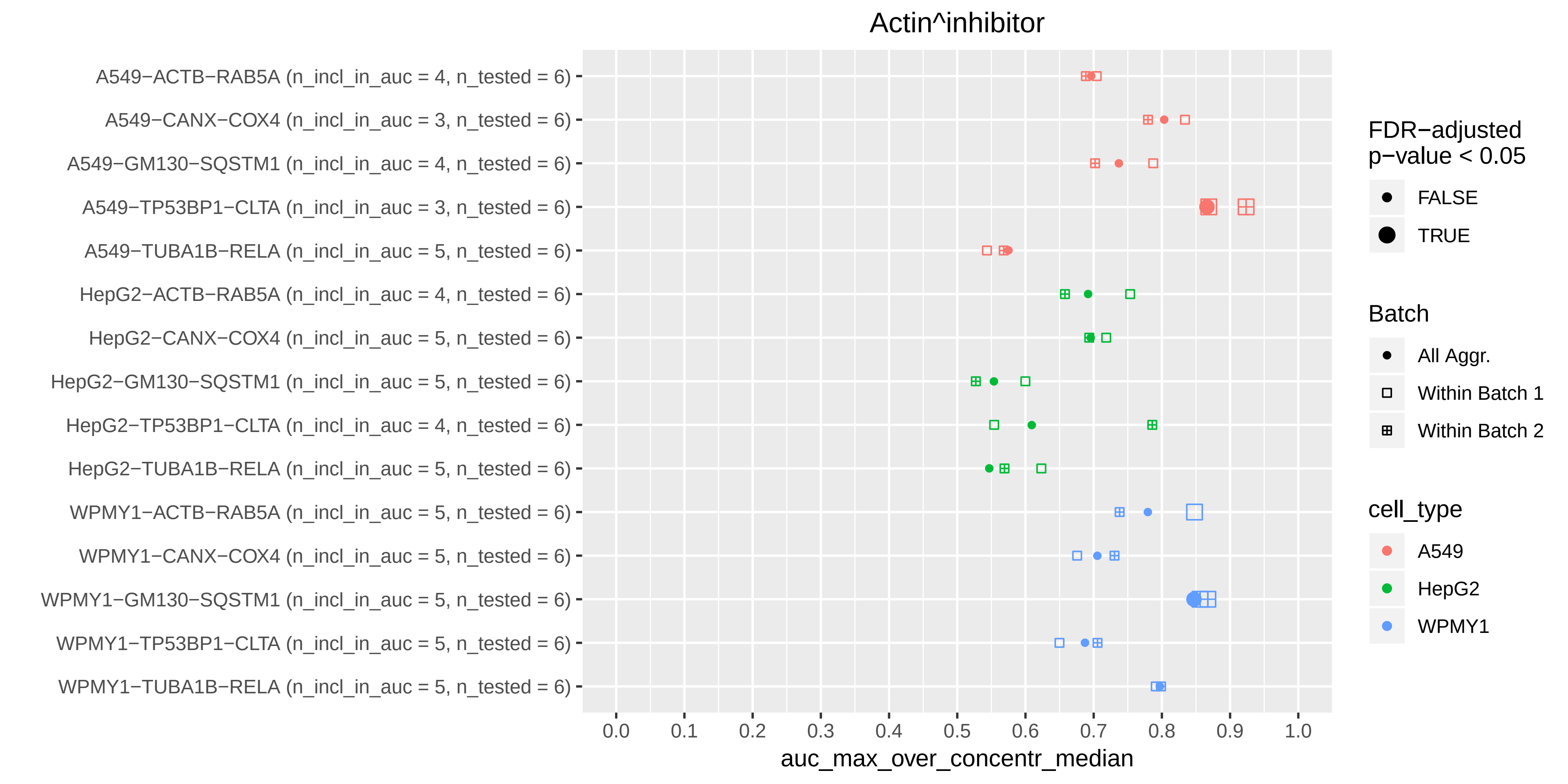
Actin^inhibitor
A549−ACTB−RAB5A (n_incl_in_auc = 4, n_tested = 6)
A549−CANX−COX4 (n_incl_in_auc = 3, n_tested = 6)
A549−GM130−SQSTM1 (n_incl_in_auc = 4, n_tested = 6)
A549−TP53BP1−CLTA (n_incl_in_auc = 3, n_tested = 6)
A549−TUBA1B−RELA (n_incl_in_auc = 5, n_tested = 6)
HepG2−ACTB−RAB5A (n_incl_in_auc = 4, n_tested = 6)
HepG2−CANX−COX4 (n_incl_in_auc = 5, n_tested = 6)
HepG2−GM130−SQSTM1 (n_incl_in_auc = 5, n_tested = 6)
HepG2−TP53BP1−CLTA (n_incl_in_auc = 4, n_tested = 6)
HepG2−TUBA1B−RELA (n_incl_in_auc = 5, n_tested = 6)
WPMY1−ACTB−RAB5A (n_incl_in_auc = 5, n_tested = 6)
WPMY1−CANX−COX4 (n_incl_in_auc = 5, n_tested = 6)
WPMY1−GM130−SQSTM1 (n_incl_in_auc = 5, n_tested = 6)
WPMY1−TP53BP1−CLTA (n_incl_in_auc = 5, n_tested = 6)
WPMY1−TUBA1B−RELA (n_incl_in_auc = 5, n_tested = 6)
0.0
0.1
0.2
0.3
0.4
0.5
0.6
0.7
0.8
0.9
1.0
auc_max_over_concentr_median
FDR−adjusted p−value < 0.05
FALSE
TRUE
Batch
All Aggr.
Within Batch 1
Within Batch 2
cell_type
A549
HepG2
WPMY1

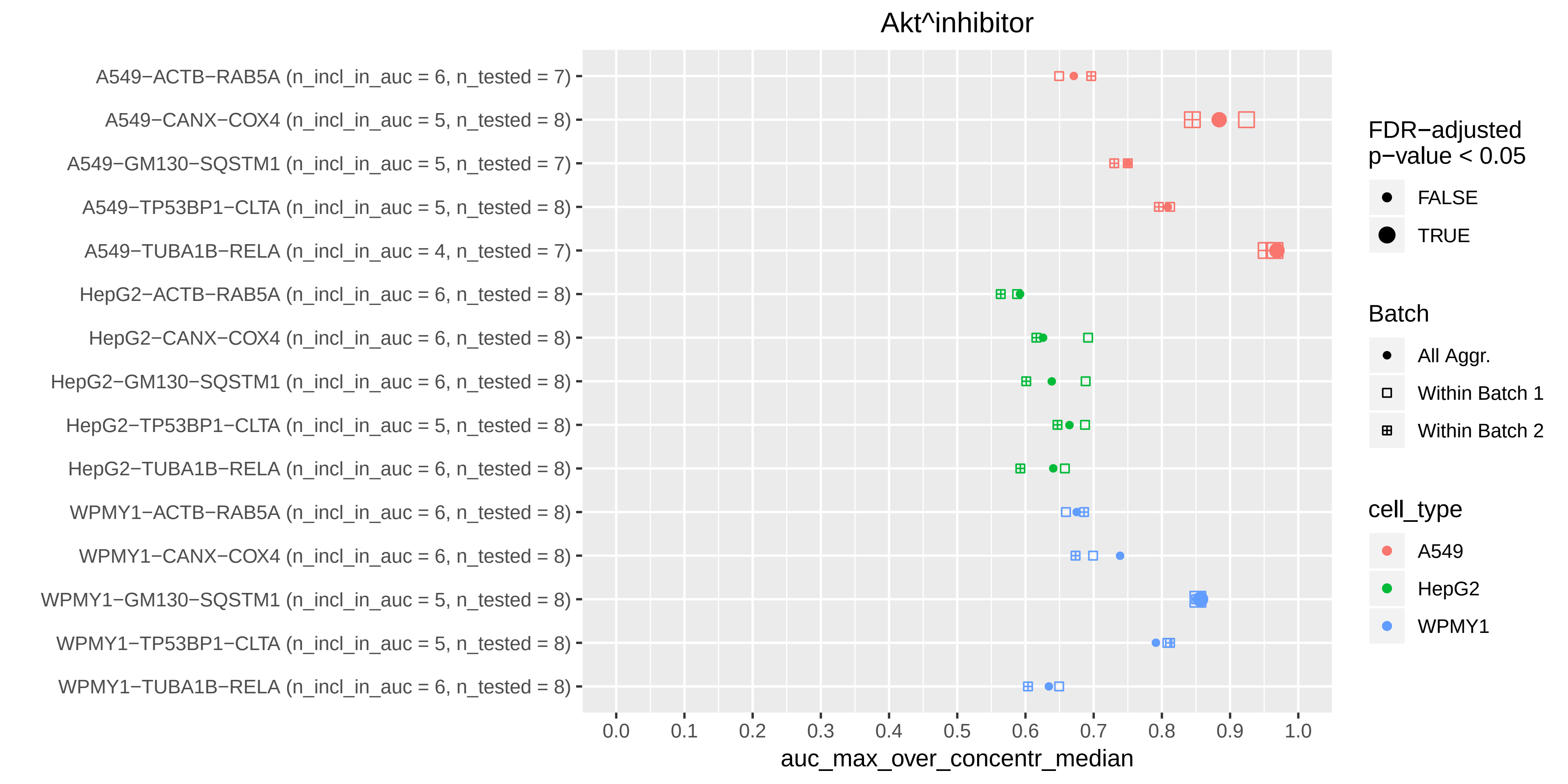

Akt^inhibitor
A549−ACTB−RAB5A (n_incl_in_auc = 6, n_tested = 7)
A549−CANX−COX4 (n_incl_in_auc = 5, n_tested = 8)
A549−GM130−SQSTM1 (n_incl_in_auc = 5, n_tested = 7)
A549−TP53BP1−CLTA (n_incl_in_auc = 5, n_tested = 8)
A549−TUBA1B−RELA (n_incl_in_auc = 4, n_tested = 7)
HepG2−ACTB−RAB5A (n_incl_in_auc = 6, n_tested = 8)
HepG2−CANX−COX4 (n_incl_in_auc = 6, n_tested = 8)
HepG2−GM130−SQSTM1 (n_incl_in_auc = 6, n_tested = 8)
HepG2−TP53BP1−CLTA (n_incl_in_auc = 5, n_tested = 8)
HepG2−TUBA1B−RELA (n_incl_in_auc = 6, n_tested = 8)
WPMY1−ACTB−RAB5A (n_incl_in_auc = 6, n_tested = 8)
WPMY1−CANX−COX4 (n_incl_in_auc = 6, n_tested = 8)
WPMY1−GM130−SQSTM1 (n_incl_in_auc = 5, n_tested = 8)
WPMY1−TP53BP1−CLTA (n_incl_in_auc = 5, n_tested = 8)
WPMY1−TUBA1B−RELA (n_incl_in_auc = 6, n_tested = 8)
0.0
0.1
0.2
0.3
0.4
0.5
0.6
0.7
0.8
0.9
1.0
auc_max_over_concentr_median
FDR−adjusted p−value < 0.05
FALSE
TRUE
Batch
All Aggr.
Within Batch 1
Within Batch 2
cell_type
A549
HepG2
WPMY1

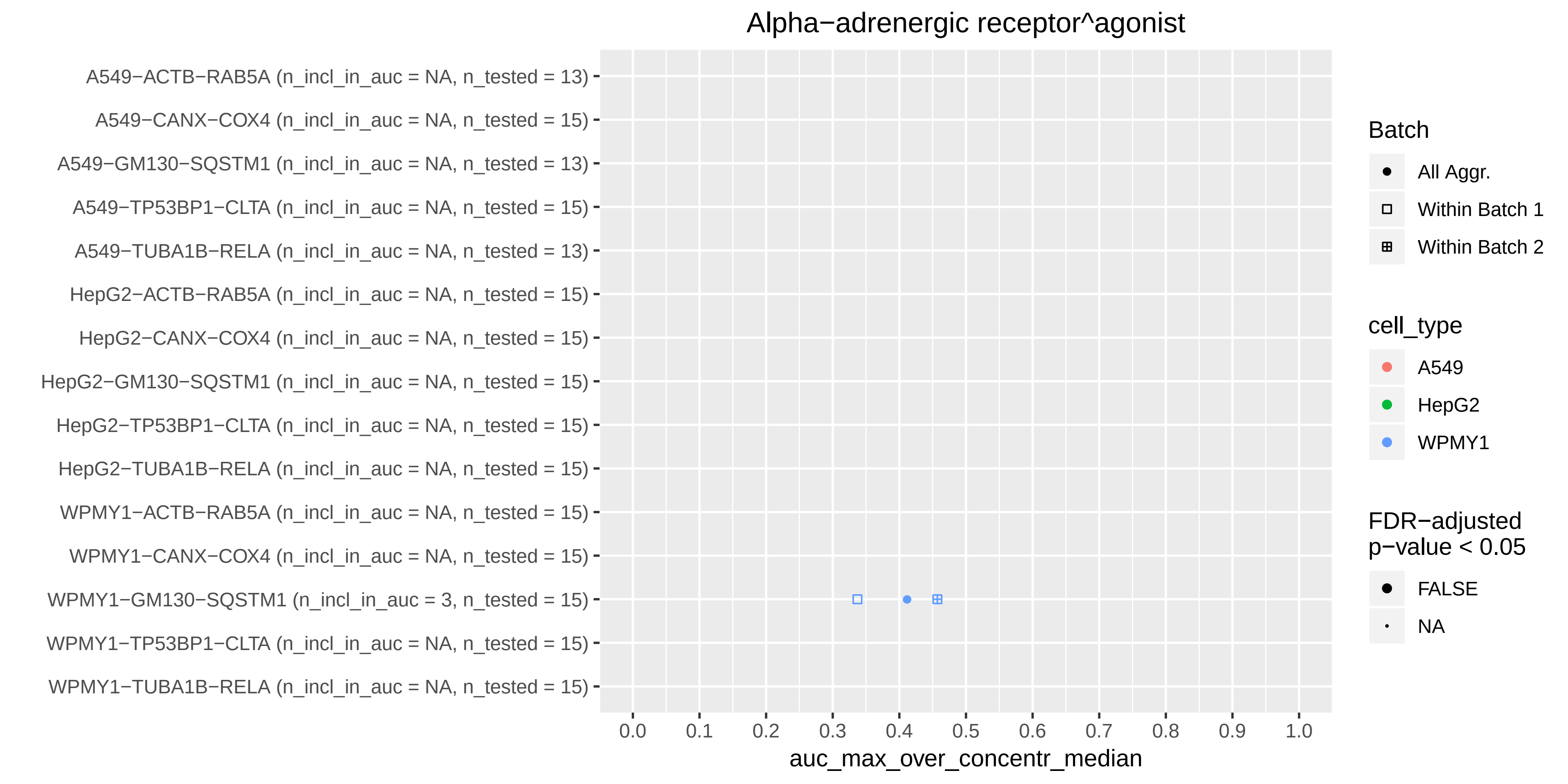

Alpha−adrenergic receptor^agonist
A549−ACTB−RAB5A (n_incl_in_auc = NA, n_tested = 13)
A549−CANX−COX4 (n_incl_in_auc = NA, n_tested = 15)
A549−GM130−SQSTM1 (n_incl_in_auc = NA, n_tested = 13)
A549−TP53BP1−CLTA (n_incl_in_auc = NA, n_tested = 15)
A549−TUBA1B−RELA (n_incl_in_auc = NA, n_tested = 13)
HepG2−ACTB−RAB5A (n_incl_in_auc = NA, n_tested = 15)
HepG2−CANX−COX4 (n_incl_in_auc = NA, n_tested = 15)
HepG2−GM130−SQSTM1 (n_incl_in_auc = NA, n_tested = 15)
HepG2−TP53BP1−CLTA (n_incl_in_auc = NA, n_tested = 15)
HepG2−TUBA1B−RELA (n_incl_in_auc = NA, n_tested = 15)
WPMY1−ACTB−RAB5A (n_incl_in_auc = NA, n_tested = 15)
WPMY1−CANX−COX4 (n_incl_in_auc = NA, n_tested = 15)
WPMY1−GM130−SQSTM1 (n_incl_in_auc = 3, n_tested = 15)
WPMY1−TP53BP1−CLTA (n_incl_in_auc = NA, n_tested = 15)
WPMY1−TUBA1B−RELA (n_incl_in_auc = NA, n_tested = 15)
0.0
0.1
0.2
0.3
0.4
0.5
0.6
0.7
0.8
0.9
1.0
auc_max_over_concentr_median
Batch
All Aggr.
Within Batch 1
Within Batch 2
cell_type
A549
HepG2
WPMY1
FDR−adjusted
p−value < 0.05
FALSE
NA

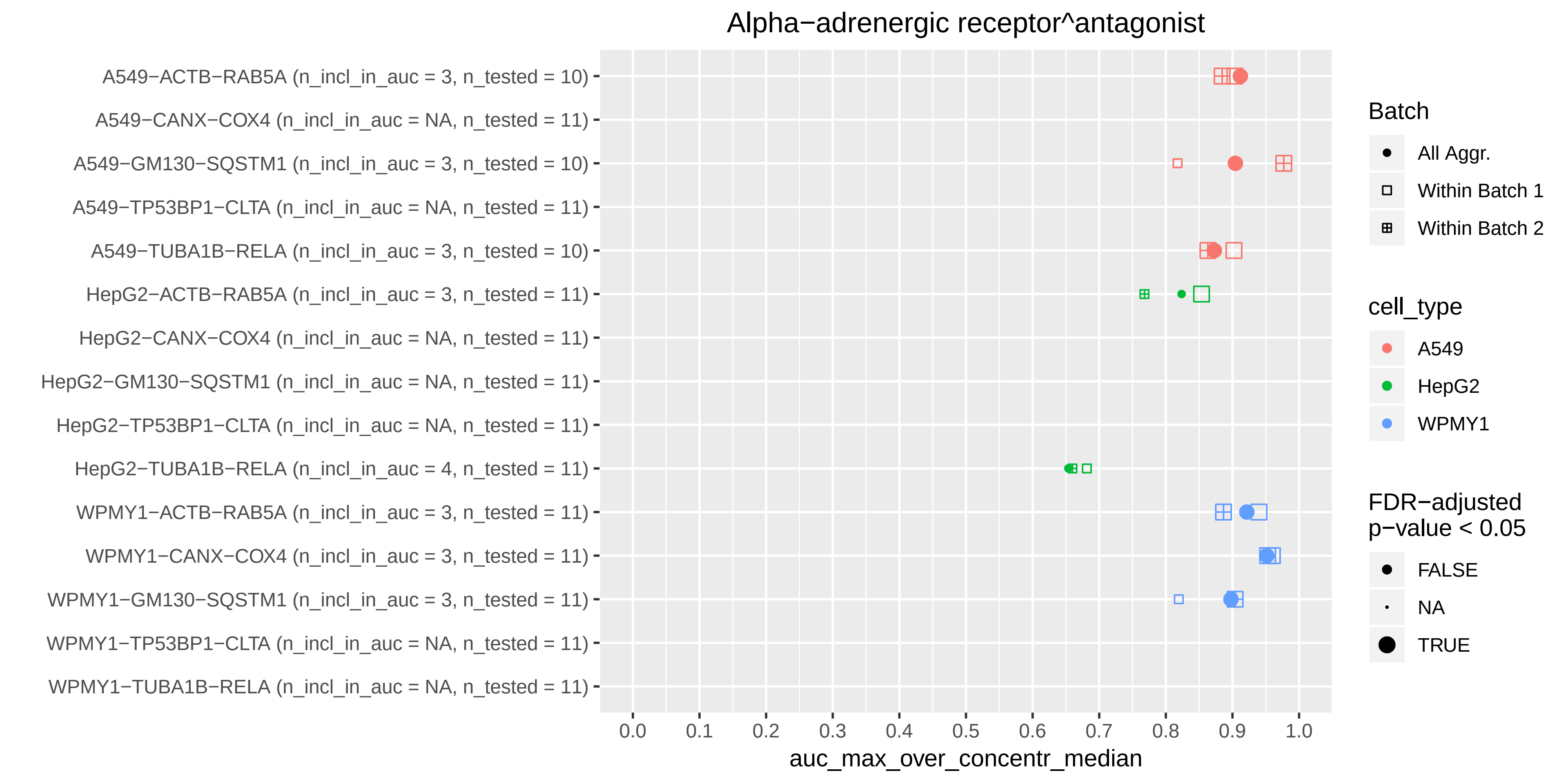
Alpha−adrenergic receptor^antagonist
A549−ACTB−RAB5A (n_incl_in_auc = 3, n_tested = 10)
A549−CANX−COX4 (n_incl_in_auc = NA, n_tested = 11)
A549−GM130−SQSTM1 (n_incl_in_auc = 3, n_tested = 10)
A549−TP53BP1−CLTA (n_incl_in_auc = NA, n_tested = 11)
A549−TUBA1B−RELA (n_incl_in_auc = 3, n_tested = 10)
HepG2−ACTB−RAB5A (n_incl_in_auc = 3, n_tested = 11)
HepG2−CANX−COX4 (n_incl_in_auc = NA, n_tested = 11)
HepG2−GM130−SQSTM1 (n_incl_in_auc = NA, n_tested = 11)
HepG2−TP53BP1−CLTA (n_incl_in_auc = NA, n_tested = 11)
HepG2−TUBA1B−RELA (n_incl_in_auc = 4, n_tested = 11)
WPMY1−ACTB−RAB5A (n_incl_in_auc = 3, n_tested = 11)
WPMY1−CANX−COX4 (n_incl_in_auc = 3, n_tested = 11)
WPMY1−GM130−SQSTM1 (n_incl_in_auc = 3, n_tested = 11)
WPMY1−TP53BP1−CLTA (n_incl_in_auc = NA, n_tested = 11)
WPMY1−TUBA1B−RELA (n_incl_in_auc = NA, n_tested = 11)
0.0
0.1
0.2
0.3
0.4
0.5
0.6
0.7
0.8
0.9
1.0
auc_max_over_concentr_median
Batch
All Aggr.
Within Batch 1
Within Batch 2
cell_type
A549
HepG2
WPMY1
FDR−adjusted
p−value < 0.05
FALSE
NA
TRUE

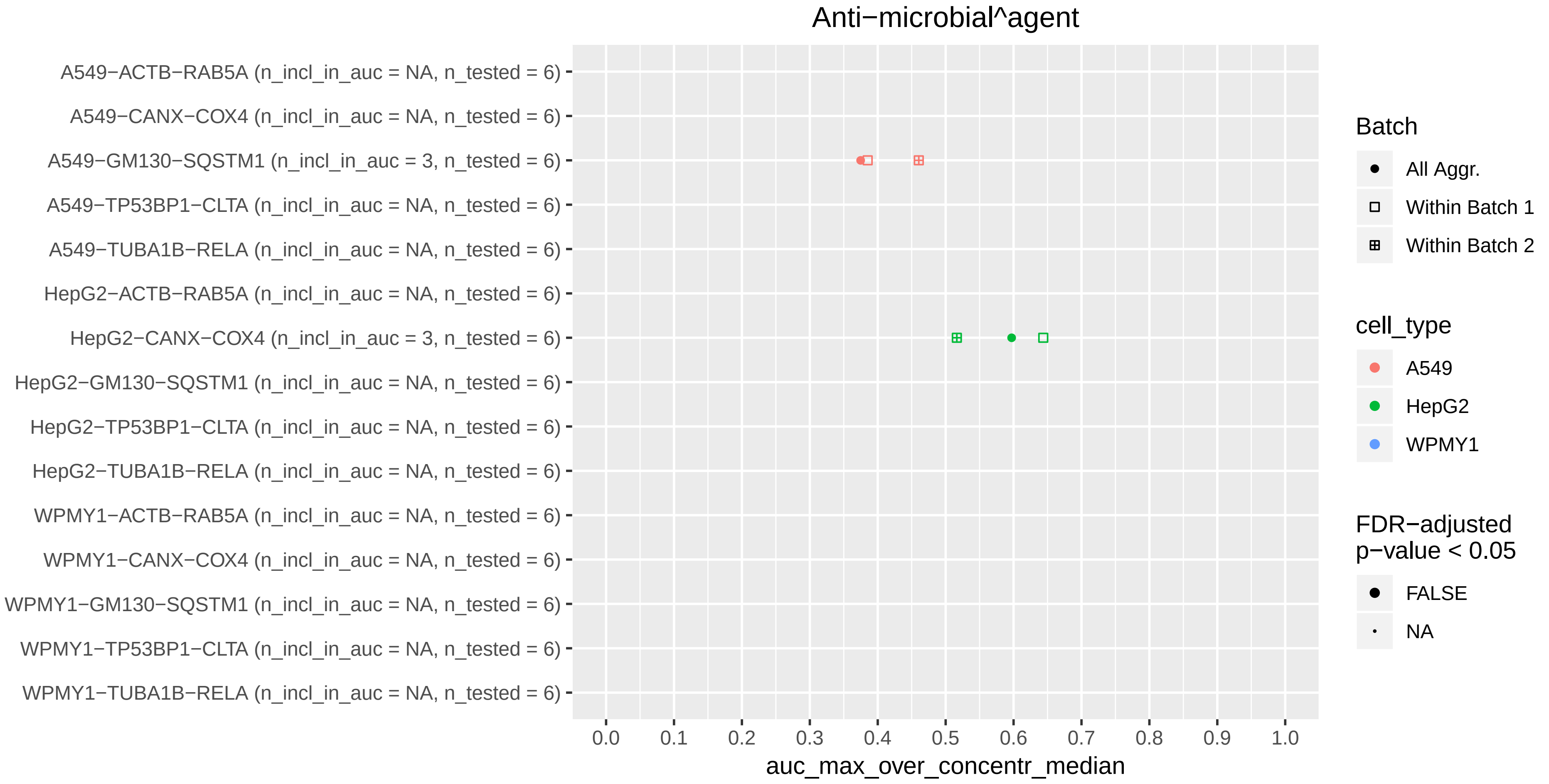
Anti−microbial^agent
A549−ACTB−RAB5A (n_incl_in_auc = NA, n_tested = 6)
A549−CANX−COX4 (n_incl_in_auc = NA, n_tested = 6)
A549−GM130−SQSTM1 (n_incl_in_auc = 3, n_tested = 6)
A549−TP53BP1−CLTA (n_incl_in_auc = NA, n_tested = 6)
A549−TUBA1B−RELA (n_incl_in_auc = NA, n_tested = 6)
HepG2−ACTB−RAB5A (n_incl_in_auc = NA, n_tested = 6)
HepG2−CANX−COX4 (n_incl_in_auc = 3, n_tested = 6)
HepG2−GM130−SQSTM1 (n_incl_in_auc = NA, n_tested = 6)
HepG2−TP53BP1−CLTA (n_incl_in_auc = NA, n_tested = 6)
HepG2−TUBA1B−RELA (n_incl_in_auc = NA, n_tested = 6)
WPMY1−ACTB−RAB5A (n_incl_in_auc = NA, n_tested = 6)
WPMY1−CANX−COX4 (n_incl_in_auc = NA, n_tested = 6)
WPMY1−GM130−SQSTM1 (n_incl_in_auc = NA, n_tested = 6)
WPMY1−TP53BP1−CLTA (n_incl_in_auc = NA, n_tested = 6)
WPMY1−TUBA1B−RELA (n_incl_in_auc = NA, n_tested = 6)
0.0 0.1 0.2 0.3 0.4 0.5 0.6 0.7 0.8 0.9 1.0
auc_max_over_concentr_median
Batch
All Aggr.
Within Batch 1
Within Batch 2
cell_type
A549
HepG2
WPMY1
FDR−adjusted p−value < 0.05
FALSE
NA

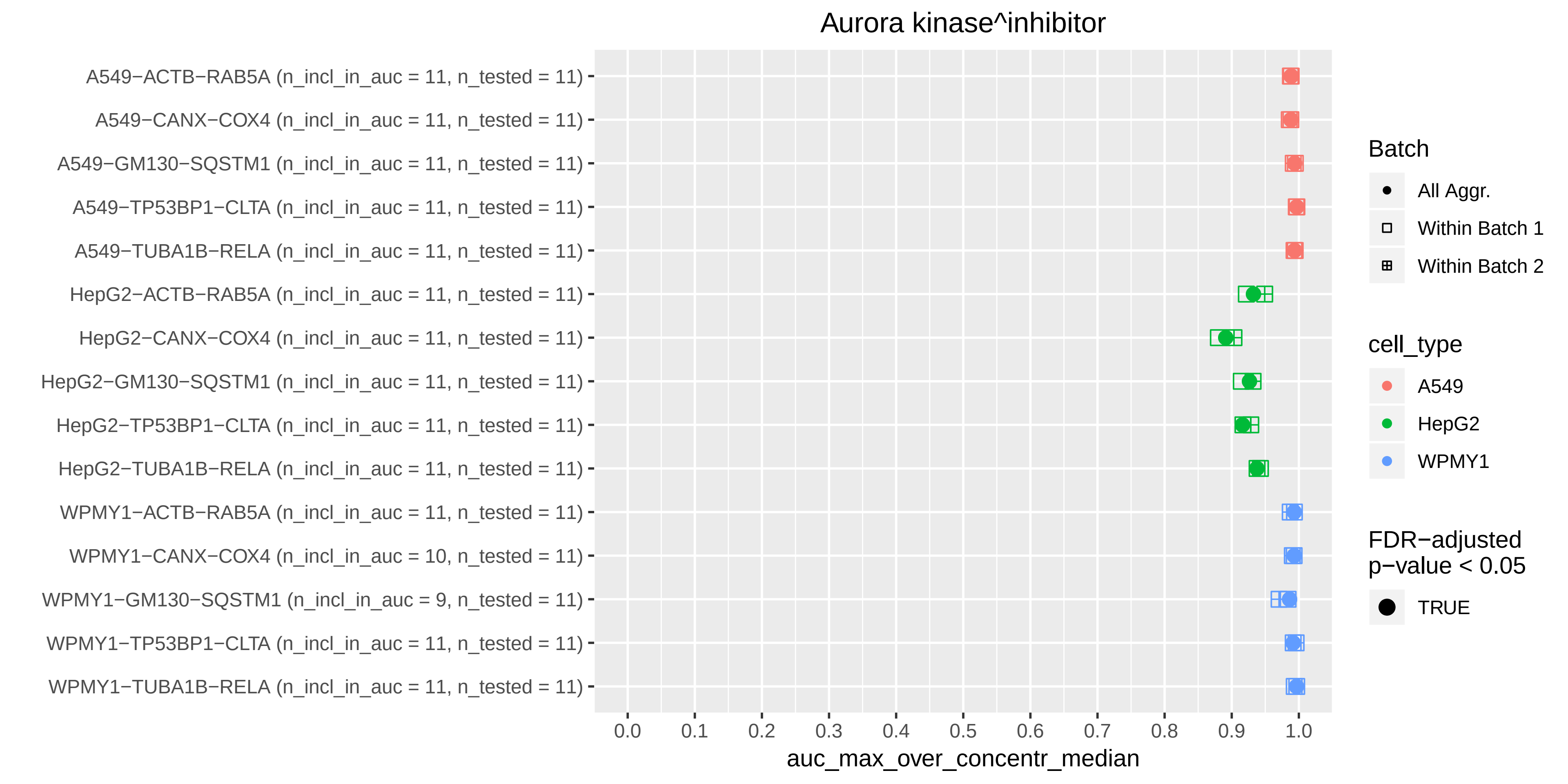

Aurora kinase^inhibitor
A549−ACTB−RAB5A (n_incl_in_auc = 11, n_tested = 11)
A549−CANX−COX4 (n_incl_in_auc = 11, n_tested = 11)
A549−GM130−SQSTM1 (n_incl_in_auc = 11, n_tested = 11)
A549−TP53BP1−CLTA (n_incl_in_auc = 11, n_tested = 11)
A549−TUBA1B−RELA (n_incl_in_auc = 11, n_tested = 11)
HepG2−ACTB−RAB5A (n_incl_in_auc = 11, n_tested = 11)
HepG2−CANX−COX4 (n_incl_in_auc = 11, n_tested = 11)
HepG2−GM130−SQSTM1 (n_incl_in_auc = 11, n_tested = 11)
HepG2−TP53BP1−CLTA (n_incl_in_auc = 11, n_tested = 11)
HepG2−TUBA1B−RELA (n_incl_in_auc = 11, n_tested = 11)
WPMY1−ACTB−RAB5A (n_incl_in_auc = 11, n_tested = 11)
WPMY1−CANX−COX4 (n_incl_in_auc = 10, n_tested = 11)
WPMY1−GM130−SQSTM1 (n_incl_in_auc = 9, n_tested = 11)
WPMY1−TP53BP1−CLTA (n_incl_in_auc = 11, n_tested = 11)
WPMY1−TUBA1B−RELA (n_incl_in_auc = 11, n_tested = 11)
0.0
0.1
0.2
0.3
0.4
0.5
0.6
0.7
0.8
0.9
1.0
auc_max_over_concentr_median
Batch
All Aggr.
Within Batch 1
Within Batch 2
cell_type
A549
HepG2
WPMY1
FDR−adjusted p−value < 0.05
TRUE

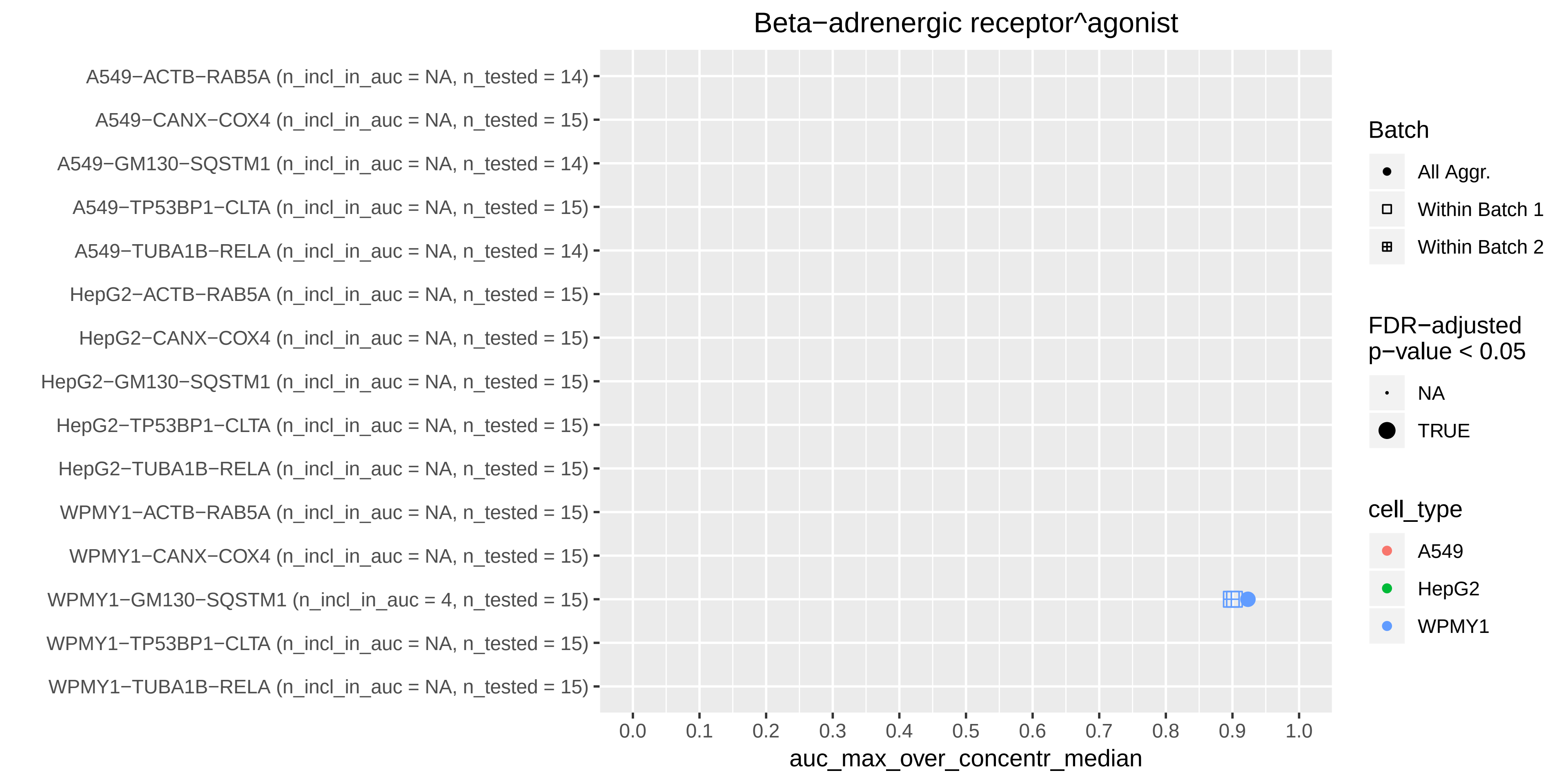

Beta−adrenergic receptor^agonist
A549−ACTB−RAB5A (n_incl_in_auc = NA, n_tested = 14)
A549−CANX−COX4 (n_incl_in_auc = NA, n_tested = 15)
A549−GM130−SQSTM1 (n_incl_in_auc = NA, n_tested = 14)
A549−TP53BP1−CLTA (n_incl_in_auc = NA, n_tested = 15)
A549−TUBA1B−RELA (n_incl_in_auc = NA, n_tested = 14)
HepG2−ACTB−RAB5A (n_incl_in_auc = NA, n_tested = 15)
HepG2−CANX−COX4 (n_incl_in_auc = NA, n_tested = 15)
HepG2−GM130−SQSTM1 (n_incl_in_auc = NA, n_tested = 15)
HepG2−TP53BP1−CLTA (n_incl_in_auc = NA, n_tested = 15)
HepG2−TUBA1B−RELA (n_incl_in_auc = NA, n_tested = 15)
WPMY1−ACTB−RAB5A (n_incl_in_auc = NA, n_tested = 15)
WPMY1−CANX−COX4 (n_incl_in_auc = NA, n_tested = 15)
WPMY1−GM130−SQSTM1 (n_incl_in_auc = 4, n_tested = 15)
WPMY1−TP53BP1−CLTA (n_incl_in_auc = NA, n_tested = 15)
WPMY1−TUBA1B−RELA (n_incl_in_auc = NA, n_tested = 15)
0.0
0.1
0.2
0.3
0.4
0.5
0.6
0.7
0.8
0.9
1.0
auc_max_over_concentr_median
Batch
All Aggr.
Within Batch 1
Within Batch 2
FDR−adjusted p−value < 0.05
NA
TRUE
cell_type
A549
HepG2
WPMY1

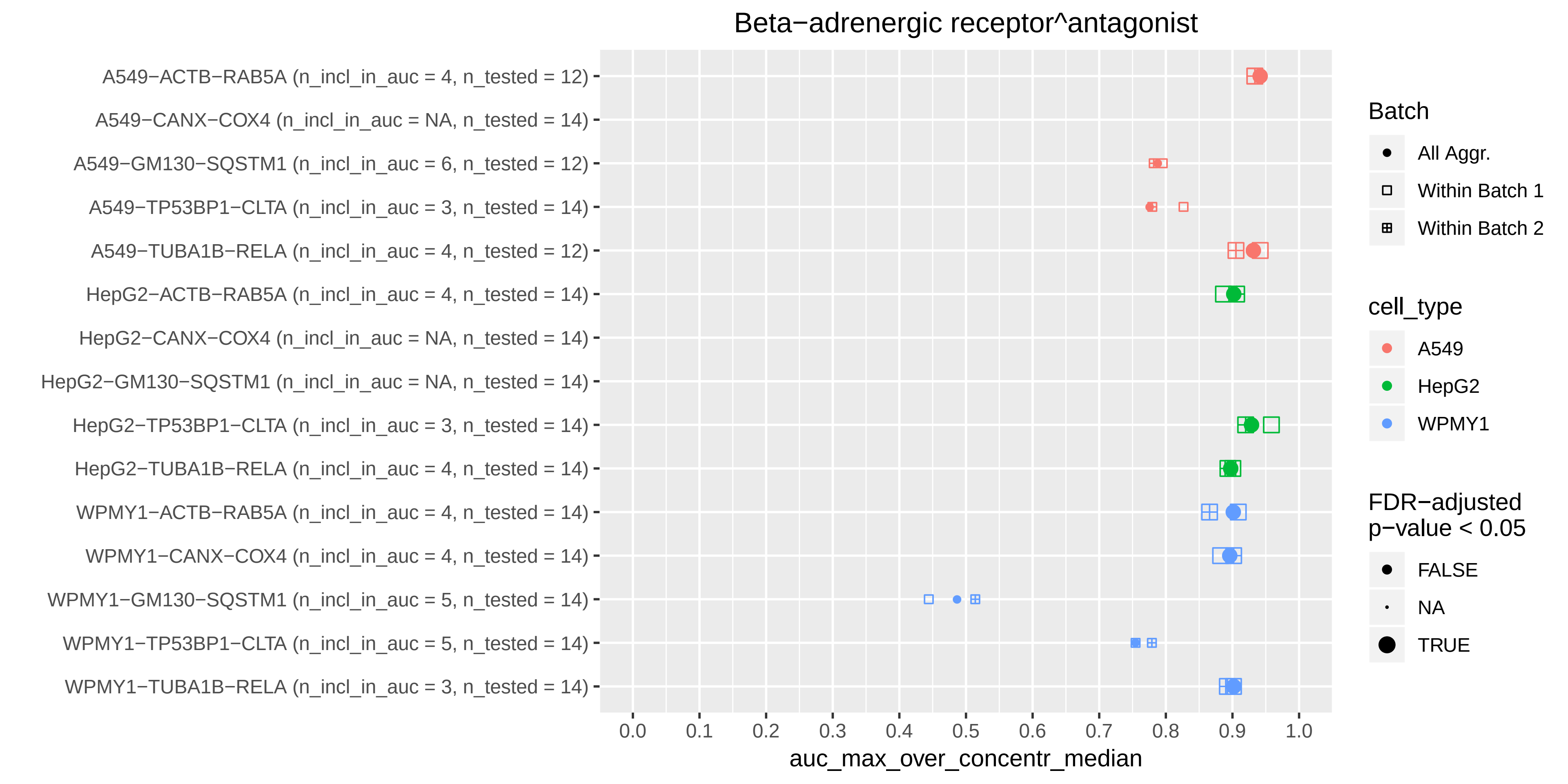
Beta−adrenergic receptor^antagonist
A549−ACTB−RAB5A (n_incl_in_auc = 4, n_tested = 12)
A549−CANX−COX4 (n_incl_in_auc = NA, n_tested = 14)
A549−GM130−SQSTM1 (n_incl_in_auc = 6, n_tested = 12)
A549−TP53BP1−CLTA (n_incl_in_auc = 3, n_tested = 14)
A549−TUBA1B−RELA (n_incl_in_auc = 4, n_tested = 12)
HepG2−ACTB−RAB5A (n_incl_in_auc = 4, n_tested = 14)
HepG2−CANX−COX4 (n_incl_in_auc = NA, n_tested = 14)
HepG2−GM130−SQSTM1 (n_incl_in_auc = NA, n_tested = 14)
HepG2−TP53BP1−CLTA (n_incl_in_auc = 3, n_tested = 14)
HepG2−TUBA1B−RELA (n_incl_in_auc = 4, n_tested = 14)
WPMY1−ACTB−RAB5A (n_incl_in_auc = 4, n_tested = 14)
WPMY1−CANX−COX4 (n_incl_in_auc = 4, n_tested = 14)
WPMY1−GM130−SQSTM1 (n_incl_in_auc = 5, n_tested = 14)
WPMY1−TP53BP1−CLTA (n_incl_in_auc = 5, n_tested = 14)
WPMY1−TUBA1B−RELA (n_incl_in_auc = 3, n_tested = 14)
0.0 0.1 0.2 0.3 0.4 0.5 0.6 0.7 0.8 0.9 1.0
auc_max_over_concentr_median
Batch
All Aggr.
Within Batch 1
Within Batch 2
cell_type
A549
HepG2
WPMY1
FDR−adjusted
p−value < 0.05
FALSE
NA
TRUE

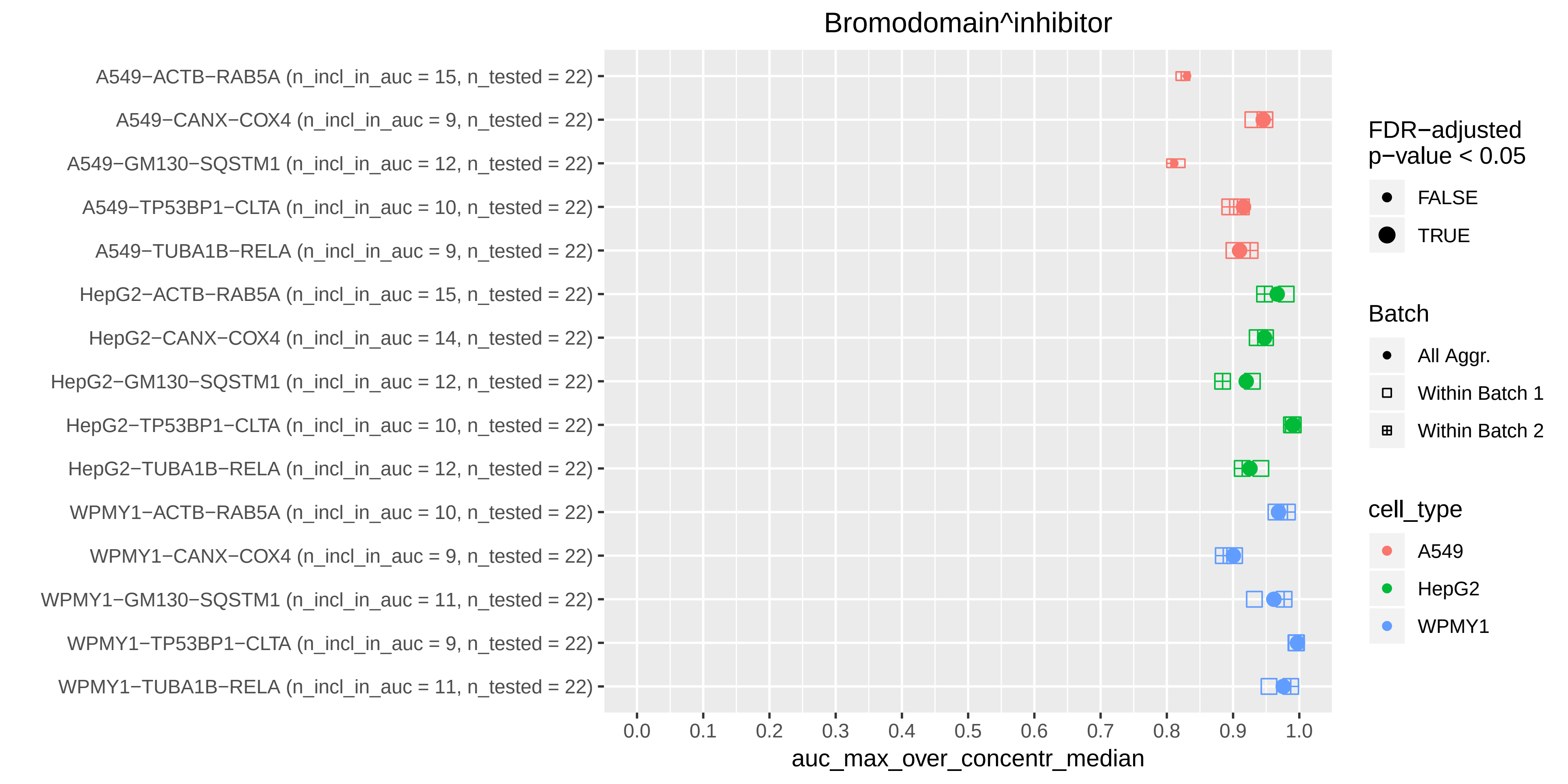

Bromodomain^inhibitor
A549−ACTB−RAB5A (n_incl_in_auc = 15, n_tested = 22)
A549−CANX−COX4 (n_incl_in_auc = 9, n_tested = 22)
A549−GM130−SQSTM1 (n_incl_in_auc = 12, n_tested = 22)
A549−TP53BP1−CLTA (n_incl_in_auc = 10, n_tested = 22)
A549−TUBA1B−RELA (n_incl_in_auc = 9, n_tested = 22)
HepG2−ACTB−RAB5A (n_incl_in_auc = 15, n_tested = 22)
HepG2−CANX−COX4 (n_incl_in_auc = 14, n_tested = 22)
HepG2−GM130−SQSTM1 (n_incl_in_auc = 12, n_tested = 22)
HepG2−TP53BP1−CLTA (n_incl_in_auc = 10, n_tested = 22)
HepG2−TUBA1B−RELA (n_incl_in_auc = 12, n_tested = 22)
WPMY1−ACTB−RAB5A (n_incl_in_auc = 10, n_tested = 22)
WPMY1−CANX−COX4 (n_incl_in_auc = 9, n_tested = 22)
WPMY1−GM130−SQSTM1 (n_incl_in_auc = 11, n_tested = 22)
WPMY1−TP53BP1−CLTA (n_incl_in_auc = 9, n_tested = 22)
WPMY1−TUBA1B−RELA (n_incl_in_auc = 11, n_tested = 22)
0.0
0.1
0.2
0.3
0.4
0.5
0.6
0.7
0.8
0.9
1.0
auc_max_over_concentr_median
FDR−adjusted p−value < 0.05
FALSE
TRUE
Batch
All Aggr.
Within Batch 1
Within Batch 2
cell_type
A549
HepG2
WPMY1

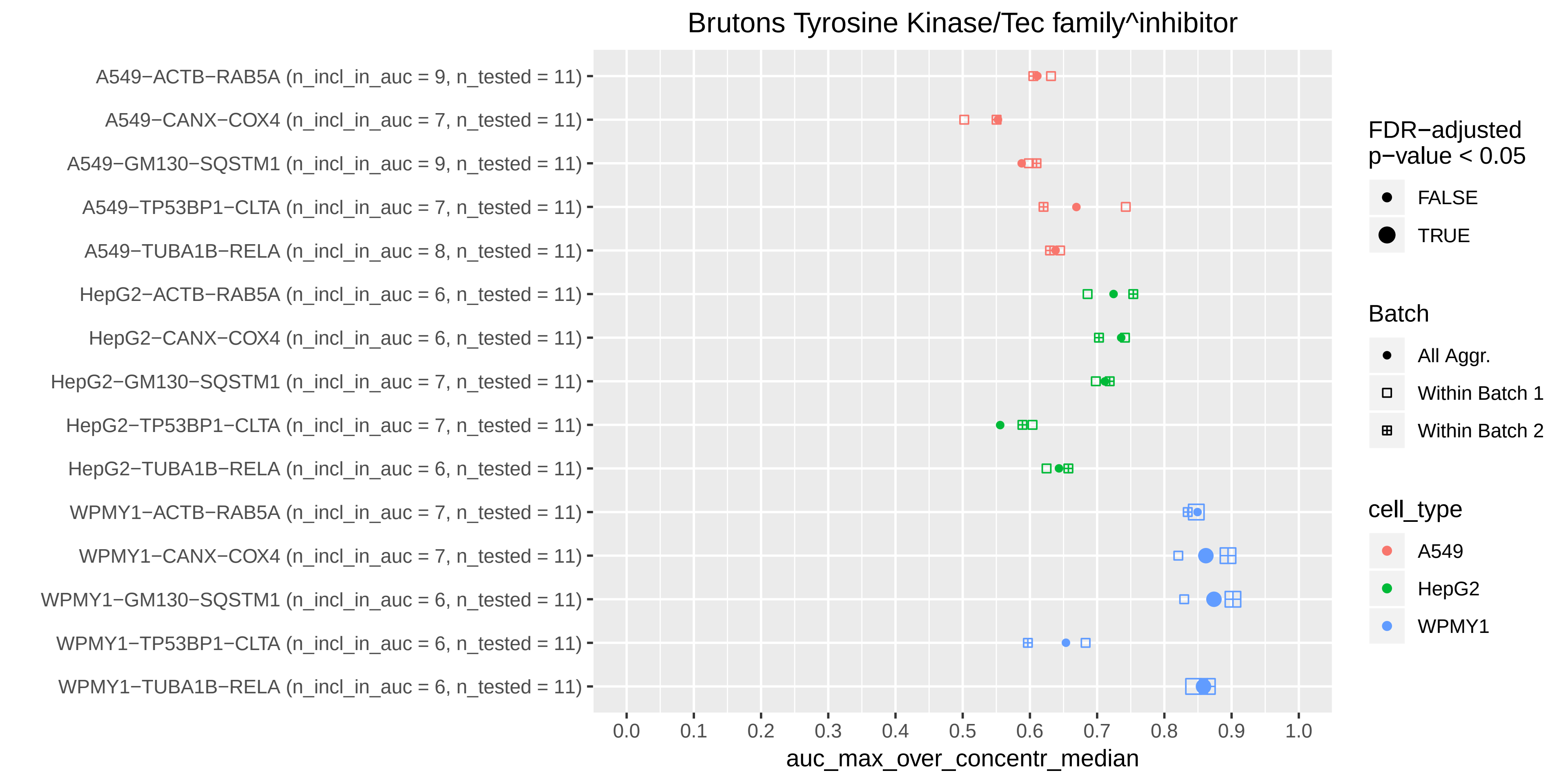

Brutons Tyrosine Kinase/Tec family^inhibitor
A549−ACTB−RAB5A (n_incl_in_auc = 9, n_tested = 11)
A549−CANX−COX4 (n_incl_in_auc = 7, n_tested = 11)
A549−GM130−SQSTM1 (n_incl_in_auc = 9, n_tested = 11)
A549−TP53BP1−CLTA (n_incl_in_auc = 7, n_tested = 11)
A549−TUBA1B−RELA (n_incl_in_auc = 8, n_tested = 11)
HepG2−ACTB−RAB5A (n_incl_in_auc = 6, n_tested = 11)
HepG2−CANX−COX4 (n_incl_in_auc = 6, n_tested = 11)
HepG2−GM130−SQSTM1 (n_incl_in_auc = 7, n_tested = 11)
HepG2−TP53BP1−CLTA (n_incl_in_auc = 7, n_tested = 11)
HepG2−TUBA1B−RELA (n_incl_in_auc = 6, n_tested = 11)
WPMY1−ACTB−RAB5A (n_incl_in_auc = 7, n_tested = 11)
WPMY1−CANX−COX4 (n_incl_in_auc = 7, n_tested = 11)
WPMY1−GM130−SQSTM1 (n_incl_in_auc = 6, n_tested = 11)
WPMY1−TP53BP1−CLTA (n_incl_in_auc = 6, n_tested = 11)
WPMY1−TUBA1B−RELA (n_incl_in_auc = 6, n_tested = 11)
0.0
0.1
0.2
0.3
0.4
0.5
0.6
0.7
0.8
0.9
1.0
auc_max_over_concentr_median
FDR−adjusted p−value < 0.05
FALSE
TRUE
Batch
All Aggr.
Within Batch 1
Within Batch 2
cell_type
A549
HepG2
WPMY1

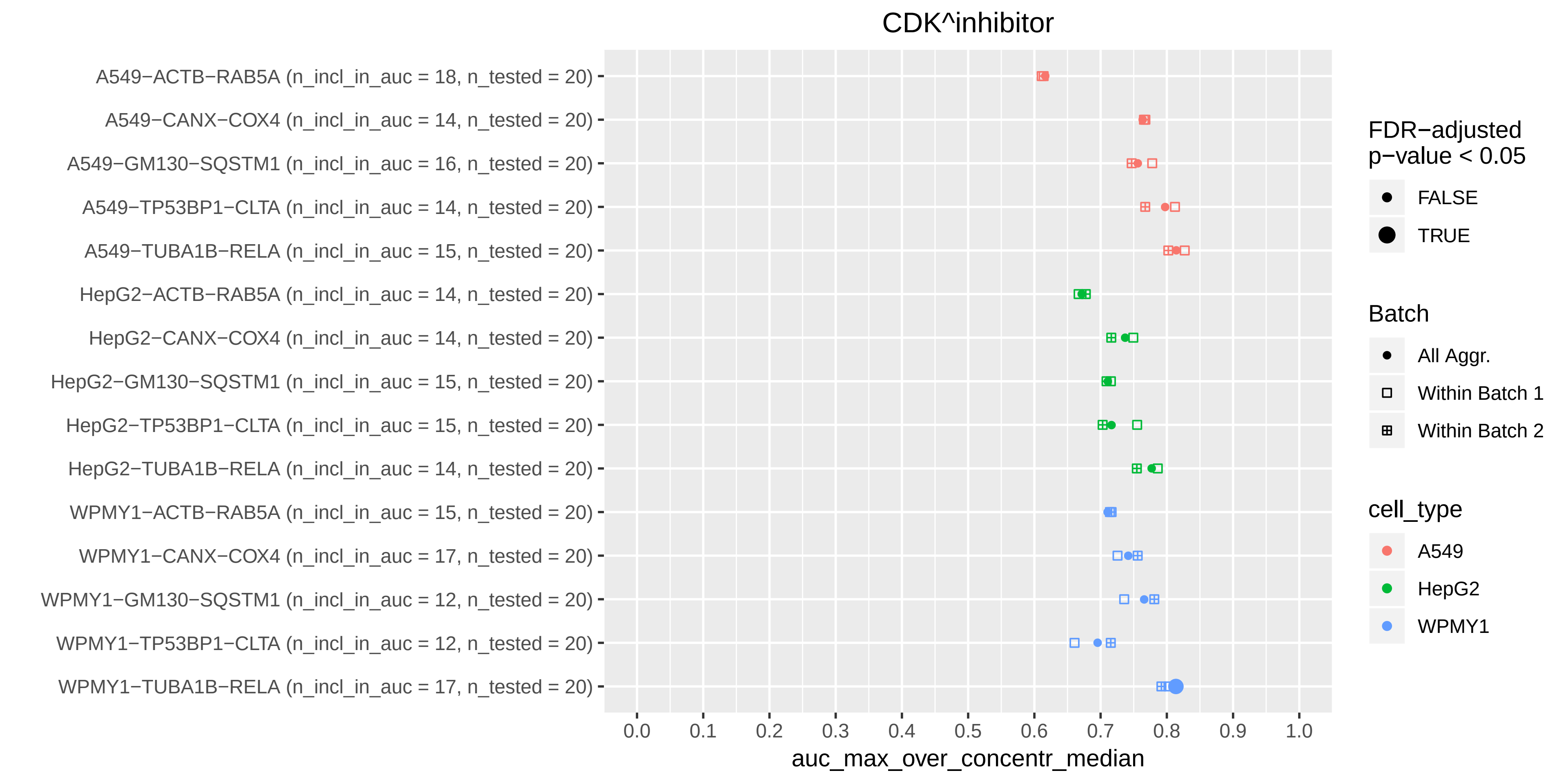

CDK^inhibitor
A549−ACTB−RAB5A (n_incl_in_auc = 18, n_tested = 20)
A549−CANX−COX4 (n_incl_in_auc = 14, n_tested = 20)
A549−GM130−SQSTM1 (n_incl_in_auc = 16, n_tested = 20)
A549−TP53BP1−CLTA (n_incl_in_auc = 14, n_tested = 20)
A549−TUBA1B−RELA (n_incl_in_auc = 15, n_tested = 20)
HepG2−ACTB−RAB5A (n_incl_in_auc = 14, n_tested = 20)
HepG2−CANX−COX4 (n_incl_in_auc = 14, n_tested = 20)
HepG2−GM130−SQSTM1 (n_incl_in_auc = 15, n_tested = 20)
HepG2−TP53BP1−CLTA (n_incl_in_auc = 15, n_tested = 20)
HepG2−TUBA1B−RELA (n_incl_in_auc = 14, n_tested = 20)
WPMY1−ACTB−RAB5A (n_incl_in_auc = 15, n_tested = 20)
WPMY1−CANX−COX4 (n_incl_in_auc = 17, n_tested = 20)
WPMY1−GM130−SQSTM1 (n_incl_in_auc = 12, n_tested = 20)
WPMY1−TP53BP1−CLTA (n_incl_in_auc = 12, n_tested = 20)
WPMY1−TUBA1B−RELA (n_incl_in_auc = 17, n_tested = 20)
0.0
0.1
0.2
0.3
0.4
0.5
0.6
0.7
0.8
0.9
1.0
auc_max_over_concentr_median
FDR−adjusted p−value < 0.05
FALSE
TRUE
Batch
All Aggr.
Within Batch 1
Within Batch 2
cell_type
A549
HepG2
WPMY1

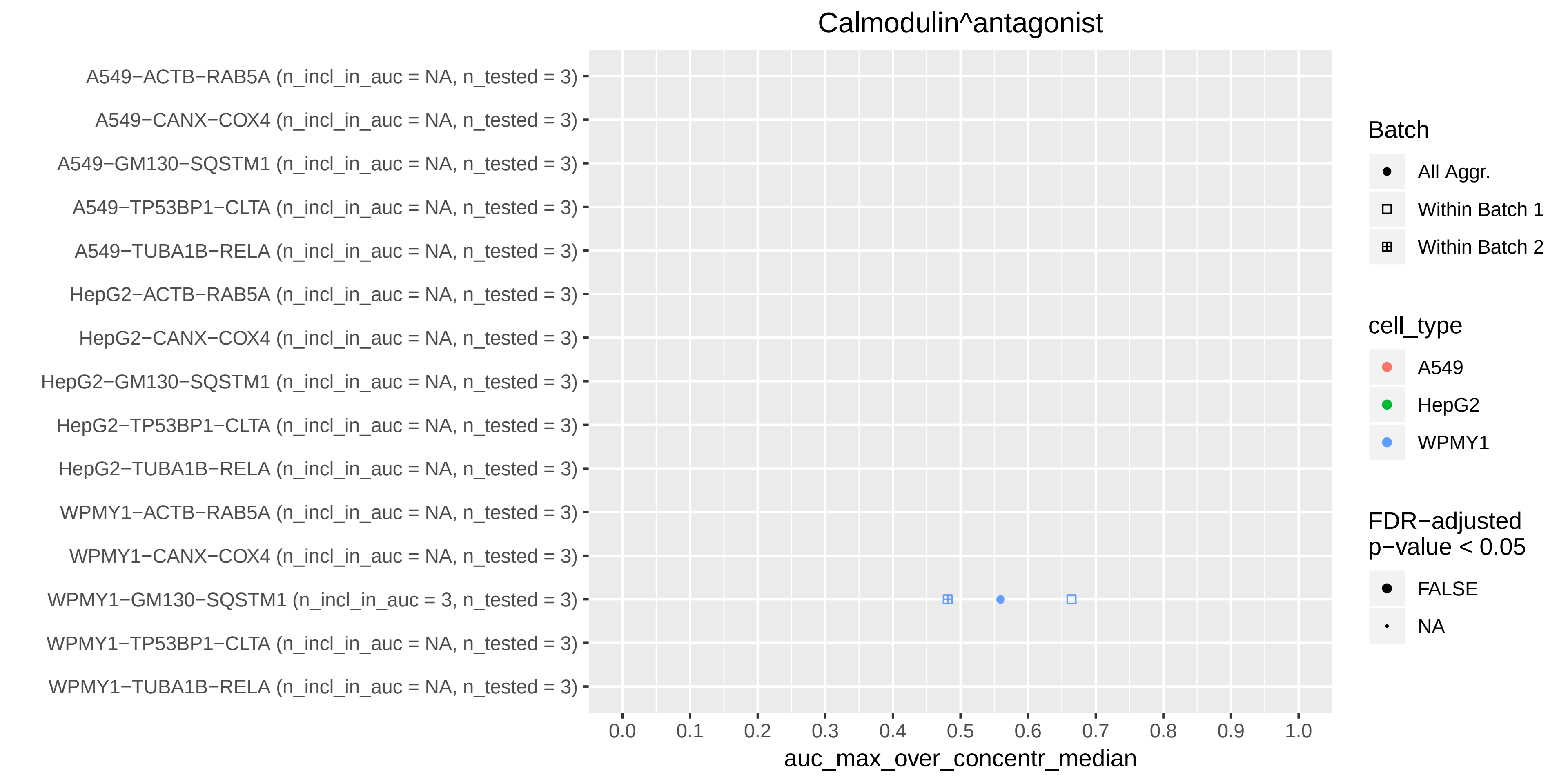

Calmodulin^antagonist
A549−ACTB−RAB5A (n_incl_in_auc = NA, n_tested = 3)
A549−CANX−COX4 (n_incl_in_auc = NA, n_tested = 3)
A549−GM130−SQSTM1 (n_incl_in_auc = NA, n_tested = 3)
A549−TP53BP1−CLTA (n_incl_in_auc = NA, n_tested = 3)
A549−TUBA1B−RELA (n_incl_in_auc = NA, n_tested = 3)
HepG2−ACTB−RAB5A (n_incl_in_auc = NA, n_tested = 3)
HepG2−CANX−COX4 (n_incl_in_auc = NA, n_tested = 3)
HepG2−GM130−SQSTM1 (n_incl_in_auc = NA, n_tested = 3)
HepG2−TP53BP1−CLTA (n_incl_in_auc = NA, n_tested = 3)
HepG2−TUBA1B−RELA (n_incl_in_auc = NA, n_tested = 3)
WPMY1−ACTB−RAB5A (n_incl_in_auc = NA, n_tested = 3)
WPMY1−CANX−COX4 (n_incl_in_auc = NA, n_tested = 3)
WPMY1−GM130−SQSTM1 (n_incl_in_auc = 3, n_tested = 3)
WPMY1−TP53BP1−CLTA (n_incl_in_auc = NA, n_tested = 3)
WPMY1−TUBA1B−RELA (n_incl_in_auc = NA, n_tested = 3)
0.0 0.1 0.2 0.3 0.4 0.5 0.6 0.7 0.8 0.9 1.0
auc_max_over_concentr_median
Batch
All Aggr.
Within Batch 1
Within Batch 2
cell_type
A549
HepG2
WPMY1
FDR−adjusted
p−value < 0.05
FALSE
NA

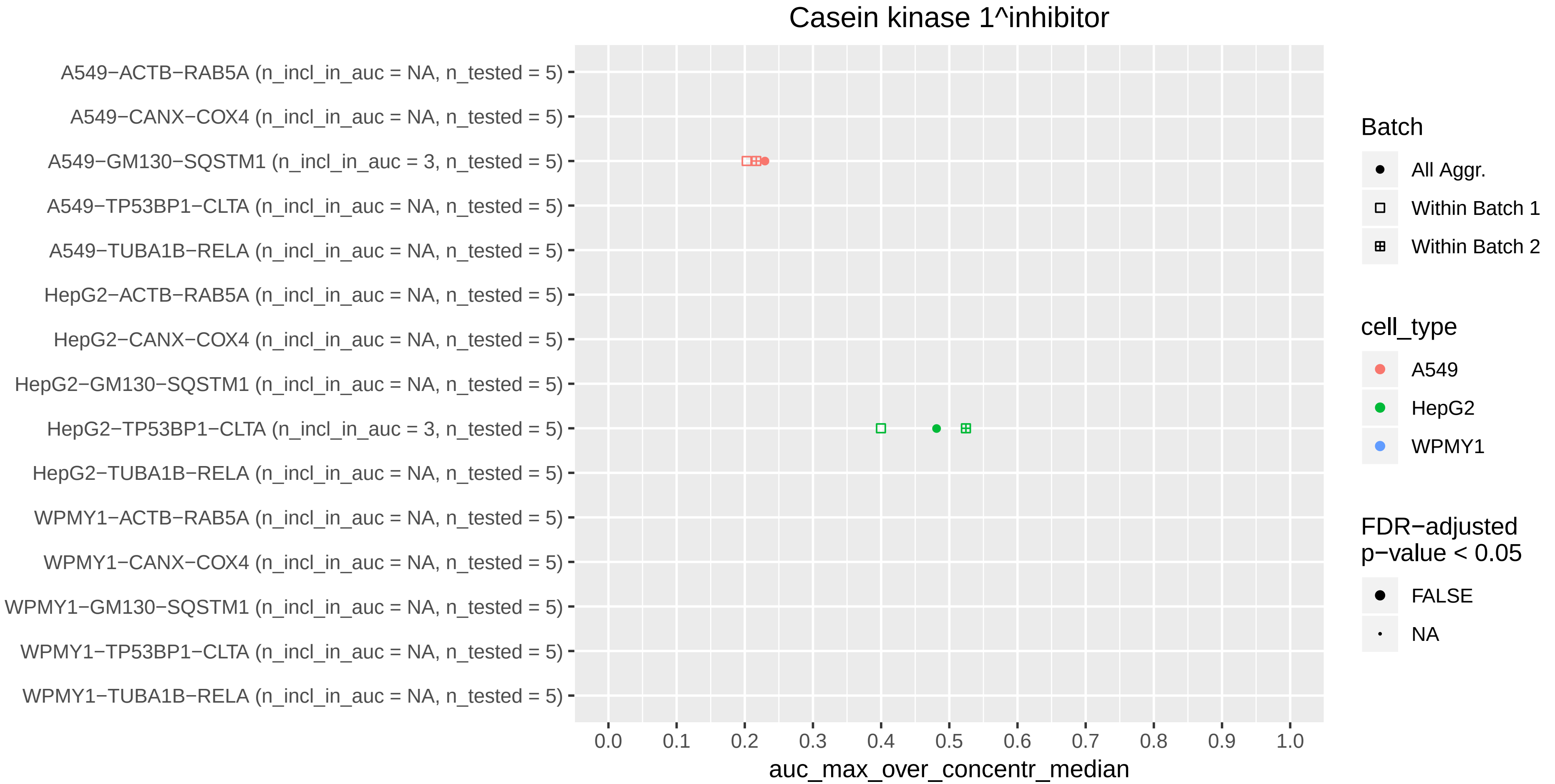

Casein kinase 1^inhibitor
A549−ACTB−RAB5A (n_incl_in_auc = NA, n_tested = 5)
A549−CANX−COX4 (n_incl_in_auc = NA, n_tested = 5)
A549−GM130−SQSTM1 (n_incl_in_auc = 3, n_tested = 5)
A549−TP53BP1−CLTA (n_incl_in_auc = NA, n_tested = 5)
A549−TUBA1B−RELA (n_incl_in_auc = NA, n_tested = 5)
HepG2−ACTB−RAB5A (n_incl_in_auc = NA, n_tested = 5)
HepG2−CANX−COX4 (n_incl_in_auc = NA, n_tested = 5)
HepG2−GM130−SQSTM1 (n_incl_in_auc = NA, n_tested = 5)
HepG2−TP53BP1−CLTA (n_incl_in_auc = 3, n_tested = 5)
HepG2−TUBA1B−RELA (n_incl_in_auc = NA, n_tested = 5)
WPMY1−ACTB−RAB5A (n_incl_in_auc = NA, n_tested = 5)
WPMY1−CANX−COX4 (n_incl_in_auc = NA, n_tested = 5)
WPMY1−GM130−SQSTM1 (n_incl_in_auc = NA, n_tested = 5)
WPMY1−TP53BP1−CLTA (n_incl_in_auc = NA, n_tested = 5)
WPMY1−TUBA1B−RELA (n_incl_in_auc = NA, n_tested = 5)
0.0
0.1
0.2
0.3
0.4
0.5
0.6
0.7
0.8
0.9
1.0
auc_max_over_concentr_median
Batch
All Aggr.
Within Batch 1
Within Batch 2
cell_type
A549
HepG2
WPMY1
FDR−adjusted p−value < 0.05
FALSE
NA

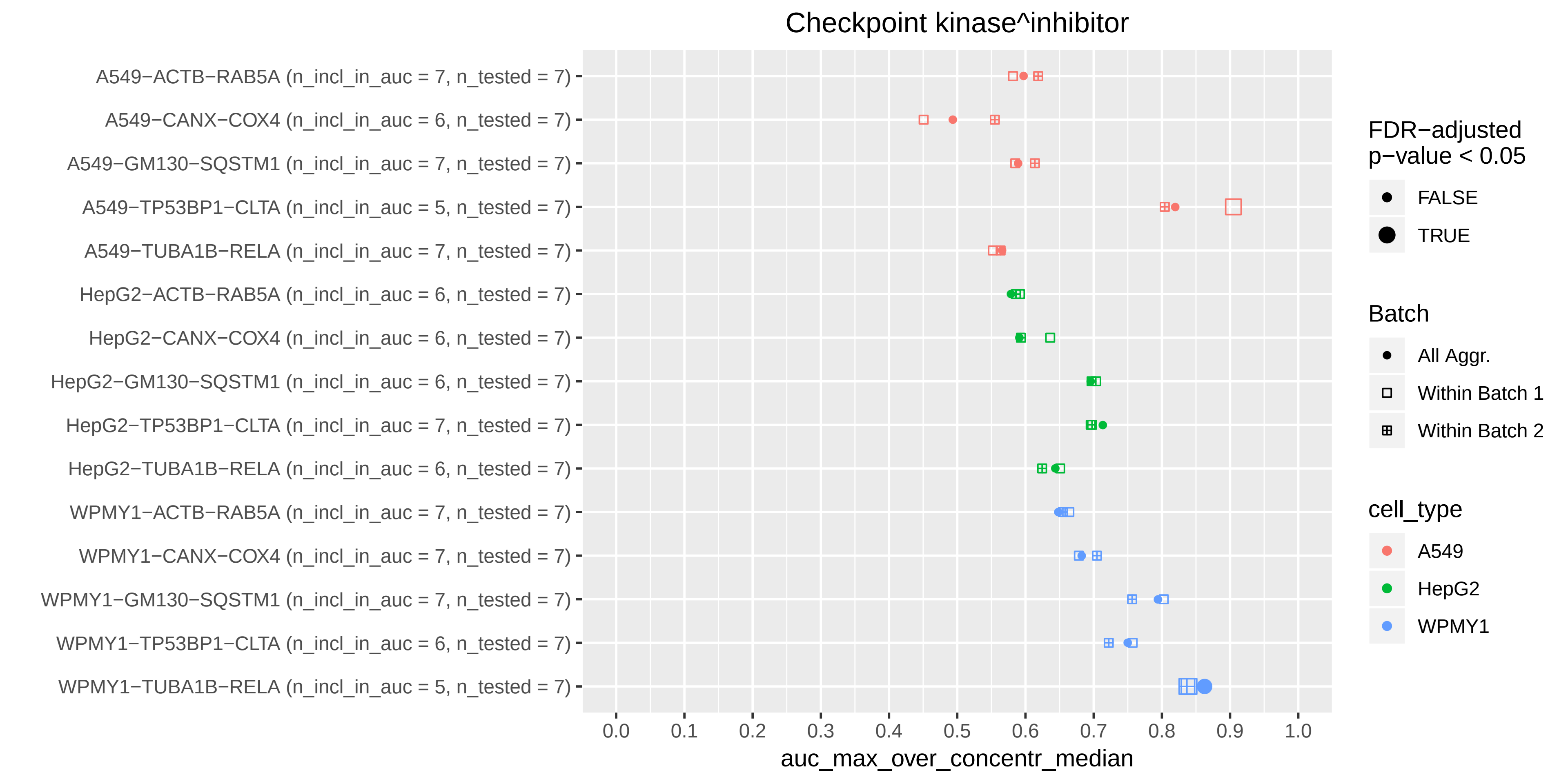
Checkpoint kinase^inhibitor
A549−ACTB−RAB5A (n_incl_in_auc = 7, n_tested = 7)
A549−CANX−COX4 (n_incl_in_auc = 6, n_tested = 7)
A549−GM130−SQSTM1 (n_incl_in_auc = 7, n_tested = 7)
A549−TP53BP1−CLTA (n_incl_in_auc = 5, n_tested = 7)
A549−TUBA1B−RELA (n_incl_in_auc = 7, n_tested = 7)
HepG2−ACTB−RAB5A (n_incl_in_auc = 6, n_tested = 7)
HepG2−CANX−COX4 (n_incl_in_auc = 6, n_tested = 7)
HepG2−GM130−SQSTM1 (n_incl_in_auc = 6, n_tested = 7)
HepG2−TP53BP1−CLTA (n_incl_in_auc = 7, n_tested = 7)
HepG2−TUBA1B−RELA (n_incl_in_auc = 6, n_tested = 7)
WPMY1−ACTB−RAB5A (n_incl_in_auc = 7, n_tested = 7)
WPMY1−CANX−COX4 (n_incl_in_auc = 7, n_tested = 7)
WPMY1−GM130−SQSTM1 (n_incl_in_auc = 7, n_tested = 7)
WPMY1−TP53BP1−CLTA (n_incl_in_auc = 6, n_tested = 7)
WPMY1−TUBA1B−RELA (n_incl_in_auc = 5, n_tested = 7)
0.0
0.1
0.2
0.3
0.4
0.5
0.6
0.7
0.8
0.9
1.0
auc_max_over_concentr_median
FDR−adjusted p−value < 0.05
FALSE
TRUE
Batch
All Aggr.
Within Batch 1
Within Batch 2
cell_type
A549
HepG2
WPMY1

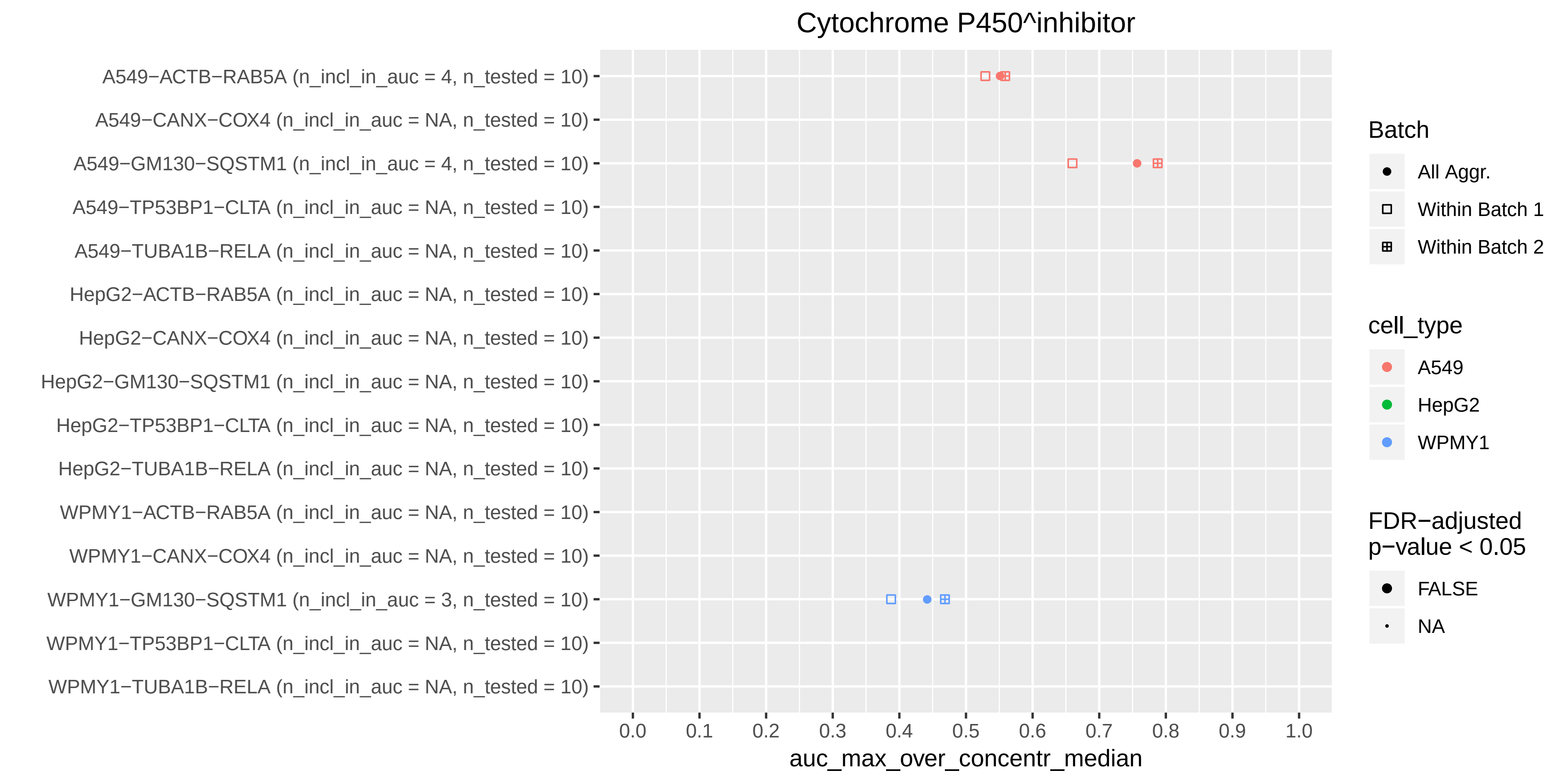
Cytochrome P450^inhibitor
A549−ACTB−RAB5A (n_incl_in_auc = 4, n_tested = 10)
A549−CANX−COX4 (n_incl_in_auc = NA, n_tested = 10)
A549−GM130−SQSTM1 (n_incl_in_auc = 4, n_tested = 10)
A549−TP53BP1−CLTA (n_incl_in_auc = NA, n_tested = 10)
A549−TUBA1B−RELA (n_incl_in_auc = NA, n_tested = 10)
HepG2−ACTB−RAB5A (n_incl_in_auc = NA, n_tested = 10)
HepG2−CANX−COX4 (n_incl_in_auc = NA, n_tested = 10)
HepG2−GM130−SQSTM1 (n_incl_in_auc = NA, n_tested = 10)
HepG2−TP53BP1−CLTA (n_incl_in_auc = NA, n_tested = 10)
HepG2−TUBA1B−RELA (n_incl_in_auc = NA, n_tested = 10)
WPMY1−ACTB−RAB5A (n_incl_in_auc = NA, n_tested = 10)
WPMY1−CANX−COX4 (n_incl_in_auc = NA, n_tested = 10)
WPMY1−GM130−SQSTM1 (n_incl_in_auc = 3, n_tested = 10)
WPMY1−TP53BP1−CLTA (n_incl_in_auc = NA, n_tested = 10)
WPMY1−TUBA1B−RELA (n_incl_in_auc = NA, n_tested = 10)
0.0
0.1
0.2
0.3
0.4
0.5
0.6
0.7
0.8
0.9
1.0
auc_max_over_concentr_median
Batch
All Aggr.
Within Batch 1
Within Batch 2
cell_type
A549
HepG2
WPMY1
FDR−adjusted p−value < 0.05
FALSE
NA

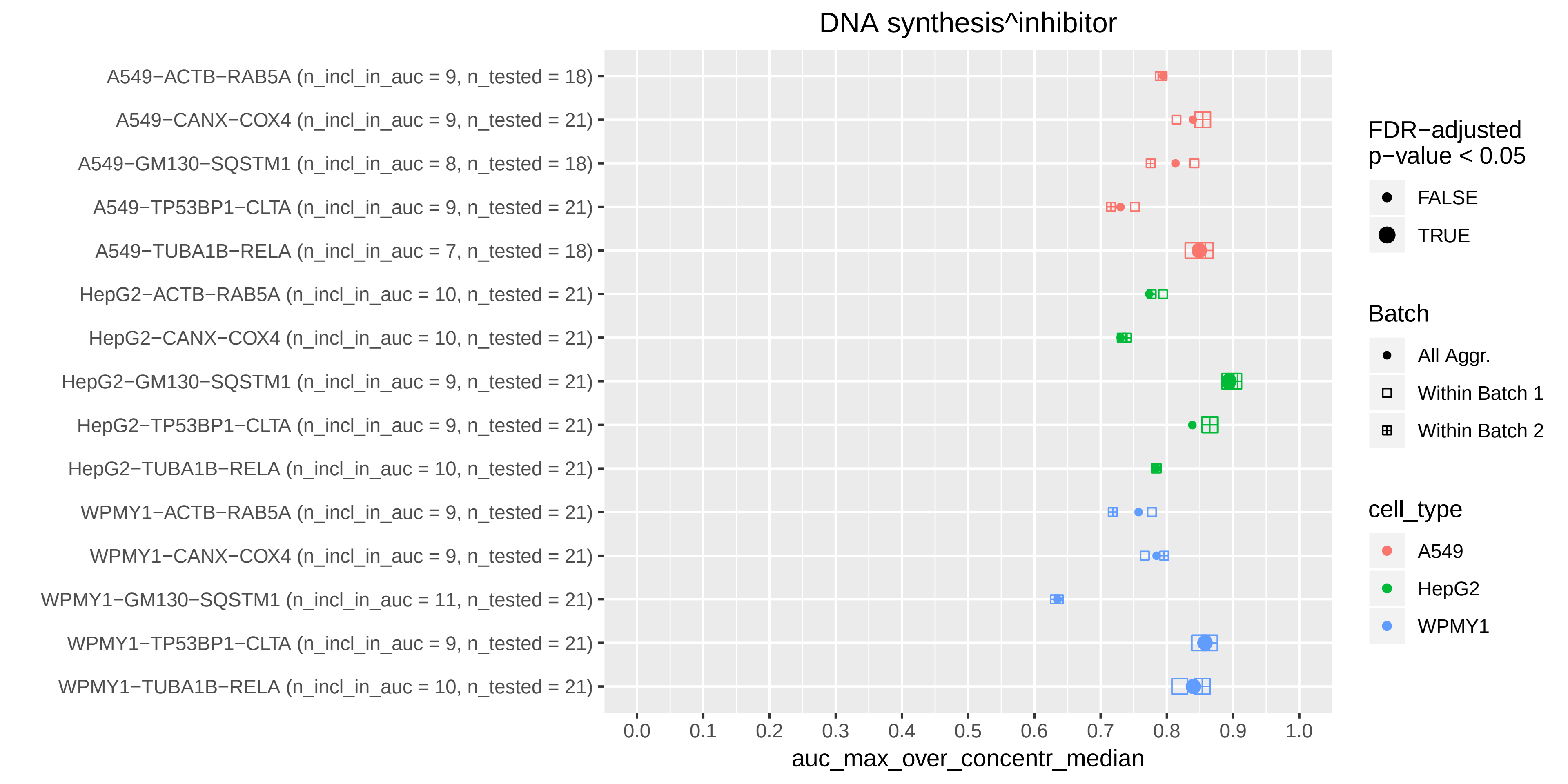
DNA synthesis^inhibitor
A549−ACTB−RAB5A (n_incl_in_auc = 9, n_tested = 18)
A549−CANX−COX4 (n_incl_in_auc = 9, n_tested = 21)
A549−GM130−SQSTM1 (n_incl_in_auc = 8, n_tested = 18)
A549−TP53BP1−CLTA (n_incl_in_auc = 9, n_tested = 21)
A549−TUBA1B−RELA (n_incl_in_auc = 7, n_tested = 18)
HepG2−ACTB−RAB5A (n_incl_in_auc = 10, n_tested = 21)
HepG2−CANX−COX4 (n_incl_in_auc = 10, n_tested = 21)
HepG2−GM130−SQSTM1 (n_incl_in_auc = 9, n_tested = 21)
HepG2−TP53BP1−CLTA (n_incl_in_auc = 9, n_tested = 21)
HepG2−TUBA1B−RELA (n_incl_in_auc = 10, n_tested = 21)
WPMY1−ACTB−RAB5A (n_incl_in_auc = 9, n_tested = 21)
WPMY1−CANX−COX4 (n_incl_in_auc = 9, n_tested = 21)
WPMY1−GM130−SQSTM1 (n_incl_in_auc = 11, n_tested = 21)
WPMY1−TP53BP1−CLTA (n_incl_in_auc = 9, n_tested = 21)
WPMY1−TUBA1B−RELA (n_incl_in_auc = 10, n_tested = 21)
0.0
0.1
0.2
0.3
0.4
0.5
0.6
0.7
0.8
0.9
1.0
auc_max_over_concentr_median
FDR−adjusted p−value < 0.05
FALSE
TRUE
Batch
All Aggr.
Within Batch 1
Within Batch 2
cell_type
A549
HepG2
WPMY1

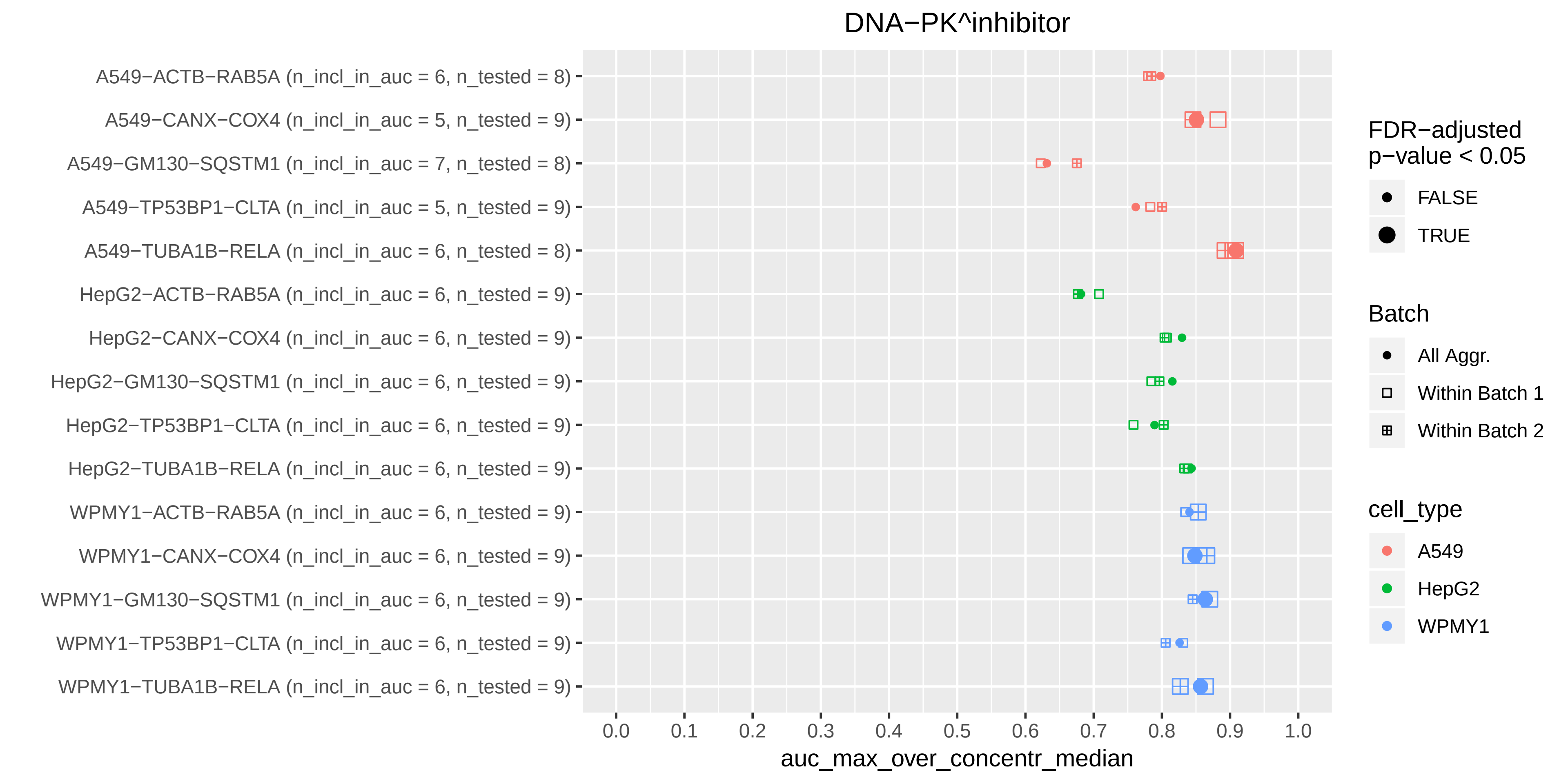

DNA−PK^inhibitor
A549−ACTB−RAB5A (n_incl_in_auc = 6, n_tested = 8)
A549−CANX−COX4 (n_incl_in_auc = 5, n_tested = 9)
A549−GM130−SQSTM1 (n_incl_in_auc = 7, n_tested = 8)
A549−TP53BP1−CLTA (n_incl_in_auc = 5, n_tested = 9)
A549−TUBA1B−RELA (n_incl_in_auc = 6, n_tested = 8)
HepG2−ACTB−RAB5A (n_incl_in_auc = 6, n_tested = 9)
HepG2−CANX−COX4 (n_incl_in_auc = 6, n_tested = 9)
HepG2−GM130−SQSTM1 (n_incl_in_auc = 6, n_tested = 9)
HepG2−TP53BP1−CLTA (n_incl_in_auc = 6, n_tested = 9)
HepG2−TUBA1B−RELA (n_incl_in_auc = 6, n_tested = 9)
WPMY1−ACTB−RAB5A (n_incl_in_auc = 6, n_tested = 9)
WPMY1−CANX−COX4 (n_incl_in_auc = 6, n_tested = 9)
WPMY1−GM130−SQSTM1 (n_incl_in_auc = 6, n_tested = 9)
WPMY1−TP53BP1−CLTA (n_incl_in_auc = 6, n_tested = 9)
WPMY1−TUBA1B−RELA (n_incl_in_auc = 6, n_tested = 9)
0.0
0.1
0.2
0.3
0.4
0.5
0.6
0.7
0.8
0.9
1.0
auc_max_over_concentr_median
FDR−adjusted p−value < 0.05
FALSE
TRUE
Batch
All Aggr.
Within Batch 1
Within Batch 2
cell_type
A549
HepG2
WPMY1

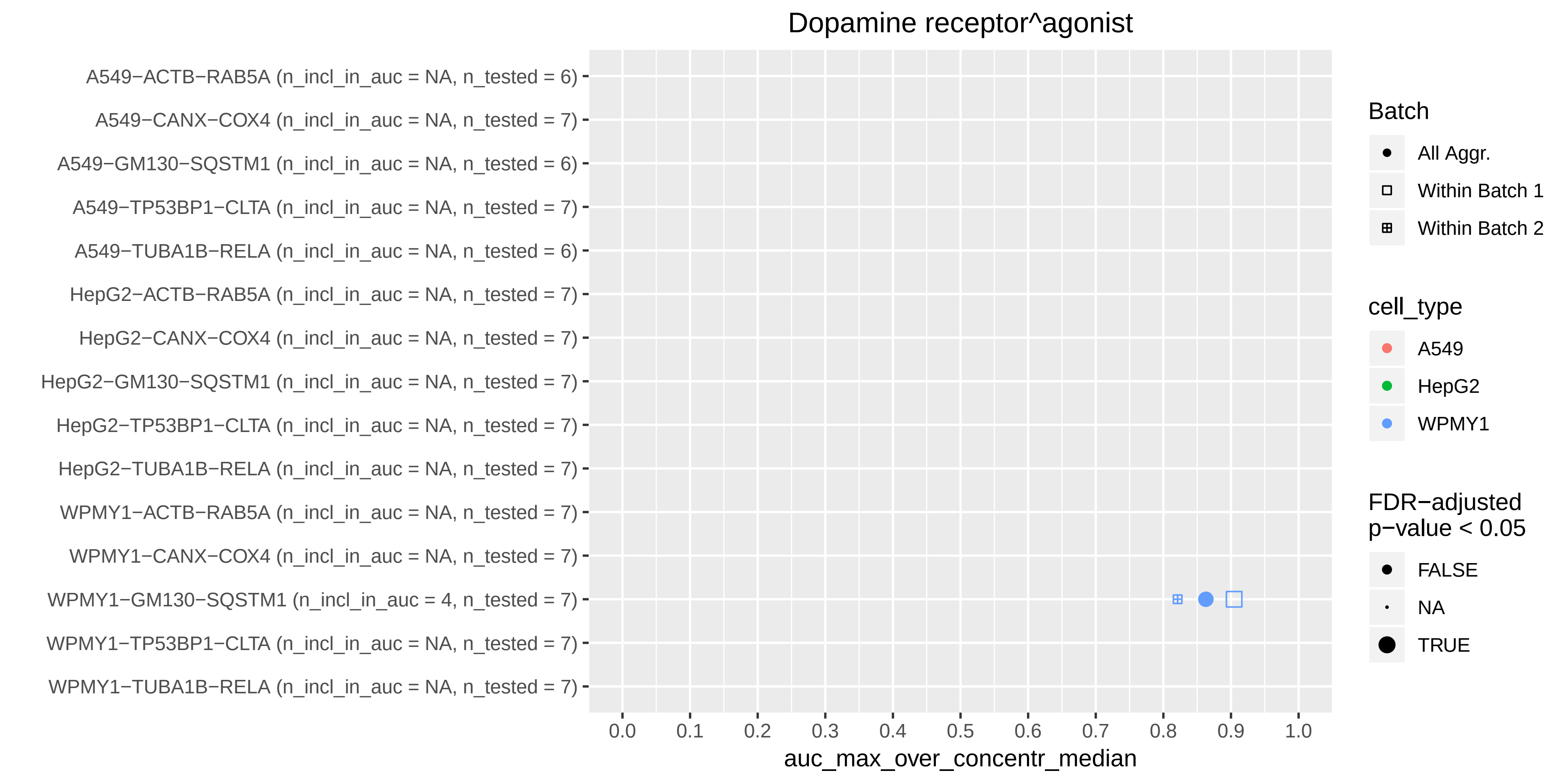

Dopamine receptor^agonist
A549−ACTB−RAB5A (n_incl_in_auc = NA, n_tested = 6)
A549−CANX−COX4 (n_incl_in_auc = NA, n_tested = 7)
A549−GM130−SQSTM1 (n_incl_in_auc = NA, n_tested = 6)
A549−TP53BP1−CLTA (n_incl_in_auc = NA, n_tested = 7)
A549−TUBA1B−RELA (n_incl_in_auc = NA, n_tested = 6)
HepG2−ACTB−RAB5A (n_incl_in_auc = NA, n_tested = 7)
HepG2−CANX−COX4 (n_incl_in_auc = NA, n_tested = 7)
HepG2−GM130−SQSTM1 (n_incl_in_auc = NA, n_tested = 7)
HepG2−TP53BP1−CLTA (n_incl_in_auc = NA, n_tested = 7)
HepG2−TUBA1B−RELA (n_incl_in_auc = NA, n_tested = 7)
WPMY1−ACTB−RAB5A (n_incl_in_auc = NA, n_tested = 7)
WPMY1−CANX−COX4 (n_incl_in_auc = NA, n_tested = 7)
WPMY1−GM130−SQSTM1 (n_incl_in_auc = 4, n_tested = 7)
WPMY1−TP53BP1−CLTA (n_incl_in_auc = NA, n_tested = 7)
WPMY1−TUBA1B−RELA (n_incl_in_auc = NA, n_tested = 7)
0.0
0.1
0.2
0.3
0.4
0.5
0.6
0.7
0.8
0.9
1.0
auc_max_over_concentr_median
Batch
All Aggr.
Within Batch 1
Within Batch 2
cell_type
A549
HepG2
WPMY1
FDR−adjusted p−value < 0.05
FALSE
NA
TRUE

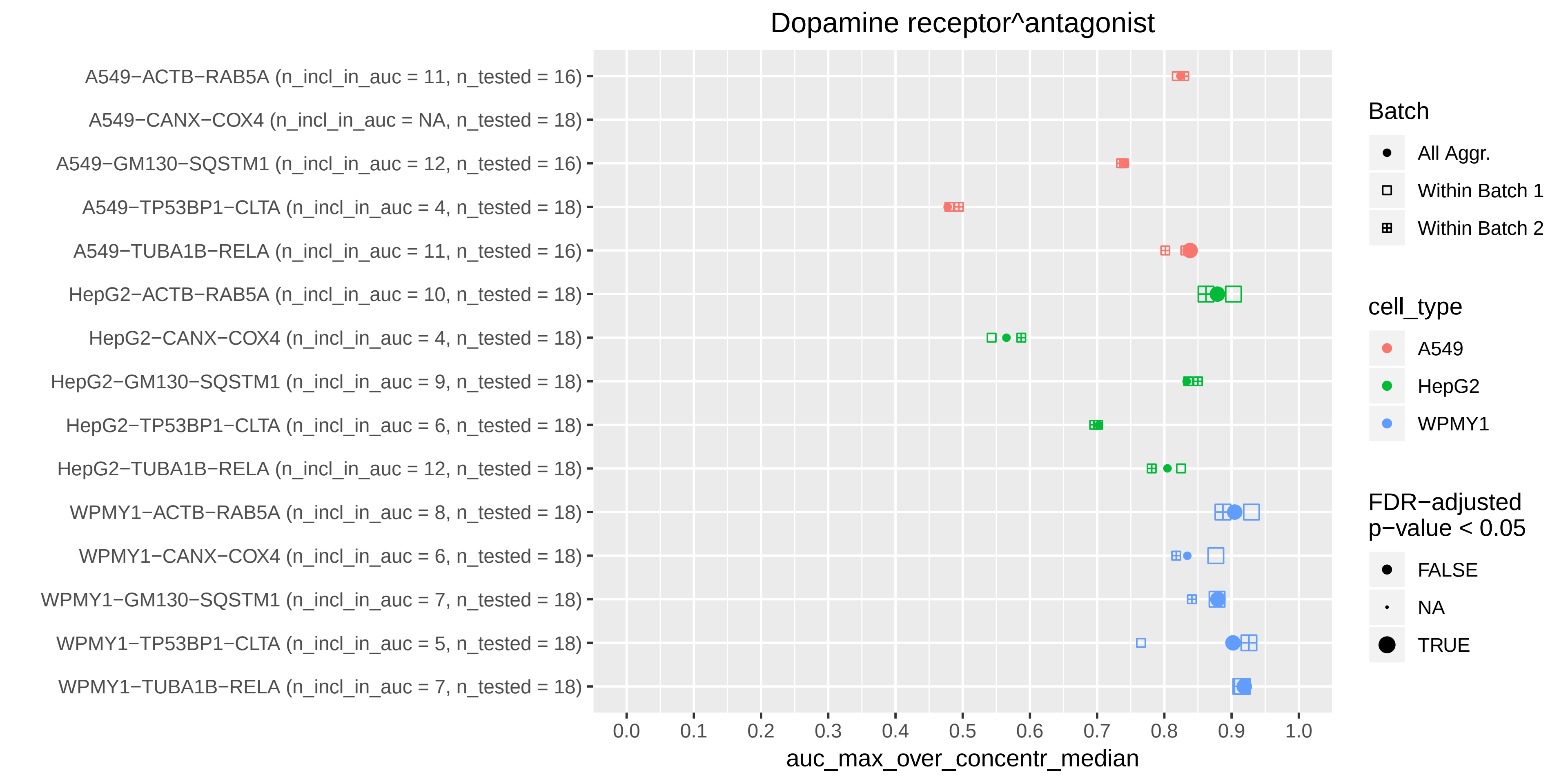
Dopamine receptor^antagonist
A549−ACTB−RAB5A (n_incl_in_auc = 11, n_tested = 16)
A549−CANX−COX4 (n_incl_in_auc = NA, n_tested = 18)
A549−GM130−SQSTM1 (n_incl_in_auc = 12, n_tested = 16)
A549−TP53BP1−CLTA (n_incl_in_auc = 4, n_tested = 18)
A549−TUBA1B−RELA (n_incl_in_auc = 11, n_tested = 16)
HepG2−ACTB−RAB5A (n_incl_in_auc = 10, n_tested = 18)
HepG2−CANX−COX4 (n_incl_in_auc = 4, n_tested = 18)
HepG2−GM130−SQSTM1 (n_incl_in_auc = 9, n_tested = 18)
HepG2−TP53BP1−CLTA (n_incl_in_auc = 6, n_tested = 18)
HepG2−TUBA1B−RELA (n_incl_in_auc = 12, n_tested = 18)
WPMY1−ACTB−RAB5A (n_incl_in_auc = 8, n_tested = 18)
WPMY1−CANX−COX4 (n_incl_in_auc = 6, n_tested = 18)
WPMY1−GM130−SQSTM1 (n_incl_in_auc = 7, n_tested = 18)
WPMY1−TP53BP1−CLTA (n_incl_in_auc = 5, n_tested = 18)
WPMY1−TUBA1B−RELA (n_incl_in_auc = 7, n_tested = 18)
0.0
0.1
0.2
0.3
0.4
0.5
0.6
0.7
0.8
0.9
1.0
auc_max_over_concentr_median
Batch
All Aggr.
Within Batch 1
Within Batch 2
cell_type
A549
HepG2
WPMY1
FDR−adjusted p−value < 0.05
FALSE
NA
TRUE

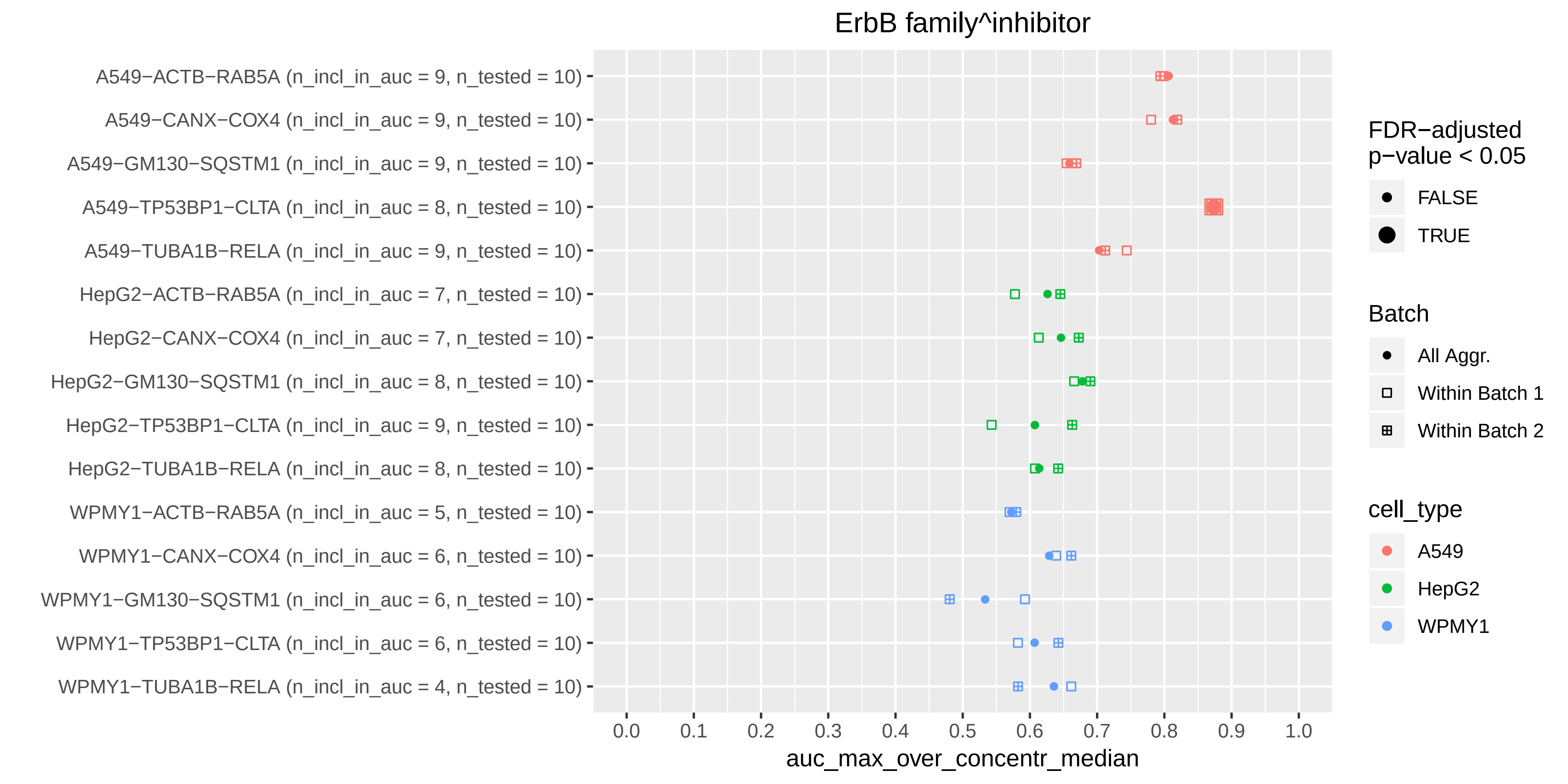

ErbB family^inhibitor
A549−ACTB−RAB5A (n_incl_in_auc = 9, n_tested = 10)
A549−CANX−COX4 (n_incl_in_auc = 9, n_tested = 10)
A549−GM130−SQSTM1 (n_incl_in_auc = 9, n_tested = 10)
A549−TP53BP1−CLTA (n_incl_in_auc = 8, n_tested = 10)
A549−TUBA1B−RELA (n_incl_in_auc = 9, n_tested = 10)
HepG2−ACTB−RAB5A (n_incl_in_auc = 7, n_tested = 10)
HepG2−CANX−COX4 (n_incl_in_auc = 7, n_tested = 10)
HepG2−GM130−SQSTM1 (n_incl_in_auc = 8, n_tested = 10)
HepG2−TP53BP1−CLTA (n_incl_in_auc = 9, n_tested = 10)
HepG2−TUBA1B−RELA (n_incl_in_auc = 8, n_tested = 10)
WPMY1−ACTB−RAB5A (n_incl_in_auc = 5, n_tested = 10)
WPMY1−CANX−COX4 (n_incl_in_auc = 6, n_tested = 10)
WPMY1−GM130−SQSTM1 (n_incl_in_auc = 6, n_tested = 10)
WPMY1−TP53BP1−CLTA (n_incl_in_auc = 6, n_tested = 10)
WPMY1−TUBA1B−RELA (n_incl_in_auc = 4, n_tested = 10)
0.0
0.1
0.2
0.3
0.4
0.5
0.6
0.7
0.8
0.9
1.0
auc_max_over_concentr_median
FDR−adjusted
p−value < 0.05
FALSE
TRUE
Batch
All Aggr.
Within Batch 1
Within Batch 2
cell_type
A549
HepG2
WPMY1

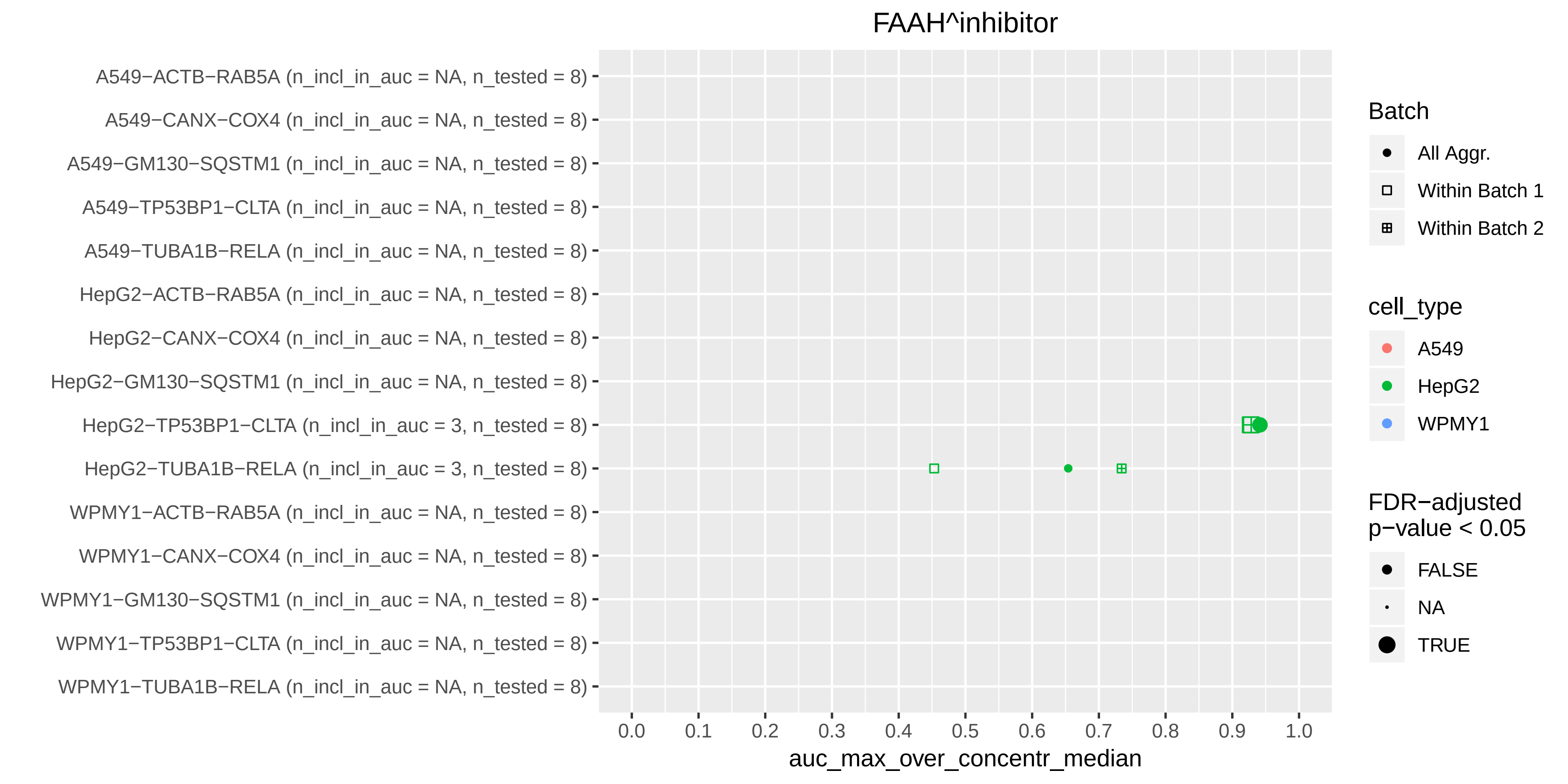
FAAH^inhibitor
A549−ACTB−RAB5A (n_incl_in_auc = NA, n_tested = 8)
A549−CANX−COX4 (n_incl_in_auc = NA, n_tested = 8)
A549−GM130−SQSTM1 (n_incl_in_auc = NA, n_tested = 8)
A549−TP53BP1−CLTA (n_incl_in_auc = NA, n_tested = 8)
A549−TUBA1B−RELA (n_incl_in_auc = NA, n_tested = 8)
HepG2−ACTB−RAB5A (n_incl_in_auc = NA, n_tested = 8)
HepG2−CANX−COX4 (n_incl_in_auc = NA, n_tested = 8)
HepG2−GM130−SQSTM1 (n_incl_in_auc = NA, n_tested = 8)
HepG2−TP53BP1−CLTA (n_incl_in_auc = 3, n_tested = 8)
HepG2−TUBA1B−RELA (n_incl_in_auc = 3, n_tested = 8)
WPMY1−ACTB−RAB5A (n_incl_in_auc = NA, n_tested = 8)
WPMY1−CANX−COX4 (n_incl_in_auc = NA, n_tested = 8)
WPMY1−GM130−SQSTM1 (n_incl_in_auc = NA, n_tested = 8)
WPMY1−TP53BP1−CLTA (n_incl_in_auc = NA, n_tested = 8)
WPMY1−TUBA1B−RELA (n_incl_in_auc = NA, n_tested = 8)
0.0
0.1
0.2
0.3
0.4
0.5
0.6
0.7
0.8
0.9
1.0
auc_max_over_concentr_median
Batch
All Aggr.
Within Batch 1
Within Batch 2
cell_type
A549
HepG2
WPMY1
FDR−adjusted
p−value < 0.05
FALSE
NA
TRUE

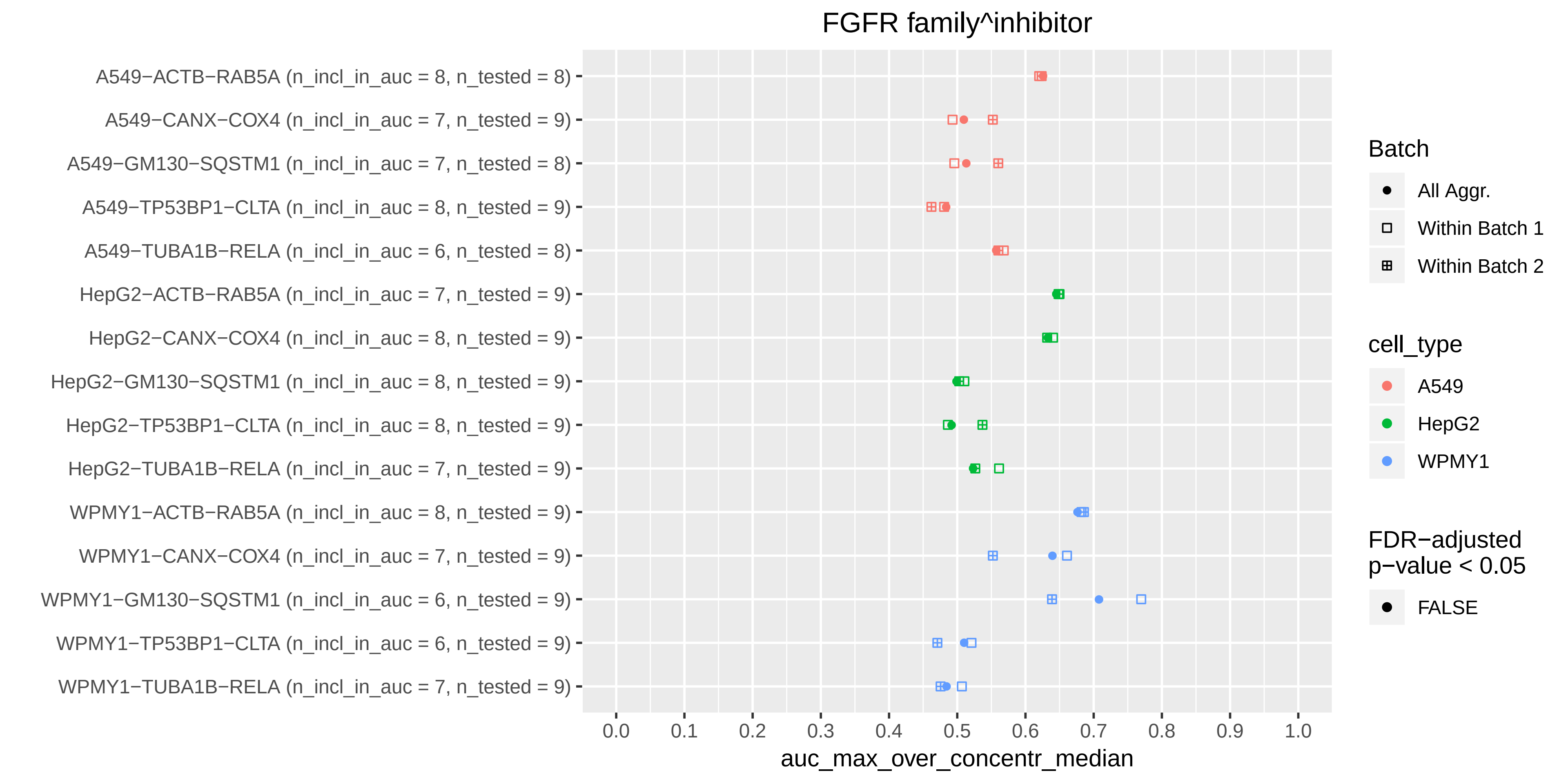

FGFR family^inhibitor
A549−ACTB−RAB5A (n_incl_in_auc = 8, n_tested = 8)
A549−CANX−COX4 (n_incl_in_auc = 7, n_tested = 9)
A549−GM130−SQSTM1 (n_incl_in_auc = 7, n_tested = 8)
A549−TP53BP1−CLTA (n_incl_in_auc = 8, n_tested = 9)
A549−TUBA1B−RELA (n_incl_in_auc = 6, n_tested = 8)
HepG2−ACTB−RAB5A (n_incl_in_auc = 7, n_tested = 9)
HepG2−CANX−COX4 (n_incl_in_auc = 8, n_tested = 9)
HepG2−GM130−SQSTM1 (n_incl_in_auc = 8, n_tested = 9)
HepG2−TP53BP1−CLTA (n_incl_in_auc = 8, n_tested = 9)
HepG2−TUBA1B−RELA (n_incl_in_auc = 7, n_tested = 9)
WPMY1−ACTB−RAB5A (n_incl_in_auc = 8, n_tested = 9)
WPMY1−CANX−COX4 (n_incl_in_auc = 7, n_tested = 9)
WPMY1−GM130−SQSTM1 (n_incl_in_auc = 6, n_tested = 9)
WPMY1−TP53BP1−CLTA (n_incl_in_auc = 6, n_tested = 9)
WPMY1−TUBA1B−RELA (n_incl_in_auc = 7, n_tested = 9)
0.0
0.1
0.2
0.3
0.4
0.5
0.6
0.7
0.8
0.9
1.0
auc_max_over_concentr_median
Batch
All Aggr.
Within Batch 1
Within Batch 2
cell_type
A549
HepG2
WPMY1
FDR−adjusted
p−value < 0.05
FALSE

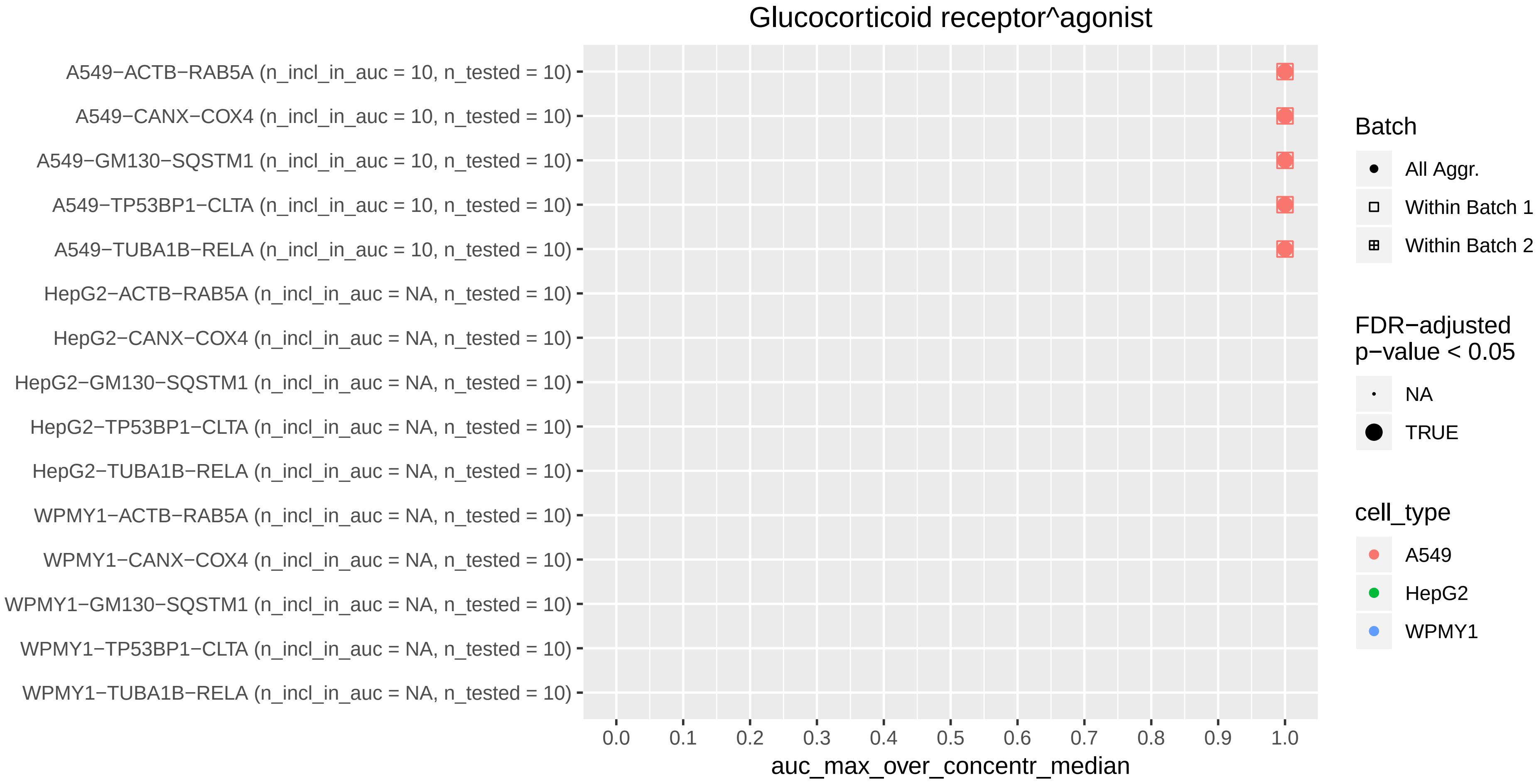
Glucocorticoid receptor^agonist
A549−ACTB−RAB5A (n_incl_in_auc = 10, n_tested = 10)
A549−CANX−COX4 (n_incl_in_auc = 10, n_tested = 10)
A549−GM130−SQSTM1 (n_incl_in_auc = 10, n_tested = 10)
A549−TP53BP1−CLTA (n_incl_in_auc = 10, n_tested = 10)
A549−TUBA1B−RELA (n_incl_in_auc = 10, n_tested = 10)
HepG2−ACTB−RAB5A (n_incl_in_auc = NA, n_tested = 10)
HepG2−CANX−COX4 (n_incl_in_auc = NA, n_tested = 10)
HepG2−GM130−SQSTM1 (n_incl_in_auc = NA, n_tested = 10)
HepG2−TP53BP1−CLTA (n_incl_in_auc = NA, n_tested = 10)
HepG2−TUBA1B−RELA (n_incl_in_auc = NA, n_tested = 10)
WPMY1−ACTB−RAB5A (n_incl_in_auc = NA, n_tested = 10)
WPMY1−CANX−COX4 (n_incl_in_auc = NA, n_tested = 10)
WPMY1−GM130−SQSTM1 (n_incl_in_auc = NA, n_tested = 10)
WPMY1−TP53BP1−CLTA (n_incl_in_auc = NA, n_tested = 10)
WPMY1−TUBA1B−RELA (n_incl_in_auc = NA, n_tested = 10)
0.0
0.1
0.2
0.3
0.4
0.5
0.6
0.7
0.8
0.9
1.0
auc_max_over_concentr_median
Batch
All Aggr.
Within Batch 1
Within Batch 2
FDR−adjusted
p−value < 0.05
NA
TRUE
cell_type
A549
HepG2
WPMY1

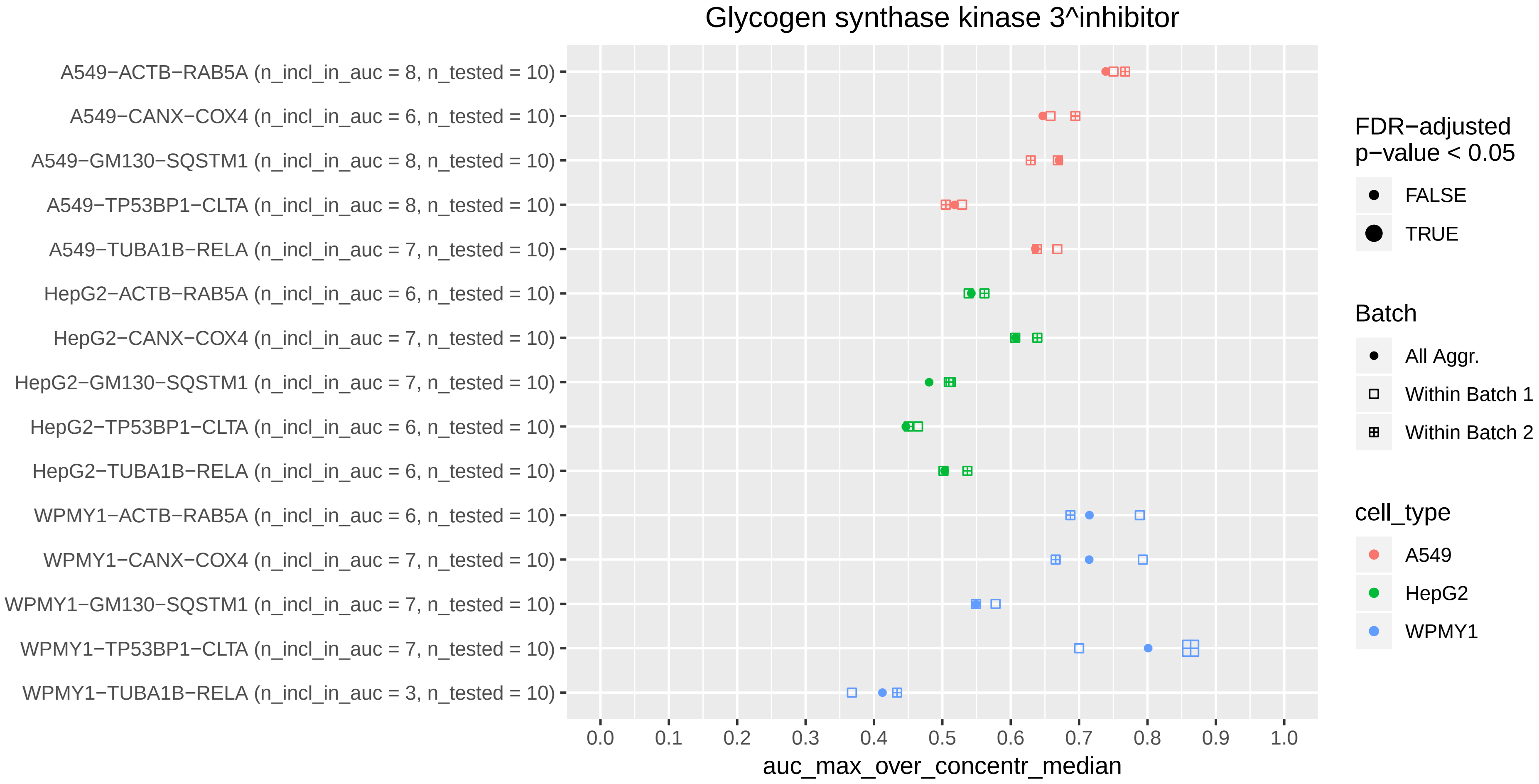

Glycogen synthase kinase 3^inhibitor
A549−ACTB−RAB5A (n_incl_in_auc = 8, n_tested = 10)
A549−CANX−COX4 (n_incl_in_auc = 6, n_tested = 10)
A549−GM130−SQSTM1 (n_incl_in_auc = 8, n_tested = 10)
A549−TP53BP1−CLTA (n_incl_in_auc = 8, n_tested = 10)
A549−TUBA1B−RELA (n_incl_in_auc = 7, n_tested = 10)
HepG2−ACTB−RAB5A (n_incl_in_auc = 6, n_tested = 10)
HepG2−CANX−COX4 (n_incl_in_auc = 7, n_tested = 10)
HepG2−GM130−SQSTM1 (n_incl_in_auc = 7, n_tested = 10)
HepG2−TP53BP1−CLTA (n_incl_in_auc = 6, n_tested = 10)
HepG2−TUBA1B−RELA (n_incl_in_auc = 6, n_tested = 10)
WPMY1−ACTB−RAB5A (n_incl_in_auc = 6, n_tested = 10)
WPMY1−CANX−COX4 (n_incl_in_auc = 7, n_tested = 10)
WPMY1−GM130−SQSTM1 (n_incl_in_auc = 7, n_tested = 10)
WPMY1−TP53BP1−CLTA (n_incl_in_auc = 7, n_tested = 10)
WPMY1−TUBA1B−RELA (n_incl_in_auc = 3, n_tested = 10)
0.0
0.1
0.2
0.3
0.4
0.5
0.6
0.7
0.8
0.9
1.0
auc_max_over_concentr_median
FDR−adjusted p−value < 0.05
FALSE
TRUE
Batch
All Aggr.
Within Batch 1
Within Batch 2
cell_type
A549
HepG2
WPMY1

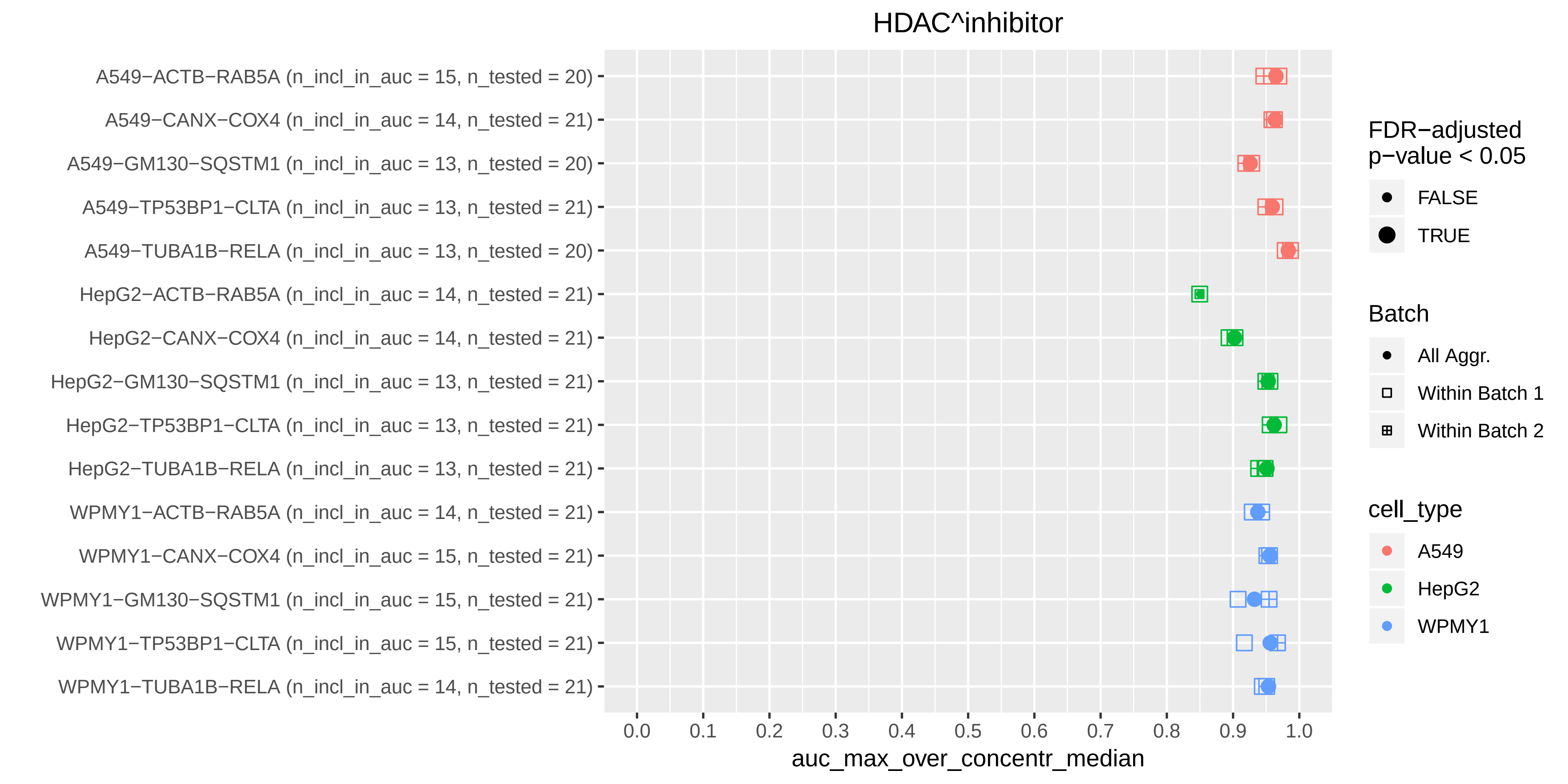

HDAC^inhibitor
A549−ACTB−RAB5A (n_incl_in_auc = 15, n_tested = 20)
A549−CANX−COX4 (n_incl_in_auc = 14, n_tested = 21)
A549−GM130−SQSTM1 (n_incl_in_auc = 13, n_tested = 20)
A549−TP53BP1−CLTA (n_incl_in_auc = 13, n_tested = 21)
A549−TUBA1B−RELA (n_incl_in_auc = 13, n_tested = 20)
HepG2−ACTB−RAB5A (n_incl_in_auc = 14, n_tested = 21)
HepG2−CANX−COX4 (n_incl_in_auc = 14, n_tested = 21)
HepG2−GM130−SQSTM1 (n_incl_in_auc = 13, n_tested = 21)
HepG2−TP53BP1−CLTA (n_incl_in_auc = 13, n_tested = 21)
HepG2−TUBA1B−RELA (n_incl_in_auc = 13, n_tested = 21)
WPMY1−ACTB−RAB5A (n_incl_in_auc = 14, n_tested = 21)
WPMY1−CANX−COX4 (n_incl_in_auc = 15, n_tested = 21)
WPMY1−GM130−SQSTM1 (n_incl_in_auc = 15, n_tested = 21)
WPMY1−TP53BP1−CLTA (n_incl_in_auc = 15, n_tested = 21)
WPMY1−TUBA1B−RELA (n_incl_in_auc = 14, n_tested = 21)
0.0 0.1 0.2 0.3 0.4 0.5 0.6 0.7 0.8 0.9 1.0
auc_max_over_concentr_median
FDR−adjusted p−value < 0.05
FALSE
TRUE
Batch
All Aggr.
Within Batch 1
Within Batch 2
cell_type
A549
HepG2
WPMY1

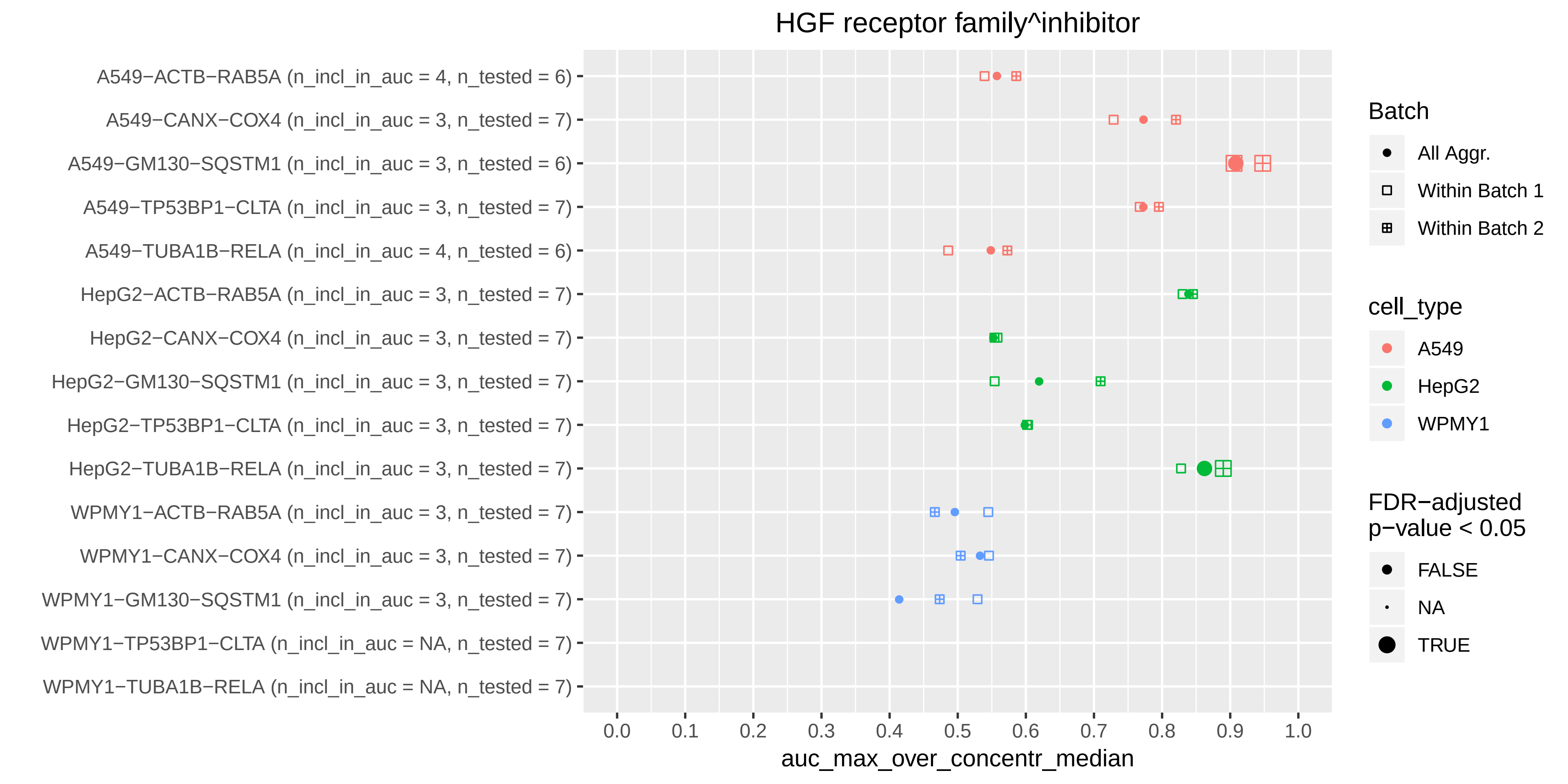
HGF receptor family^inhibitor
A549−ACTB−RAB5A (n_incl_in_auc = 4, n_tested = 6)
A549−CANX−COX4 (n_incl_in_auc = 3, n_tested = 7)
A549−GM130−SQSTM1 (n_incl_in_auc = 3, n_tested = 6)
A549−TP53BP1−CLTA (n_incl_in_auc = 3, n_tested = 7)
A549−TUBA1B−RELA (n_incl_in_auc = 4, n_tested = 6)
HepG2−ACTB−RAB5A (n_incl_in_auc = 3, n_tested = 7)
HepG2−CANX−COX4 (n_incl_in_auc = 3, n_tested = 7)
HepG2−GM130−SQSTM1 (n_incl_in_auc = 3, n_tested = 7)
HepG2−TP53BP1−CLTA (n_incl_in_auc = 3, n_tested = 7)
HepG2−TUBA1B−RELA (n_incl_in_auc = 3, n_tested = 7)
WPMY1−ACTB−RAB5A (n_incl_in_auc = 3, n_tested = 7)
WPMY1−CANX−COX4 (n_incl_in_auc = 3, n_tested = 7)
WPMY1−GM130−SQSTM1 (n_incl_in_auc = 3, n_tested = 7)
WPMY1−TP53BP1−CLTA (n_incl_in_auc = NA, n_tested = 7)
WPMY1−TUBA1B−RELA (n_incl_in_auc = NA, n_tested = 7)
0.0
0.1
0.2
0.3
0.4
0.5
0.6
0.7
0.8
0.9
1.0
auc_max_over_concentr_median
Batch
All Aggr.
Within Batch 1
Within Batch 2
cell_type
A549
HepG2
WPMY1
FDR−adjusted p−value < 0.05
FALSE
NA
TRUE

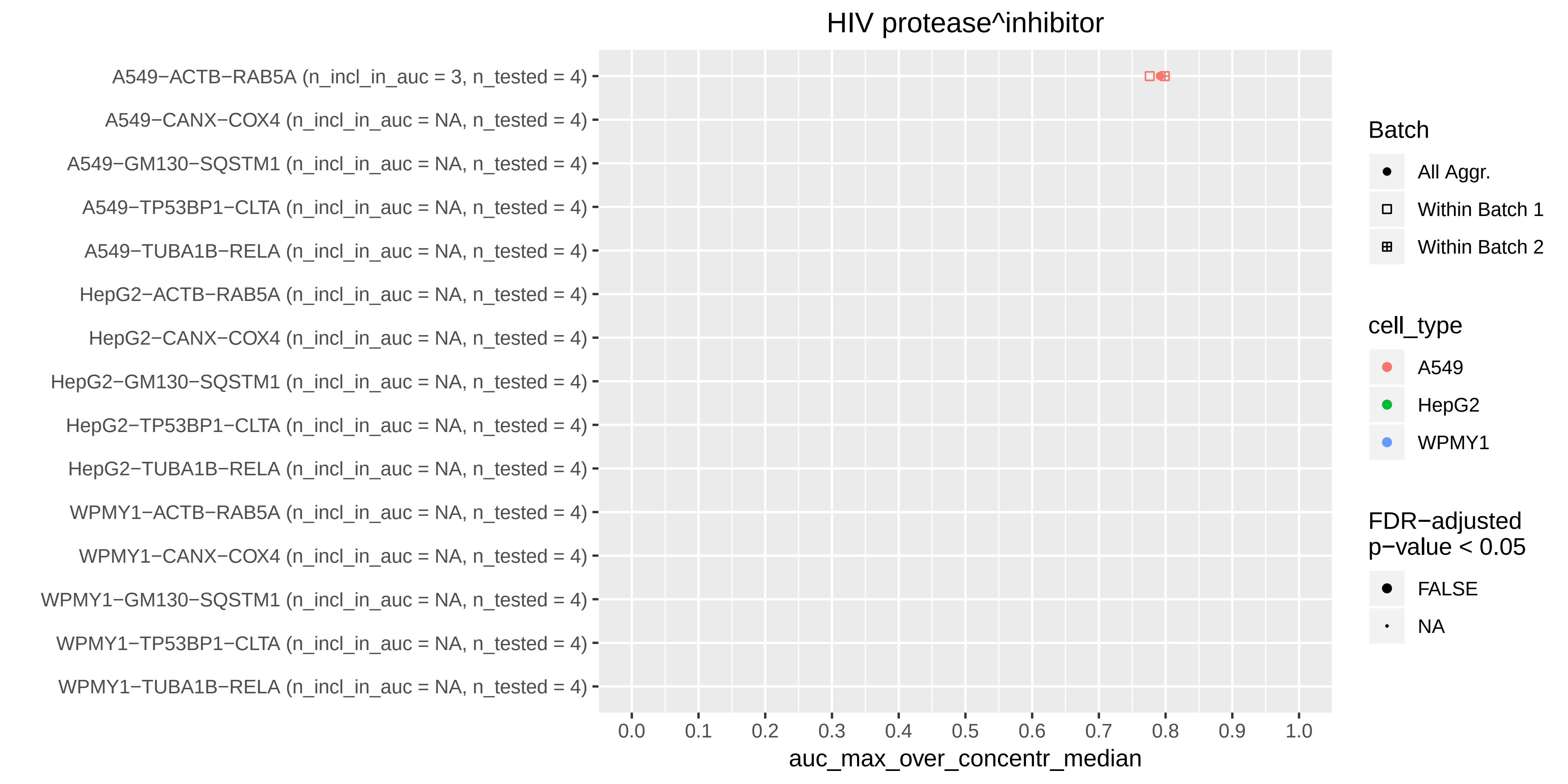
HIV protease^inhibitor
A549−ACTB−RAB5A (n_incl_in_auc = 3, n_tested = 4)
A549−CANX−COX4 (n_incl_in_auc = NA, n_tested = 4)
A549−GM130−SQSTM1 (n_incl_in_auc = NA, n_tested = 4)
A549−TP53BP1−CLTA (n_incl_in_auc = NA, n_tested = 4)
A549−TUBA1B−RELA (n_incl_in_auc = NA, n_tested = 4)
HepG2−ACTB−RAB5A (n_incl_in_auc = NA, n_tested = 4)
HepG2−CANX−COX4 (n_incl_in_auc = NA, n_tested = 4)
HepG2−GM130−SQSTM1 (n_incl_in_auc = NA, n_tested = 4)
HepG2−TP53BP1−CLTA (n_incl_in_auc = NA, n_tested = 4)
HepG2−TUBA1B−RELA (n_incl_in_auc = NA, n_tested = 4)
WPMY1−ACTB−RAB5A (n_incl_in_auc = NA, n_tested = 4)
WPMY1−CANX−COX4 (n_incl_in_auc = NA, n_tested = 4)
WPMY1−GM130−SQSTM1 (n_incl_in_auc = NA, n_tested = 4)
WPMY1−TP53BP1−CLTA (n_incl_in_auc = NA, n_tested = 4)
WPMY1−TUBA1B−RELA (n_incl_in_auc = NA, n_tested = 4)
0.0
0.1
0.2
0.3
0.4
0.5
0.6
0.7
0.8
0.9
1.0
auc_max_over_concentr_median
Batch
All Aggr.
Within Batch 1
Within Batch 2
cell_type
A549
HepG2
WPMY1
FDR−adjusted p−value < 0.05
FALSE
NA

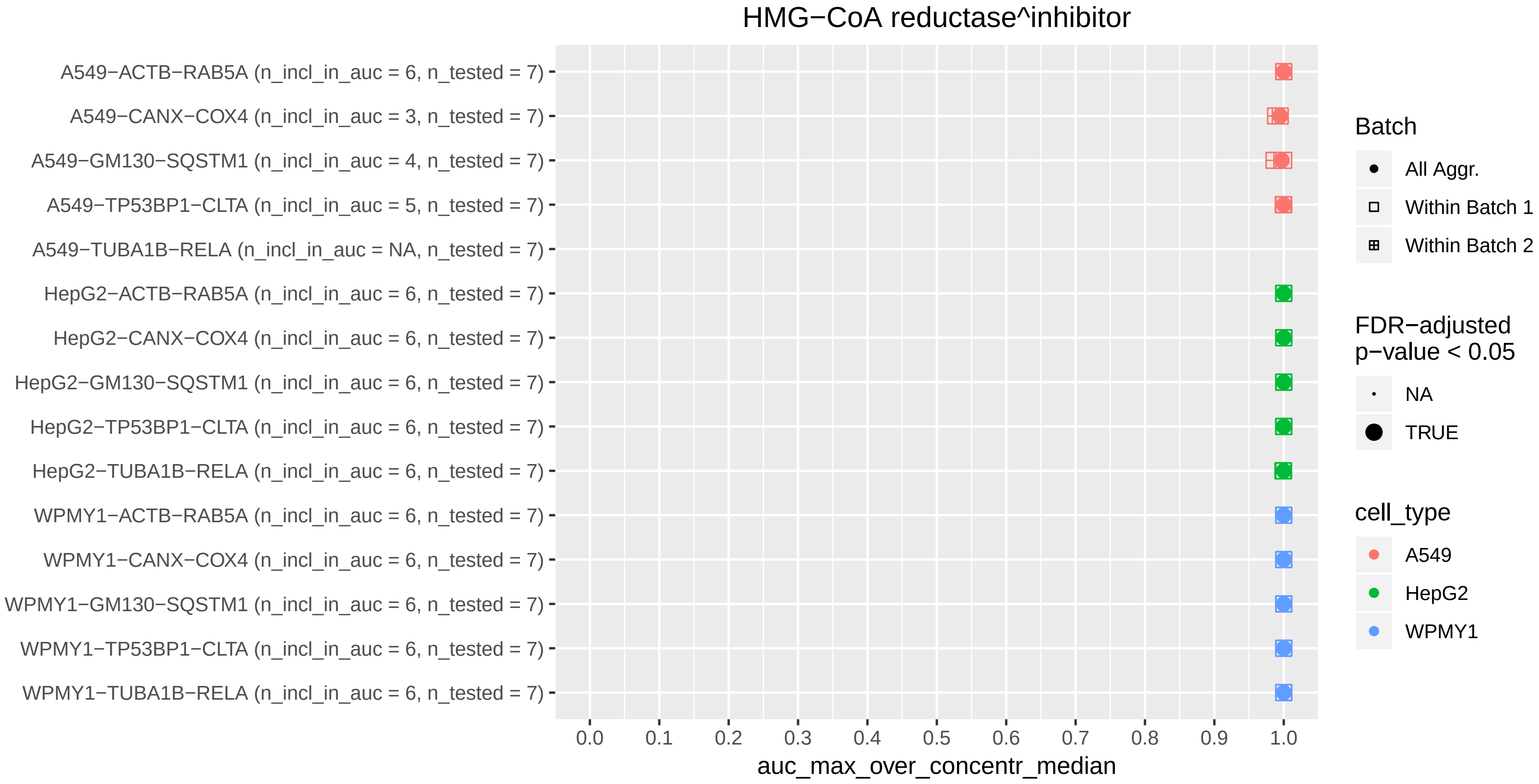

HMG−CoA reductase^inhibitor
A549−ACTB−RAB5A (n_incl_in_auc = 6, n_tested = 7)
A549−CANX−COX4 (n_incl_in_auc = 3, n_tested = 7)
A549−GM130−SQSTM1 (n_incl_in_auc = 4, n_tested = 7)
A549−TP53BP1−CLTA (n_incl_in_auc = 5, n_tested = 7)
A549−TUBA1B−RELA (n_incl_in_auc = NA, n_tested = 7)
HepG2−ACTB−RAB5A (n_incl_in_auc = 6, n_tested = 7)
HepG2−CANX−COX4 (n_incl_in_auc = 6, n_tested = 7)
HepG2−GM130−SQSTM1 (n_incl_in_auc = 6, n_tested = 7)
HepG2−TP53BP1−CLTA (n_incl_in_auc = 6, n_tested = 7)
HepG2−TUBA1B−RELA (n_incl_in_auc = 6, n_tested = 7)
WPMY1−ACTB−RAB5A (n_incl_in_auc = 6, n_tested = 7)
WPMY1−CANX−COX4 (n_incl_in_auc = 6, n_tested = 7)
WPMY1−GM130−SQSTM1 (n_incl_in_auc = 6, n_tested = 7)
WPMY1−TP53BP1−CLTA (n_incl_in_auc = 6, n_tested = 7)
WPMY1−TUBA1B−RELA (n_incl_in_auc = 6, n_tested = 7)
0.0
0.1
0.2
0.3
0.4
0.5
0.6
0.7
0.8
0.9
1.0
auc_max_over_concentr_median
Batch
All Aggr.
Within Batch 1
Within Batch 2
FDR−adjusted
p−value < 0.05
NA
TRUE
cell_type
A549
HepG2
WPMY1

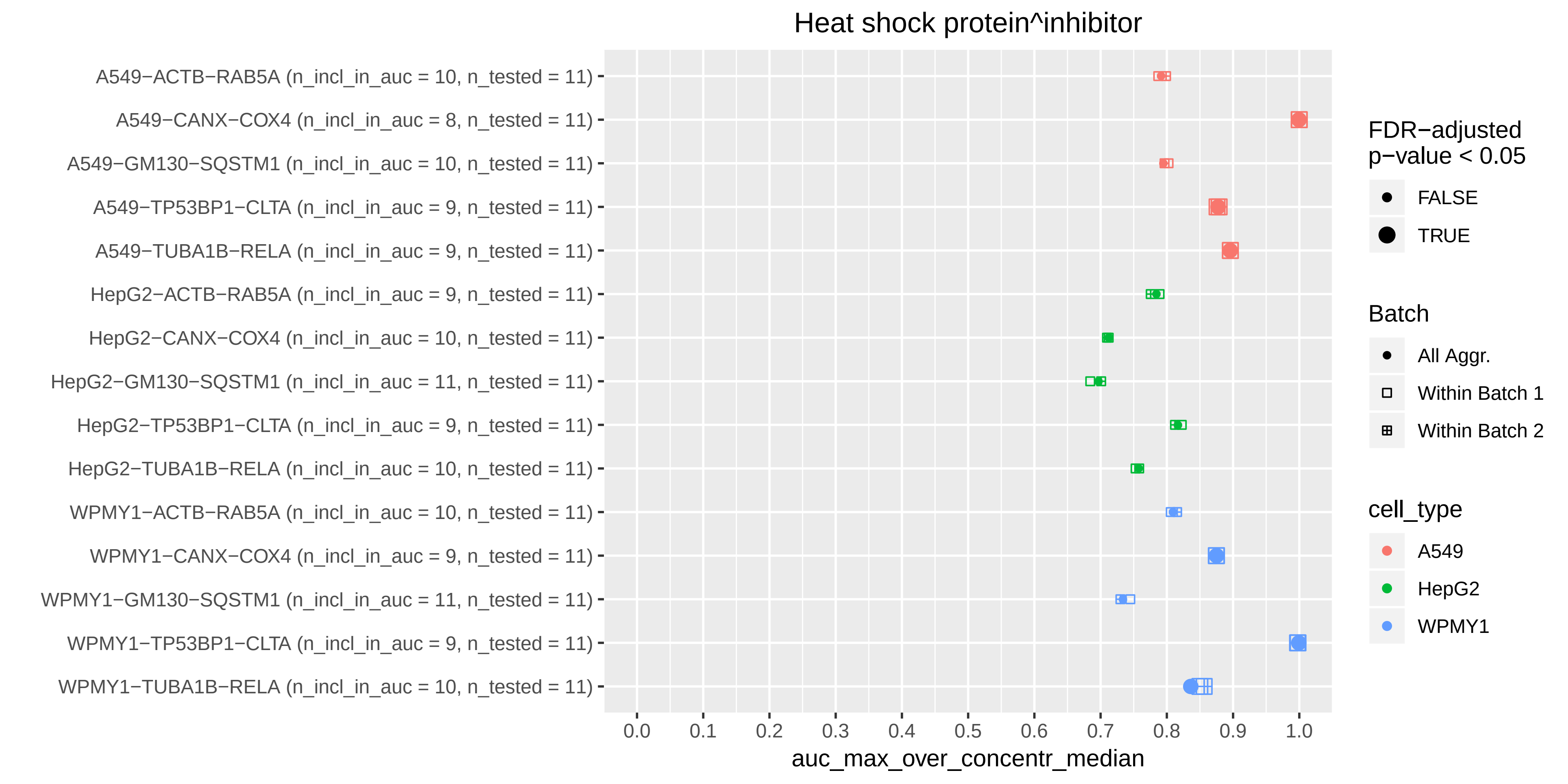

Heat shock protein^inhibitor
A549−ACTB−RAB5A (n_incl_in_auc = 10, n_tested = 11)
A549−CANX−COX4 (n_incl_in_auc = 8, n_tested = 11)
A549−GM130−SQSTM1 (n_incl_in_auc = 10, n_tested = 11)
A549−TP53BP1−CLTA (n_incl_in_auc = 9, n_tested = 11)
A549−TUBA1B−RELA (n_incl_in_auc = 9, n_tested = 11)
HepG2−ACTB−RAB5A (n_incl_in_auc = 9, n_tested = 11)
HepG2−CANX−COX4 (n_incl_in_auc = 10, n_tested = 11)
HepG2−GM130−SQSTM1 (n_incl_in_auc = 11, n_tested = 11)
HepG2−TP53BP1−CLTA (n_incl_in_auc = 9, n_tested = 11)
HepG2−TUBA1B−RELA (n_incl_in_auc = 10, n_tested = 11)
WPMY1−ACTB−RAB5A (n_incl_in_auc = 10, n_tested = 11)
WPMY1−CANX−COX4 (n_incl_in_auc = 9, n_tested = 11)
WPMY1−GM130−SQSTM1 (n_incl_in_auc = 11, n_tested = 11)
WPMY1−TP53BP1−CLTA (n_incl_in_auc = 9, n_tested = 11)
WPMY1−TUBA1B−RELA (n_incl_in_auc = 10, n_tested = 11)
0.0
0.1
0.2
0.3
0.4
0.5
0.6
0.7
0.8
0.9
1.0
auc_max_over_concentr_median
FDR−adjusted p−value < 0.05
FALSE
TRUE
Batch
All Aggr.
Within Batch 1
Within Batch 2
cell_type
A549
HepG2
WPMY1

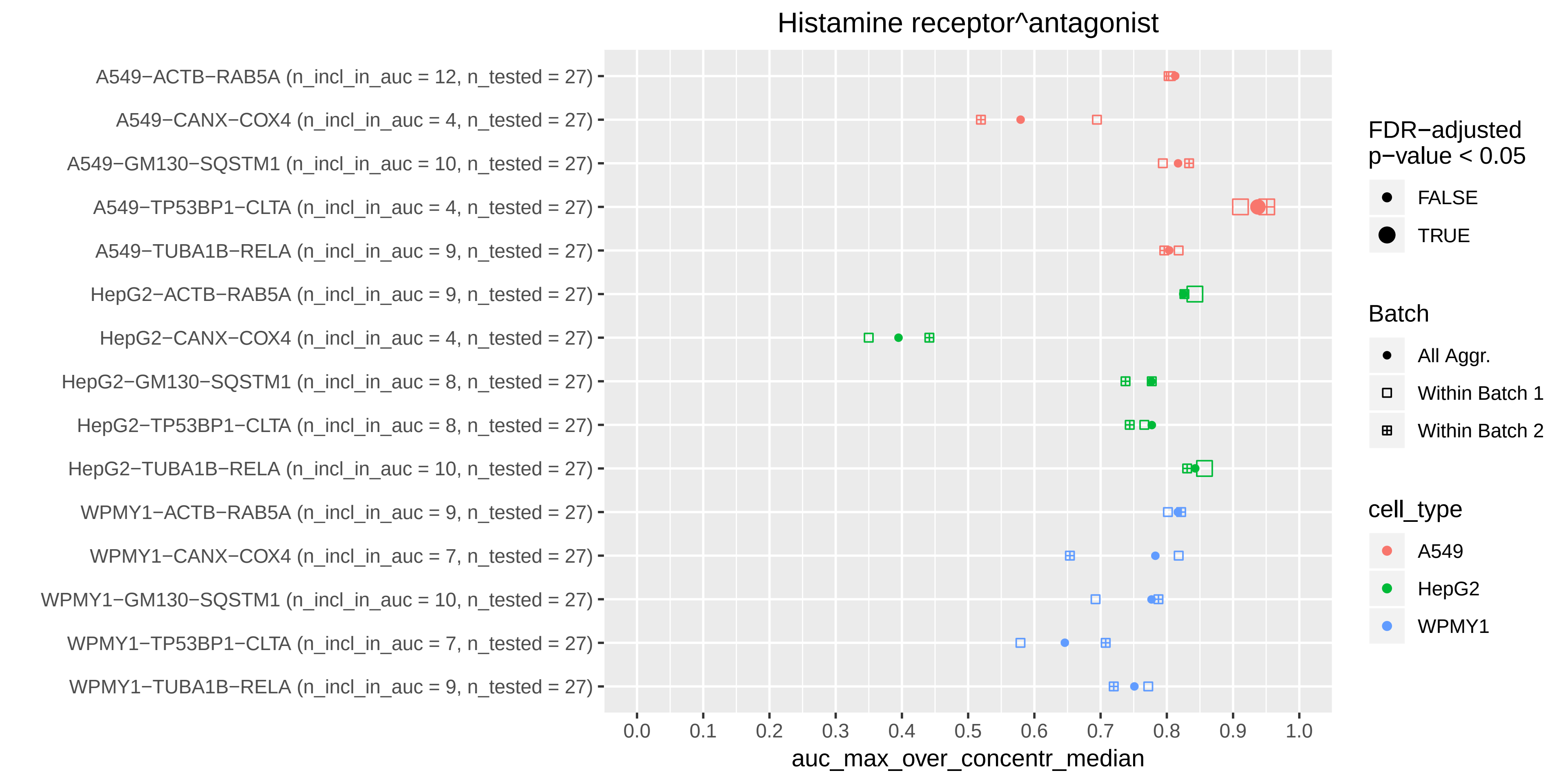

Histamine receptor^antagonist
A549−ACTB−RAB5A (n_incl_in_auc = 12, n_tested = 27)
A549−CANX−COX4 (n_incl_in_auc = 4, n_tested = 27)
A549−GM130−SQSTM1 (n_incl_in_auc = 10, n_tested = 27)
A549−TP53BP1−CLTA (n_incl_in_auc = 4, n_tested = 27)
A549−TUBA1B−RELA (n_incl_in_auc = 9, n_tested = 27)
HepG2−ACTB−RAB5A (n_incl_in_auc = 9, n_tested = 27)
HepG2−CANX−COX4 (n_incl_in_auc = 4, n_tested = 27)
HepG2−GM130−SQSTM1 (n_incl_in_auc = 8, n_tested = 27)
HepG2−TP53BP1−CLTA (n_incl_in_auc = 8, n_tested = 27)
HepG2−TUBA1B−RELA (n_incl_in_auc = 10, n_tested = 27)
WPMY1−ACTB−RAB5A (n_incl_in_auc = 9, n_tested = 27)
WPMY1−CANX−COX4 (n_incl_in_auc = 7, n_tested = 27)
WPMY1−GM130−SQSTM1 (n_incl_in_auc = 10, n_tested = 27)
WPMY1−TP53BP1−CLTA (n_incl_in_auc = 7, n_tested = 27)
WPMY1−TUBA1B−RELA (n_incl_in_auc = 9, n_tested = 27)
0.0
0.1
0.2
0.3
0.4
0.5
0.6
0.7
0.8
0.9
1.0
auc_max_over_concentr_median
FDR−adjusted p−value < 0.05
FALSE
TRUE
Batch
All Aggr.
Within Batch 1
Within Batch 2
cell_type
A549
HepG2
WPMY1

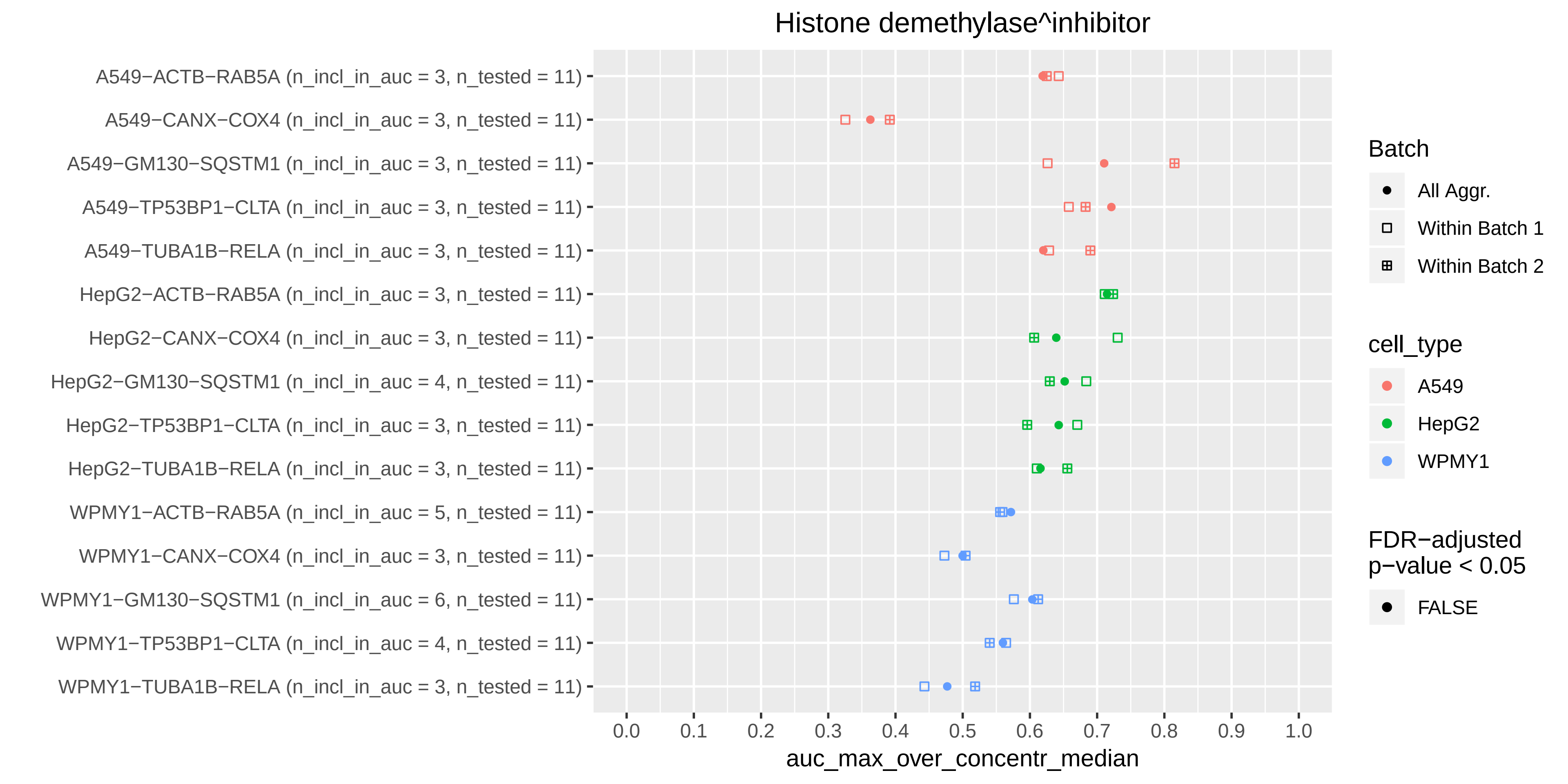

Histone demethylase^inhibitor
A549−ACTB−RAB5A (n_incl_in_auc = 3, n_tested = 11)
A549−CANX−COX4 (n_incl_in_auc = 3, n_tested = 11)
A549−GM130−SQSTM1 (n_incl_in_auc = 3, n_tested = 11)
A549−TP53BP1−CLTA (n_incl_in_auc = 3, n_tested = 11)
A549−TUBA1B−RELA (n_incl_in_auc = 3, n_tested = 11)
HepG2−ACTB−RAB5A (n_incl_in_auc = 3, n_tested = 11)
HepG2−CANX−COX4 (n_incl_in_auc = 3, n_tested = 11)
HepG2−GM130−SQSTM1 (n_incl_in_auc = 4, n_tested = 11)
HepG2−TP53BP1−CLTA (n_incl_in_auc = 3, n_tested = 11)
HepG2−TUBA1B−RELA (n_incl_in_auc = 3, n_tested = 11)
WPMY1−ACTB−RAB5A (n_incl_in_auc = 5, n_tested = 11)
WPMY1−CANX−COX4 (n_incl_in_auc = 3, n_tested = 11)
WPMY1−GM130−SQSTM1 (n_incl_in_auc = 6, n_tested = 11)
WPMY1−TP53BP1−CLTA (n_incl_in_auc = 4, n_tested = 11)
WPMY1−TUBA1B−RELA (n_incl_in_auc = 3, n_tested = 11)
0.0
0.1
0.2
0.3
0.4
0.5
0.6
0.7
0.8
0.9
1.0
auc_max_over_concentr_median
Batch
All Aggr.
Within Batch 1
Within Batch 2
cell_type
A549
HepG2
WPMY1
FDR−adjusted p−value < 0.05
FALSE

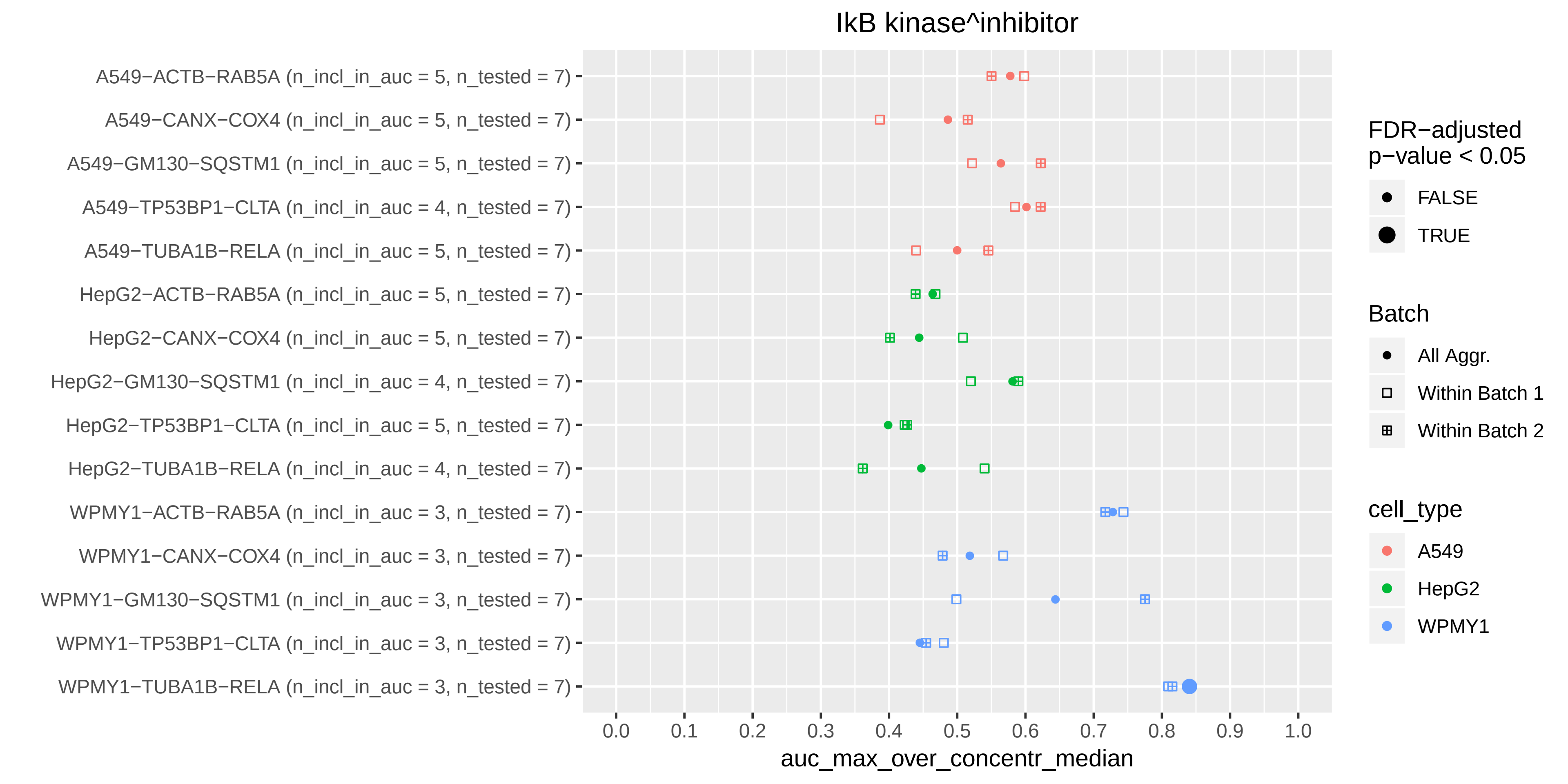

IkB kinase^inhibitor
A549−ACTB−RAB5A (n_incl_in_auc = 5, n_tested = 7)
A549−CANX−COX4 (n_incl_in_auc = 5, n_tested = 7)
A549−GM130−SQSTM1 (n_incl_in_auc = 5, n_tested = 7)
A549−TP53BP1−CLTA (n_incl_in_auc = 4, n_tested = 7)
A549−TUBA1B−RELA (n_incl_in_auc = 5, n_tested = 7)
HepG2−ACTB−RAB5A (n_incl_in_auc = 5, n_tested = 7)
HepG2−CANX−COX4 (n_incl_in_auc = 5, n_tested = 7)
HepG2−GM130−SQSTM1 (n_incl_in_auc = 4, n_tested = 7)
HepG2−TP53BP1−CLTA (n_incl_in_auc = 5, n_tested = 7)
HepG2−TUBA1B−RELA (n_incl_in_auc = 4, n_tested = 7)
WPMY1−ACTB−RAB5A (n_incl_in_auc = 3, n_tested = 7)
WPMY1−CANX−COX4 (n_incl_in_auc = 3, n_tested = 7)
WPMY1−GM130−SQSTM1 (n_incl_in_auc = 3, n_tested = 7)
WPMY1−TP53BP1−CLTA (n_incl_in_auc = 3, n_tested = 7)
WPMY1−TUBA1B−RELA (n_incl_in_auc = 3, n_tested = 7)
0.0 0.1 0.2 0.3 0.4 0.5 0.6 0.7 0.8 0.9 1.0
auc_max_over_concentr_median
FDR−adjusted p−value < 0.05
FALSE
TRUE
Batch
All Aggr.
Within Batch 1
Within Batch 2
cell_type
A549
HepG2
WPMY1

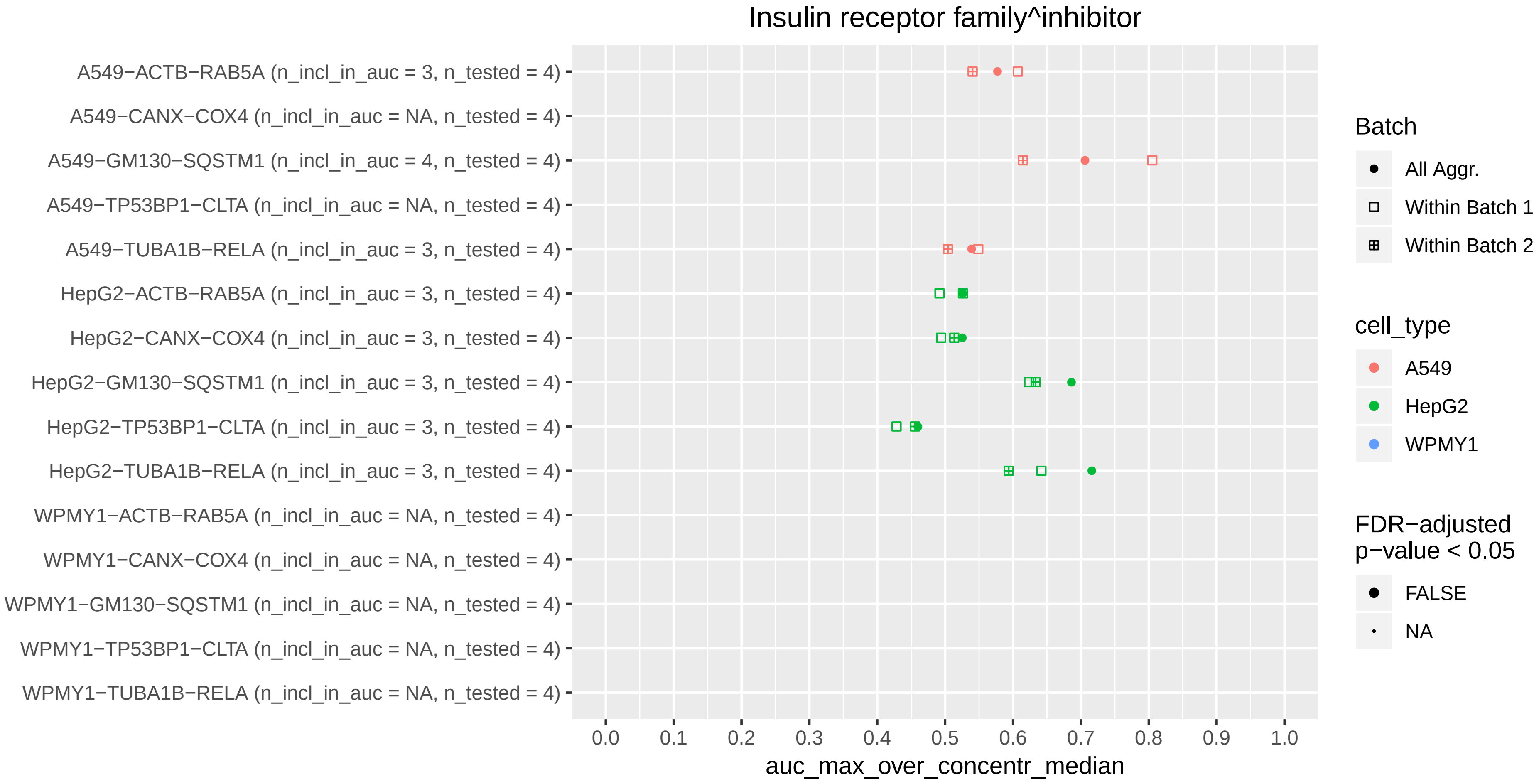

Insulin receptor family^inhibitor
A549−ACTB−RAB5A (n_incl_in_auc = 3, n_tested = 4)
A549−CANX−COX4 (n_incl_in_auc = NA, n_tested = 4)
A549−GM130−SQSTM1 (n_incl_in_auc = 4, n_tested = 4)
A549−TP53BP1−CLTA (n_incl_in_auc = NA, n_tested = 4)
A549−TUBA1B−RELA (n_incl_in_auc = 3, n_tested = 4)
HepG2−ACTB−RAB5A (n_incl_in_auc = 3, n_tested = 4)
HepG2−CANX−COX4 (n_incl_in_auc = 3, n_tested = 4)
HepG2−GM130−SQSTM1 (n_incl_in_auc = 3, n_tested = 4)
HepG2−TP53BP1−CLTA (n_incl_in_auc = 3, n_tested = 4)
HepG2−TUBA1B−RELA (n_incl_in_auc = 3, n_tested = 4)
WPMY1−ACTB−RAB5A (n_incl_in_auc = NA, n_tested = 4)
WPMY1−CANX−COX4 (n_incl_in_auc = NA, n_tested = 4)
WPMY1−GM130−SQSTM1 (n_incl_in_auc = NA, n_tested = 4)
WPMY1−TP53BP1−CLTA (n_incl_in_auc = NA, n_tested = 4)
WPMY1−TUBA1B−RELA (n_incl_in_auc = NA, n_tested = 4)
0.0
0.1
0.2
0.3
0.4
0.5
0.6
0.7
0.8
0.9
1.0
auc_max_over_concentr_median
Batch
All Aggr.
Within Batch 1
Within Batch 2
cell_type
A549
HepG2
WPMY1
FDR−adjusted p−value < 0.05
FALSE
NA

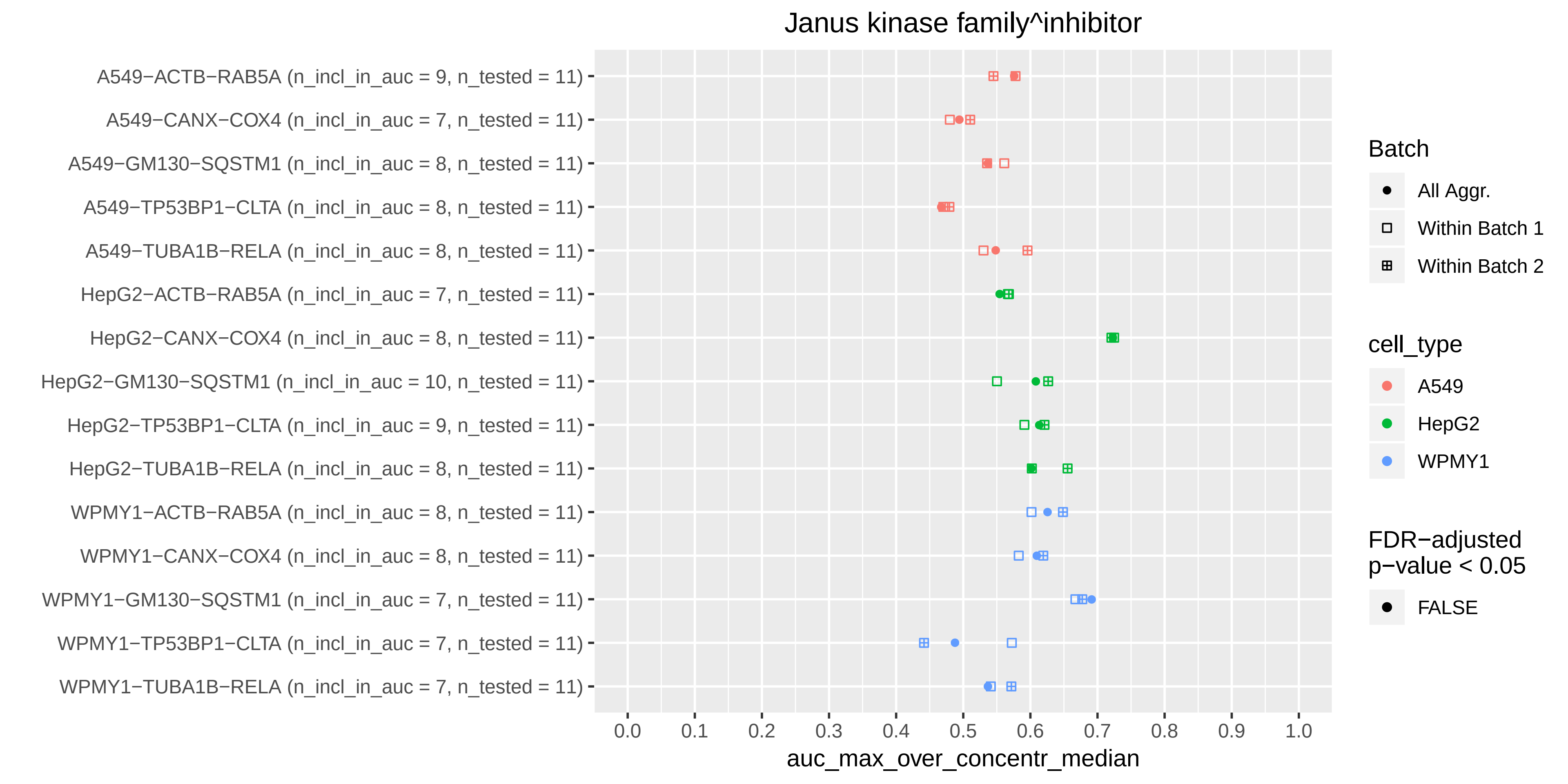

Janus kinase family^inhibitor
A549−ACTB−RAB5A (n_incl_in_auc = 9, n_tested = 11)
A549−CANX−COX4 (n_incl_in_auc = 7, n_tested = 11)
A549−GM130−SQSTM1 (n_incl_in_auc = 8, n_tested = 11)
A549−TP53BP1−CLTA (n_incl_in_auc = 8, n_tested = 11)
A549−TUBA1B−RELA (n_incl_in_auc = 8, n_tested = 11)
HepG2−ACTB−RAB5A (n_incl_in_auc = 7, n_tested = 11)
HepG2−CANX−COX4 (n_incl_in_auc = 8, n_tested = 11)
HepG2−GM130−SQSTM1 (n_incl_in_auc = 10, n_tested = 11)
HepG2−TP53BP1−CLTA (n_incl_in_auc = 9, n_tested = 11)
HepG2−TUBA1B−RELA (n_incl_in_auc = 8, n_tested = 11)
WPMY1−ACTB−RAB5A (n_incl_in_auc = 8, n_tested = 11)
WPMY1−CANX−COX4 (n_incl_in_auc = 8, n_tested = 11)
WPMY1−GM130−SQSTM1 (n_incl_in_auc = 7, n_tested = 11)
WPMY1−TP53BP1−CLTA (n_incl_in_auc = 7, n_tested = 11)
WPMY1−TUBA1B−RELA (n_incl_in_auc = 7, n_tested = 11)
0.0
0.1
0.2
0.3
0.4
0.5
0.6
0.7
0.8
0.9
1.0
auc_max_over_concentr_median
Batch
All Aggr.
Within Batch 1
Within Batch 2
cell_type
A549
HepG2
WPMY1
FDR−adjusted p−value < 0.05
FALSE

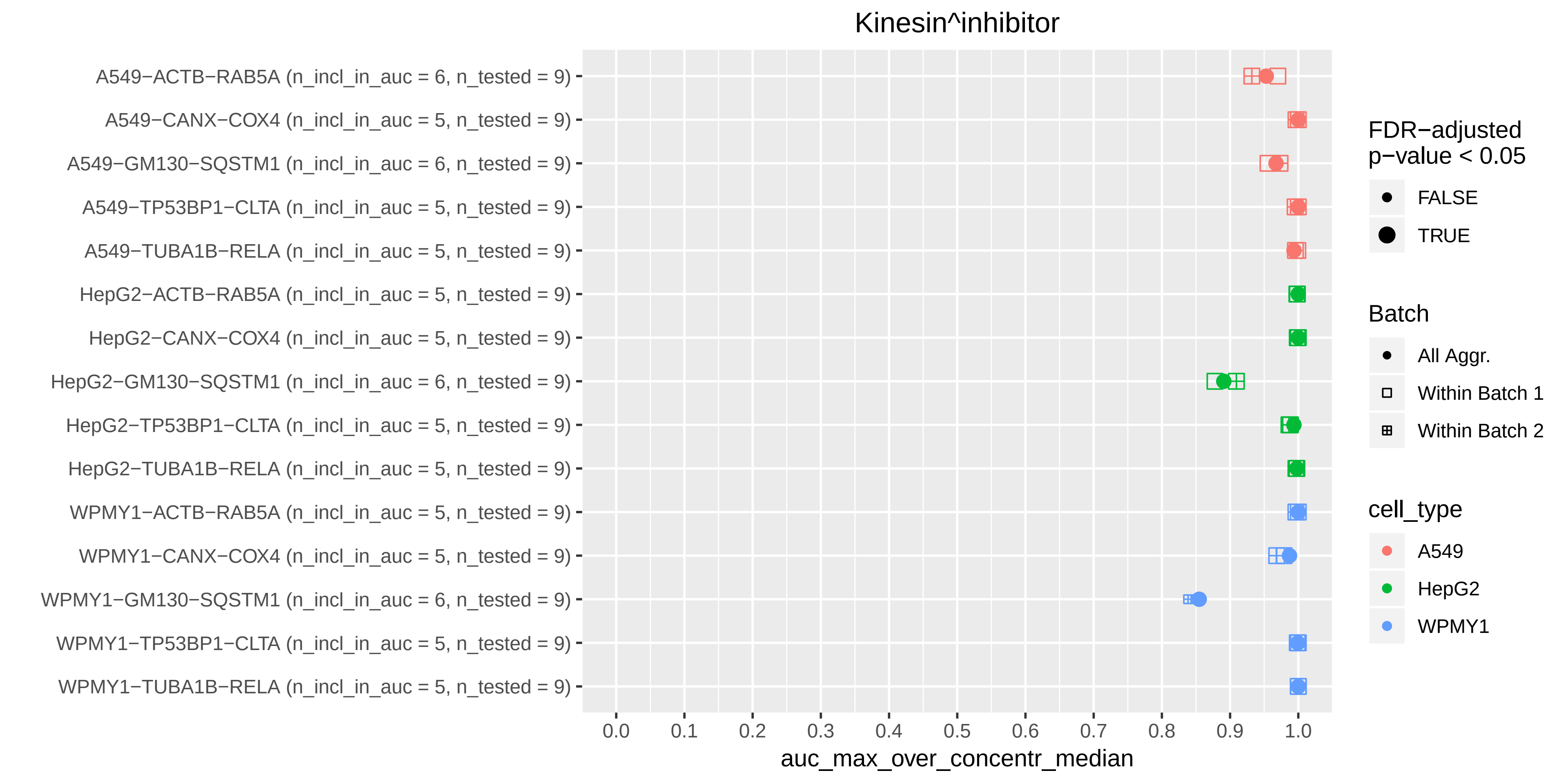

Kinesin^inhibitor
A549−ACTB−RAB5A (n_incl_in_auc = 6, n_tested = 9)
A549−CANX−COX4 (n_incl_in_auc = 5, n_tested = 9)
A549−GM130−SQSTM1 (n_incl_in_auc = 6, n_tested = 9)
A549−TP53BP1−CLTA (n_incl_in_auc = 5, n_tested = 9)
A549−TUBA1B−RELA (n_incl_in_auc = 5, n_tested = 9)
HepG2−ACTB−RAB5A (n_incl_in_auc = 5, n_tested = 9)
HepG2−CANX−COX4 (n_incl_in_auc = 5, n_tested = 9)
HepG2−GM130−SQSTM1 (n_incl_in_auc = 6, n_tested = 9)
HepG2−TP53BP1−CLTA (n_incl_in_auc = 5, n_tested = 9)
HepG2−TUBA1B−RELA (n_incl_in_auc = 5, n_tested = 9)
WPMY1−ACTB−RAB5A (n_incl_in_auc = 5, n_tested = 9)
WPMY1−CANX−COX4 (n_incl_in_auc = 5, n_tested = 9)
WPMY1−GM130−SQSTM1 (n_incl_in_auc = 6, n_tested = 9)
WPMY1−TP53BP1−CLTA (n_incl_in_auc = 5, n_tested = 9)
WPMY1−TUBA1B−RELA (n_incl_in_auc = 5, n_tested = 9)
0.0
0.1
0.2
0.3
0.4
0.5
0.6
0.7
0.8
0.9
1.0
auc_max_over_concentr_median
FDR−adjusted p−value < 0.05
FALSE
TRUE
Batch
All Aggr.
Within Batch 1
Within Batch 2
cell_type
A549
HepG2
WPMY1

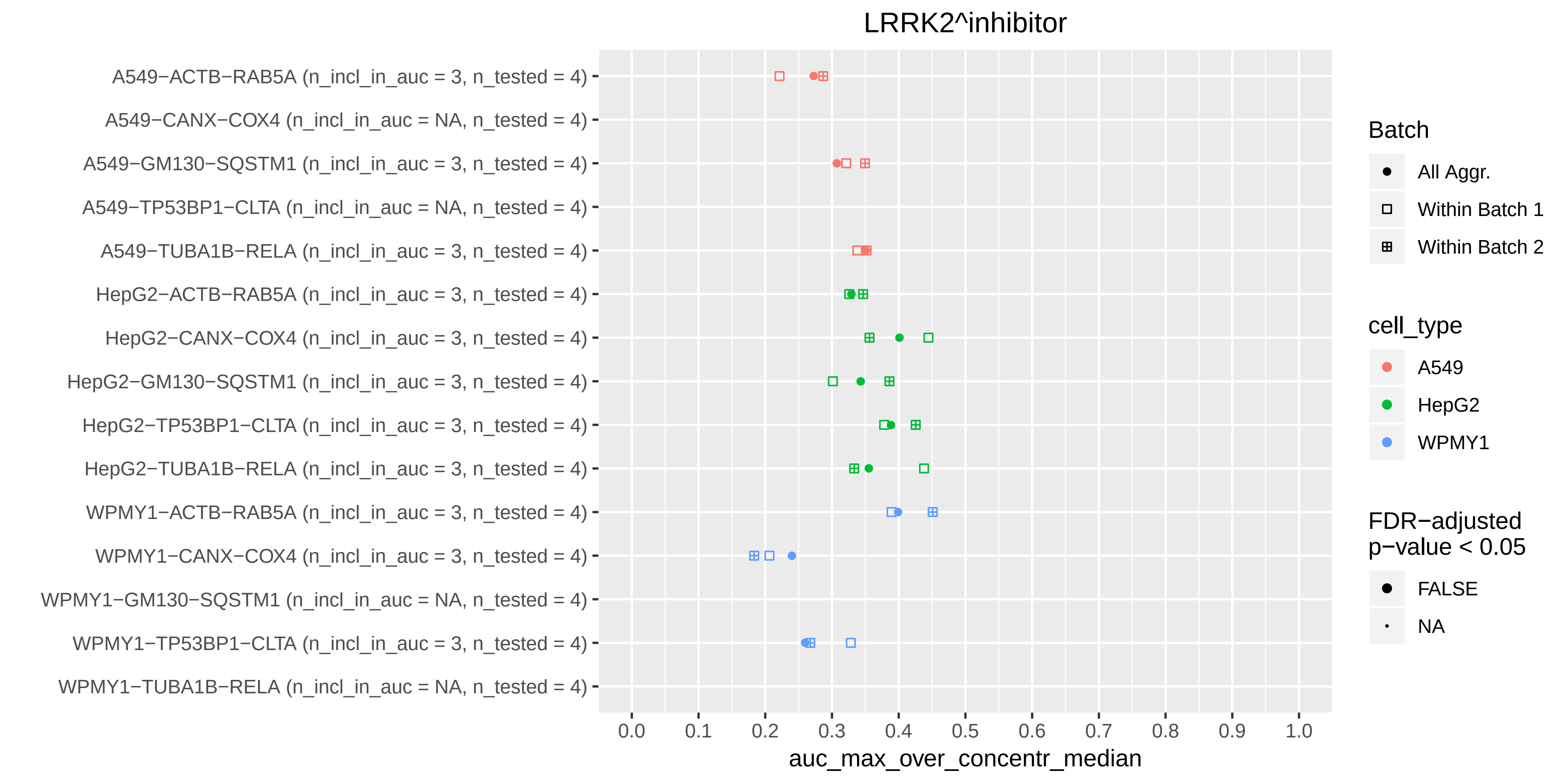
LRRK2^inhibitor
A549−ACTB−RAB5A (n_incl_in_auc = 3, n_tested = 4)
A549−CANX−COX4 (n_incl_in_auc = NA, n_tested = 4)
A549−GM130−SQSTM1 (n_incl_in_auc = 3, n_tested = 4)
A549−TP53BP1−CLTA (n_incl_in_auc = NA, n_tested = 4)
A549−TUBA1B−RELA (n_incl_in_auc = 3, n_tested = 4)
HepG2−ACTB−RAB5A (n_incl_in_auc = 3, n_tested = 4)
HepG2−CANX−COX4 (n_incl_in_auc = 3, n_tested = 4)
HepG2−GM130−SQSTM1 (n_incl_in_auc = 3, n_tested = 4)
HepG2−TP53BP1−CLTA (n_incl_in_auc = 3, n_tested = 4)
HepG2−TUBA1B−RELA (n_incl_in_auc = 3, n_tested = 4)
WPMY1−ACTB−RAB5A (n_incl_in_auc = 3, n_tested = 4)
WPMY1−CANX−COX4 (n_incl_in_auc = 3, n_tested = 4)
WPMY1−GM130−SQSTM1 (n_incl_in_auc = NA, n_tested = 4)
WPMY1−TP53BP1−CLTA (n_incl_in_auc = 3, n_tested = 4)
WPMY1−TUBA1B−RELA (n_incl_in_auc = NA, n_tested = 4)
0.0
0.1
0.2
0.3
0.4
0.5
0.6
0.7
0.8
0.9
1.0
auc_max_over_concentr_median
Batch
All Aggr.
Within Batch 1
Within Batch 2
cell_type
A549
HepG2
WPMY1
FDR−adjusted
p−value < 0.05
FALSE
NA

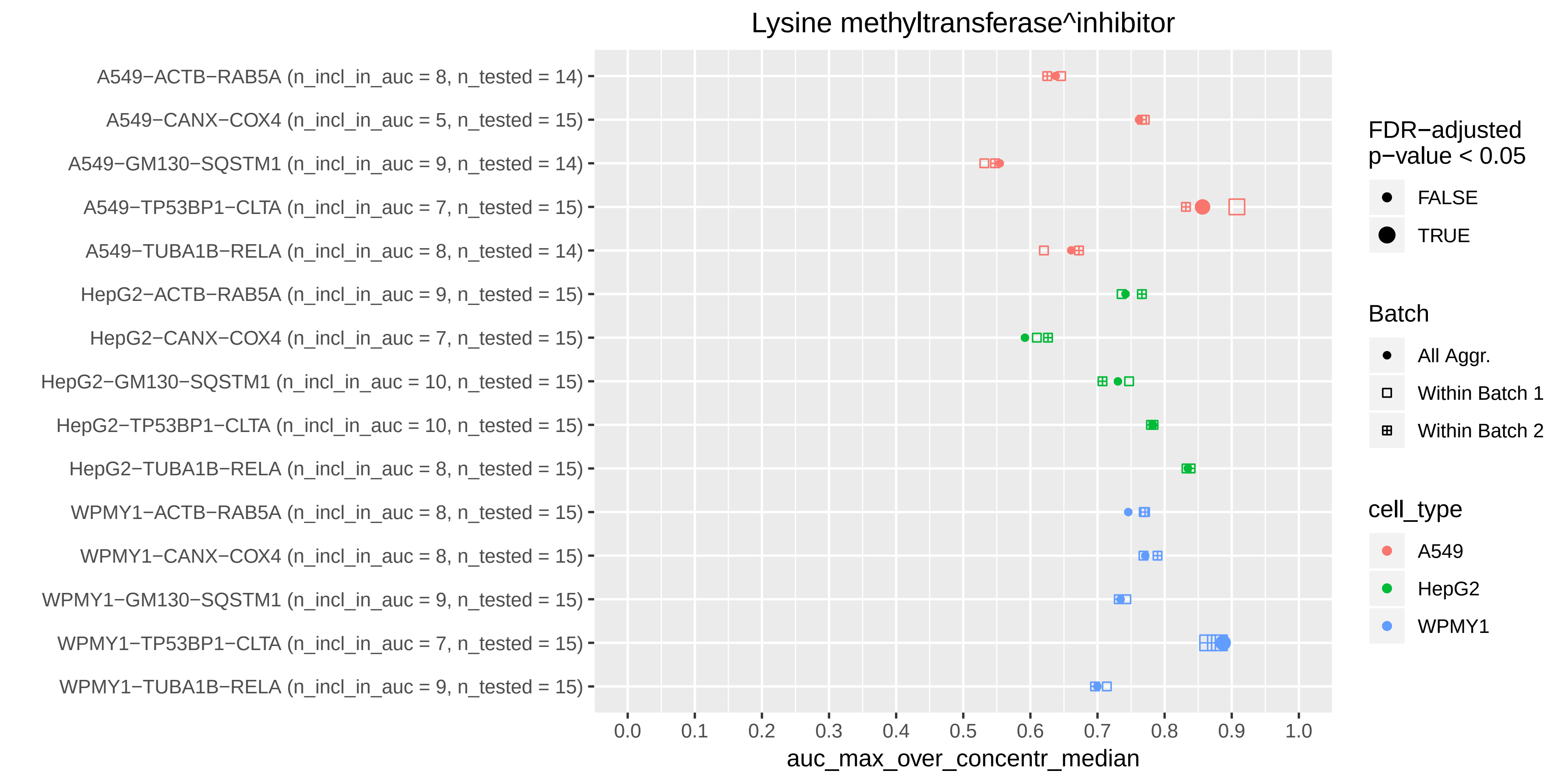

Lysine methyltransferase^inhibitor
A549−ACTB−RAB5A (n_incl_in_auc = 8, n_tested = 14)
A549−CANX−COX4 (n_incl_in_auc = 5, n_tested = 15)
A549−GM130−SQSTM1 (n_incl_in_auc = 9, n_tested = 14)
A549−TP53BP1−CLTA (n_incl_in_auc = 7, n_tested = 15)
A549−TUBA1B−RELA (n_incl_in_auc = 8, n_tested = 14)
HepG2−ACTB−RAB5A (n_incl_in_auc = 9, n_tested = 15)
HepG2−CANX−COX4 (n_incl_in_auc = 7, n_tested = 15)
HepG2−GM130−SQSTM1 (n_incl_in_auc = 10, n_tested = 15)
HepG2−TP53BP1−CLTA (n_incl_in_auc = 10, n_tested = 15)
HepG2−TUBA1B−RELA (n_incl_in_auc = 8, n_tested = 15)
WPMY1−ACTB−RAB5A (n_incl_in_auc = 8, n_tested = 15)
WPMY1−CANX−COX4 (n_incl_in_auc = 8, n_tested = 15)
WPMY1−GM130−SQSTM1 (n_incl_in_auc = 9, n_tested = 15)
WPMY1−TP53BP1−CLTA (n_incl_in_auc = 7, n_tested = 15)
WPMY1−TUBA1B−RELA (n_incl_in_auc = 9, n_tested = 15)
0.0
0.1
0.2
0.3
0.4
0.5
0.6
0.7
0.8
0.9
1.0
auc_max_over_concentr_median
FDR−adjusted p−value < 0.05
FALSE
TRUE
Batch
All Aggr.
Within Batch 1
Within Batch 2
cell_type
A549
HepG2
WPMY1

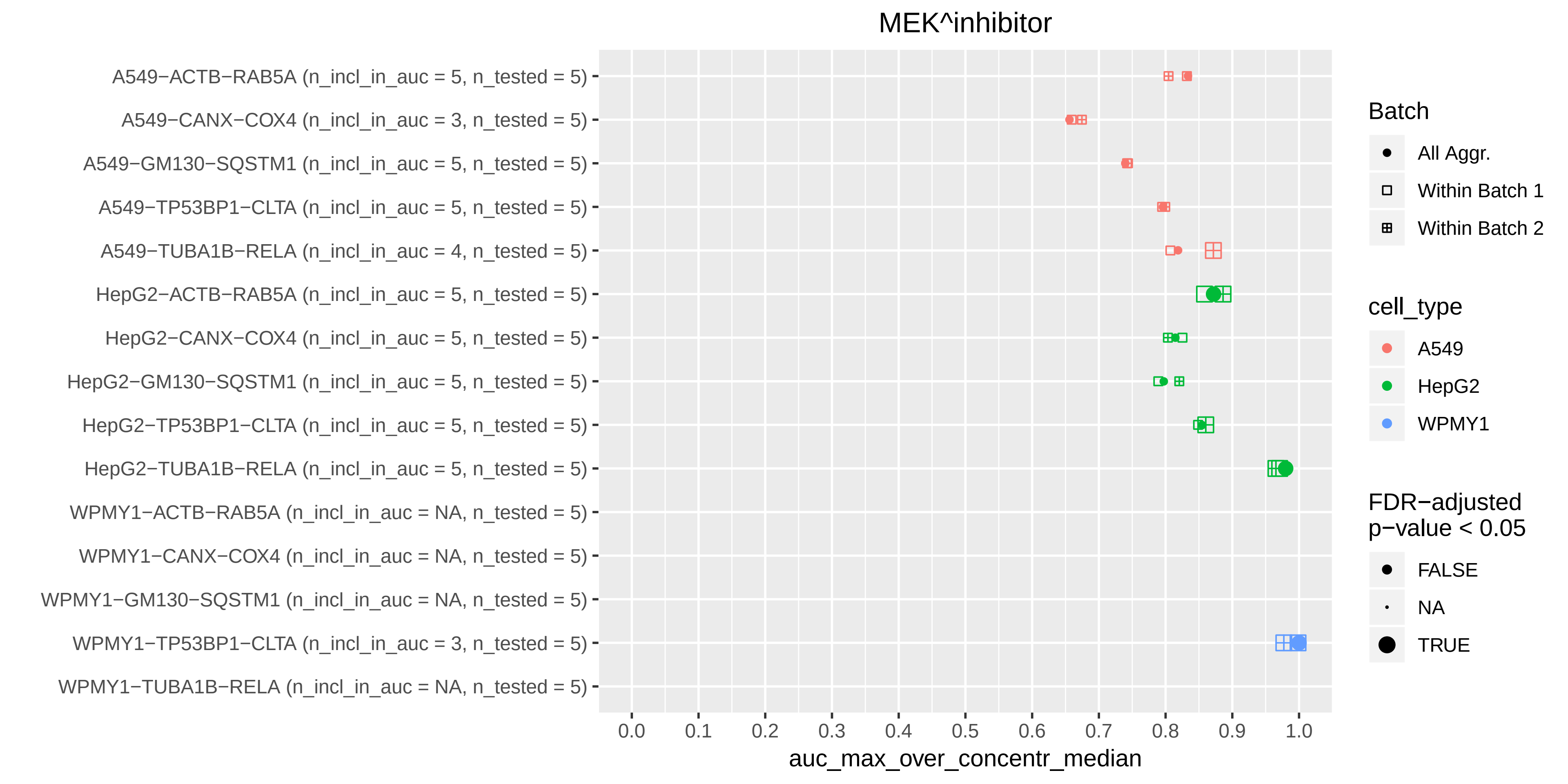

MEK^inhibitor
A549−ACTB−RAB5A (n_incl_in_auc = 5, n_tested = 5)
A549−CANX−COX4 (n_incl_in_auc = 3, n_tested = 5)
A549−GM130−SQSTM1 (n_incl_in_auc = 5, n_tested = 5)
A549−TP53BP1−CLTA (n_incl_in_auc = 5, n_tested = 5)
A549−TUBA1B−RELA (n_incl_in_auc = 4, n_tested = 5)
HepG2−ACTB−RAB5A (n_incl_in_auc = 5, n_tested = 5)
HepG2−CANX−COX4 (n_incl_in_auc = 5, n_tested = 5)
HepG2−GM130−SQSTM1 (n_incl_in_auc = 5, n_tested = 5)
HepG2−TP53BP1−CLTA (n_incl_in_auc = 5, n_tested = 5)
HepG2−TUBA1B−RELA (n_incl_in_auc = 5, n_tested = 5)
WPMY1−ACTB−RAB5A (n_incl_in_auc = NA, n_tested = 5)
WPMY1−CANX−COX4 (n_incl_in_auc = NA, n_tested = 5)
WPMY1−GM130−SQSTM1 (n_incl_in_auc = NA, n_tested = 5)
WPMY1−TP53BP1−CLTA (n_incl_in_auc = 3, n_tested = 5)
WPMY1−TUBA1B−RELA (n_incl_in_auc = NA, n_tested = 5)
0.0 0.1 0.2 0.3 0.4 0.5 0.6 0.7 0.8 0.9 1.0
auc_max_over_concentr_median
Batch
All Aggr.
Within Batch 1
Within Batch 2
cell_type
A549
HepG2
WPMY1
FDR−adjusted p−value < 0.05
FALSE
NA
TRUE

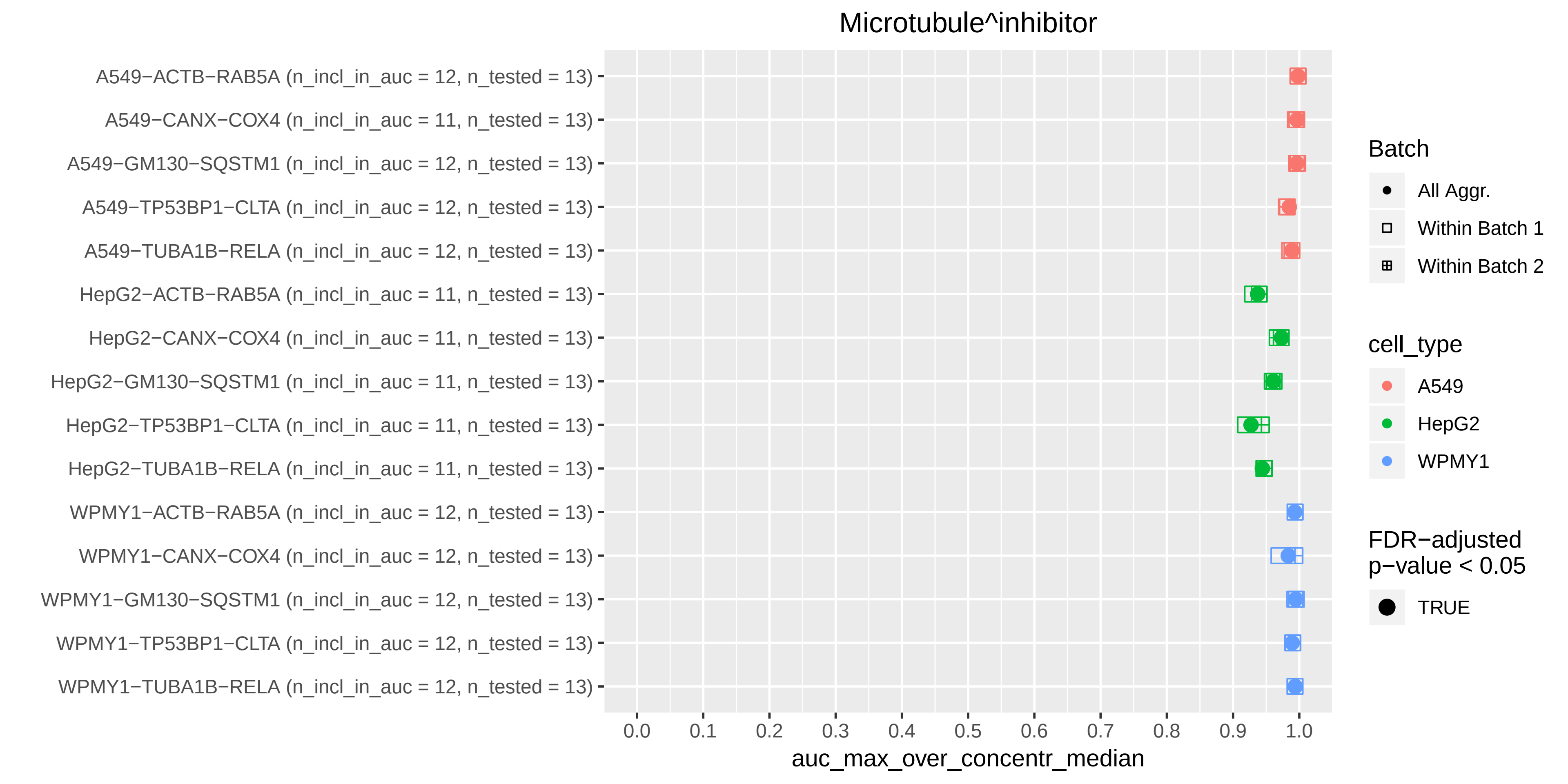
Microtubule^inhibitor
A549−ACTB−RAB5A (n_incl_in_auc = 12, n_tested = 13)
A549−CANX−COX4 (n_incl_in_auc = 11, n_tested = 13)
A549−GM130−SQSTM1 (n_incl_in_auc = 12, n_tested = 13)
A549−TP53BP1−CLTA (n_incl_in_auc = 12, n_tested = 13)
A549−TUBA1B−RELA (n_incl_in_auc = 12, n_tested = 13)
HepG2−ACTB−RAB5A (n_incl_in_auc = 11, n_tested = 13)
HepG2−CANX−COX4 (n_incl_in_auc = 11, n_tested = 13)
HepG2−GM130−SQSTM1 (n_incl_in_auc = 11, n_tested = 13)
HepG2−TP53BP1−CLTA (n_incl_in_auc = 11, n_tested = 13)
HepG2−TUBA1B−RELA (n_incl_in_auc = 11, n_tested = 13)
WPMY1−ACTB−RAB5A (n_incl_in_auc = 12, n_tested = 13)
WPMY1−CANX−COX4 (n_incl_in_auc = 12, n_tested = 13)
WPMY1−GM130−SQSTM1 (n_incl_in_auc = 12, n_tested = 13)
WPMY1−TP53BP1−CLTA (n_incl_in_auc = 12, n_tested = 13)
WPMY1−TUBA1B−RELA (n_incl_in_auc = 12, n_tested = 13)
0.0
0.1
0.2
0.3
0.4
0.5
0.6
0.7
0.8
0.9
1.0
auc_max_over_concentr_median
Batch
All Aggr.
Within Batch 1
Within Batch 2
cell_type
A549
HepG2
WPMY1
FDR−adjusted
p−value < 0.05
TRUE

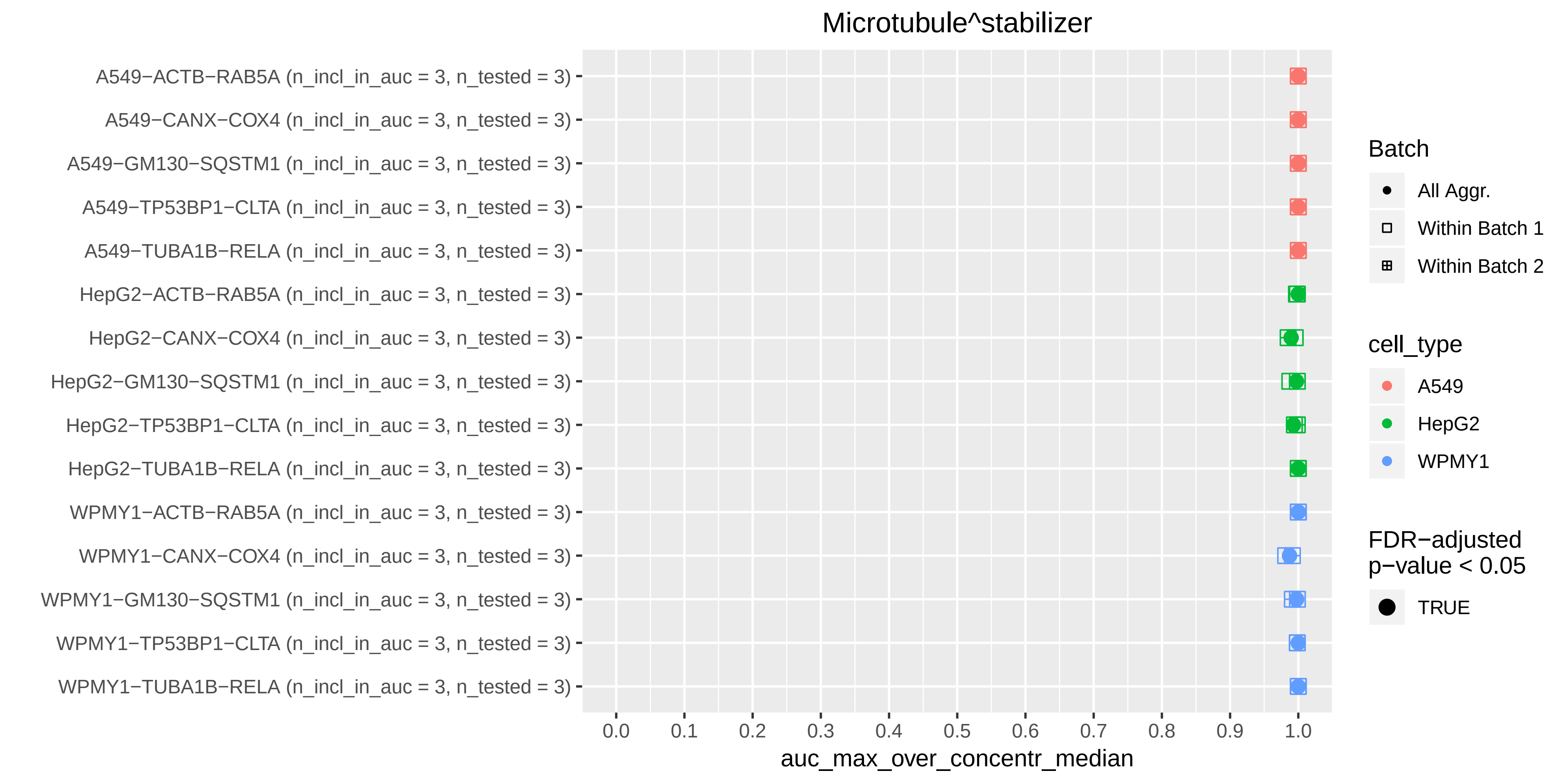
Microtubule^stabilizer
A549−ACTB−RAB5A (n_incl_in_auc = 3, n_tested = 3)
A549−CANX−COX4 (n_incl_in_auc = 3, n_tested = 3)
A549−GM130−SQSTM1 (n_incl_in_auc = 3, n_tested = 3)
A549−TP53BP1−CLTA (n_incl_in_auc = 3, n_tested = 3)
A549−TUBA1B−RELA (n_incl_in_auc = 3, n_tested = 3)
HepG2−ACTB−RAB5A (n_incl_in_auc = 3, n_tested = 3)
HepG2−CANX−COX4 (n_incl_in_auc = 3, n_tested = 3)
HepG2−GM130−SQSTM1 (n_incl_in_auc = 3, n_tested = 3)
HepG2−TP53BP1−CLTA (n_incl_in_auc = 3, n_tested = 3)
HepG2−TUBA1B−RELA (n_incl_in_auc = 3, n_tested = 3)
WPMY1−ACTB−RAB5A (n_incl_in_auc = 3, n_tested = 3)
WPMY1−CANX−COX4 (n_incl_in_auc = 3, n_tested = 3)
WPMY1−GM130−SQSTM1 (n_incl_in_auc = 3, n_tested = 3)
WPMY1−TP53BP1−CLTA (n_incl_in_auc = 3, n_tested = 3)
WPMY1−TUBA1B−RELA (n_incl_in_auc = 3, n_tested = 3)
0.0
0.1
0.2
0.3
0.4
0.5
0.6
0.7
0.8
0.9
1.0
auc_max_over_concentr_median
Batch
All Aggr.
Within Batch 1
Within Batch 2
cell_type
A549
HepG2
WPMY1
FDR−adjusted p−value < 0.05
TRUE

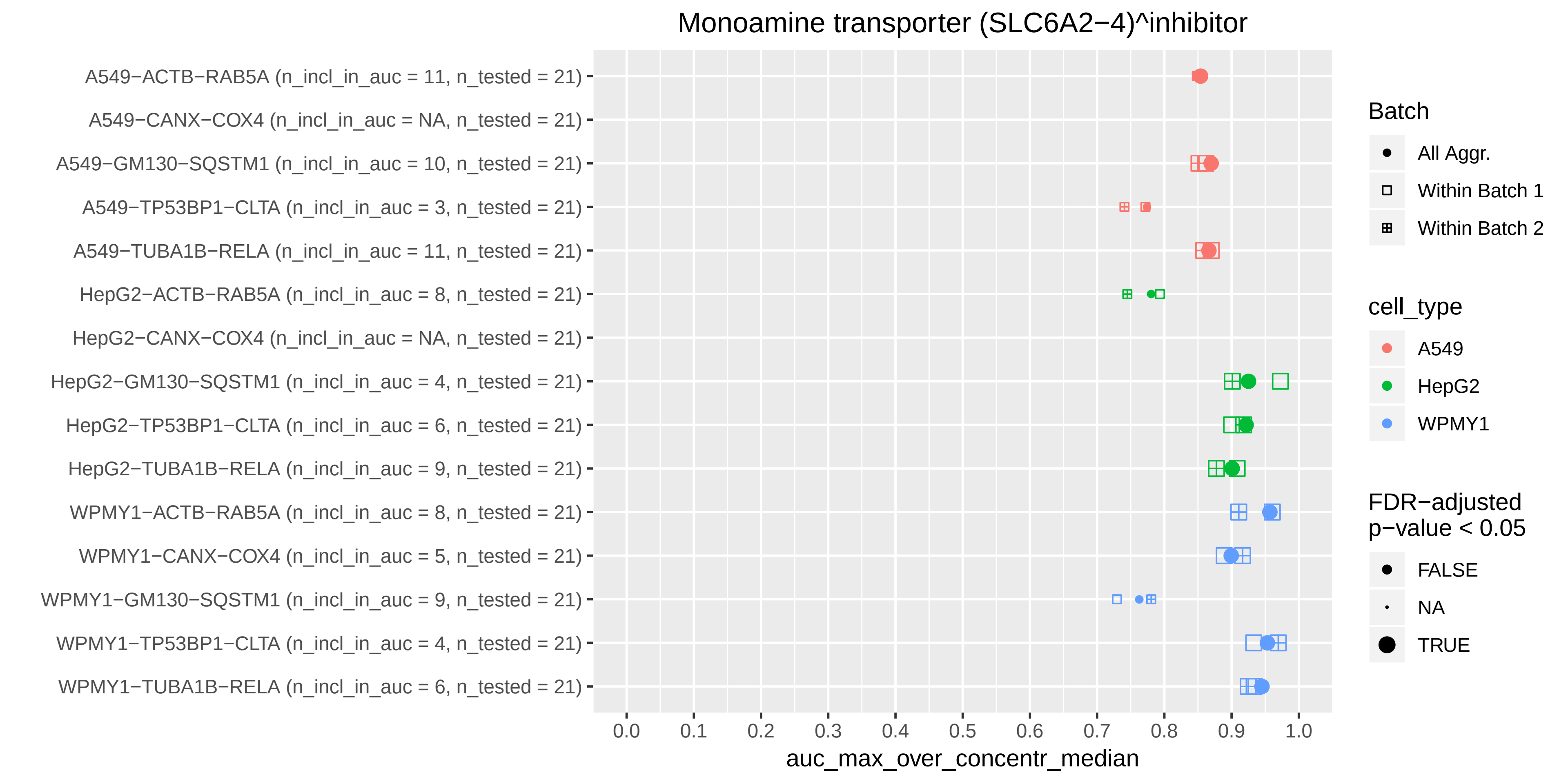
Monoamine transporter (SLC6A2−4)^inhibitor
A549−ACTB−RAB5A (n_incl_in_auc = 11, n_tested = 21)
A549−CANX−COX4 (n_incl_in_auc = NA, n_tested = 21)
A549−GM130−SQSTM1 (n_incl_in_auc = 10, n_tested = 21)
A549−TP53BP1−CLTA (n_incl_in_auc = 3, n_tested = 21)
A549−TUBA1B−RELA (n_incl_in_auc = 11, n_tested = 21)
HepG2−ACTB−RAB5A (n_incl_in_auc = 8, n_tested = 21)
HepG2−CANX−COX4 (n_incl_in_auc = NA, n_tested = 21)
HepG2−GM130−SQSTM1 (n_incl_in_auc = 4, n_tested = 21)
HepG2−TP53BP1−CLTA (n_incl_in_auc = 6, n_tested = 21)
HepG2−TUBA1B−RELA (n_incl_in_auc = 9, n_tested = 21)
WPMY1−ACTB−RAB5A (n_incl_in_auc = 8, n_tested = 21)
WPMY1−CANX−COX4 (n_incl_in_auc = 5, n_tested = 21)
WPMY1−GM130−SQSTM1 (n_incl_in_auc = 9, n_tested = 21)
WPMY1−TP53BP1−CLTA (n_incl_in_auc = 4, n_tested = 21)
WPMY1−TUBA1B−RELA (n_incl_in_auc = 6, n_tested = 21)
0.0
0.1
0.2
0.3
0.4
0.5
0.6
0.7
0.8
0.9
1.0
auc_max_over_concentr_median
Batch
All Aggr.
Within Batch 1
Within Batch 2
cell_type
A549
HepG2
WPMY1
FDR−adjusted p−value < 0.05
FALSE
NA
TRUE

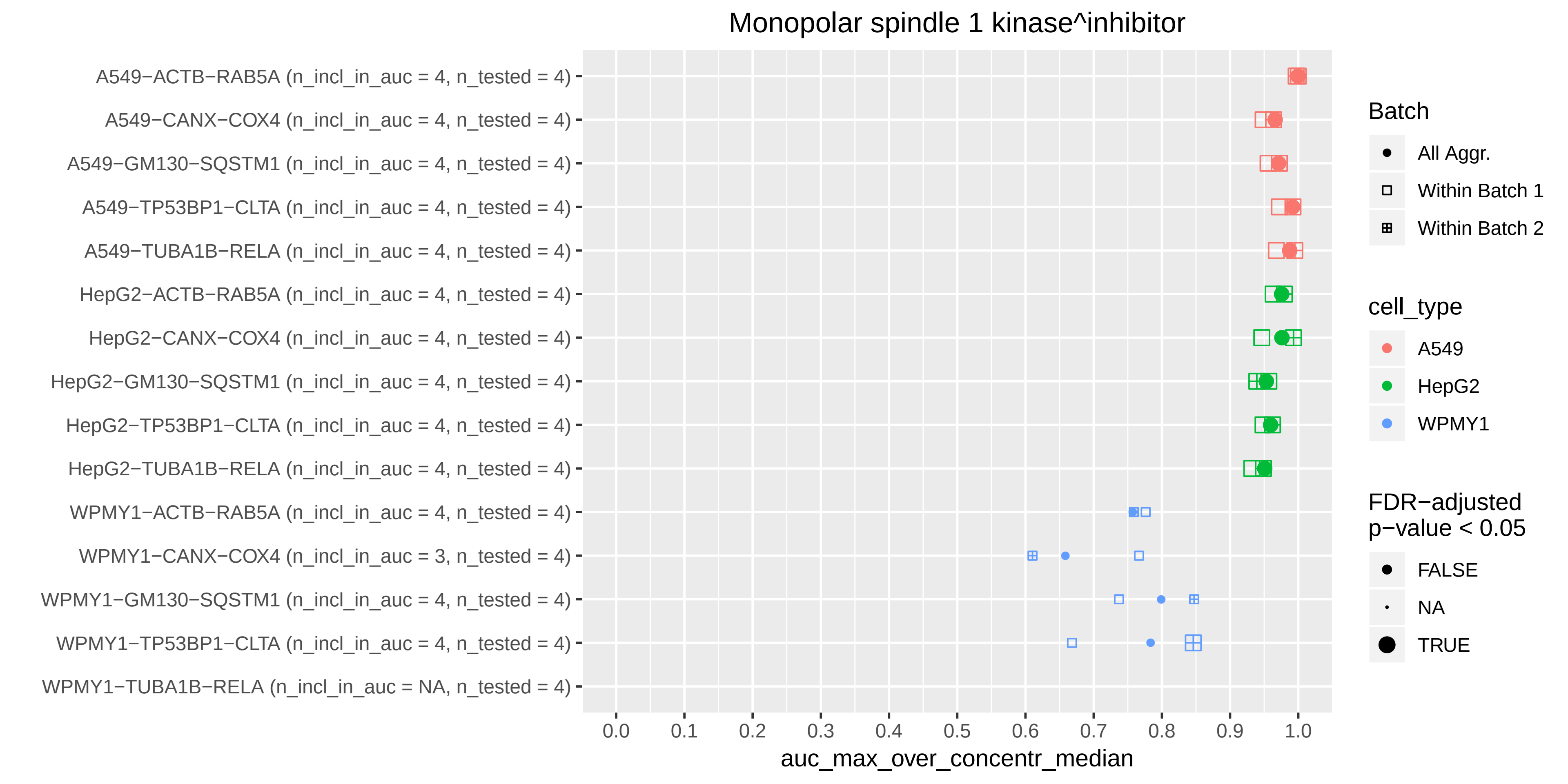

Monopolar spindle 1 kinase^inhibitor
A549−ACTB−RAB5A (n_incl_in_auc = 4, n_tested = 4)
A549−CANX−COX4 (n_incl_in_auc = 4, n_tested = 4)
A549−GM130−SQSTM1 (n_incl_in_auc = 4, n_tested = 4)
A549−TP53BP1−CLTA (n_incl_in_auc = 4, n_tested = 4)
A549−TUBA1B−RELA (n_incl_in_auc = 4, n_tested = 4)
HepG2−ACTB−RAB5A (n_incl_in_auc = 4, n_tested = 4)
HepG2−CANX−COX4 (n_incl_in_auc = 4, n_tested = 4)
HepG2−GM130−SQSTM1 (n_incl_in_auc = 4, n_tested = 4)
HepG2−TP53BP1−CLTA (n_incl_in_auc = 4, n_tested = 4)
HepG2−TUBA1B−RELA (n_incl_in_auc = 4, n_tested = 4)
WPMY1−ACTB−RAB5A (n_incl_in_auc = 4, n_tested = 4)
WPMY1−CANX−COX4 (n_incl_in_auc = 3, n_tested = 4)
WPMY1−GM130−SQSTM1 (n_incl_in_auc = 4, n_tested = 4)
WPMY1−TP53BP1−CLTA (n_incl_in_auc = 4, n_tested = 4)
WPMY1−TUBA1B−RELA (n_incl_in_auc = NA, n_tested = 4)
0.0
0.1
0.2
0.3
0.4
0.5
0.6
0.7
0.8
0.9
1.0
auc_max_over_concentr_median
Batch
All Aggr.
Within Batch 1
Within Batch 2
cell_type
A549
HepG2
WPMY1
FDR−adjusted p−value < 0.05
FALSE
NA
TRUE

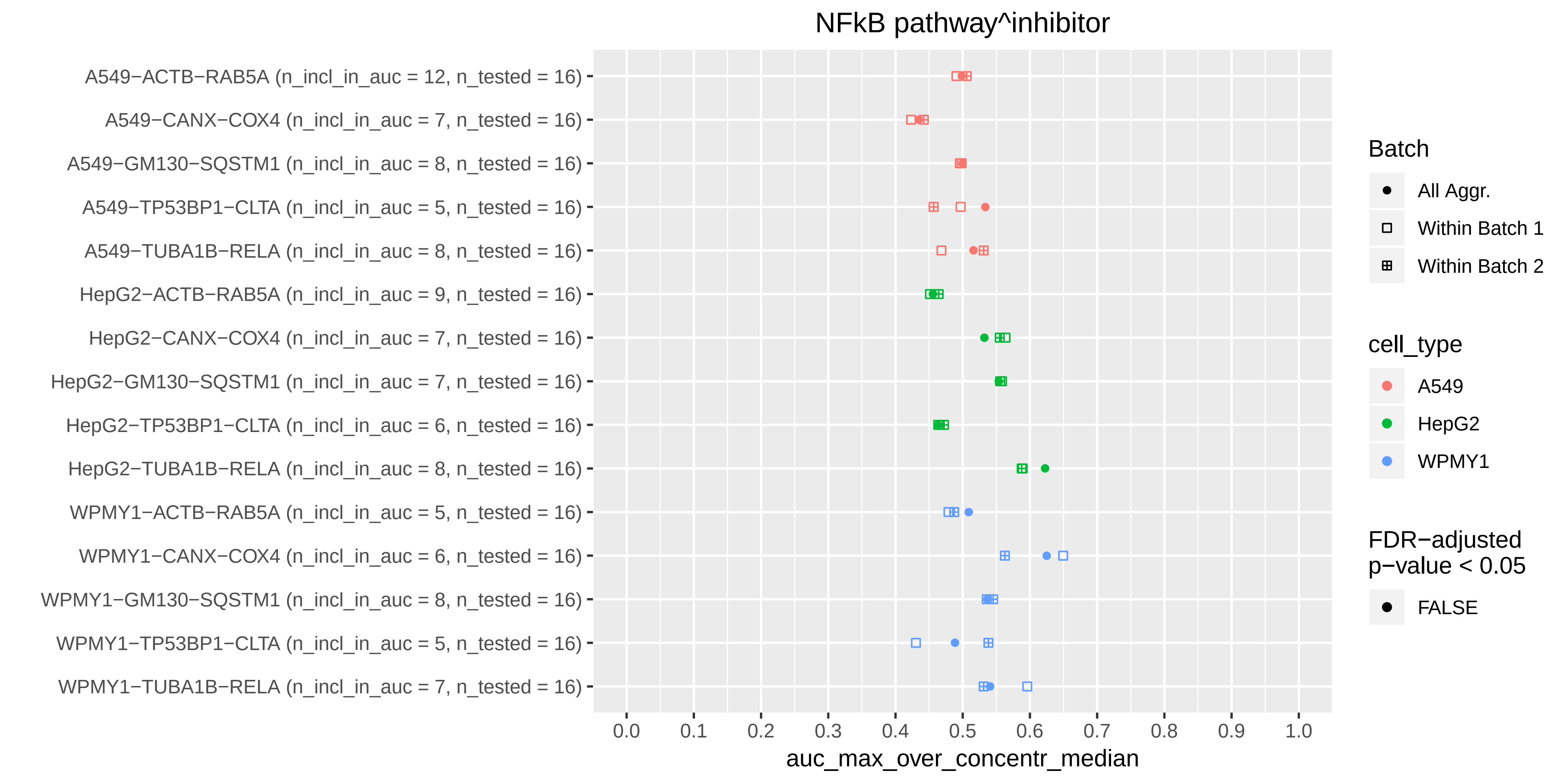

NFkB pathway^inhibitor
A549−ACTB−RAB5A (n_incl_in_auc = 12, n_tested = 16)
A549−CANX−COX4 (n_incl_in_auc = 7, n_tested = 16)
A549−GM130−SQSTM1 (n_incl_in_auc = 8, n_tested = 16)
A549−TP53BP1−CLTA (n_incl_in_auc = 5, n_tested = 16)
A549−TUBA1B−RELA (n_incl_in_auc = 8, n_tested = 16)
HepG2−ACTB−RAB5A (n_incl_in_auc = 9, n_tested = 16)
HepG2−CANX−COX4 (n_incl_in_auc = 7, n_tested = 16)
HepG2−GM130−SQSTM1 (n_incl_in_auc = 7, n_tested = 16)
HepG2−TP53BP1−CLTA (n_incl_in_auc = 6, n_tested = 16)
HepG2−TUBA1B−RELA (n_incl_in_auc = 8, n_tested = 16)
WPMY1−ACTB−RAB5A (n_incl_in_auc = 5, n_tested = 16)
WPMY1−CANX−COX4 (n_incl_in_auc = 6, n_tested = 16)
WPMY1−GM130−SQSTM1 (n_incl_in_auc = 8, n_tested = 16)
WPMY1−TP53BP1−CLTA (n_incl_in_auc = 5, n_tested = 16)
WPMY1−TUBA1B−RELA (n_incl_in_auc = 7, n_tested = 16)
0.0
0.1
0.2
0.3
0.4
0.5
0.6
0.7
0.8
0.9
1.0
auc_max_over_concentr_median
Batch
All Aggr.
Within Batch 1
Within Batch 2
cell_type
A549
HepG2
WPMY1
FDR−adjusted p−value < 0.05
FALSE

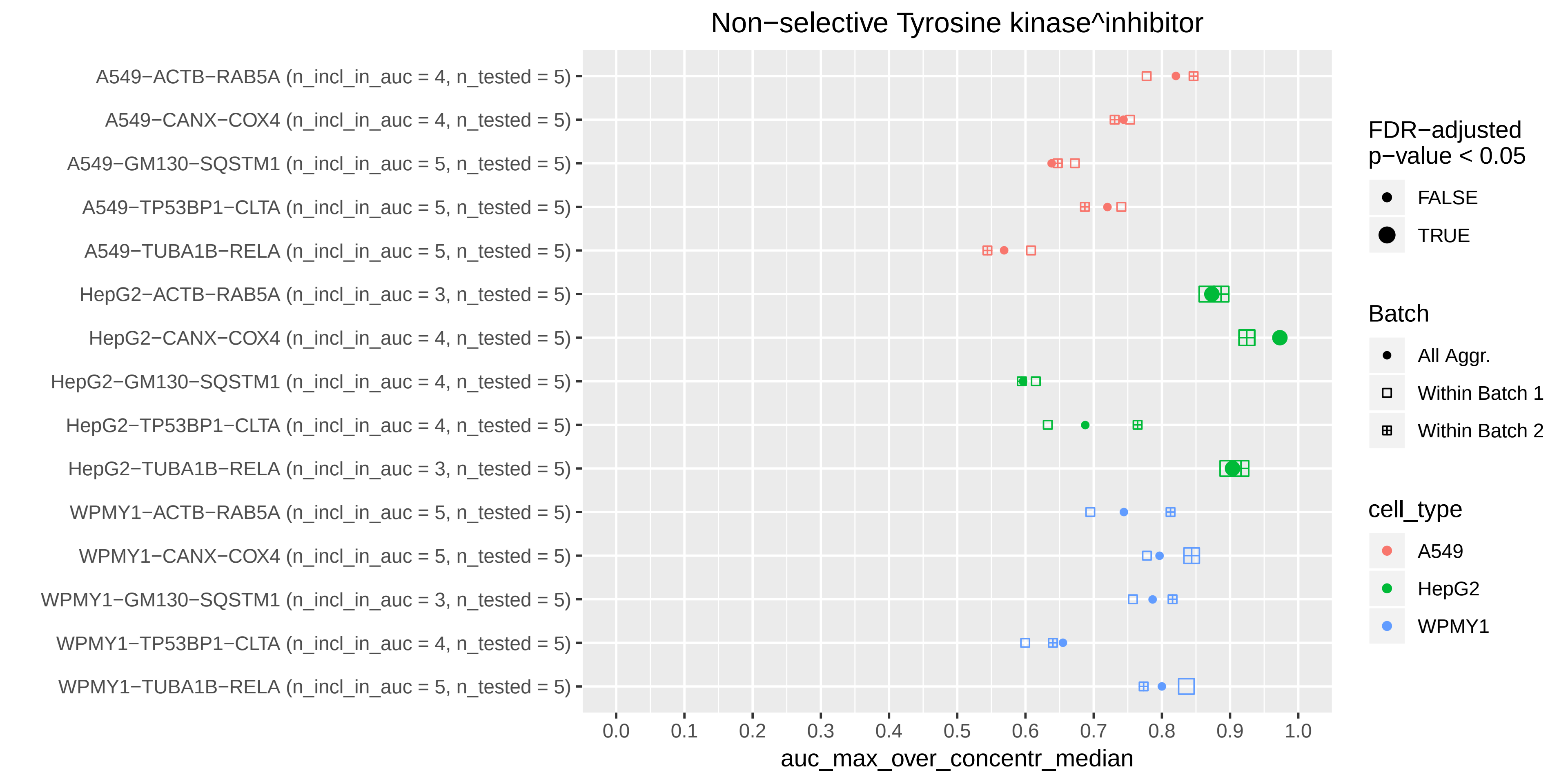

Non−selective Tyrosine kinase^inhibitor
A549−ACTB−RAB5A (n_incl_in_auc = 4, n_tested = 5)
A549−CANX−COX4 (n_incl_in_auc = 4, n_tested = 5)
A549−GM130−SQSTM1 (n_incl_in_auc = 5, n_tested = 5)
A549−TP53BP1−CLTA (n_incl_in_auc = 5, n_tested = 5)
A549−TUBA1B−RELA (n_incl_in_auc = 5, n_tested = 5)
HepG2−ACTB−RAB5A (n_incl_in_auc = 3, n_tested = 5)
HepG2−CANX−COX4 (n_incl_in_auc = 4, n_tested = 5)
HepG2−GM130−SQSTM1 (n_incl_in_auc = 4, n_tested = 5)
HepG2−TP53BP1−CLTA (n_incl_in_auc = 4, n_tested = 5)
HepG2−TUBA1B−RELA (n_incl_in_auc = 3, n_tested = 5)
WPMY1−ACTB−RAB5A (n_incl_in_auc = 5, n_tested = 5)
WPMY1−CANX−COX4 (n_incl_in_auc = 5, n_tested = 5)
WPMY1−GM130−SQSTM1 (n_incl_in_auc = 3, n_tested = 5)
WPMY1−TP53BP1−CLTA (n_incl_in_auc = 4, n_tested = 5)
WPMY1−TUBA1B−RELA (n_incl_in_auc = 5, n_tested = 5)
0.0
0.1
0.2
0.3
0.4
0.5
0.6
0.7
0.8
0.9
1.0
auc_max_over_concentr_median
FDR−adjusted p−value < 0.05
FALSE
TRUE
Batch
All Aggr.
Within Batch 1
Within Batch 2
cell_type
A549
HepG2
WPMY1

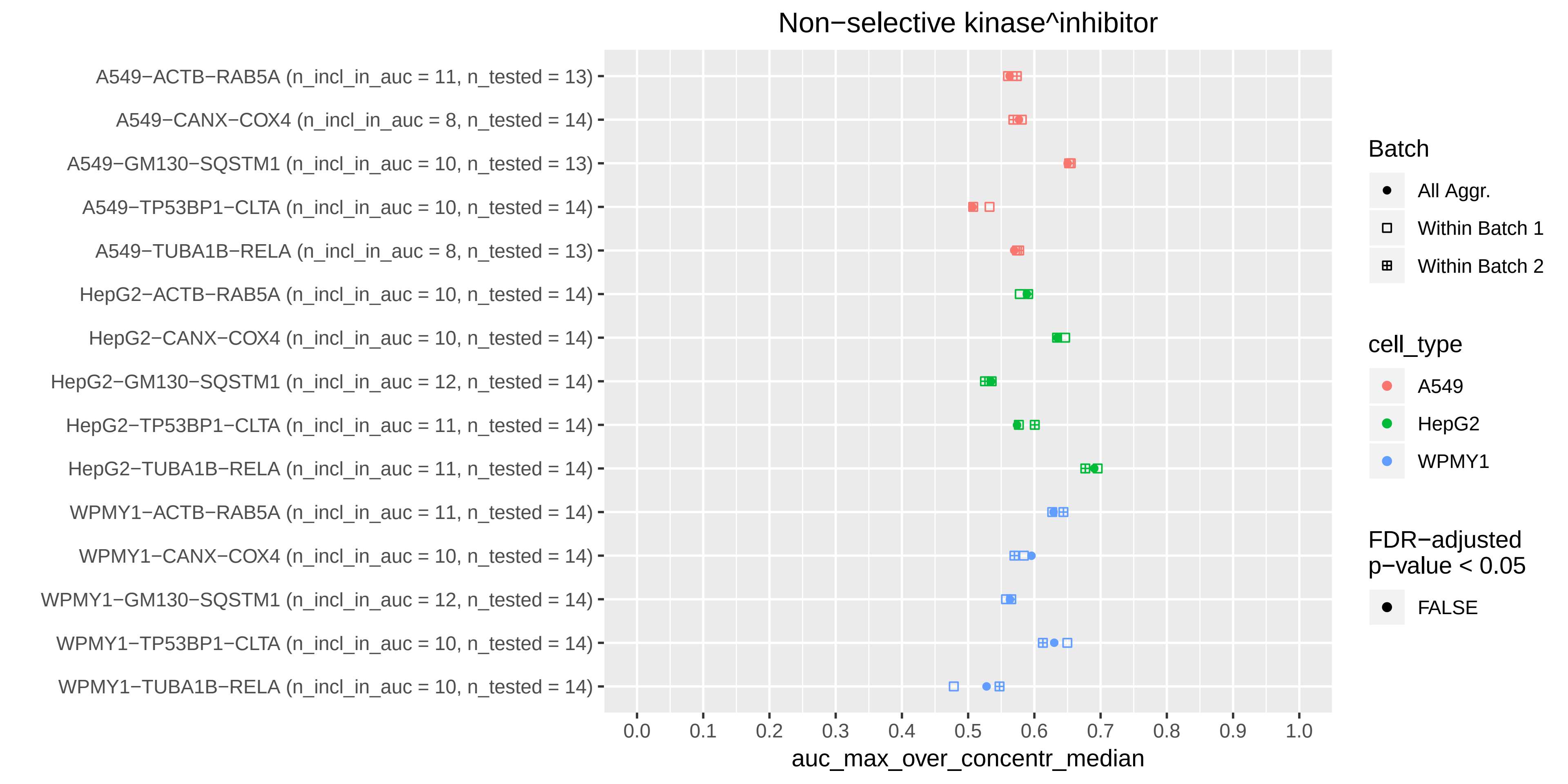

Non−selective kinase^inhibitor
A549−ACTB−RAB5A (n_incl_in_auc = 11, n_tested = 13)
A549−CANX−COX4 (n_incl_in_auc = 8, n_tested = 14)
A549−GM130−SQSTM1 (n_incl_in_auc = 10, n_tested = 13)
A549−TP53BP1−CLTA (n_incl_in_auc = 10, n_tested = 14)
A549−TUBA1B−RELA (n_incl_in_auc = 8, n_tested = 13)
HepG2−ACTB−RAB5A (n_incl_in_auc = 10, n_tested = 14)
HepG2−CANX−COX4 (n_incl_in_auc = 10, n_tested = 14)
HepG2−GM130−SQSTM1 (n_incl_in_auc = 12, n_tested = 14)
HepG2−TP53BP1−CLTA (n_incl_in_auc = 11, n_tested = 14)
HepG2−TUBA1B−RELA (n_incl_in_auc = 11, n_tested = 14)
WPMY1−ACTB−RAB5A (n_incl_in_auc = 11, n_tested = 14)
WPMY1−CANX−COX4 (n_incl_in_auc = 10, n_tested = 14)
WPMY1−GM130−SQSTM1 (n_incl_in_auc = 12, n_tested = 14)
WPMY1−TP53BP1−CLTA (n_incl_in_auc = 10, n_tested = 14)
WPMY1−TUBA1B−RELA (n_incl_in_auc = 10, n_tested = 14)
0.0 0.1 0.2 0.3 0.4 0.5 0.6 0.7 0.8 0.9 1.0
auc_max_over_concentr_median
Batch
All Aggr.
Within Batch 1
Within Batch 2
cell_type
A549
HepG2
WPMY1
FDR−adjusted p−value < 0.05
FALSE

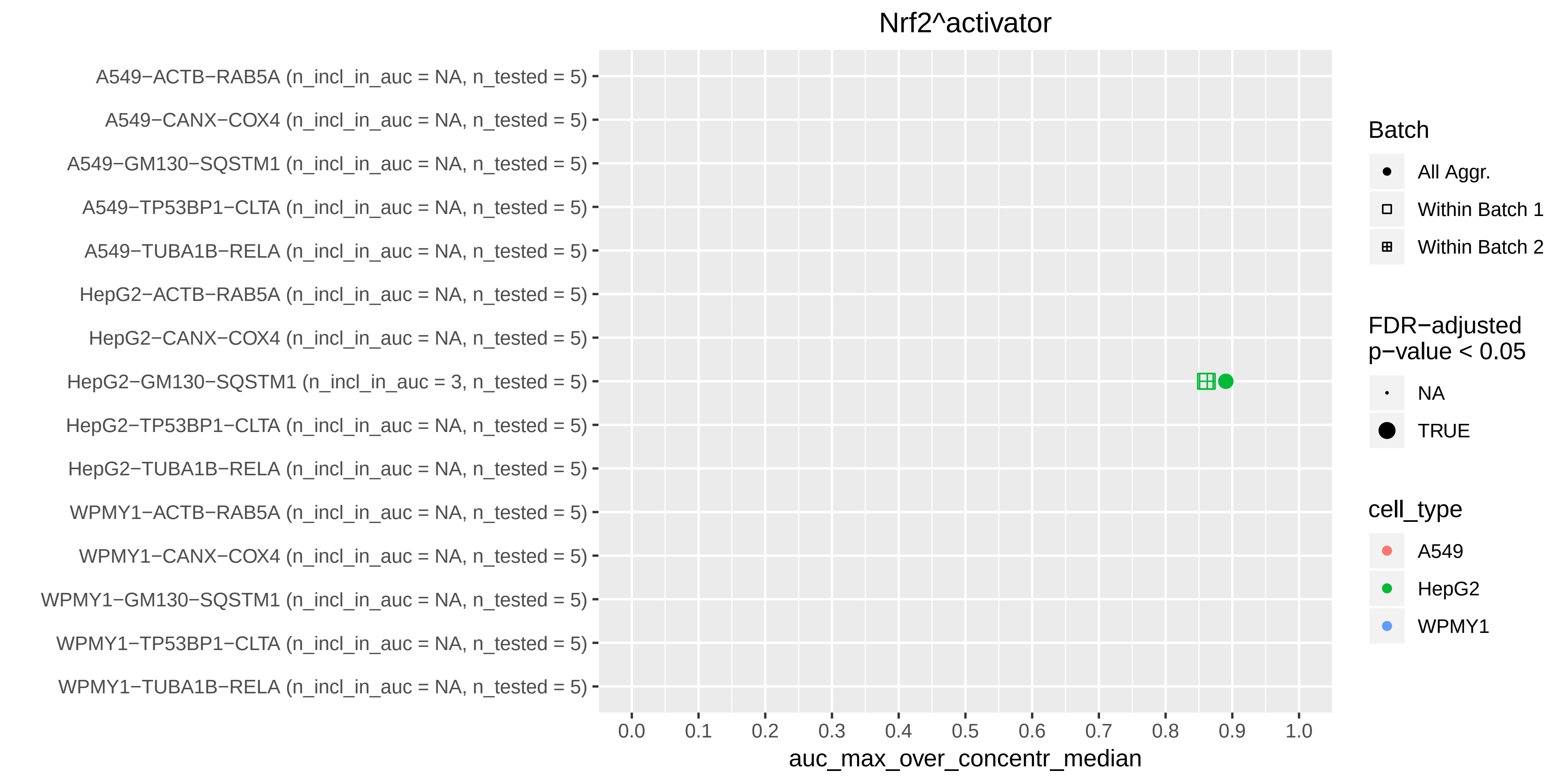

Nrf2^activator
A549−ACTB−RAB5A (n_incl_in_auc = NA, n_tested = 5)
A549−CANX−COX4 (n_incl_in_auc = NA, n_tested = 5)
A549−GM130−SQSTM1 (n_incl_in_auc = NA, n_tested = 5)
A549−TP53BP1−CLTA (n_incl_in_auc = NA, n_tested = 5)
A549−TUBA1B−RELA (n_incl_in_auc = NA, n_tested = 5)
HepG2−ACTB−RAB5A (n_incl_in_auc = NA, n_tested = 5)
HepG2−CANX−COX4 (n_incl_in_auc = NA, n_tested = 5)
HepG2−GM130−SQSTM1 (n_incl_in_auc = 3, n_tested = 5)
HepG2−TP53BP1−CLTA (n_incl_in_auc = NA, n_tested = 5)
HepG2−TUBA1B−RELA (n_incl_in_auc = NA, n_tested = 5)
WPMY1−ACTB−RAB5A (n_incl_in_auc = NA, n_tested = 5)
WPMY1−CANX−COX4 (n_incl_in_auc = NA, n_tested = 5)
WPMY1−GM130−SQSTM1 (n_incl_in_auc = NA, n_tested = 5)
WPMY1−TP53BP1−CLTA (n_incl_in_auc = NA, n_tested = 5)
WPMY1−TUBA1B−RELA (n_incl_in_auc = NA, n_tested = 5)
0.0
0.1
0.2
0.3
0.4
0.5
0.6
0.7
0.8
0.9
1.0
auc_max_over_concentr_median
Batch
All Aggr.
Within Batch 1
Within Batch 2
FDR−adjusted
p−value < 0.05
NA
TRUE
cell_type
A549
HepG2
WPMY1

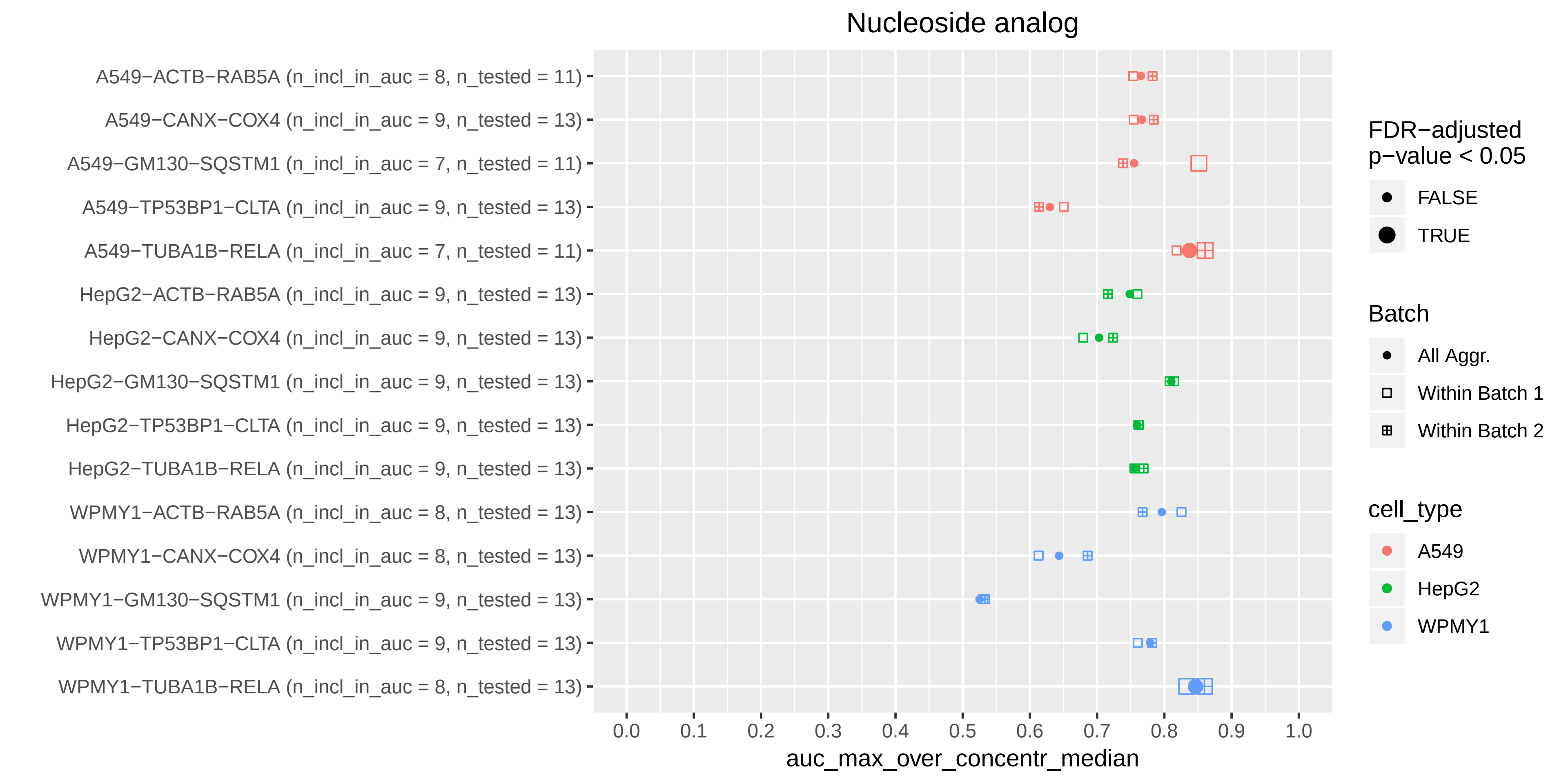

Nucleoside analog
A549−ACTB−RAB5A (n_incl_in_auc = 8, n_tested = 11)
A549−CANX−COX4 (n_incl_in_auc = 9, n_tested = 13)
A549−GM130−SQSTM1 (n_incl_in_auc = 7, n_tested = 11)
A549−TP53BP1−CLTA (n_incl_in_auc = 9, n_tested = 13)
A549−TUBA1B−RELA (n_incl_in_auc = 7, n_tested = 11)
HepG2−ACTB−RAB5A (n_incl_in_auc = 9, n_tested = 13)
HepG2−CANX−COX4 (n_incl_in_auc = 9, n_tested = 13)
HepG2−GM130−SQSTM1 (n_incl_in_auc = 9, n_tested = 13)
HepG2−TP53BP1−CLTA (n_incl_in_auc = 9, n_tested = 13)
HepG2−TUBA1B−RELA (n_incl_in_auc = 9, n_tested = 13)
WPMY1−ACTB−RAB5A (n_incl_in_auc = 8, n_tested = 13)
WPMY1−CANX−COX4 (n_incl_in_auc = 8, n_tested = 13)
WPMY1−GM130−SQSTM1 (n_incl_in_auc = 9, n_tested = 13)
WPMY1−TP53BP1−CLTA (n_incl_in_auc = 9, n_tested = 13)
WPMY1−TUBA1B−RELA (n_incl_in_auc = 8, n_tested = 13)
0.0
0.1
0.2
0.3
0.4
0.5
0.6
0.7
0.8
0.9
1.0
auc_max_over_concentr_median
FDR−adjusted p−value < 0.05
FALSE
TRUE
Batch
All Aggr.
Within Batch 1
Within Batch 2
cell_type
A549
HepG2
WPMY1

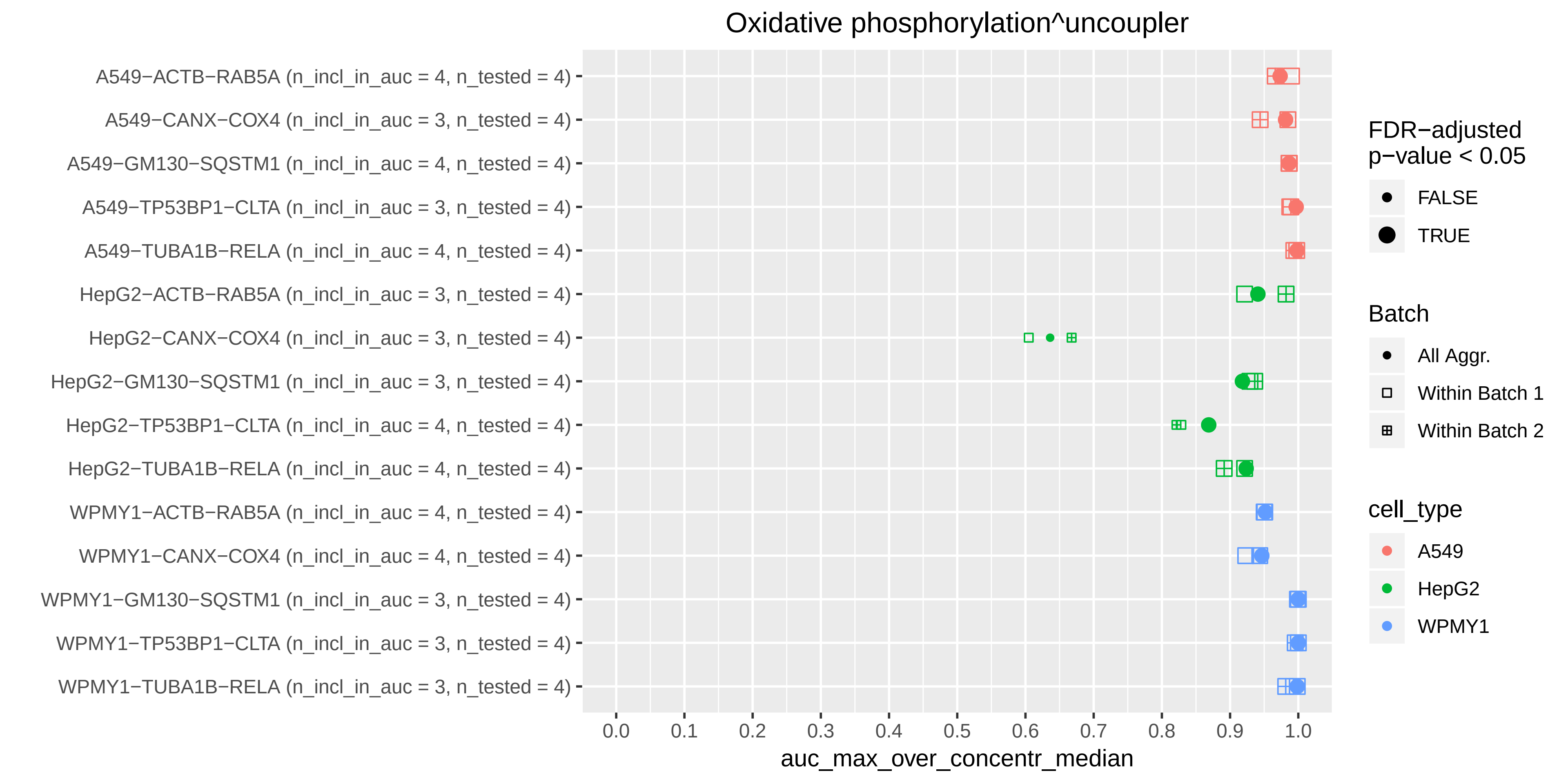

Oxidative phosphorylation^uncoupler
A549−ACTB−RAB5A (n_incl_in_auc = 4, n_tested = 4)
A549−CANX−COX4 (n_incl_in_auc = 3, n_tested = 4)
A549−GM130−SQSTM1 (n_incl_in_auc = 4, n_tested = 4)
A549−TP53BP1−CLTA (n_incl_in_auc = 3, n_tested = 4)
A549−TUBA1B−RELA (n_incl_in_auc = 4, n_tested = 4)
HepG2−ACTB−RAB5A (n_incl_in_auc = 3, n_tested = 4)
HepG2−CANX−COX4 (n_incl_in_auc = 3, n_tested = 4)
HepG2−GM130−SQSTM1 (n_incl_in_auc = 3, n_tested = 4)
HepG2−TP53BP1−CLTA (n_incl_in_auc = 4, n_tested = 4)
HepG2−TUBA1B−RELA (n_incl_in_auc = 4, n_tested = 4)
WPMY1−ACTB−RAB5A (n_incl_in_auc = 4, n_tested = 4)
WPMY1−CANX−COX4 (n_incl_in_auc = 4, n_tested = 4)
WPMY1−GM130−SQSTM1 (n_incl_in_auc = 3, n_tested = 4)
WPMY1−TP53BP1−CLTA (n_incl_in_auc = 3, n_tested = 4)
WPMY1−TUBA1B−RELA (n_incl_in_auc = 3, n_tested = 4)
0.0
0.1
0.2
0.3
0.4
0.5
0.6
0.7
0.8
0.9
1.0
auc_max_over_concentr_median
FDR−adjusted p−value < 0.05
FALSE
TRUE
Batch
All Aggr.
Within Batch 1
Within Batch 2
cell_type
A549
HepG2
WPMY1

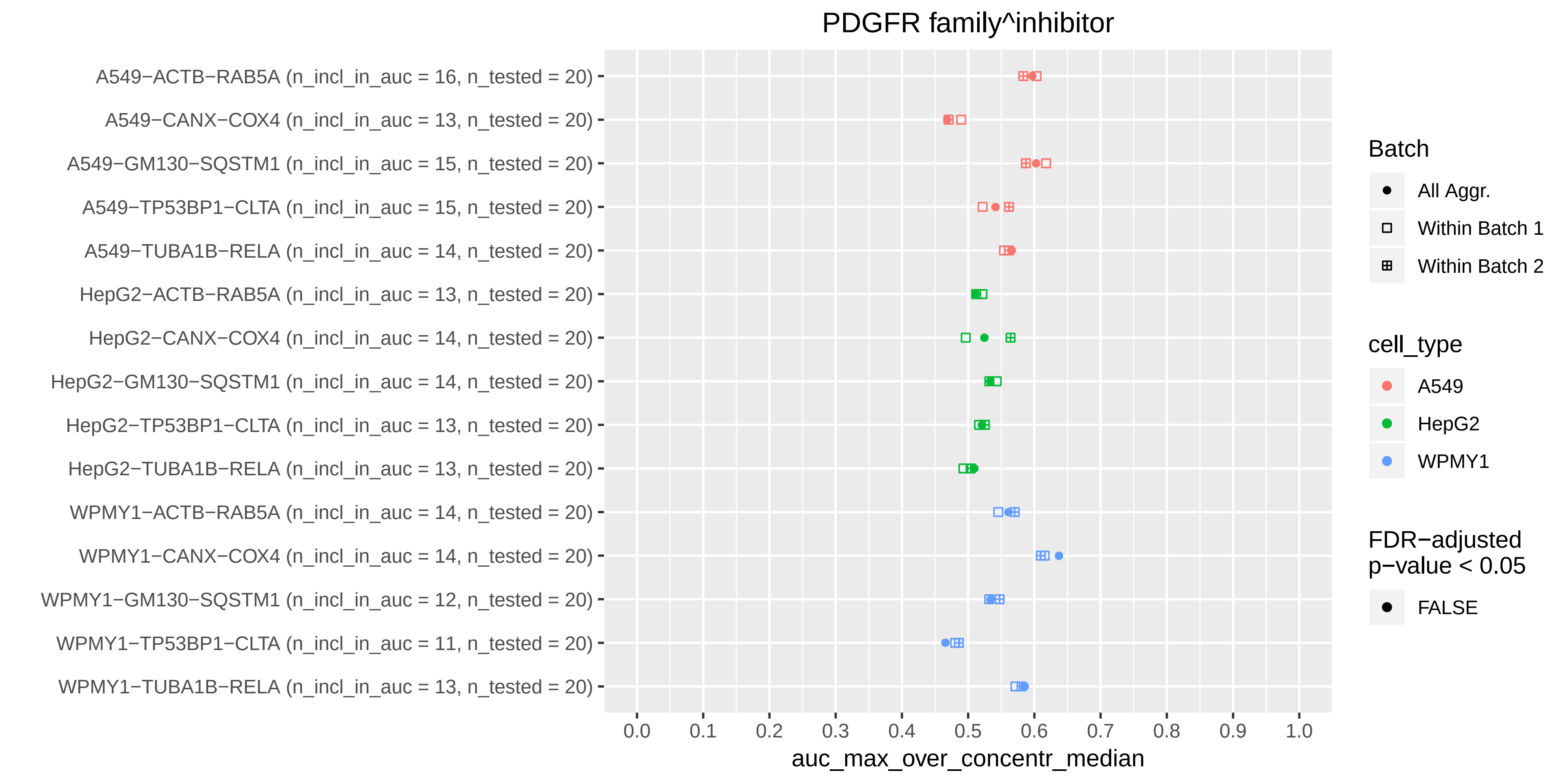

PDGFR family^inhibitor
A549−ACTB−RAB5A (n_incl_in_auc = 16, n_tested = 20)
A549−CANX−COX4 (n_incl_in_auc = 13, n_tested = 20)
A549−GM130−SQSTM1 (n_incl_in_auc = 15, n_tested = 20)
A549−TP53BP1−CLTA (n_incl_in_auc = 15, n_tested = 20)
A549−TUBA1B−RELA (n_incl_in_auc = 14, n_tested = 20)
HepG2−ACTB−RAB5A (n_incl_in_auc = 13, n_tested = 20)
HepG2−CANX−COX4 (n_incl_in_auc = 14, n_tested = 20)
HepG2−GM130−SQSTM1 (n_incl_in_auc = 14, n_tested = 20)
HepG2−TP53BP1−CLTA (n_incl_in_auc = 13, n_tested = 20)
HepG2−TUBA1B−RELA (n_incl_in_auc = 13, n_tested = 20)
WPMY1−ACTB−RAB5A (n_incl_in_auc = 14, n_tested = 20)
WPMY1−CANX−COX4 (n_incl_in_auc = 14, n_tested = 20)
WPMY1−GM130−SQSTM1 (n_incl_in_auc = 12, n_tested = 20)
WPMY1−TP53BP1−CLTA (n_incl_in_auc = 11, n_tested = 20)
WPMY1−TUBA1B−RELA (n_incl_in_auc = 13, n_tested = 20)
0.0
0.1
0.2
0.3
0.4
0.5
0.6
0.7
0.8
0.9
1.0
auc_max_over_concentr_median
Batch
All Aggr.
Within Batch 1
Within Batch 2
cell_type
A549
HepG2
WPMY1
FDR−adjusted p−value < 0.05
FALSE

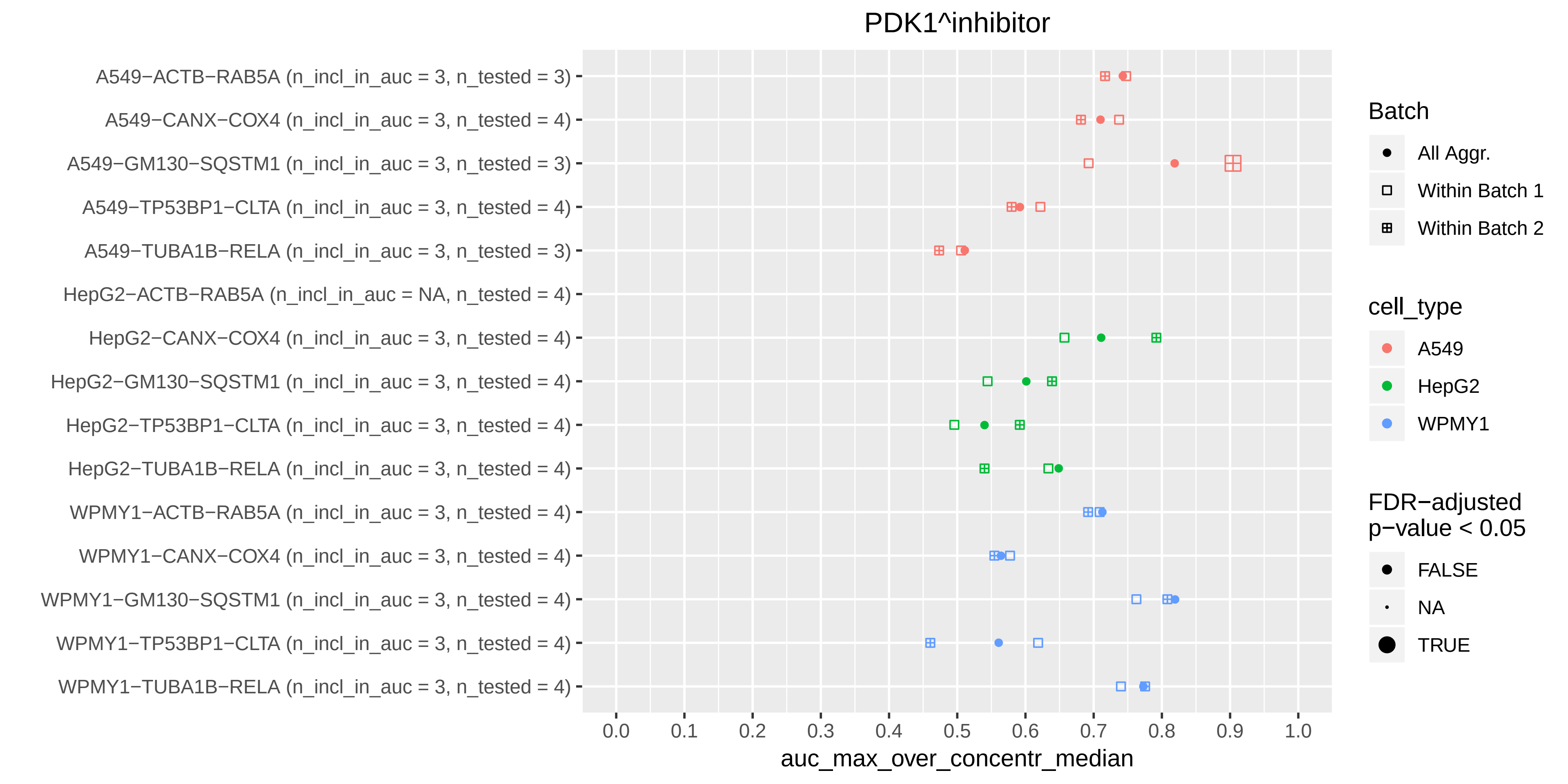
PDK1^inhibitor
A549−ACTB−RAB5A (n_incl_in_auc = 3, n_tested = 3)
A549−CANX−COX4 (n_incl_in_auc = 3, n_tested = 4)
A549−GM130−SQSTM1 (n_incl_in_auc = 3, n_tested = 3)
A549−TP53BP1−CLTA (n_incl_in_auc = 3, n_tested = 4)
A549−TUBA1B−RELA (n_incl_in_auc = 3, n_tested = 3)
HepG2−ACTB−RAB5A (n_incl_in_auc = NA, n_tested = 4)
HepG2−CANX−COX4 (n_incl_in_auc = 3, n_tested = 4)
HepG2−GM130−SQSTM1 (n_incl_in_auc = 3, n_tested = 4)
HepG2−TP53BP1−CLTA (n_incl_in_auc = 3, n_tested = 4)
HepG2−TUBA1B−RELA (n_incl_in_auc = 3, n_tested = 4)
WPMY1−ACTB−RAB5A (n_incl_in_auc = 3, n_tested = 4)
WPMY1−CANX−COX4 (n_incl_in_auc = 3, n_tested = 4)
WPMY1−GM130−SQSTM1 (n_incl_in_auc = 3, n_tested = 4)
WPMY1−TP53BP1−CLTA (n_incl_in_auc = 3, n_tested = 4)
WPMY1−TUBA1B−RELA (n_incl_in_auc = 3, n_tested = 4)
0.0
0.1
0.2
0.3
0.4
0.5
0.6
0.7
0.8
0.9
1.0
auc_max_over_concentr_median
Batch
All Aggr.
Within Batch 1
Within Batch 2
cell_type
A549
HepG2
WPMY1
FDR−adjusted p−value < 0.05
FALSE
NA
TRUE

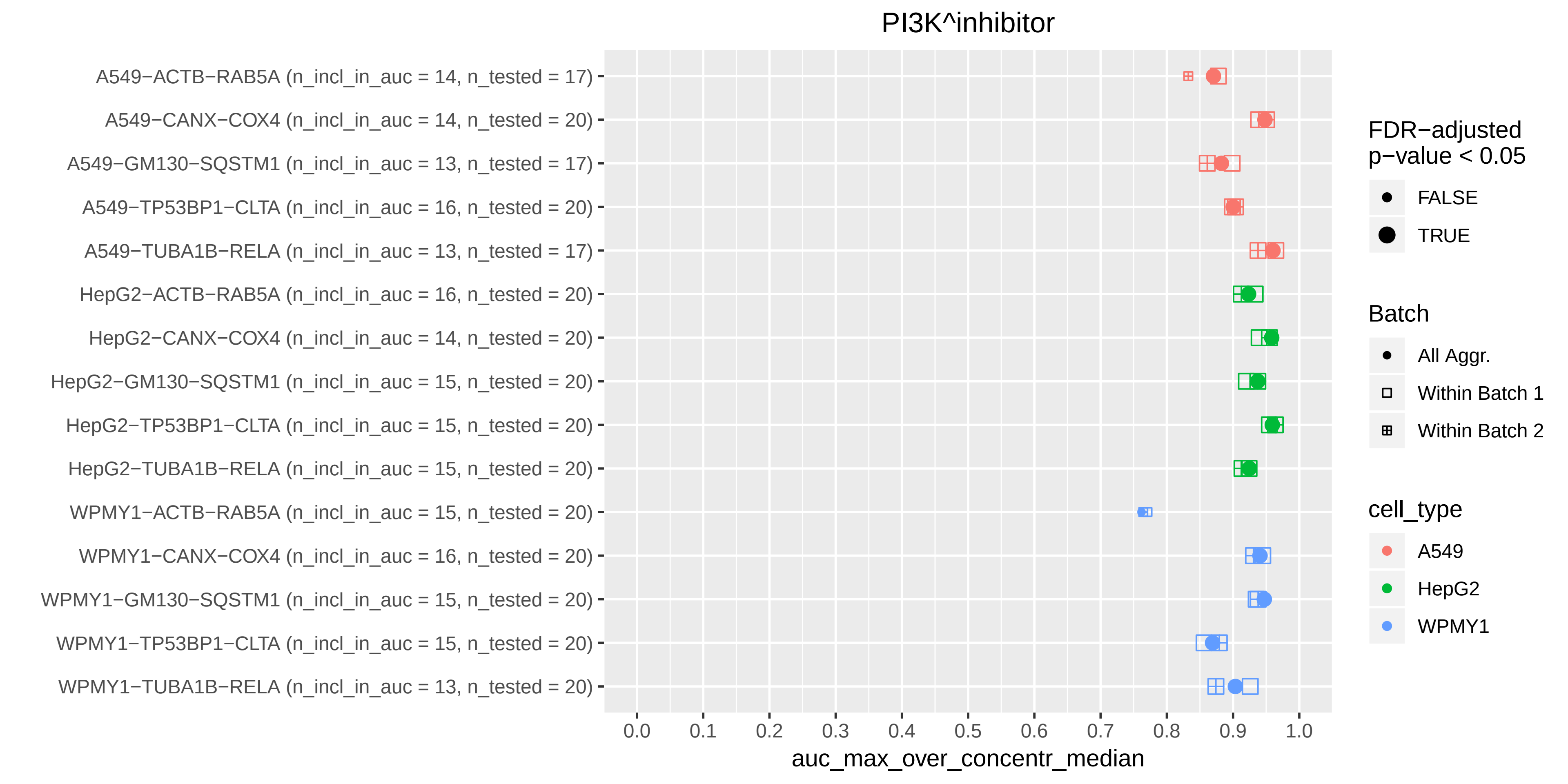

PI3K^inhibitor
A549−ACTB−RAB5A (n_incl_in_auc = 14, n_tested = 17)
A549−CANX−COX4 (n_incl_in_auc = 14, n_tested = 20)
A549−GM130−SQSTM1 (n_incl_in_auc = 13, n_tested = 17)
A549−TP53BP1−CLTA (n_incl_in_auc = 16, n_tested = 20)
A549−TUBA1B−RELA (n_incl_in_auc = 13, n_tested = 17)
HepG2−ACTB−RAB5A (n_incl_in_auc = 16, n_tested = 20)
HepG2−CANX−COX4 (n_incl_in_auc = 14, n_tested = 20)
HepG2−GM130−SQSTM1 (n_incl_in_auc = 15, n_tested = 20)
HepG2−TP53BP1−CLTA (n_incl_in_auc = 15, n_tested = 20)
HepG2−TUBA1B−RELA (n_incl_in_auc = 15, n_tested = 20)
WPMY1−ACTB−RAB5A (n_incl_in_auc = 15, n_tested = 20)
WPMY1−CANX−COX4 (n_incl_in_auc = 16, n_tested = 20)
WPMY1−GM130−SQSTM1 (n_incl_in_auc = 15, n_tested = 20)
WPMY1−TP53BP1−CLTA (n_incl_in_auc = 15, n_tested = 20)
WPMY1−TUBA1B−RELA (n_incl_in_auc = 13, n_tested = 20)
0.0
0.1
0.2
0.3
0.4
0.5
0.6
0.7
0.8
0.9
1.0
auc_max_over_concentr_median
FDR−adjusted p−value < 0.05
FALSE
TRUE
Batch
All Aggr.
Within Batch 1
Within Batch 2
cell_type
A549
HepG2
WPMY1

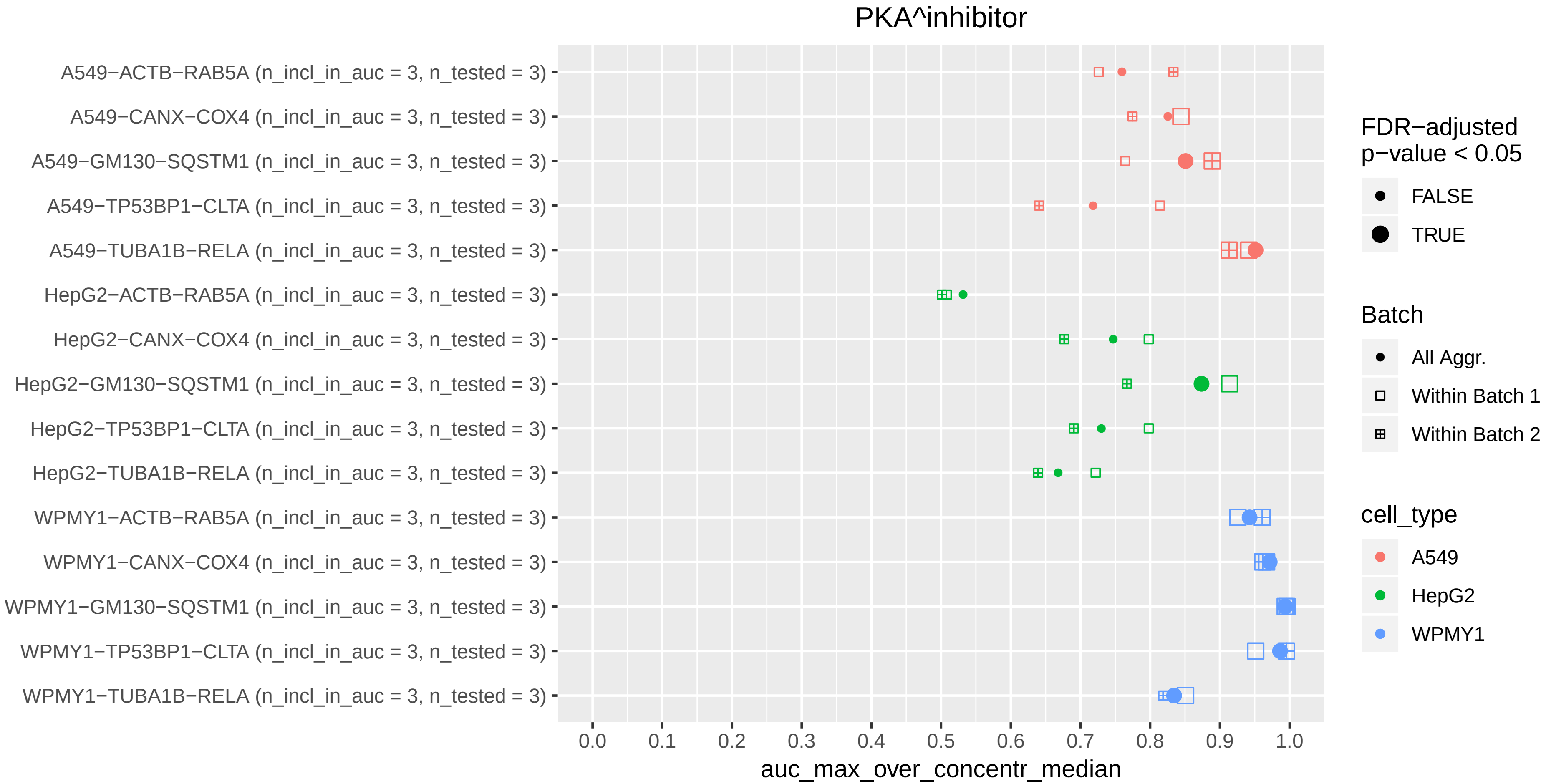

PKA^inhibitor
A549−ACTB−RAB5A (n_incl_in_auc = 3, n_tested = 3)
A549−CANX−COX4 (n_incl_in_auc = 3, n_tested = 3)
A549−GM130−SQSTM1 (n_incl_in_auc = 3, n_tested = 3)
A549−TP53BP1−CLTA (n_incl_in_auc = 3, n_tested = 3)
A549−TUBA1B−RELA (n_incl_in_auc = 3, n_tested = 3)
HepG2−ACTB−RAB5A (n_incl_in_auc = 3, n_tested = 3)
HepG2−CANX−COX4 (n_incl_in_auc = 3, n_tested = 3)
HepG2−GM130−SQSTM1 (n_incl_in_auc = 3, n_tested = 3)
HepG2−TP53BP1−CLTA (n_incl_in_auc = 3, n_tested = 3)
HepG2−TUBA1B−RELA (n_incl_in_auc = 3, n_tested = 3)
WPMY1−ACTB−RAB5A (n_incl_in_auc = 3, n_tested = 3)
WPMY1−CANX−COX4 (n_incl_in_auc = 3, n_tested = 3)
WPMY1−GM130−SQSTM1 (n_incl_in_auc = 3, n_tested = 3)
WPMY1−TP53BP1−CLTA (n_incl_in_auc = 3, n_tested = 3)
WPMY1−TUBA1B−RELA (n_incl_in_auc = 3, n_tested = 3)
0.0
0.1
0.2
0.3
0.4
0.5
0.6
0.7
0.8
0.9
1.0
auc_max_over_concentr_median
FDR−adjusted p−value < 0.05
FALSE
TRUE
Batch
All Aggr.
Within Batch 1
Within Batch 2
cell_type
A549
HepG2
WPMY1

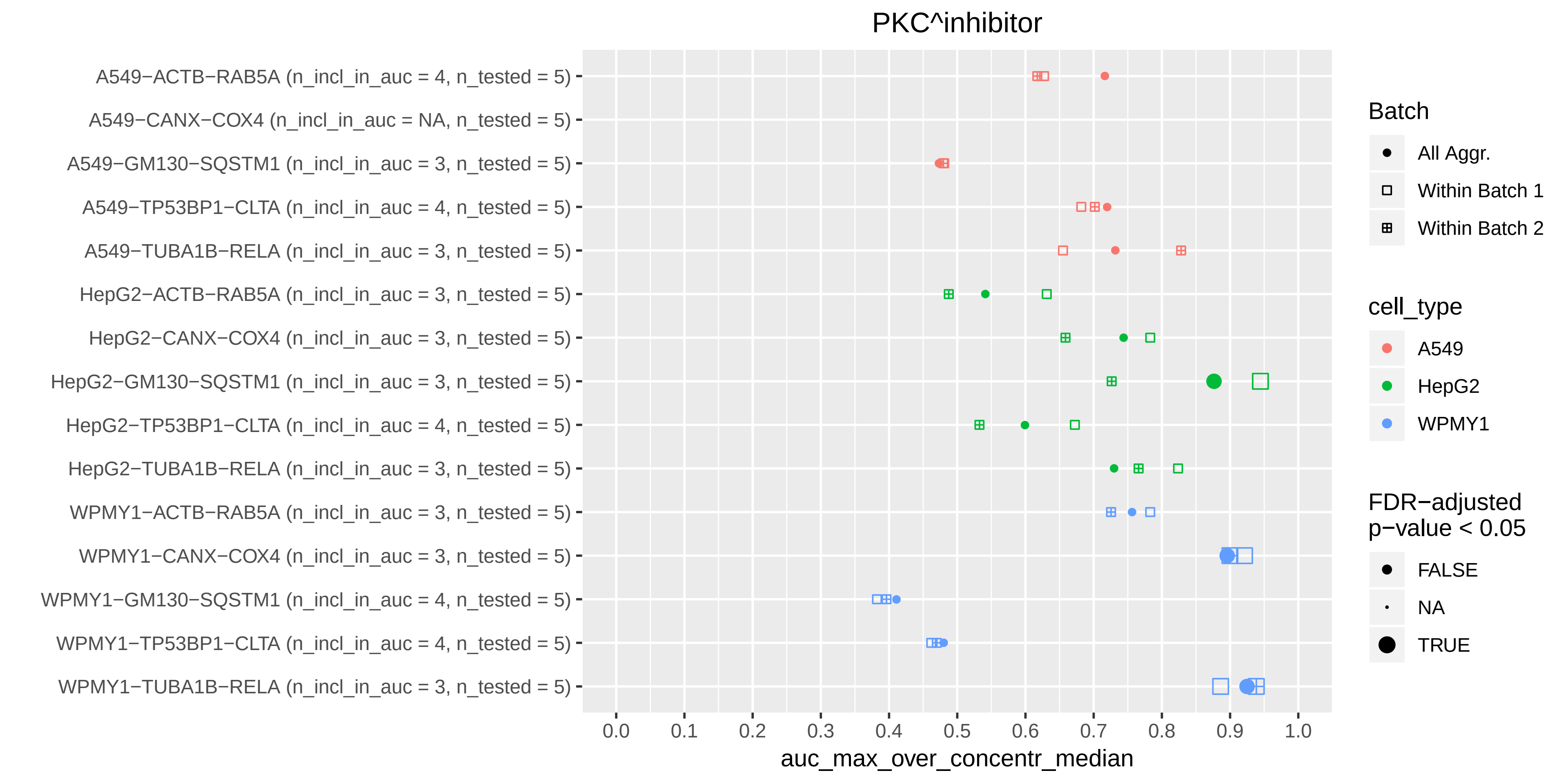

PKC^inhibitor
A549−ACTB−RAB5A (n_incl_in_auc = 4, n_tested = 5)
A549−CANX−COX4 (n_incl_in_auc = NA, n_tested = 5)
A549−GM130−SQSTM1 (n_incl_in_auc = 3, n_tested = 5)
A549−TP53BP1−CLTA (n_incl_in_auc = 4, n_tested = 5)
A549−TUBA1B−RELA (n_incl_in_auc = 3, n_tested = 5)
HepG2−ACTB−RAB5A (n_incl_in_auc = 3, n_tested = 5)
HepG2−CANX−COX4 (n_incl_in_auc = 3, n_tested = 5)
HepG2−GM130−SQSTM1 (n_incl_in_auc = 3, n_tested = 5)
HepG2−TP53BP1−CLTA (n_incl_in_auc = 4, n_tested = 5)
HepG2−TUBA1B−RELA (n_incl_in_auc = 3, n_tested = 5)
WPMY1−ACTB−RAB5A (n_incl_in_auc = 3, n_tested = 5)
WPMY1−CANX−COX4 (n_incl_in_auc = 3, n_tested = 5)
WPMY1−GM130−SQSTM1 (n_incl_in_auc = 4, n_tested = 5)
WPMY1−TP53BP1−CLTA (n_incl_in_auc = 4, n_tested = 5)
WPMY1−TUBA1B−RELA (n_incl_in_auc = 3, n_tested = 5)
0.0
0.1
0.2
0.3
0.4
0.5
0.6
0.7
0.8
0.9
1.0
auc_max_over_concentr_median
Batch
All Aggr.
Within Batch 1
Within Batch 2
cell_type
A549
HepG2
WPMY1
FDR−adjusted p−value < 0.05
FALSE
NA
TRUE

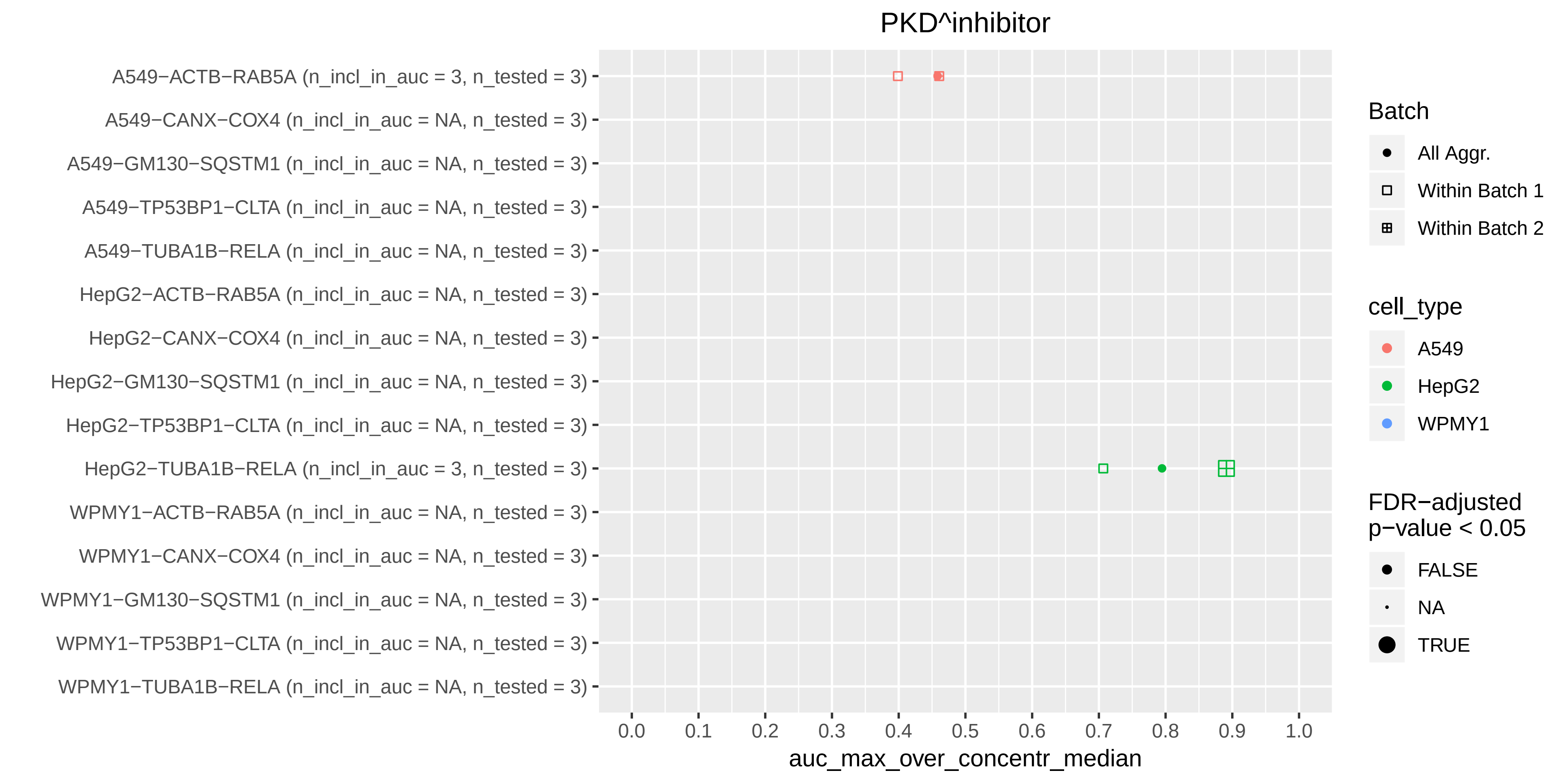
PKD^inhibitor
A549−ACTB−RAB5A (n_incl_in_auc = 3, n_tested = 3)
A549−CANX−COX4 (n_incl_in_auc = NA, n_tested = 3)
A549−GM130−SQSTM1 (n_incl_in_auc = NA, n_tested = 3)
A549−TP53BP1−CLTA (n_incl_in_auc = NA, n_tested = 3)
A549−TUBA1B−RELA (n_incl_in_auc = NA, n_tested = 3)
HepG2−ACTB−RAB5A (n_incl_in_auc = NA, n_tested = 3)
HepG2−CANX−COX4 (n_incl_in_auc = NA, n_tested = 3)
HepG2−GM130−SQSTM1 (n_incl_in_auc = NA, n_tested = 3)
HepG2−TP53BP1−CLTA (n_incl_in_auc = NA, n_tested = 3)
HepG2−TUBA1B−RELA (n_incl_in_auc = 3, n_tested = 3)
WPMY1−ACTB−RAB5A (n_incl_in_auc = NA, n_tested = 3)
WPMY1−CANX−COX4 (n_incl_in_auc = NA, n_tested = 3)
WPMY1−GM130−SQSTM1 (n_incl_in_auc = NA, n_tested = 3)
WPMY1−TP53BP1−CLTA (n_incl_in_auc = NA, n_tested = 3)
WPMY1−TUBA1B−RELA (n_incl_in_auc = NA, n_tested = 3)
0.0
0.1
0.2
0.3
0.4
0.5
0.6
0.7
0.8
0.9
1.0
auc_max_over_concentr_median
Batch
All Aggr.
Within Batch 1
Within Batch 2
cell_type
A549
HepG2
WPMY1
FDR−adjusted
p−value < 0.05
FALSE
NA
TRUE

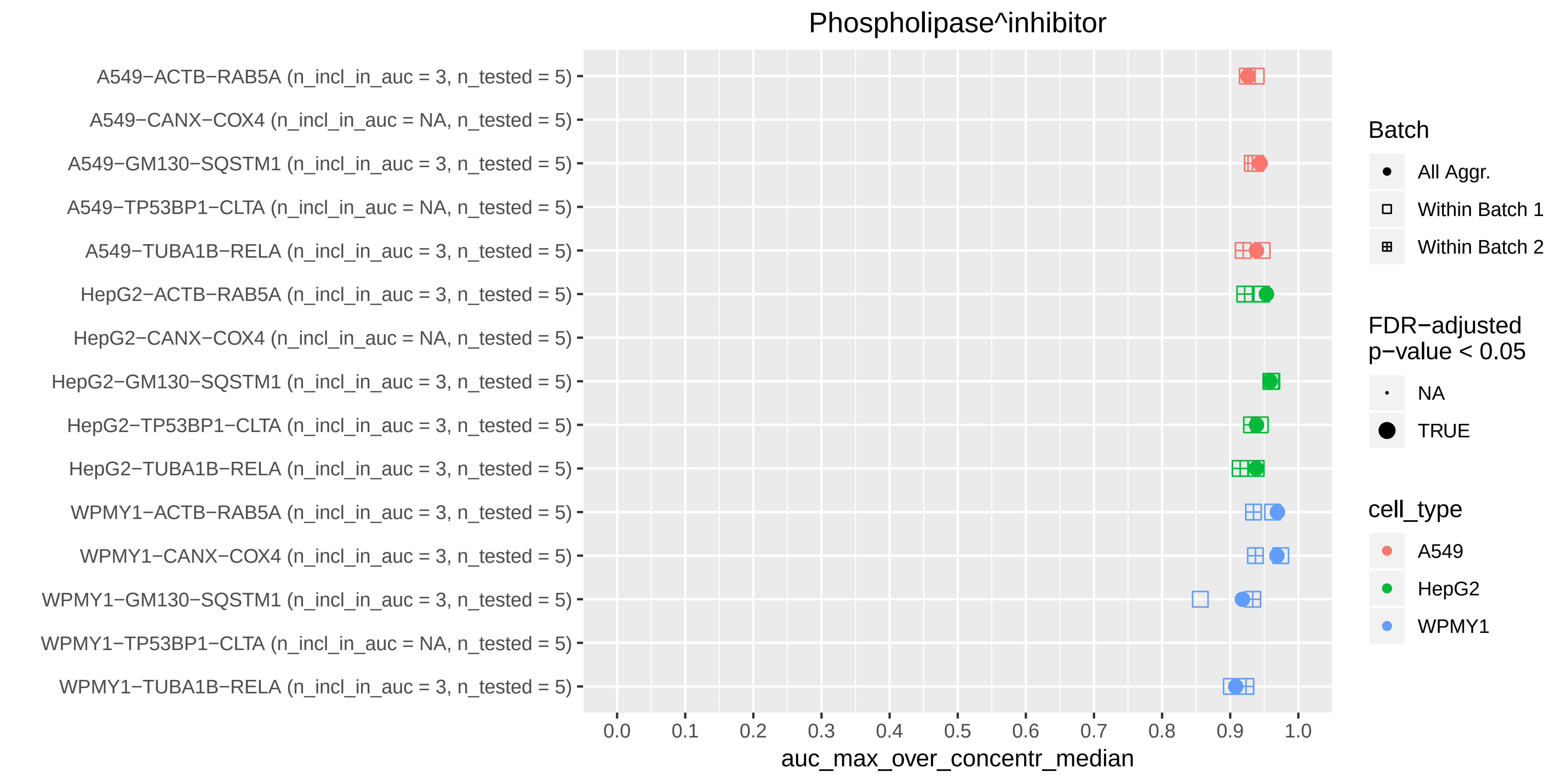

Phospholipase^inhibitor
A549−ACTB−RAB5A (n_incl_in_auc = 3, n_tested = 5)
A549−CANX−COX4 (n_incl_in_auc = NA, n_tested = 5)
A549−GM130−SQSTM1 (n_incl_in_auc = 3, n_tested = 5)
A549−TP53BP1−CLTA (n_incl_in_auc = NA, n_tested = 5)
A549−TUBA1B−RELA (n_incl_in_auc = 3, n_tested = 5)
HepG2−ACTB−RAB5A (n_incl_in_auc = 3, n_tested = 5)
HepG2−CANX−COX4 (n_incl_in_auc = NA, n_tested = 5)
HepG2−GM130−SQSTM1 (n_incl_in_auc = 3, n_tested = 5)
HepG2−TP53BP1−CLTA (n_incl_in_auc = 3, n_tested = 5)
HepG2−TUBA1B−RELA (n_incl_in_auc = 3, n_tested = 5)
WPMY1−ACTB−RAB5A (n_incl_in_auc = 3, n_tested = 5)
WPMY1−CANX−COX4 (n_incl_in_auc = 3, n_tested = 5)
WPMY1−GM130−SQSTM1 (n_incl_in_auc = 3, n_tested = 5)
WPMY1−TP53BP1−CLTA (n_incl_in_auc = NA, n_tested = 5)
WPMY1−TUBA1B−RELA (n_incl_in_auc = 3, n_tested = 5)
0.0
0.1
0.2
0.3
0.4
0.5
0.6
0.7
0.8
0.9
1.0
auc_max_over_concentr_median
Batch
All Aggr.
Within Batch 1
Within Batch 2
FDR−adjusted
p−value < 0.05
NA
TRUE
cell_type
A549
HepG2
WPMY1

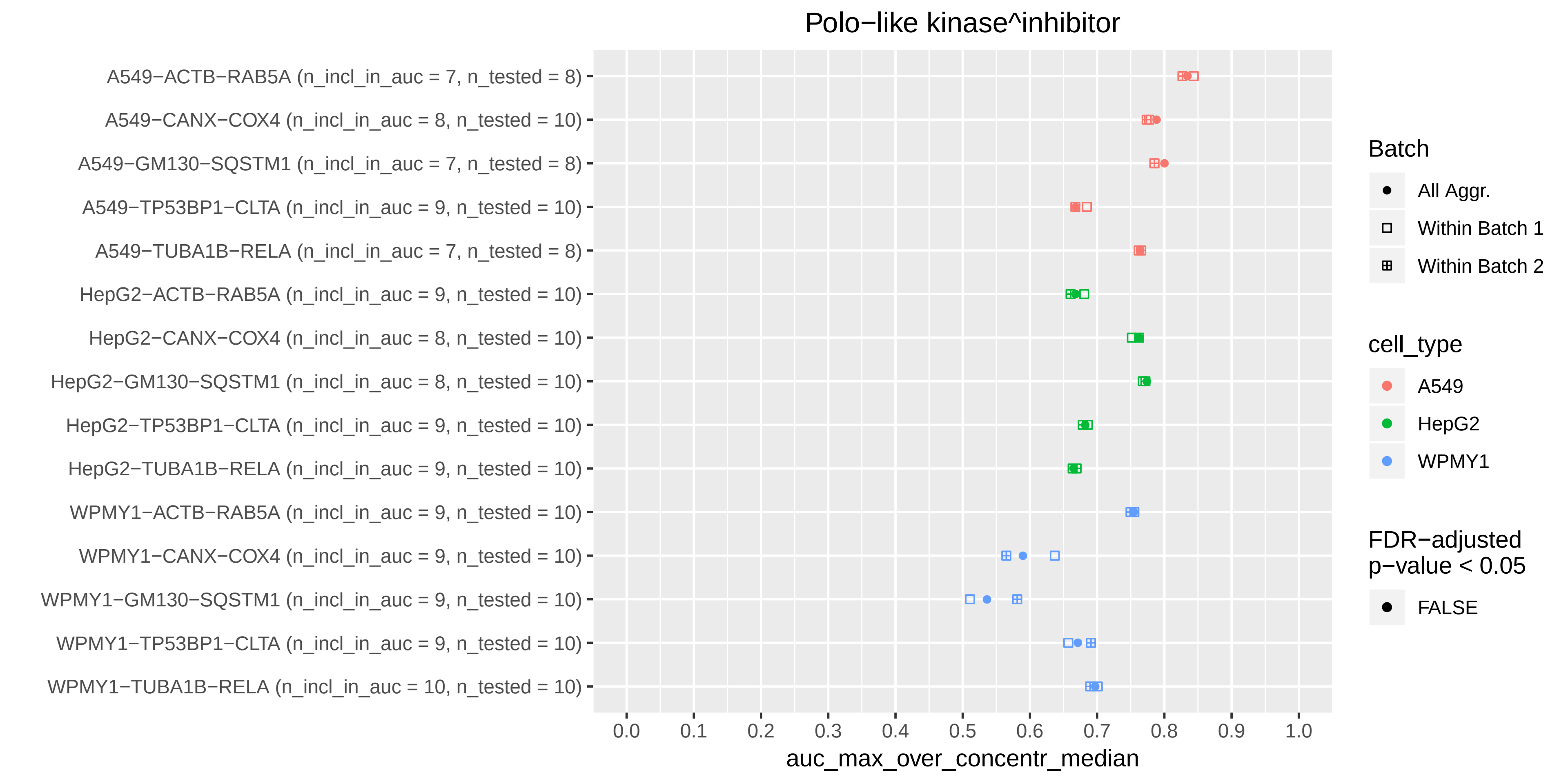
Polo−like kinase^inhibitor
A549−ACTB−RAB5A (n_incl_in_auc = 7, n_tested = 8)
A549−CANX−COX4 (n_incl_in_auc = 8, n_tested = 10)
A549−GM130−SQSTM1 (n_incl_in_auc = 7, n_tested = 8)
A549−TP53BP1−CLTA (n_incl_in_auc = 9, n_tested = 10)
A549−TUBA1B−RELA (n_incl_in_auc = 7, n_tested = 8)
HepG2−ACTB−RAB5A (n_incl_in_auc = 9, n_tested = 10)
HepG2−CANX−COX4 (n_incl_in_auc = 8, n_tested = 10)
HepG2−GM130−SQSTM1 (n_incl_in_auc = 8, n_tested = 10)
HepG2−TP53BP1−CLTA (n_incl_in_auc = 9, n_tested = 10)
HepG2−TUBA1B−RELA (n_incl_in_auc = 9, n_tested = 10)
WPMY1−ACTB−RAB5A (n_incl_in_auc = 9, n_tested = 10)
WPMY1−CANX−COX4 (n_incl_in_auc = 9, n_tested = 10)
WPMY1−GM130−SQSTM1 (n_incl_in_auc = 9, n_tested = 10)
WPMY1−TP53BP1−CLTA (n_incl_in_auc = 9, n_tested = 10)
WPMY1−TUBA1B−RELA (n_incl_in_auc = 10, n_tested = 10)
0.0
0.1
0.2
0.3
0.4
0.5
0.6
0.7
0.8
0.9
1.0
auc_max_over_concentr_median
Batch
All Aggr.
Within Batch 1
Within Batch 2
cell_type
A549
HepG2
WPMY1
FDR−adjusted p−value < 0.05
FALSE

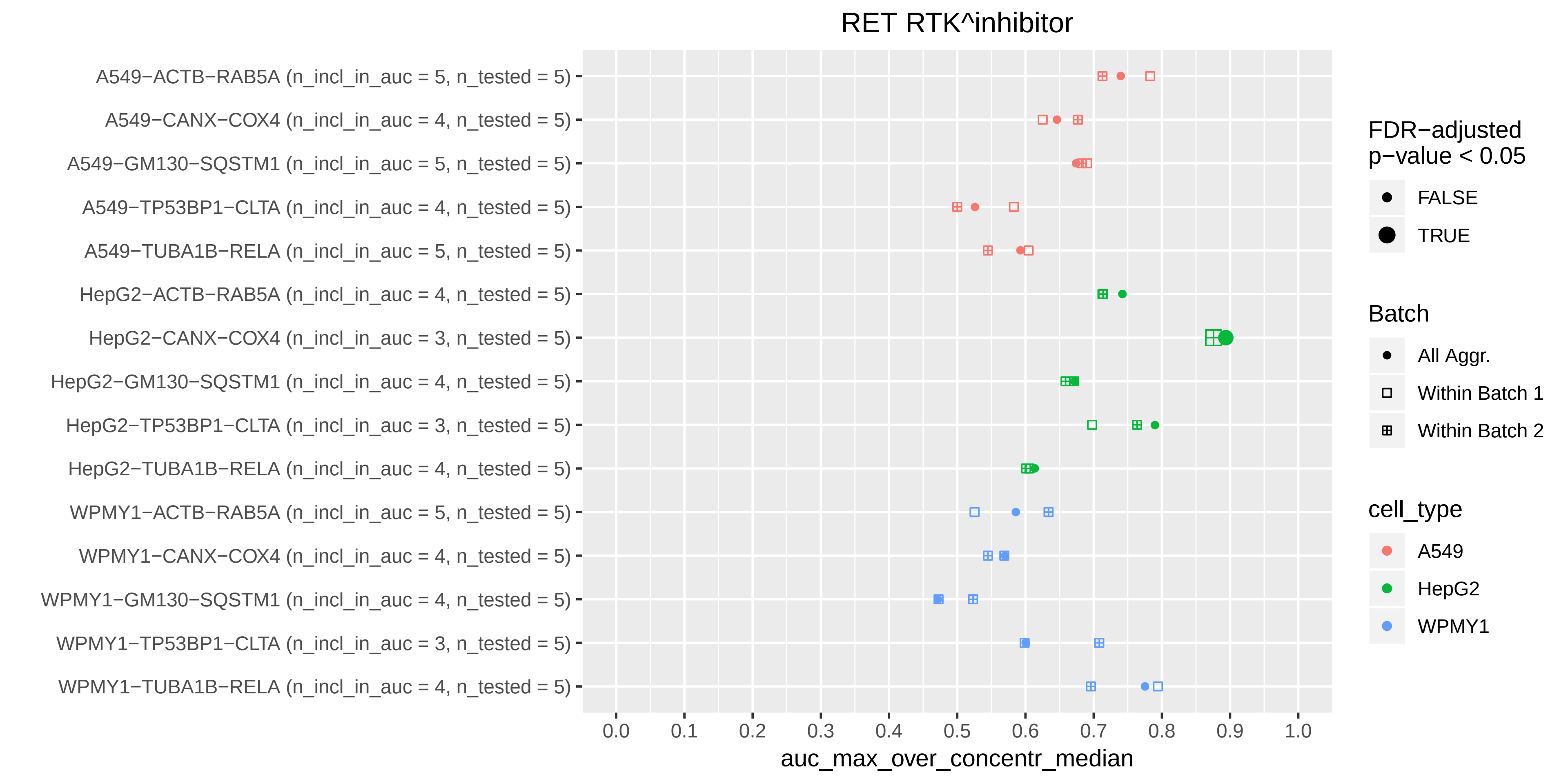

RET RTK^inhibitor
A549−ACTB−RAB5A (n_incl_in_auc = 5, n_tested = 5)
A549−CANX−COX4 (n_incl_in_auc = 4, n_tested = 5)
A549−GM130−SQSTM1 (n_incl_in_auc = 5, n_tested = 5)
A549−TP53BP1−CLTA (n_incl_in_auc = 4, n_tested = 5)
A549−TUBA1B−RELA (n_incl_in_auc = 5, n_tested = 5)
HepG2−ACTB−RAB5A (n_incl_in_auc = 4, n_tested = 5)
HepG2−CANX−COX4 (n_incl_in_auc = 3, n_tested = 5)
HepG2−GM130−SQSTM1 (n_incl_in_auc = 4, n_tested = 5)
HepG2−TP53BP1−CLTA (n_incl_in_auc = 3, n_tested = 5)
HepG2−TUBA1B−RELA (n_incl_in_auc = 4, n_tested = 5)
WPMY1−ACTB−RAB5A (n_incl_in_auc = 5, n_tested = 5)
WPMY1−CANX−COX4 (n_incl_in_auc = 4, n_tested = 5)
WPMY1−GM130−SQSTM1 (n_incl_in_auc = 4, n_tested = 5)
WPMY1−TP53BP1−CLTA (n_incl_in_auc = 3, n_tested = 5)
WPMY1−TUBA1B−RELA (n_incl_in_auc = 4, n_tested = 5)
0.0
0.1
0.2
0.3
0.4
0.5
0.6
0.7
0.8
0.9
1.0
auc_max_over_concentr_median
FDR−adjusted
p−value < 0.05
FALSE
TRUE
Batch
All Aggr.
Within Batch 1
Within Batch 2
cell_type
A549
HepG2
WPMY1

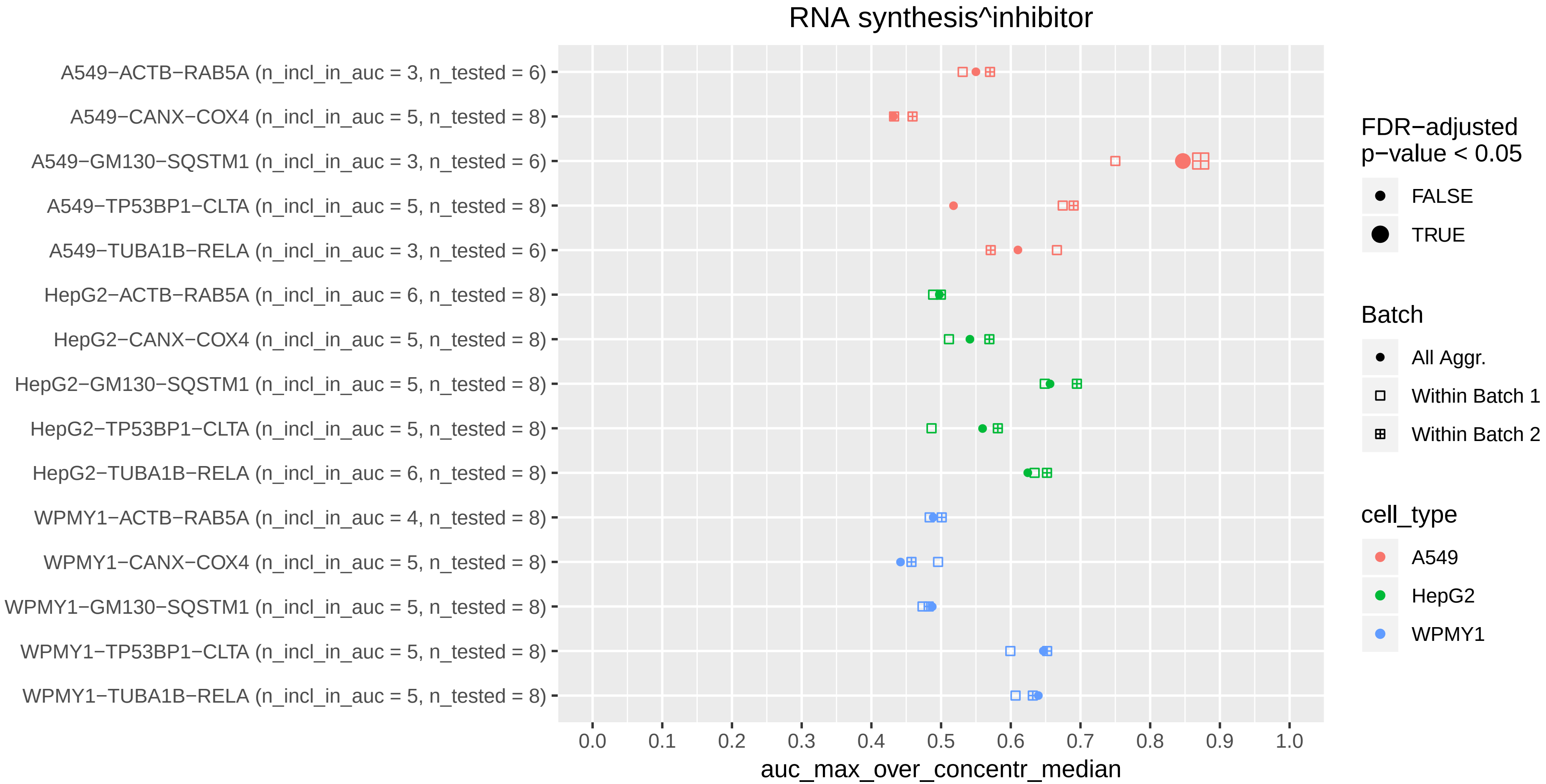
RNA synthesis^inhibitor
A549−ACTB−RAB5A (n_incl_in_auc = 3, n_tested = 6)
A549−CANX−COX4 (n_incl_in_auc = 5, n_tested = 8)
A549−GM130−SQSTM1 (n_incl_in_auc = 3, n_tested = 6)
A549−TP53BP1−CLTA (n_incl_in_auc = 5, n_tested = 8)
A549−TUBA1B−RELA (n_incl_in_auc = 3, n_tested = 6)
HepG2−ACTB−RAB5A (n_incl_in_auc = 6, n_tested = 8)
HepG2−CANX−COX4 (n_incl_in_auc = 5, n_tested = 8)
HepG2−GM130−SQSTM1 (n_incl_in_auc = 5, n_tested = 8)
HepG2−TP53BP1−CLTA (n_incl_in_auc = 5, n_tested = 8)
HepG2−TUBA1B−RELA (n_incl_in_auc = 6, n_tested = 8)
WPMY1−ACTB−RAB5A (n_incl_in_auc = 4, n_tested = 8)
WPMY1−CANX−COX4 (n_incl_in_auc = 5, n_tested = 8)
WPMY1−GM130−SQSTM1 (n_incl_in_auc = 5, n_tested = 8)
WPMY1−TP53BP1−CLTA (n_incl_in_auc = 5, n_tested = 8)
WPMY1−TUBA1B−RELA (n_incl_in_auc = 5, n_tested = 8)
0.0 0.1 0.2 0.3 0.4 0.5 0.6 0.7 0.8 0.9 1.0
auc_max_over_concentr_median
FDR−adjusted p−value < 0.05
FALSE
TRUE
Batch
All Aggr.
Within Batch 1
Within Batch 2
cell_type
A549
HepG2
WPMY1

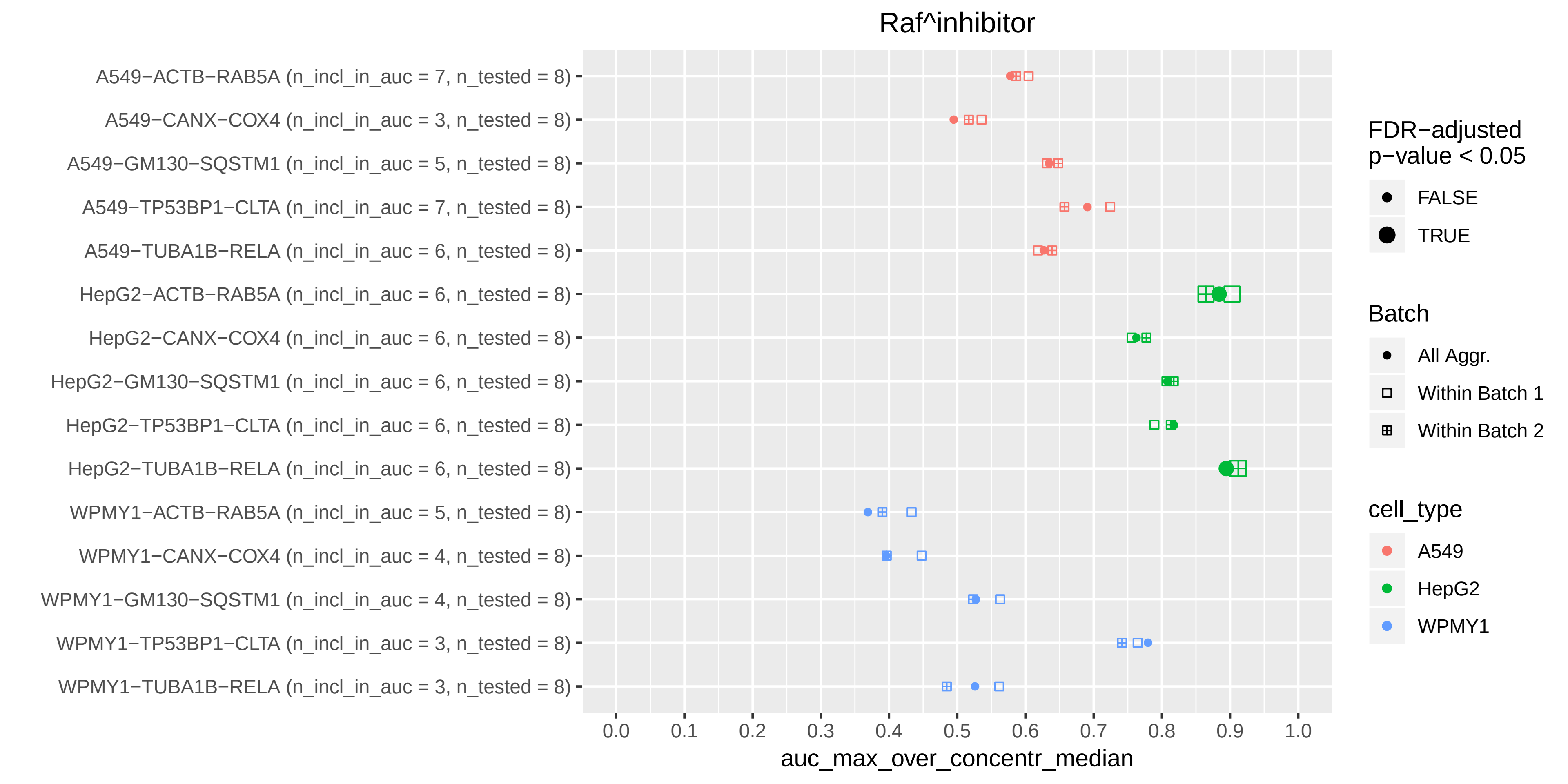

Raf^inhibitor
A549−ACTB−RAB5A (n_incl_in_auc = 7, n_tested = 8)
A549−CANX−COX4 (n_incl_in_auc = 3, n_tested = 8)
A549−GM130−SQSTM1 (n_incl_in_auc = 5, n_tested = 8)
A549−TP53BP1−CLTA (n_incl_in_auc = 7, n_tested = 8)
A549−TUBA1B−RELA (n_incl_in_auc = 6, n_tested = 8)
HepG2−ACTB−RAB5A (n_incl_in_auc = 6, n_tested = 8)
HepG2−CANX−COX4 (n_incl_in_auc = 6, n_tested = 8)
HepG2−GM130−SQSTM1 (n_incl_in_auc = 6, n_tested = 8)
HepG2−TP53BP1−CLTA (n_incl_in_auc = 6, n_tested = 8)
HepG2−TUBA1B−RELA (n_incl_in_auc = 6, n_tested = 8)
WPMY1−ACTB−RAB5A (n_incl_in_auc = 5, n_tested = 8)
WPMY1−CANX−COX4 (n_incl_in_auc = 4, n_tested = 8)
WPMY1−GM130−SQSTM1 (n_incl_in_auc = 4, n_tested = 8)
WPMY1−TP53BP1−CLTA (n_incl_in_auc = 3, n_tested = 8)
WPMY1−TUBA1B−RELA (n_incl_in_auc = 3, n_tested = 8)
0.0
0.1
0.2
0.3
0.4
0.5
0.6
0.7
0.8
0.9
1.0
auc_max_over_concentr_median
FDR−adjusted p−value < 0.05
FALSE
TRUE
Batch
All Aggr.
Within Batch 1
Within Batch 2
cell_type
A549
HepG2
WPMY1

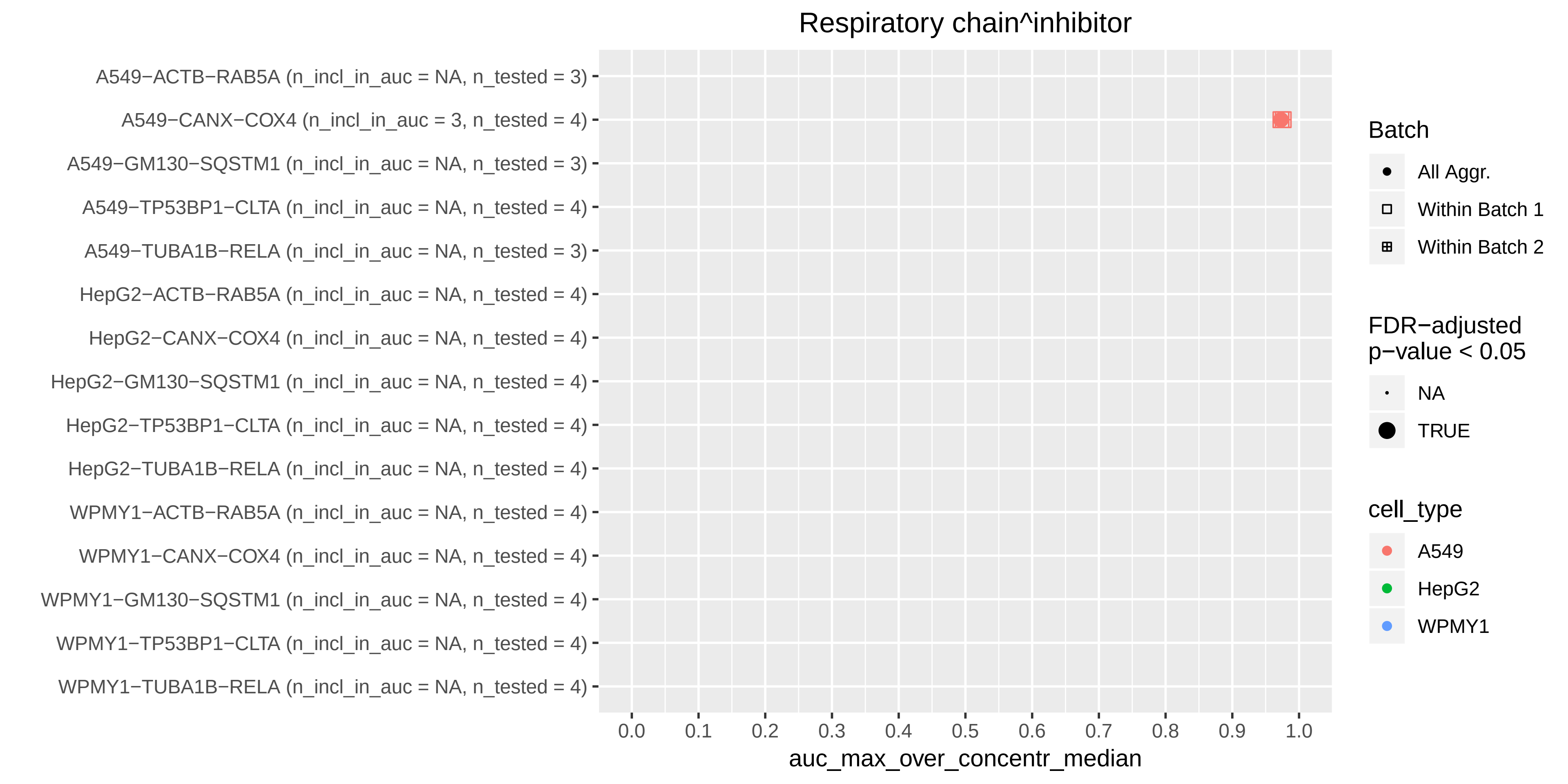
Respiratory chain^inhibitor
A549−ACTB−RAB5A (n_incl_in_auc = NA, n_tested = 3)
A549−CANX−COX4 (n_incl_in_auc = 3, n_tested = 4)
A549−GM130−SQSTM1 (n_incl_in_auc = NA, n_tested = 3)
A549−TP53BP1−CLTA (n_incl_in_auc = NA, n_tested = 4)
A549−TUBA1B−RELA (n_incl_in_auc = NA, n_tested = 3)
HepG2−ACTB−RAB5A (n_incl_in_auc = NA, n_tested = 4)
HepG2−CANX−COX4 (n_incl_in_auc = NA, n_tested = 4)
HepG2−GM130−SQSTM1 (n_incl_in_auc = NA, n_tested = 4)
HepG2−TP53BP1−CLTA (n_incl_in_auc = NA, n_tested = 4)
HepG2−TUBA1B−RELA (n_incl_in_auc = NA, n_tested = 4)
WPMY1−ACTB−RAB5A (n_incl_in_auc = NA, n_tested = 4)
WPMY1−CANX−COX4 (n_incl_in_auc = NA, n_tested = 4)
WPMY1−GM130−SQSTM1 (n_incl_in_auc = NA, n_tested = 4)
WPMY1−TP53BP1−CLTA (n_incl_in_auc = NA, n_tested = 4)
WPMY1−TUBA1B−RELA (n_incl_in_auc = NA, n_tested = 4)
0.0 0.1 0.2 0.3 0.4 0.5 0.6 0.7 0.8 0.9 1.0
auc_max_over_concentr_median
Batch
All Aggr.
Within Batch 1
Within Batch 2
FDR−adjusted p−value < 0.05
NA
TRUE
cell_type
A549
HepG2
WPMY1

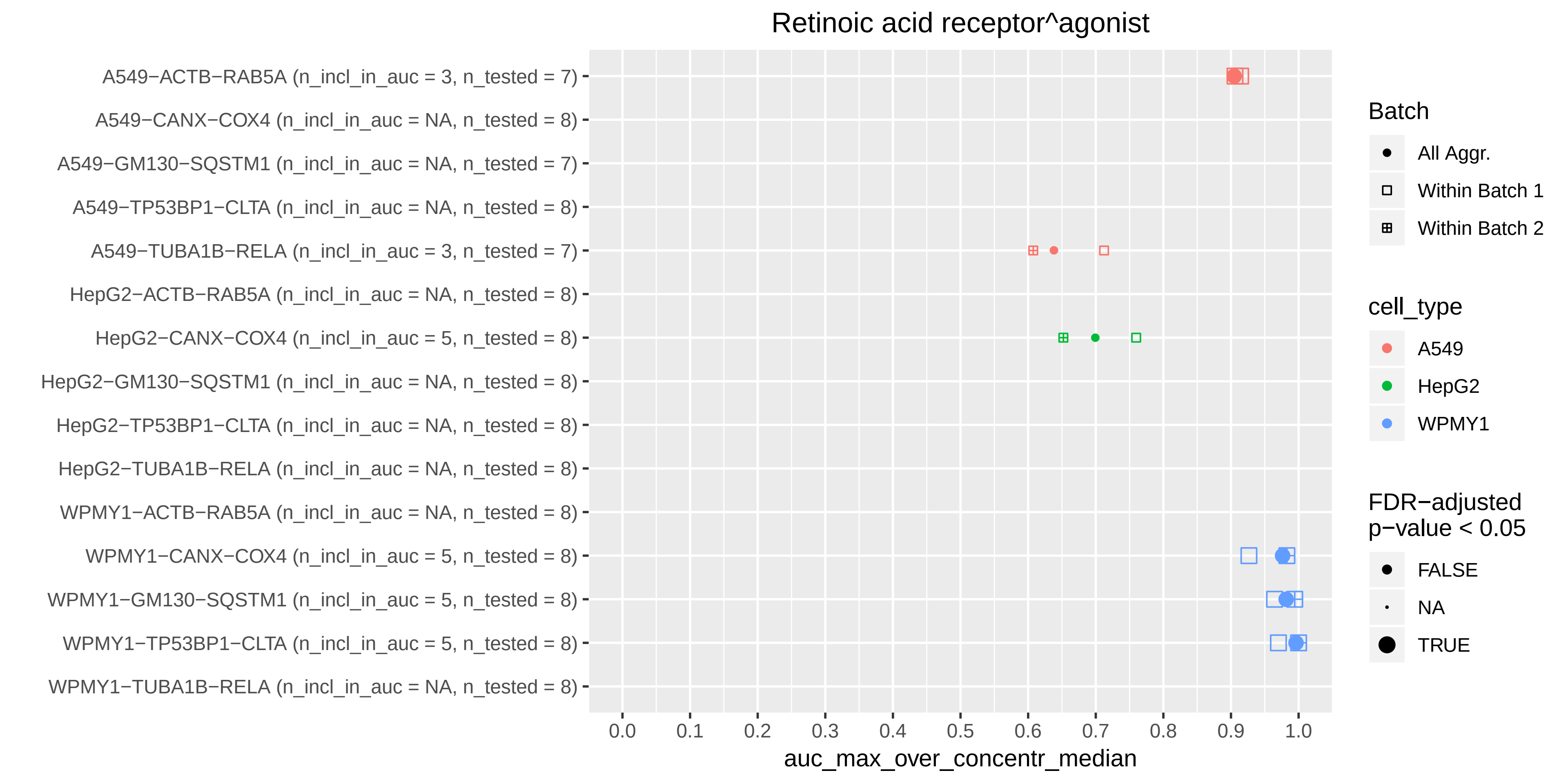
Retinoic acid receptor^agonist
A549−ACTB−RAB5A (n_incl_in_auc = 3, n_tested = 7)
A549−CANX−COX4 (n_incl_in_auc = NA, n_tested = 8)
A549−GM130−SQSTM1 (n_incl_in_auc = NA, n_tested = 7)
A549−TP53BP1−CLTA (n_incl_in_auc = NA, n_tested = 8)
A549−TUBA1B−RELA (n_incl_in_auc = 3, n_tested = 7)
HepG2−ACTB−RAB5A (n_incl_in_auc = NA, n_tested = 8)
HepG2−CANX−COX4 (n_incl_in_auc = 5, n_tested = 8)
HepG2−GM130−SQSTM1 (n_incl_in_auc = NA, n_tested = 8)
HepG2−TP53BP1−CLTA (n_incl_in_auc = NA, n_tested = 8)
HepG2−TUBA1B−RELA (n_incl_in_auc = NA, n_tested = 8)
WPMY1−ACTB−RAB5A (n_incl_in_auc = NA, n_tested = 8)
WPMY1−CANX−COX4 (n_incl_in_auc = 5, n_tested = 8)
WPMY1−GM130−SQSTM1 (n_incl_in_auc = 5, n_tested = 8)
WPMY1−TP53BP1−CLTA (n_incl_in_auc = 5, n_tested = 8)
WPMY1−TUBA1B−RELA (n_incl_in_auc = NA, n_tested = 8)
0.0
0.1
0.2
0.3
0.4
0.5
0.6
0.7
0.8
0.9
1.0
auc_max_over_concentr_median
Batch
All Aggr.
Within Batch 1
Within Batch 2
cell_type
A549
HepG2
WPMY1
FDR−adjusted
p−value < 0.05
FALSE
NA
TRUE

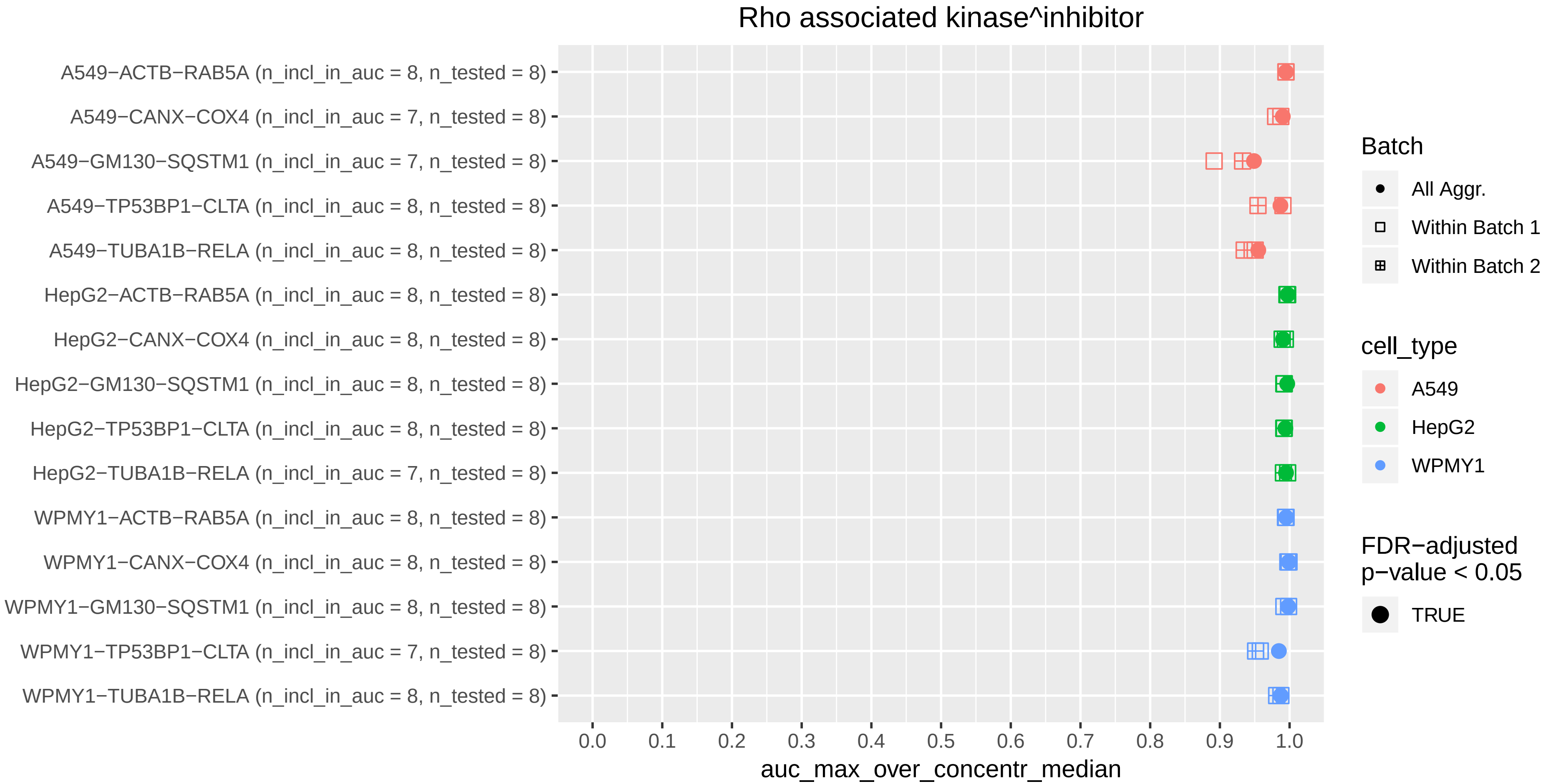

Rho associated kinase^inhibitor
A549−ACTB−RAB5A (n_incl_in_auc = 8, n_tested = 8)
A549−CANX−COX4 (n_incl_in_auc = 7, n_tested = 8)
A549−GM130−SQSTM1 (n_incl_in_auc = 7, n_tested = 8)
A549−TP53BP1−CLTA (n_incl_in_auc = 8, n_tested = 8)
A549−TUBA1B−RELA (n_incl_in_auc = 8, n_tested = 8)
HepG2−ACTB−RAB5A (n_incl_in_auc = 8, n_tested = 8)
HepG2−CANX−COX4 (n_incl_in_auc = 8, n_tested = 8)
HepG2−GM130−SQSTM1 (n_incl_in_auc = 8, n_tested = 8)
HepG2−TP53BP1−CLTA (n_incl_in_auc = 8, n_tested = 8)
HepG2−TUBA1B−RELA (n_incl_in_auc = 7, n_tested = 8)
WPMY1−ACTB−RAB5A (n_incl_in_auc = 8, n_tested = 8)
WPMY1−CANX−COX4 (n_incl_in_auc = 8, n_tested = 8)
WPMY1−GM130−SQSTM1 (n_incl_in_auc = 8, n_tested = 8)
WPMY1−TP53BP1−CLTA (n_incl_in_auc = 7, n_tested = 8)
WPMY1−TUBA1B−RELA (n_incl_in_auc = 8, n_tested = 8)
0.0
0.1
0.2
0.3
0.4
0.5
0.6
0.7
0.8
0.9
1.0
auc_max_over_concentr_median
Batch
All Aggr.
Within Batch 1
Within Batch 2
cell_type
A549
HepG2
WPMY1
FDR−adjusted p−value < 0.05
TRUE

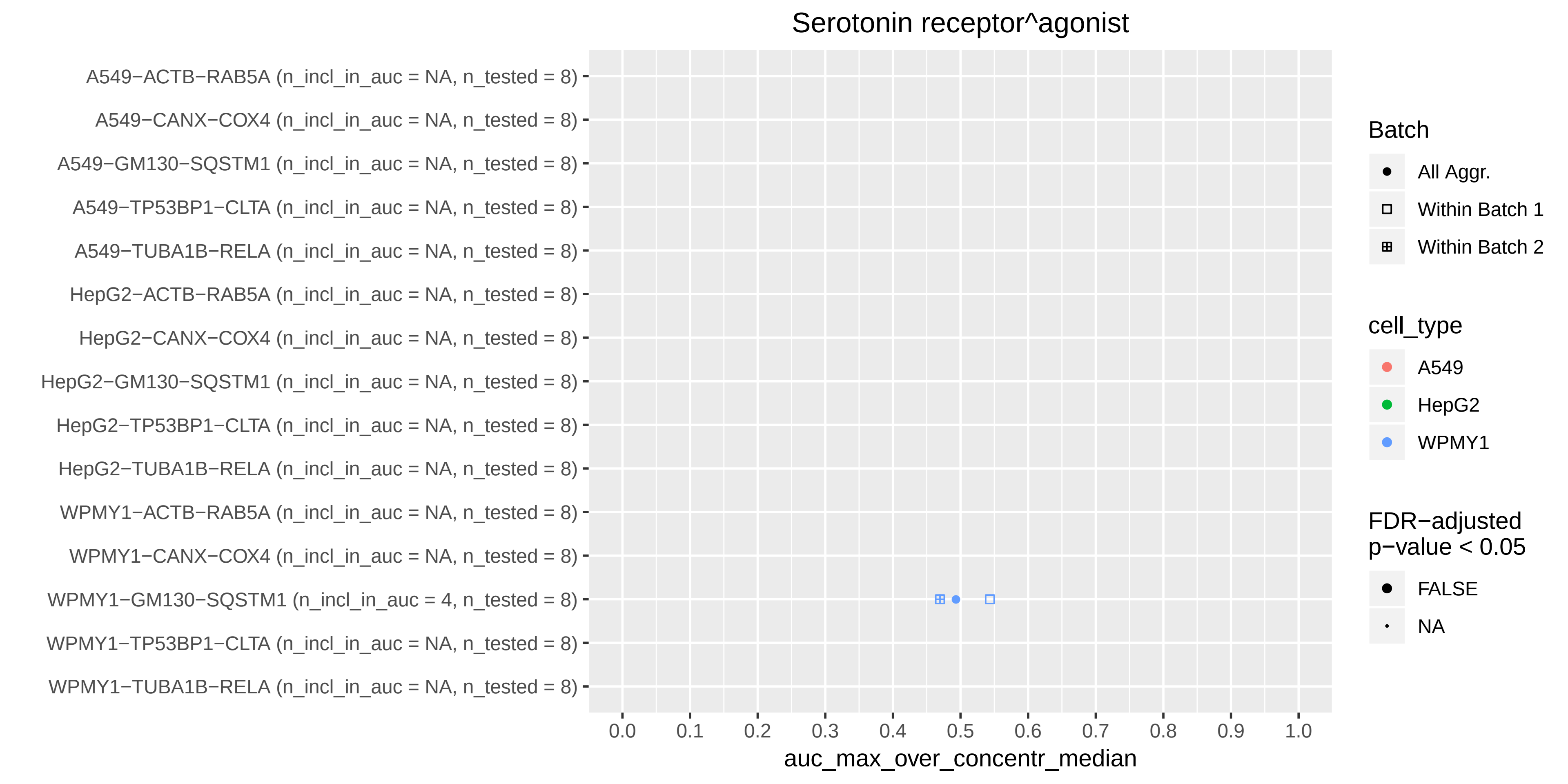

Serotonin receptor^agonist
A549−ACTB−RAB5A (n_incl_in_auc = NA, n_tested = 8)
A549−CANX−COX4 (n_incl_in_auc = NA, n_tested = 8)
A549−GM130−SQSTM1 (n_incl_in_auc = NA, n_tested = 8)
A549−TP53BP1−CLTA (n_incl_in_auc = NA, n_tested = 8)
A549−TUBA1B−RELA (n_incl_in_auc = NA, n_tested = 8)
HepG2−ACTB−RAB5A (n_incl_in_auc = NA, n_tested = 8)
HepG2−CANX−COX4 (n_incl_in_auc = NA, n_tested = 8)
HepG2−GM130−SQSTM1 (n_incl_in_auc = NA, n_tested = 8)
HepG2−TP53BP1−CLTA (n_incl_in_auc = NA, n_tested = 8)
HepG2−TUBA1B−RELA (n_incl_in_auc = NA, n_tested = 8)
WPMY1−ACTB−RAB5A (n_incl_in_auc = NA, n_tested = 8)
WPMY1−CANX−COX4 (n_incl_in_auc = NA, n_tested = 8)
WPMY1−GM130−SQSTM1 (n_incl_in_auc = 4, n_tested = 8)
WPMY1−TP53BP1−CLTA (n_incl_in_auc = NA, n_tested = 8)
WPMY1−TUBA1B−RELA (n_incl_in_auc = NA, n_tested = 8)
0.0 0.1 0.2 0.3 0.4 0.5 0.6 0.7 0.8 0.9 1.0
auc_max_over_concentr_median
Batch
All Aggr.
Within Batch 1
Within Batch 2
cell_type
A549
HepG2
WPMY1
FDR−adjusted p−value < 0.05
FALSE
NA

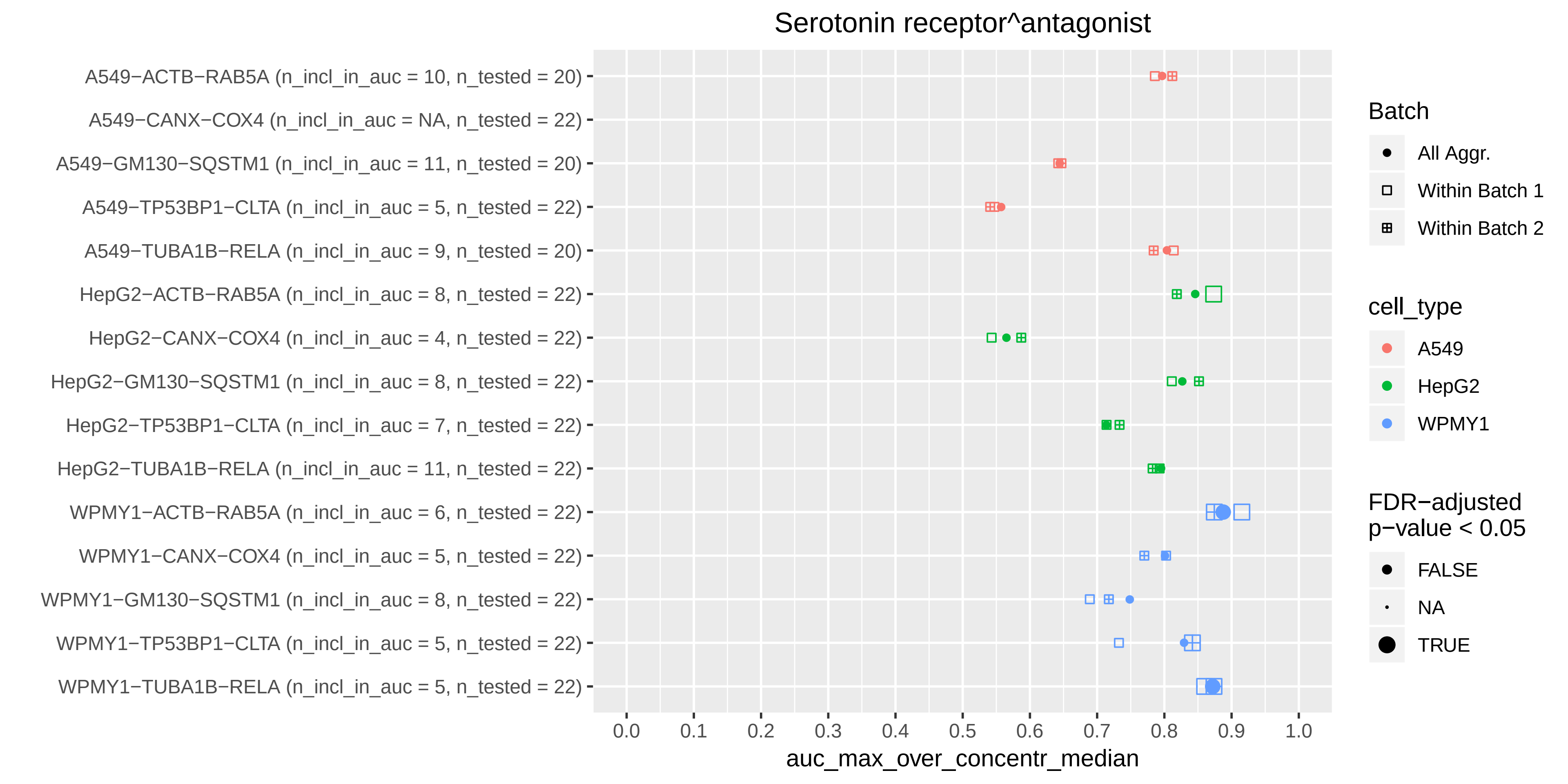
Serotonin receptor^antagonist
A549−ACTB−RAB5A (n_incl_in_auc = 10, n_tested = 20)
A549−CANX−COX4 (n_incl_in_auc = NA, n_tested = 22)
A549−GM130−SQSTM1 (n_incl_in_auc = 11, n_tested = 20)
A549−TP53BP1−CLTA (n_incl_in_auc = 5, n_tested = 22)
A549−TUBA1B−RELA (n_incl_in_auc = 9, n_tested = 20)
HepG2−ACTB−RAB5A (n_incl_in_auc = 8, n_tested = 22)
HepG2−CANX−COX4 (n_incl_in_auc = 4, n_tested = 22)
HepG2−GM130−SQSTM1 (n_incl_in_auc = 8, n_tested = 22)
HepG2−TP53BP1−CLTA (n_incl_in_auc = 7, n_tested = 22)
HepG2−TUBA1B−RELA (n_incl_in_auc = 11, n_tested = 22)
WPMY1−ACTB−RAB5A (n_incl_in_auc = 6, n_tested = 22)
WPMY1−CANX−COX4 (n_incl_in_auc = 5, n_tested = 22)
WPMY1−GM130−SQSTM1 (n_incl_in_auc = 8, n_tested = 22)
WPMY1−TP53BP1−CLTA (n_incl_in_auc = 5, n_tested = 22)
WPMY1−TUBA1B−RELA (n_incl_in_auc = 5, n_tested = 22)
0.0
0.1
0.2
0.3
0.4
0.5
0.6
0.7
0.8
0.9
1.0
auc_max_over_concentr_median
Batch
All Aggr.
Within Batch 1
Within Batch 2
cell_type
A549
HepG2
WPMY1
FDR−adjusted
p−value < 0.05
FALSE
NA
TRUE

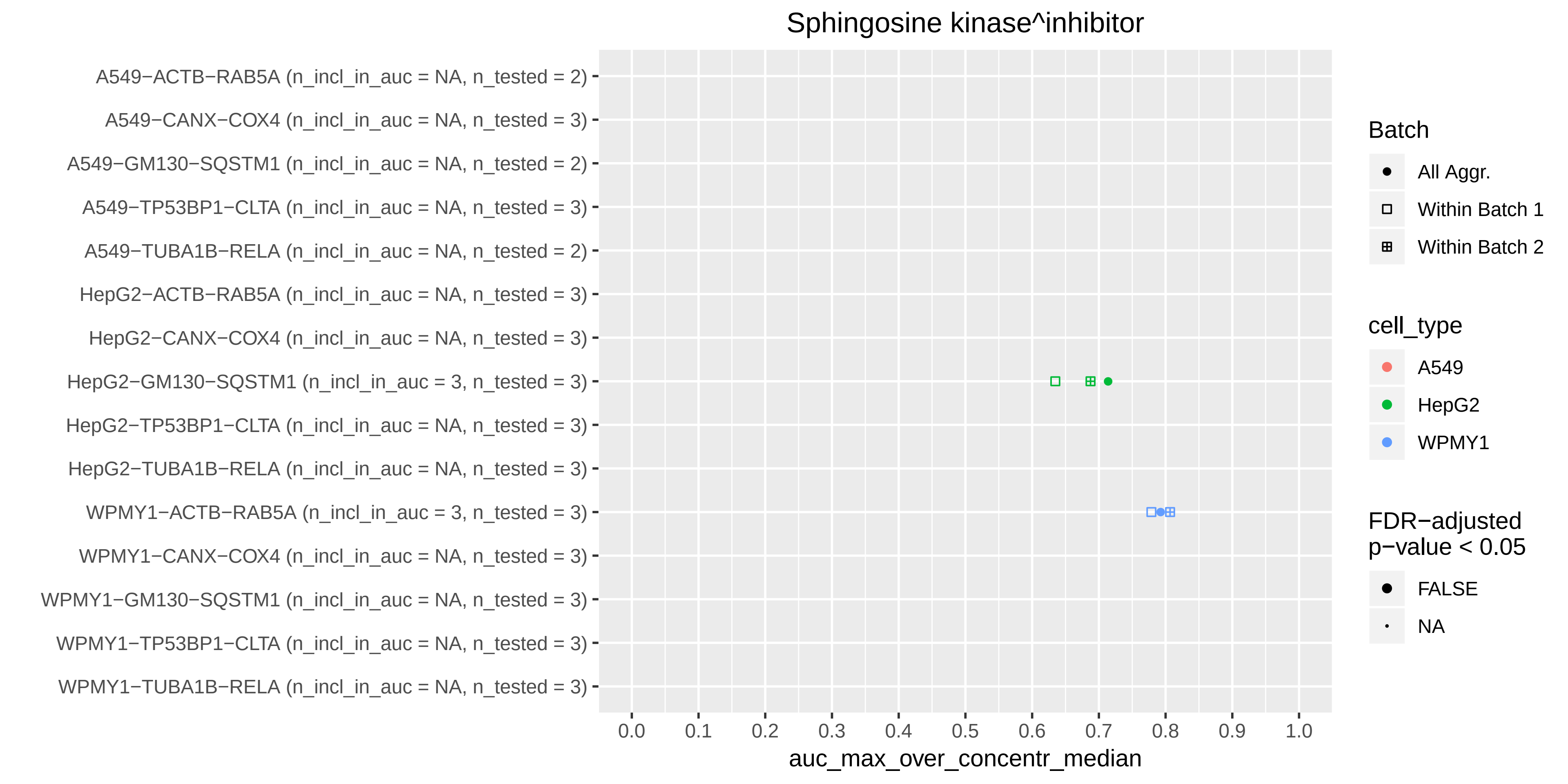

Sphingosine kinase^inhibitor
A549−ACTB−RAB5A (n_incl_in_auc = NA, n_tested = 2)
A549−CANX−COX4 (n_incl_in_auc = NA, n_tested = 3)
A549−GM130−SQSTM1 (n_incl_in_auc = NA, n_tested = 2)
A549−TP53BP1−CLTA (n_incl_in_auc = NA, n_tested = 3)
A549−TUBA1B−RELA (n_incl_in_auc = NA, n_tested = 2)
HepG2−ACTB−RAB5A (n_incl_in_auc = NA, n_tested = 3)
HepG2−CANX−COX4 (n_incl_in_auc = NA, n_tested = 3)
HepG2−GM130−SQSTM1 (n_incl_in_auc = 3, n_tested = 3)
HepG2−TP53BP1−CLTA (n_incl_in_auc = NA, n_tested = 3)
HepG2−TUBA1B−RELA (n_incl_in_auc = NA, n_tested = 3)
WPMY1−ACTB−RAB5A (n_incl_in_auc = 3, n_tested = 3)
WPMY1−CANX−COX4 (n_incl_in_auc = NA, n_tested = 3)
WPMY1−GM130−SQSTM1 (n_incl_in_auc = NA, n_tested = 3)
WPMY1−TP53BP1−CLTA (n_incl_in_auc = NA, n_tested = 3)
WPMY1−TUBA1B−RELA (n_incl_in_auc = NA, n_tested = 3)
0.0
0.1
0.2
0.3
0.4
0.5
0.6
0.7
0.8
0.9
1.0
auc_max_over_concentr_median
Batch
All Aggr.
Within Batch 1
Within Batch 2
cell_type
A549
HepG2
WPMY1
FDR−adjusted
p−value < 0.05
FALSE
NA

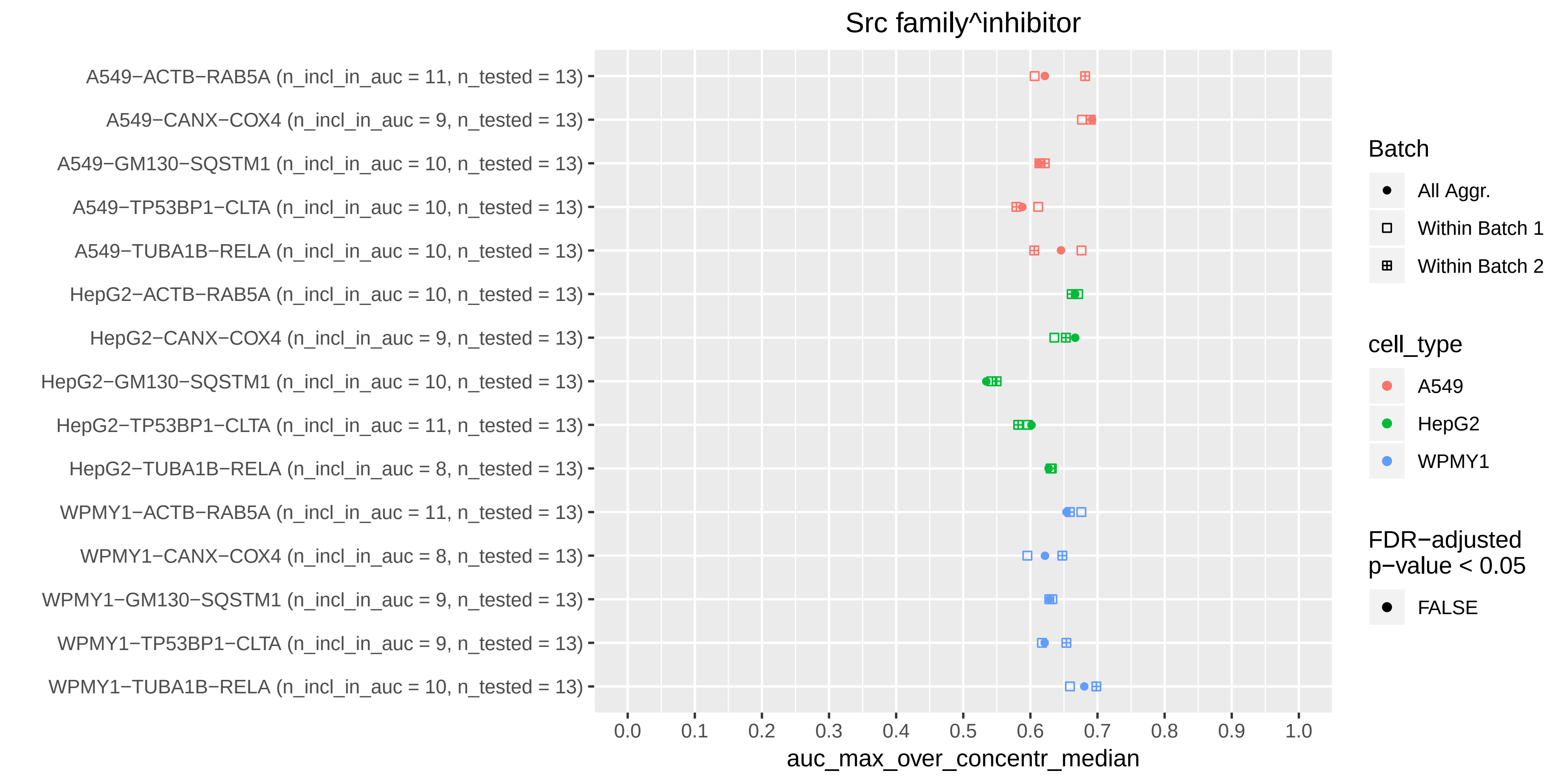
Src family^inhibitor
A549−ACTB−RAB5A (n_incl_in_auc = 11, n_tested = 13)
A549−CANX−COX4 (n_incl_in_auc = 9, n_tested = 13)
A549−GM130−SQSTM1 (n_incl_in_auc = 10, n_tested = 13)
A549−TP53BP1−CLTA (n_incl_in_auc = 10, n_tested = 13)
A549−TUBA1B−RELA (n_incl_in_auc = 10, n_tested = 13)
HepG2−ACTB−RAB5A (n_incl_in_auc = 10, n_tested = 13)
HepG2−CANX−COX4 (n_incl_in_auc = 9, n_tested = 13)
HepG2−GM130−SQSTM1 (n_incl_in_auc = 10, n_tested = 13)
HepG2−TP53BP1−CLTA (n_incl_in_auc = 11, n_tested = 13)
HepG2−TUBA1B−RELA (n_incl_in_auc = 8, n_tested = 13)
WPMY1−ACTB−RAB5A (n_incl_in_auc = 11, n_tested = 13)
WPMY1−CANX−COX4 (n_incl_in_auc = 8, n_tested = 13)
WPMY1−GM130−SQSTM1 (n_incl_in_auc = 9, n_tested = 13)
WPMY1−TP53BP1−CLTA (n_incl_in_auc = 9, n_tested = 13)
WPMY1−TUBA1B−RELA (n_incl_in_auc = 10, n_tested = 13)
0.0 0.1 0.2 0.3 0.4 0.5 0.6 0.7 0.8 0.9 1.0
auc_max_over_concentr_median
Batch
All Aggr.
Within Batch 1
Within Batch 2
cell_type
A549
HepG2
WPMY1
FDR−adjusted p−value < 0.05
FALSE

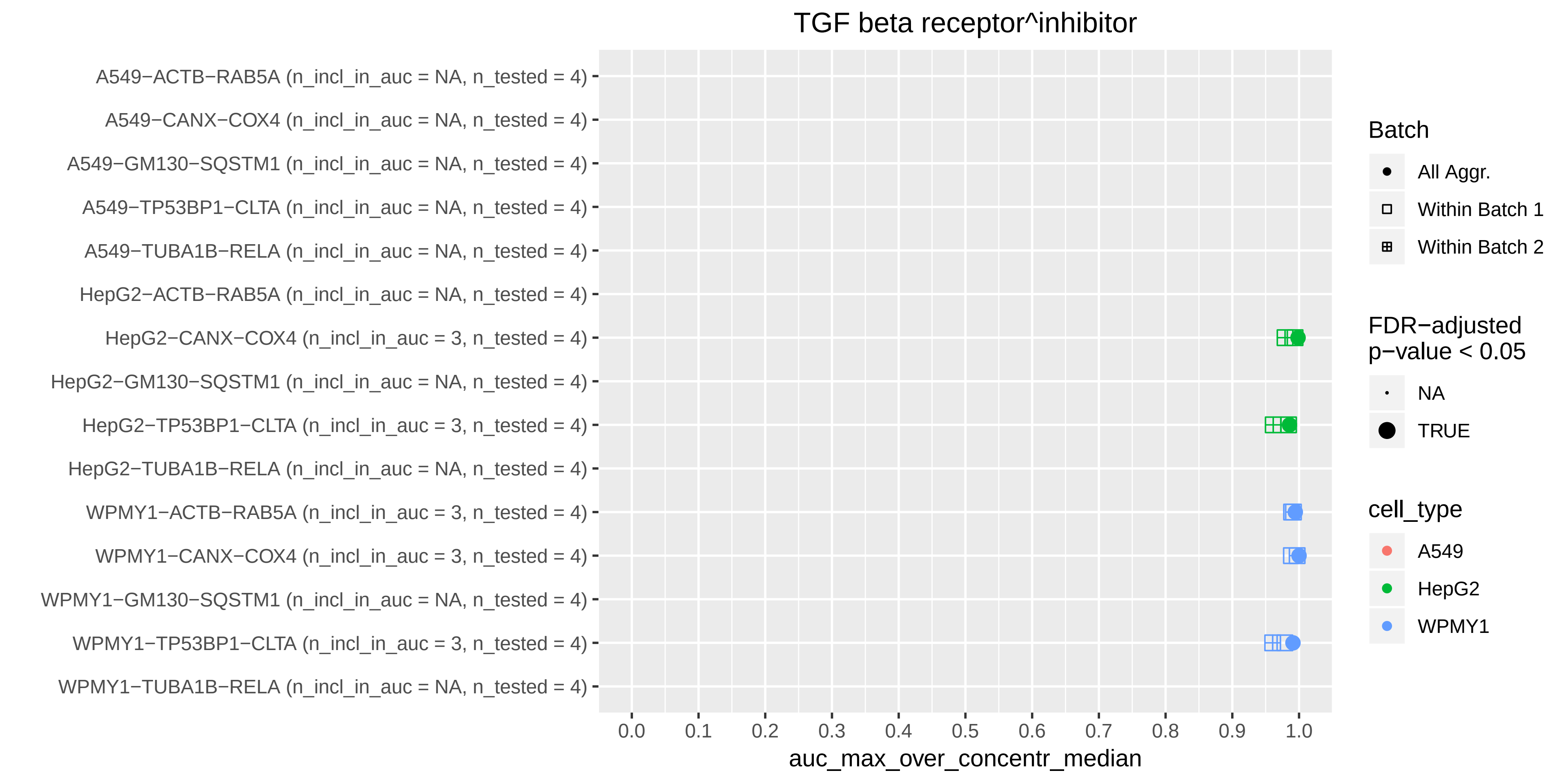

TGF beta receptor^inhibitor
A549−ACTB−RAB5A (n_incl_in_auc = NA, n_tested = 4)
A549−CANX−COX4 (n_incl_in_auc = NA, n_tested = 4)
A549−GM130−SQSTM1 (n_incl_in_auc = NA, n_tested = 4)
A549−TP53BP1−CLTA (n_incl_in_auc = NA, n_tested = 4)
A549−TUBA1B−RELA (n_incl_in_auc = NA, n_tested = 4)
HepG2−ACTB−RAB5A (n_incl_in_auc = NA, n_tested = 4)
HepG2−CANX−COX4 (n_incl_in_auc = 3, n_tested = 4)
HepG2−GM130−SQSTM1 (n_incl_in_auc = NA, n_tested = 4)
HepG2−TP53BP1−CLTA (n_incl_in_auc = 3, n_tested = 4)
HepG2−TUBA1B−RELA (n_incl_in_auc = NA, n_tested = 4)
WPMY1−ACTB−RAB5A (n_incl_in_auc = 3, n_tested = 4)
WPMY1−CANX−COX4 (n_incl_in_auc = 3, n_tested = 4)
WPMY1−GM130−SQSTM1 (n_incl_in_auc = NA, n_tested = 4)
WPMY1−TP53BP1−CLTA (n_incl_in_auc = 3, n_tested = 4)
WPMY1−TUBA1B−RELA (n_incl_in_auc = NA, n_tested = 4)
0.0
0.1
0.2
0.3
0.4
0.5
0.6
0.7
0.8
0.9
1.0
auc_max_over_concentr_median
Batch
All Aggr.
Within Batch 1
Within Batch 2
FDR−adjusted
p−value < 0.05
NA
TRUE
cell_type
A549
HepG2
WPMY1

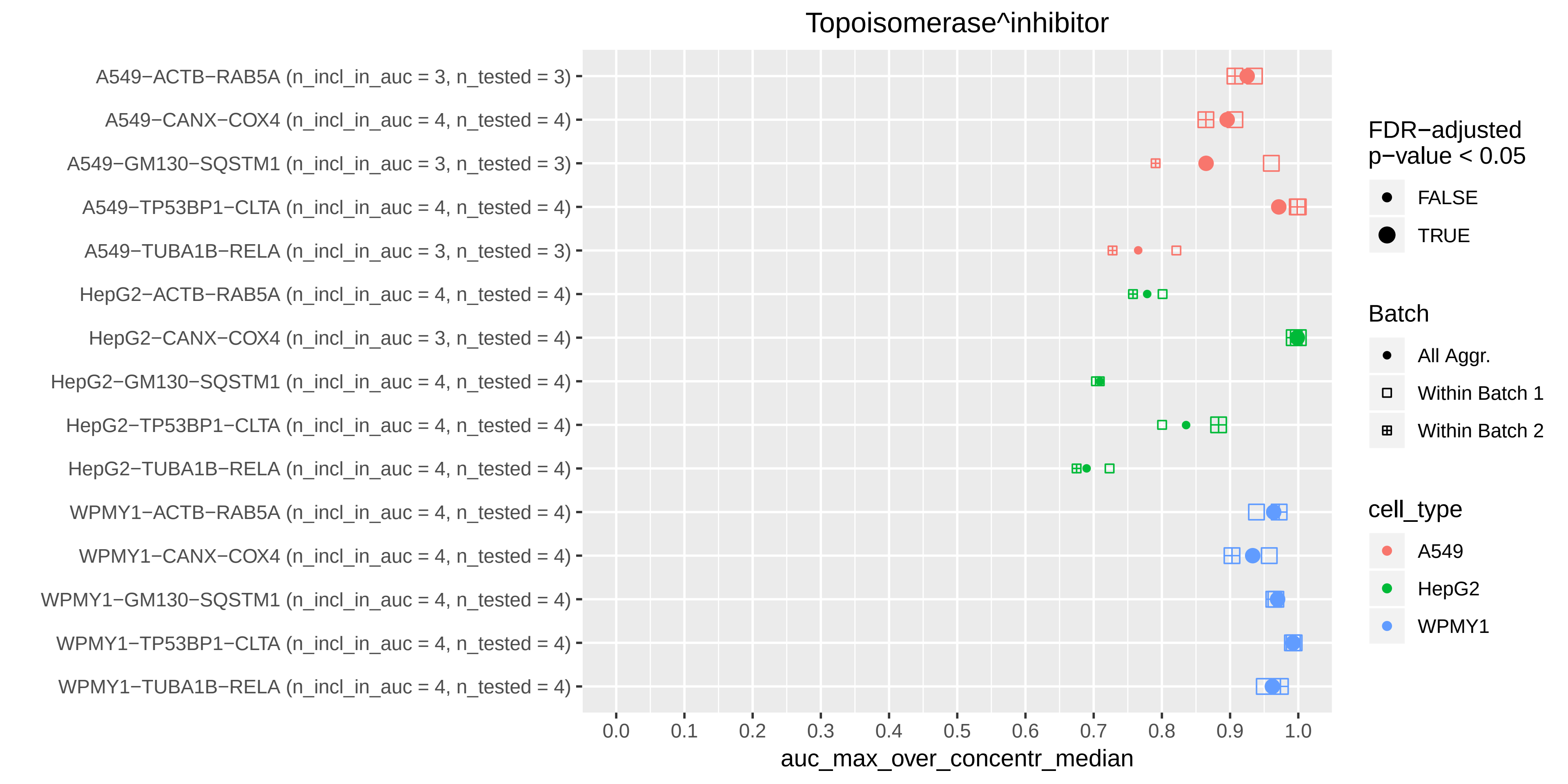

Topoisomerase^inhibitor
A549−ACTB−RAB5A (n_incl_in_auc = 3, n_tested = 3)
A549−CANX−COX4 (n_incl_in_auc = 4, n_tested = 4)
A549−GM130−SQSTM1 (n_incl_in_auc = 3, n_tested = 3)
A549−TP53BP1−CLTA (n_incl_in_auc = 4, n_tested = 4)
A549−TUBA1B−RELA (n_incl_in_auc = 3, n_tested = 3)
HepG2−ACTB−RAB5A (n_incl_in_auc = 4, n_tested = 4)
HepG2−CANX−COX4 (n_incl_in_auc = 3, n_tested = 4)
HepG2−GM130−SQSTM1 (n_incl_in_auc = 4, n_tested = 4)
HepG2−TP53BP1−CLTA (n_incl_in_auc = 4, n_tested = 4)
HepG2−TUBA1B−RELA (n_incl_in_auc = 4, n_tested = 4)
WPMY1−ACTB−RAB5A (n_incl_in_auc = 4, n_tested = 4)
WPMY1−CANX−COX4 (n_incl_in_auc = 4, n_tested = 4)
WPMY1−GM130−SQSTM1 (n_incl_in_auc = 4, n_tested = 4)
WPMY1−TP53BP1−CLTA (n_incl_in_auc = 4, n_tested = 4)
WPMY1−TUBA1B−RELA (n_incl_in_auc = 4, n_tested = 4)
0.0
0.1
0.2
0.3
0.4
0.5
0.6
0.7
0.8
0.9
1.0
auc_max_over_concentr_median
FDR−adjusted p−value < 0.05
FALSE
TRUE
Batch
All Aggr.
Within Batch 1
Within Batch 2
cell_type
A549
HepG2
WPMY1

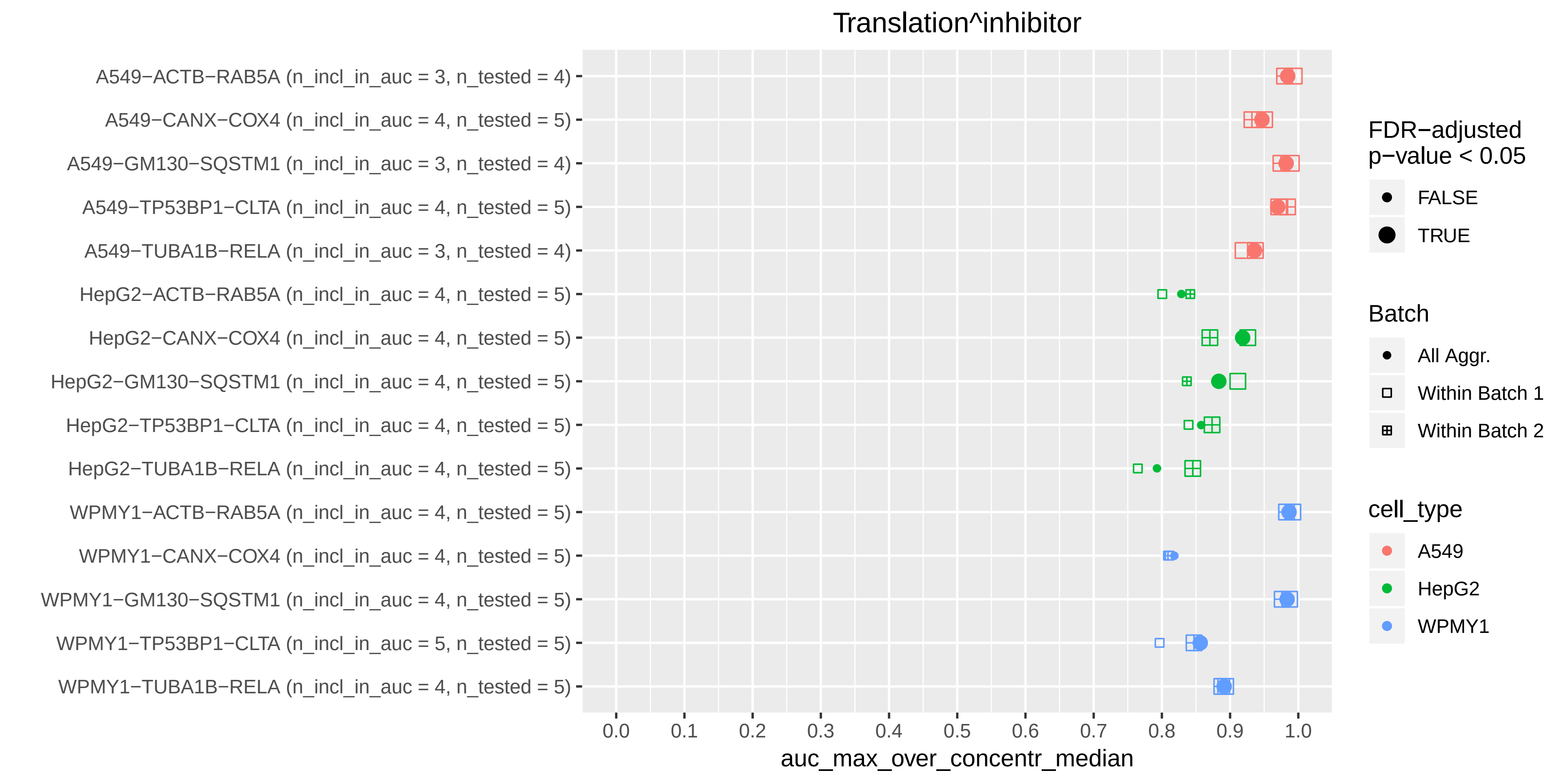
Translation^inhibitor
A549−ACTB−RAB5A (n_incl_in_auc = 3, n_tested = 4)
A549−CANX−COX4 (n_incl_in_auc = 4, n_tested = 5)
A549−GM130−SQSTM1 (n_incl_in_auc = 3, n_tested = 4)
A549−TP53BP1−CLTA (n_incl_in_auc = 4, n_tested = 5)
A549−TUBA1B−RELA (n_incl_in_auc = 3, n_tested = 4)
HepG2−ACTB−RAB5A (n_incl_in_auc = 4, n_tested = 5)
HepG2−CANX−COX4 (n_incl_in_auc = 4, n_tested = 5)
HepG2−GM130−SQSTM1 (n_incl_in_auc = 4, n_tested = 5)
HepG2−TP53BP1−CLTA (n_incl_in_auc = 4, n_tested = 5)
HepG2−TUBA1B−RELA (n_incl_in_auc = 4, n_tested = 5)
WPMY1−ACTB−RAB5A (n_incl_in_auc = 4, n_tested = 5)
WPMY1−CANX−COX4 (n_incl_in_auc = 4, n_tested = 5)
WPMY1−GM130−SQSTM1 (n_incl_in_auc = 4, n_tested = 5)
WPMY1−TP53BP1−CLTA (n_incl_in_auc = 5, n_tested = 5)
WPMY1−TUBA1B−RELA (n_incl_in_auc = 4, n_tested = 5)
0.0
0.1
0.2
0.3
0.4
0.5
0.6
0.7
0.8
0.9
1.0
auc_max_over_concentr_median
FDR−adjusted p−value < 0.05
FALSE
TRUE
Batch
All Aggr.
Within Batch 1
Within Batch 2
cell_type
A549
HepG2
WPMY1

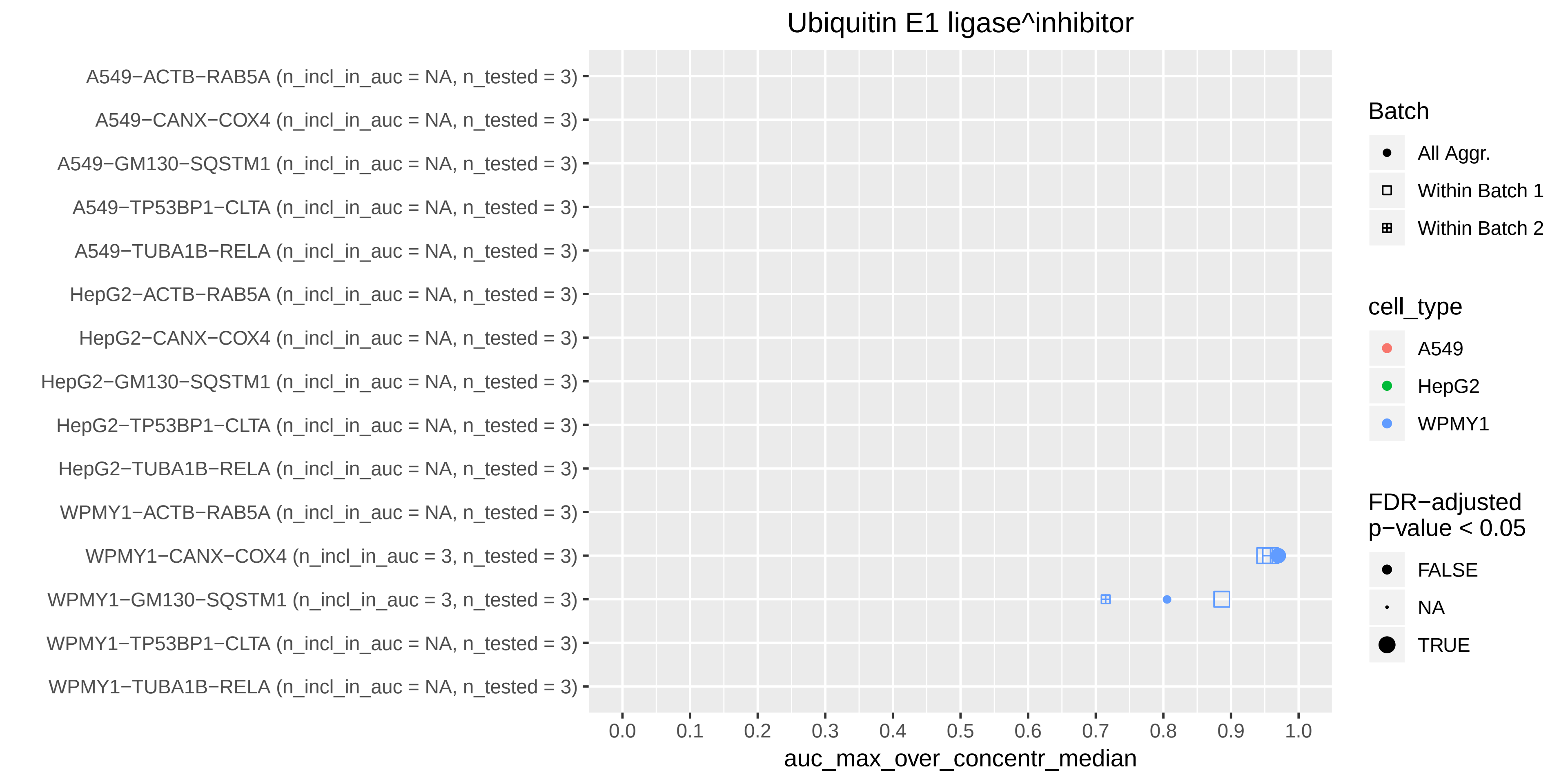

Ubiquitin E1 ligase^inhibitor
A549−ACTB−RAB5A (n_incl_in_auc = NA, n_tested = 3)
A549−CANX−COX4 (n_incl_in_auc = NA, n_tested = 3)
A549−GM130−SQSTM1 (n_incl_in_auc = NA, n_tested = 3)
A549−TP53BP1−CLTA (n_incl_in_auc = NA, n_tested = 3)
A549−TUBA1B−RELA (n_incl_in_auc = NA, n_tested = 3)
HepG2−ACTB−RAB5A (n_incl_in_auc = NA, n_tested = 3)
HepG2−CANX−COX4 (n_incl_in_auc = NA, n_tested = 3)
HepG2−GM130−SQSTM1 (n_incl_in_auc = NA, n_tested = 3)
HepG2−TP53BP1−CLTA (n_incl_in_auc = NA, n_tested = 3)
HepG2−TUBA1B−RELA (n_incl_in_auc = NA, n_tested = 3)
WPMY1−ACTB−RAB5A (n_incl_in_auc = NA, n_tested = 3)
WPMY1−CANX−COX4 (n_incl_in_auc = 3, n_tested = 3)
WPMY1−GM130−SQSTM1 (n_incl_in_auc = 3, n_tested = 3)
WPMY1−TP53BP1−CLTA (n_incl_in_auc = NA, n_tested = 3)
WPMY1−TUBA1B−RELA (n_incl_in_auc = NA, n_tested = 3)
0.0
0.1
0.2
0.3
0.4
0.5
0.6
0.7
0.8
0.9
1.0
auc_max_over_concentr_median
Batch
All Aggr.
Within Batch 1
Within Batch 2
cell_type
A549
HepG2
WPMY1
FDR−adjusted p−value < 0.05
FALSE
NA
TRUE

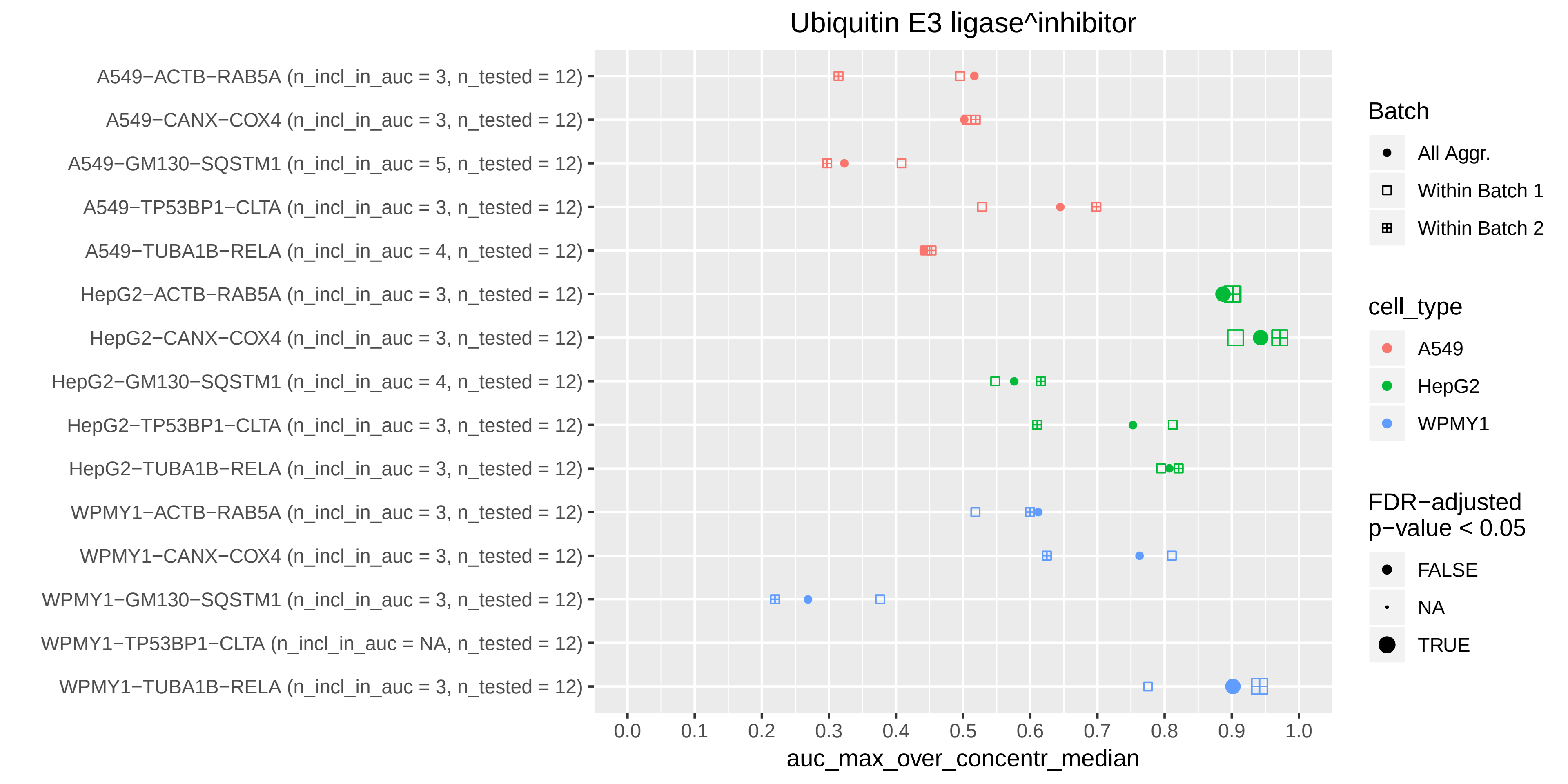

Ubiquitin E3 ligase^inhibitor
A549−ACTB−RAB5A (n_incl_in_auc = 3, n_tested = 12)
A549−CANX−COX4 (n_incl_in_auc = 3, n_tested = 12)
A549−GM130−SQSTM1 (n_incl_in_auc = 5, n_tested = 12)
A549−TP53BP1−CLTA (n_incl_in_auc = 3, n_tested = 12)
A549−TUBA1B−RELA (n_incl_in_auc = 4, n_tested = 12)
HepG2−ACTB−RAB5A (n_incl_in_auc = 3, n_tested = 12)
HepG2−CANX−COX4 (n_incl_in_auc = 3, n_tested = 12)
HepG2−GM130−SQSTM1 (n_incl_in_auc = 4, n_tested = 12)
HepG2−TP53BP1−CLTA (n_incl_in_auc = 3, n_tested = 12)
HepG2−TUBA1B−RELA (n_incl_in_auc = 3, n_tested = 12)
WPMY1−ACTB−RAB5A (n_incl_in_auc = 3, n_tested = 12)
WPMY1−CANX−COX4 (n_incl_in_auc = 3, n_tested = 12)
WPMY1−GM130−SQSTM1 (n_incl_in_auc = 3, n_tested = 12)
WPMY1−TP53BP1−CLTA (n_incl_in_auc = NA, n_tested = 12)
WPMY1−TUBA1B−RELA (n_incl_in_auc = 3, n_tested = 12)
0.0
0.1
0.2
0.3
0.4
0.5
0.6
0.7
0.8
0.9
1.0
auc_max_over_concentr_median
Batch
All Aggr.
Within Batch 1
Within Batch 2
cell_type
A549
HepG2
WPMY1
FDR−adjusted p−value < 0.05
FALSE
NA
TRUE

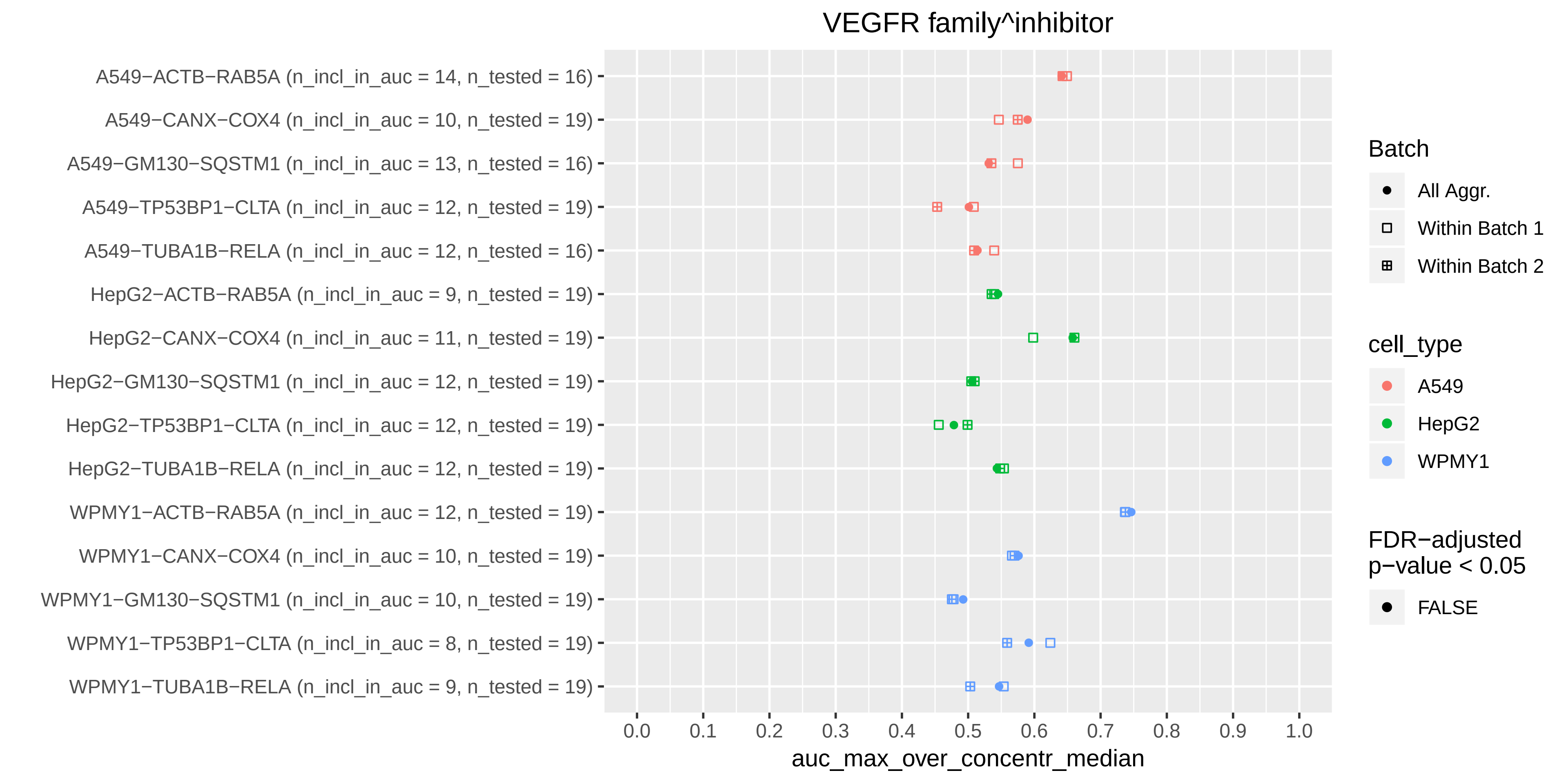

VEGFR family^inhibitor
A549−ACTB−RAB5A (n_incl_in_auc = 14, n_tested = 16)
A549−CANX−COX4 (n_incl_in_auc = 10, n_tested = 19)
A549−GM130−SQSTM1 (n_incl_in_auc = 13, n_tested = 16)
A549−TP53BP1−CLTA (n_incl_in_auc = 12, n_tested = 19)
A549−TUBA1B−RELA (n_incl_in_auc = 12, n_tested = 16)
HepG2−ACTB−RAB5A (n_incl_in_auc = 9, n_tested = 19)
HepG2−CANX−COX4 (n_incl_in_auc = 11, n_tested = 19)
HepG2−GM130−SQSTM1 (n_incl_in_auc = 12, n_tested = 19)
HepG2−TP53BP1−CLTA (n_incl_in_auc = 12, n_tested = 19)
HepG2−TUBA1B−RELA (n_incl_in_auc = 12, n_tested = 19)
WPMY1−ACTB−RAB5A (n_incl_in_auc = 12, n_tested = 19)
WPMY1−CANX−COX4 (n_incl_in_auc = 10, n_tested = 19)
WPMY1−GM130−SQSTM1 (n_incl_in_auc = 10, n_tested = 19)
WPMY1−TP53BP1−CLTA (n_incl_in_auc = 8, n_tested = 19)
WPMY1−TUBA1B−RELA (n_incl_in_auc = 9, n_tested = 19)
0.0
0.1
0.2
0.3
0.4
0.5
0.6
0.7
0.8
0.9
1.0
auc_max_over_concentr_median
Batch
All Aggr.
Within Batch 1
Within Batch 2
cell_type
A549
HepG2
WPMY1
FDR−adjusted p−value < 0.05
FALSE

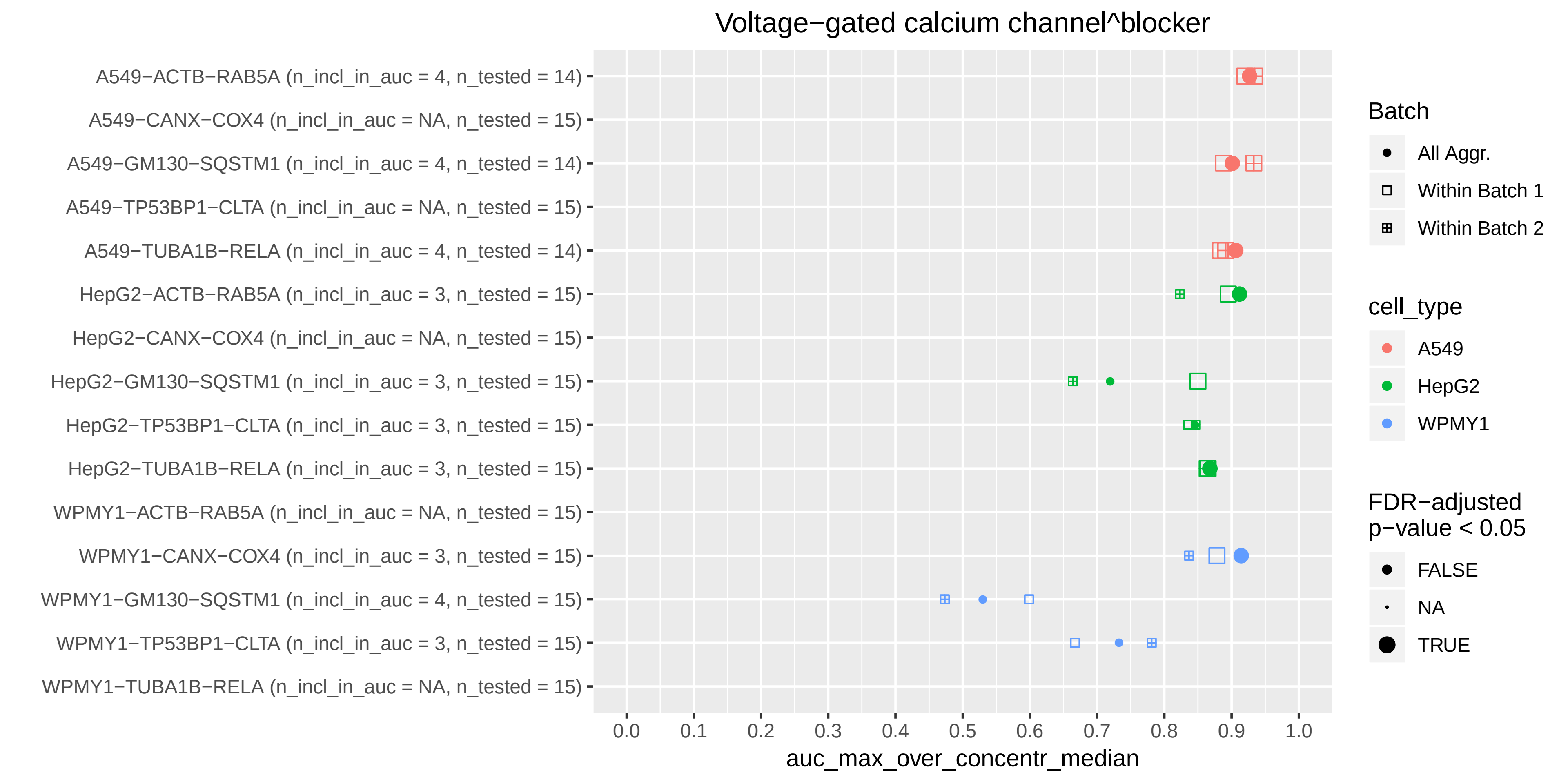
Voltage−gated calcium channel^blocker
A549−ACTB−RAB5A (n_incl_in_auc = 4, n_tested = 14)
A549−CANX−COX4 (n_incl_in_auc = NA, n_tested = 15)
A549−GM130−SQSTM1 (n_incl_in_auc = 4, n_tested = 14)
A549−TP53BP1−CLTA (n_incl_in_auc = NA, n_tested = 15)
A549−TUBA1B−RELA (n_incl_in_auc = 4, n_tested = 14)
HepG2−ACTB−RAB5A (n_incl_in_auc = 3, n_tested = 15)
HepG2−CANX−COX4 (n_incl_in_auc = NA, n_tested = 15)
HepG2−GM130−SQSTM1 (n_incl_in_auc = 3, n_tested = 15)
HepG2−TP53BP1−CLTA (n_incl_in_auc = 3, n_tested = 15)
HepG2−TUBA1B−RELA (n_incl_in_auc = 3, n_tested = 15)
WPMY1−ACTB−RAB5A (n_incl_in_auc = NA, n_tested = 15)
WPMY1−CANX−COX4 (n_incl_in_auc = 3, n_tested = 15)
WPMY1−GM130−SQSTM1 (n_incl_in_auc = 4, n_tested = 15)
WPMY1−TP53BP1−CLTA (n_incl_in_auc = 3, n_tested = 15)
WPMY1−TUBA1B−RELA (n_incl_in_auc = NA, n_tested = 15)
0.0
0.1
0.2
0.3
0.4
0.5
0.6
0.7
0.8
0.9
1.0
auc_max_over_concentr_median
Batch
All Aggr.
Within Batch 1
Within Batch 2
cell_type
A549
HepG2
WPMY1
FDR−adjusted
p−value < 0.05
FALSE
NA
TRUE

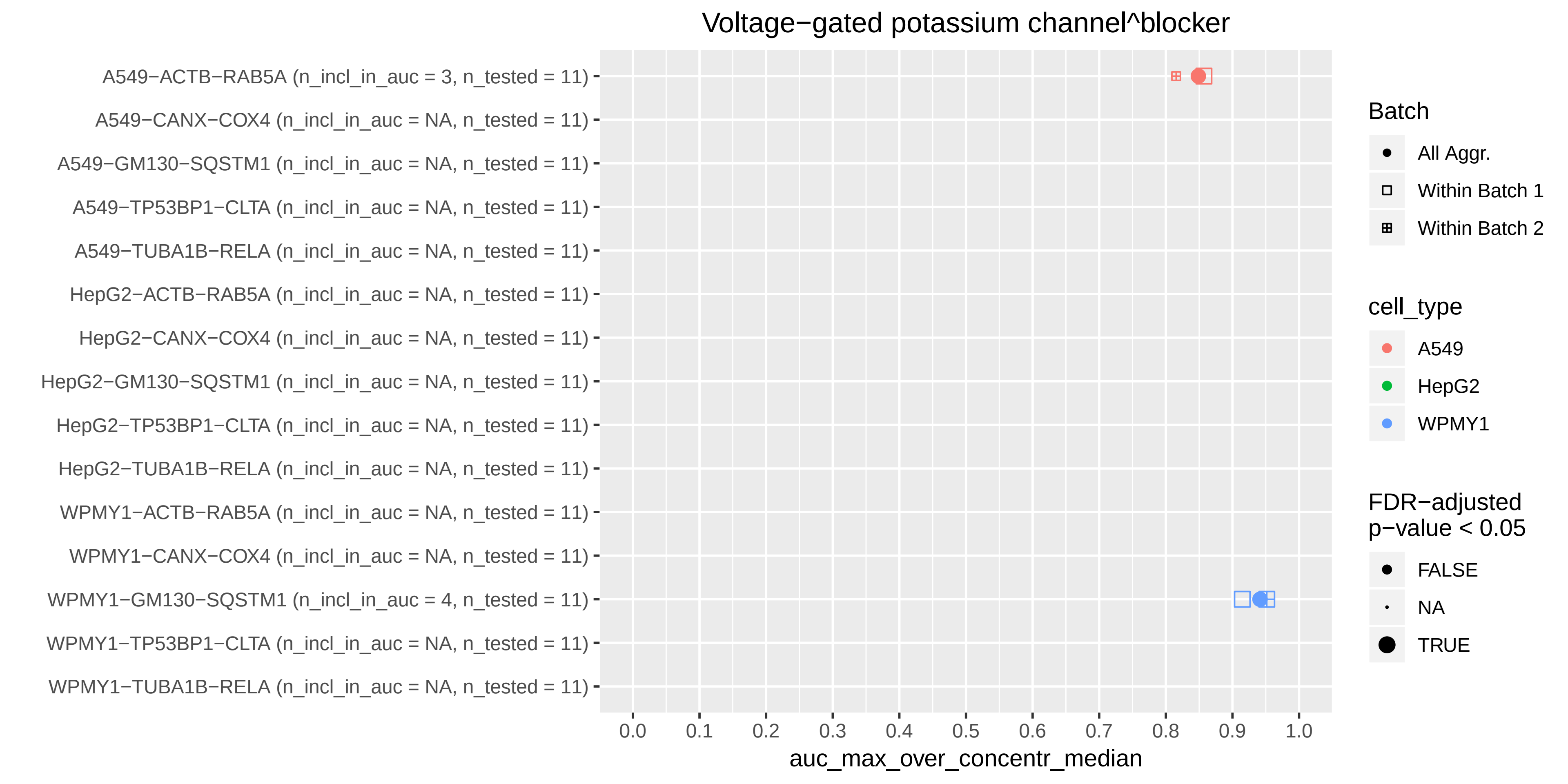
Voltage−gated potassium channel^blocker
A549−ACTB−RAB5A (n_incl_in_auc = 3, n_tested = 11)
A549−CANX−COX4 (n_incl_in_auc = NA, n_tested = 11)
A549−GM130−SQSTM1 (n_incl_in_auc = NA, n_tested = 11)
A549−TP53BP1−CLTA (n_incl_in_auc = NA, n_tested = 11)
A549−TUBA1B−RELA (n_incl_in_auc = NA, n_tested = 11)
HepG2−ACTB−RAB5A (n_incl_in_auc = NA, n_tested = 11)
HepG2−CANX−COX4 (n_incl_in_auc = NA, n_tested = 11)
HepG2−GM130−SQSTM1 (n_incl_in_auc = NA, n_tested = 11)
HepG2−TP53BP1−CLTA (n_incl_in_auc = NA, n_tested = 11)
HepG2−TUBA1B−RELA (n_incl_in_auc = NA, n_tested = 11)
WPMY1−ACTB−RAB5A (n_incl_in_auc = NA, n_tested = 11)
WPMY1−CANX−COX4 (n_incl_in_auc = NA, n_tested = 11)
WPMY1−GM130−SQSTM1 (n_incl_in_auc = 4, n_tested = 11)
WPMY1−TP53BP1−CLTA (n_incl_in_auc = NA, n_tested = 11)
WPMY1−TUBA1B−RELA (n_incl_in_auc = NA, n_tested = 11)
0.0
0.1
0.2
0.3
0.4
0.5
0.6
0.7
0.8
0.9
1.0
auc_max_over_concentr_median
Batch
All Aggr.
Within Batch 1
Within Batch 2
cell_type
A549
HepG2
WPMY1
FDR−adjusted
p−value < 0.05
FALSE
NA
TRUE

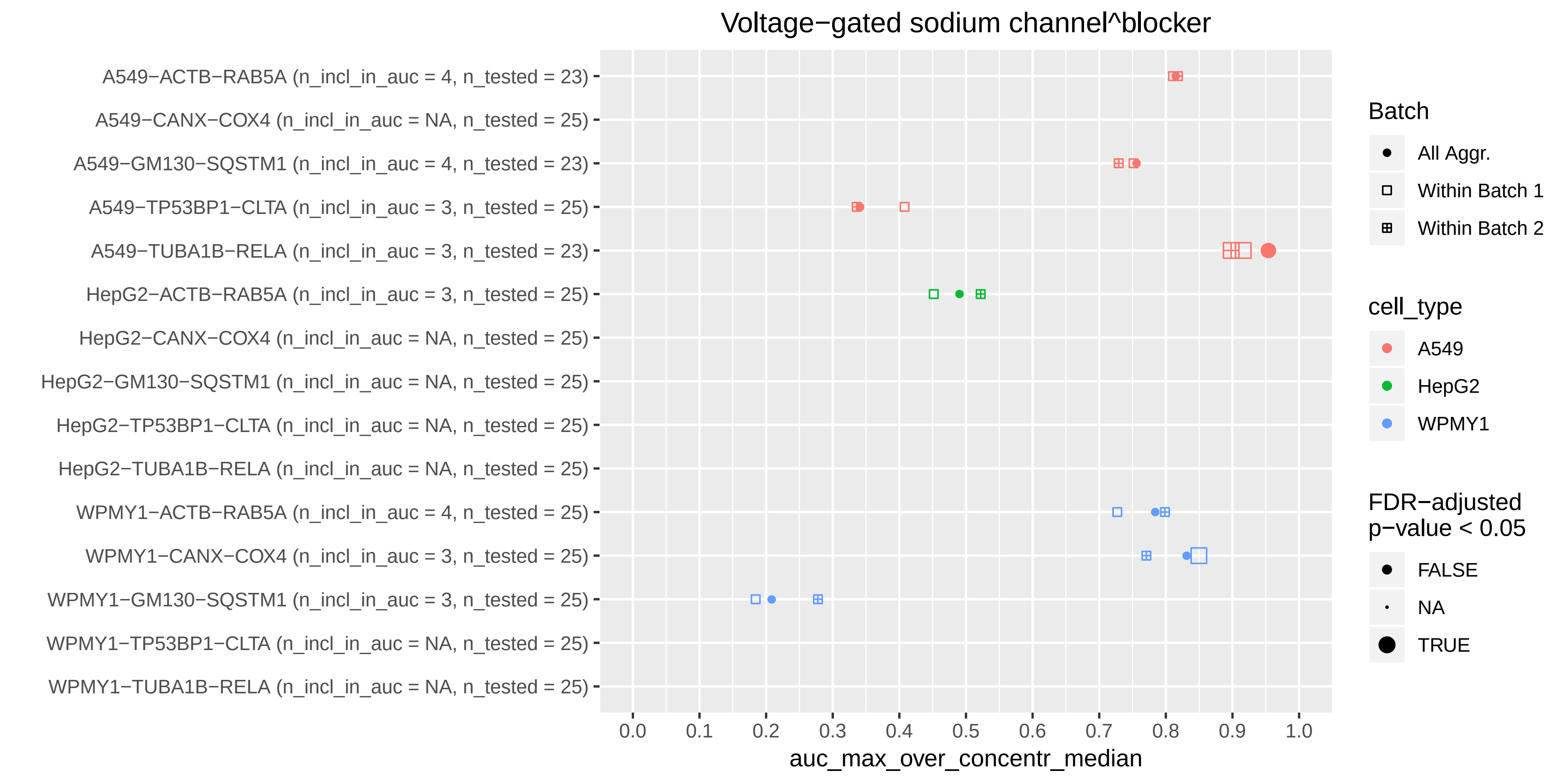
Voltage−gated sodium channel^blocker
A549−ACTB−RAB5A (n_incl_in_auc = 4, n_tested = 23)
A549−CANX−COX4 (n_incl_in_auc = NA, n_tested = 25)
A549−GM130−SQSTM1 (n_incl_in_auc = 4, n_tested = 23)
A549−TP53BP1−CLTA (n_incl_in_auc = 3, n_tested = 25)
A549−TUBA1B−RELA (n_incl_in_auc = 3, n_tested = 23)
HepG2−ACTB−RAB5A (n_incl_in_auc = 3, n_tested = 25)
HepG2−CANX−COX4 (n_incl_in_auc = NA, n_tested = 25)
HepG2−GM130−SQSTM1 (n_incl_in_auc = NA, n_tested = 25)
HepG2−TP53BP1−CLTA (n_incl_in_auc = NA, n_tested = 25)
HepG2−TUBA1B−RELA (n_incl_in_auc = NA, n_tested = 25)
WPMY1−ACTB−RAB5A (n_incl_in_auc = 4, n_tested = 25)
WPMY1−CANX−COX4 (n_incl_in_auc = 3, n_tested = 25)
WPMY1−GM130−SQSTM1 (n_incl_in_auc = 3, n_tested = 25)
WPMY1−TP53BP1−CLTA (n_incl_in_auc = NA, n_tested = 25)
WPMY1−TUBA1B−RELA (n_incl_in_auc = NA, n_tested = 25)
0.0
0.1
0.2
0.3
0.4
0.5
0.6
0.7
0.8
0.9
1.0
auc_max_over_concentr_median
Batch
All Aggr.
Within Batch 1
Within Batch 2
cell_type
A549
HepG2
WPMY1
FDR−adjusted
p−value < 0.05
FALSE
NA
TRUE

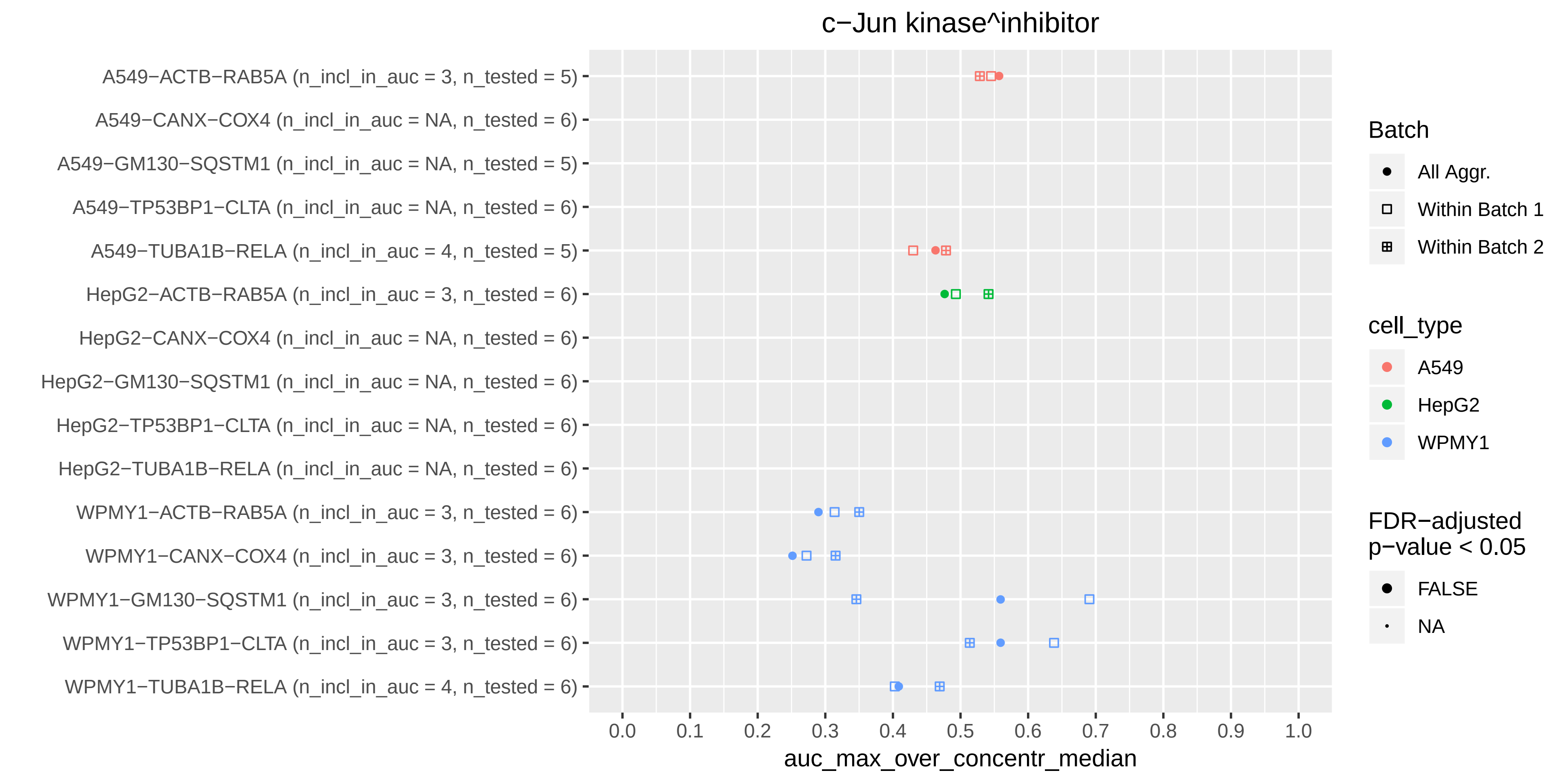
c−Jun kinase^inhibitor
A549−ACTB−RAB5A (n_incl_in_auc = 3, n_tested = 5)
A549−CANX−COX4 (n_incl_in_auc = NA, n_tested = 6)
A549−GM130−SQSTM1 (n_incl_in_auc = NA, n_tested = 5)
A549−TP53BP1−CLTA (n_incl_in_auc = NA, n_tested = 6)
A549−TUBA1B−RELA (n_incl_in_auc = 4, n_tested = 5)
HepG2−ACTB−RAB5A (n_incl_in_auc = 3, n_tested = 6)
HepG2−CANX−COX4 (n_incl_in_auc = NA, n_tested = 6)
HepG2−GM130−SQSTM1 (n_incl_in_auc = NA, n_tested = 6)
HepG2−TP53BP1−CLTA (n_incl_in_auc = NA, n_tested = 6)
HepG2−TUBA1B−RELA (n_incl_in_auc = NA, n_tested = 6)
WPMY1−ACTB−RAB5A (n_incl_in_auc = 3, n_tested = 6)
WPMY1−CANX−COX4 (n_incl_in_auc = 3, n_tested = 6)
WPMY1−GM130−SQSTM1 (n_incl_in_auc = 3, n_tested = 6)
WPMY1−TP53BP1−CLTA (n_incl_in_auc = 3, n_tested = 6)
WPMY1−TUBA1B−RELA (n_incl_in_auc = 4, n_tested = 6)
0.0
0.1
0.2
0.3
0.4
0.5
0.6
0.7
0.8
0.9
1.0
auc_max_over_concentr_median
Batch
All Aggr.
Within Batch 1
Within Batch 2
cell_type
A549
HepG2
WPMY1
FDR−adjusted
p−value < 0.05
FALSE
NA

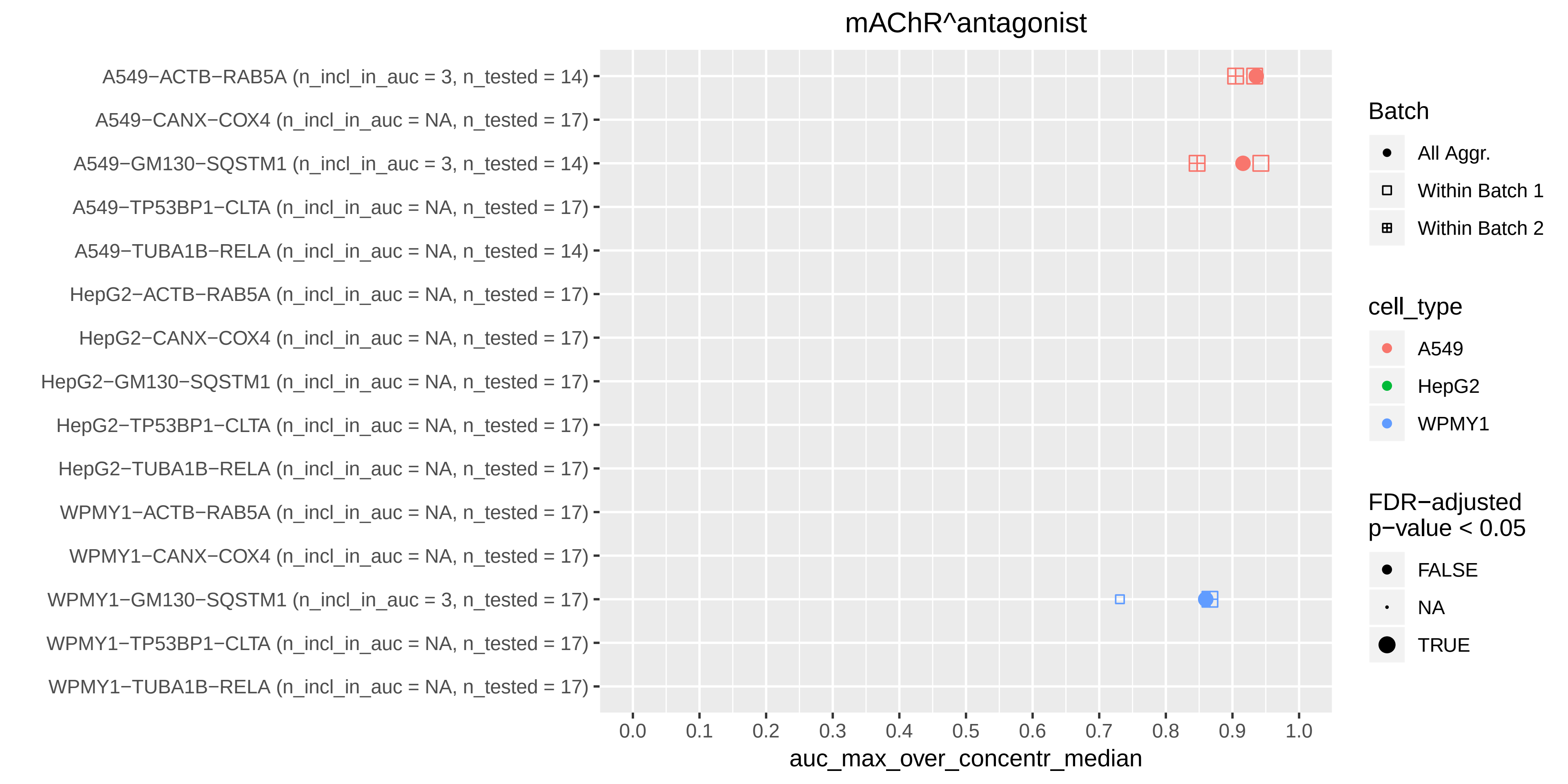

mAChR^antagonist
A549−ACTB−RAB5A (n_incl_in_auc = 3, n_tested = 14)
A549−CANX−COX4 (n_incl_in_auc = NA, n_tested = 17)
A549−GM130−SQSTM1 (n_incl_in_auc = 3, n_tested = 14)
A549−TP53BP1−CLTA (n_incl_in_auc = NA, n_tested = 17)
A549−TUBA1B−RELA (n_incl_in_auc = NA, n_tested = 14)
HepG2−ACTB−RAB5A (n_incl_in_auc = NA, n_tested = 17)
HepG2−CANX−COX4 (n_incl_in_auc = NA, n_tested = 17)
HepG2−GM130−SQSTM1 (n_incl_in_auc = NA, n_tested = 17)
HepG2−TP53BP1−CLTA (n_incl_in_auc = NA, n_tested = 17)
HepG2−TUBA1B−RELA (n_incl_in_auc = NA, n_tested = 17)
WPMY1−ACTB−RAB5A (n_incl_in_auc = NA, n_tested = 17)
WPMY1−CANX−COX4 (n_incl_in_auc = NA, n_tested = 17)
WPMY1−GM130−SQSTM1 (n_incl_in_auc = 3, n_tested = 17)
WPMY1−TP53BP1−CLTA (n_incl_in_auc = NA, n_tested = 17)
WPMY1−TUBA1B−RELA (n_incl_in_auc = NA, n_tested = 17)
0.0
0.1
0.2
0.3
0.4
0.5
0.6
0.7
0.8
0.9
1.0
auc_max_over_concentr_median
Batch
All Aggr.
Within Batch 1
Within Batch 2
cell_type
A549
HepG2
WPMY1
FDR−adjusted p−value < 0.05
FALSE
NA
TRUE

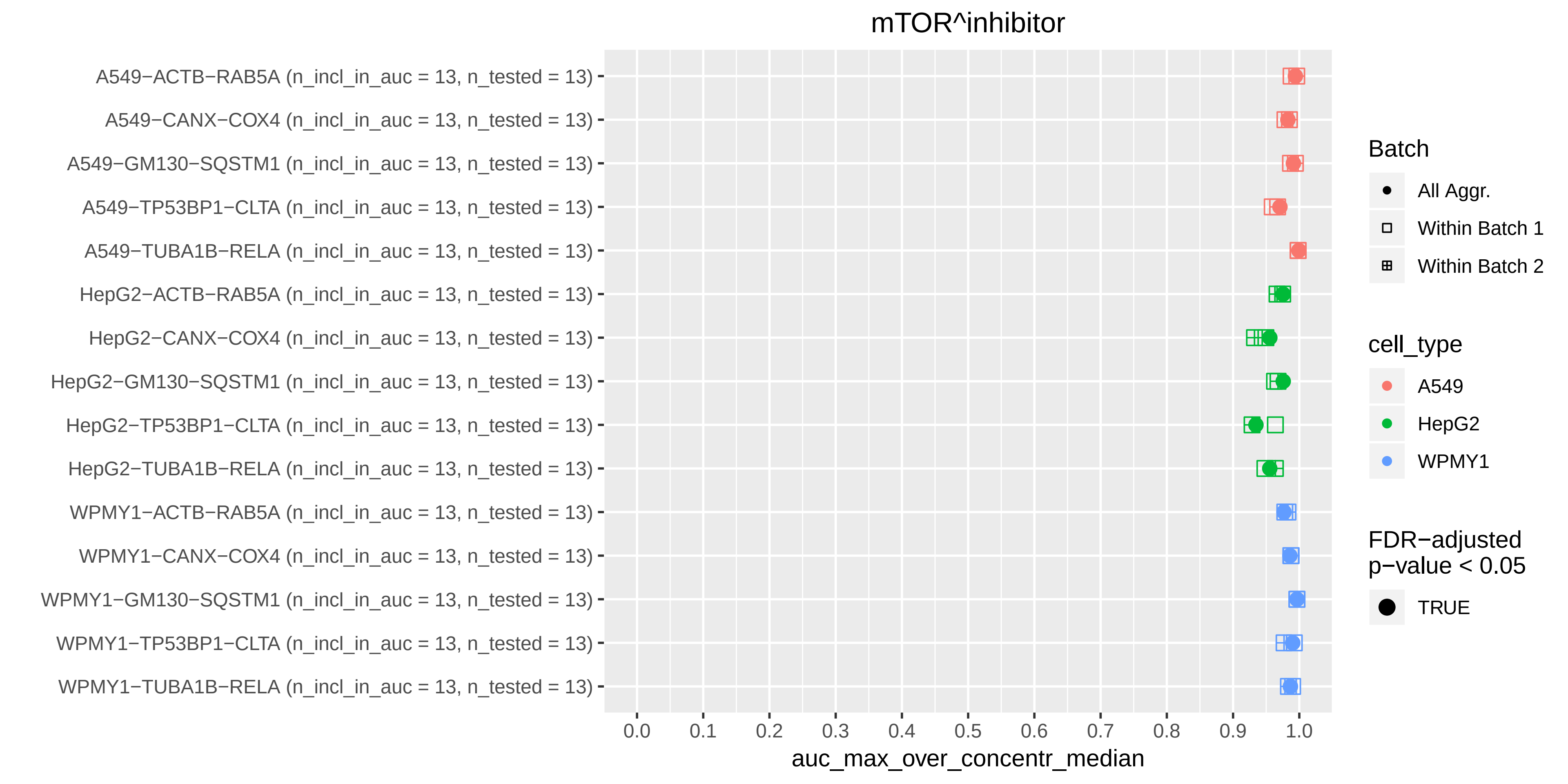

mTOR^inhibitor
A549−ACTB−RAB5A (n_incl_in_auc = 13, n_tested = 13)
A549−CANX−COX4 (n_incl_in_auc = 13, n_tested = 13)
A549−GM130−SQSTM1 (n_incl_in_auc = 13, n_tested = 13)
A549−TP53BP1−CLTA (n_incl_in_auc = 13, n_tested = 13)
A549−TUBA1B−RELA (n_incl_in_auc = 13, n_tested = 13)
HepG2−ACTB−RAB5A (n_incl_in_auc = 13, n_tested = 13)
HepG2−CANX−COX4 (n_incl_in_auc = 13, n_tested = 13)
HepG2−GM130−SQSTM1 (n_incl_in_auc = 13, n_tested = 13)
HepG2−TP53BP1−CLTA (n_incl_in_auc = 13, n_tested = 13)
HepG2−TUBA1B−RELA (n_incl_in_auc = 13, n_tested = 13)
WPMY1−ACTB−RAB5A (n_incl_in_auc = 13, n_tested = 13)
WPMY1−CANX−COX4 (n_incl_in_auc = 13, n_tested = 13)
WPMY1−GM130−SQSTM1 (n_incl_in_auc = 13, n_tested = 13)
WPMY1−TP53BP1−CLTA (n_incl_in_auc = 13, n_tested = 13)
WPMY1−TUBA1B−RELA (n_incl_in_auc = 13, n_tested = 13)
0.0
0.1
0.2
0.3
0.4
0.5
0.6
0.7
0.8
0.9
1.0
auc_max_over_concentr_median
Batch
All Aggr.
Within Batch 1
Within Batch 2
cell_type
A549
HepG2
WPMY1
FDR−adjusted p−value < 0.05
TRUE

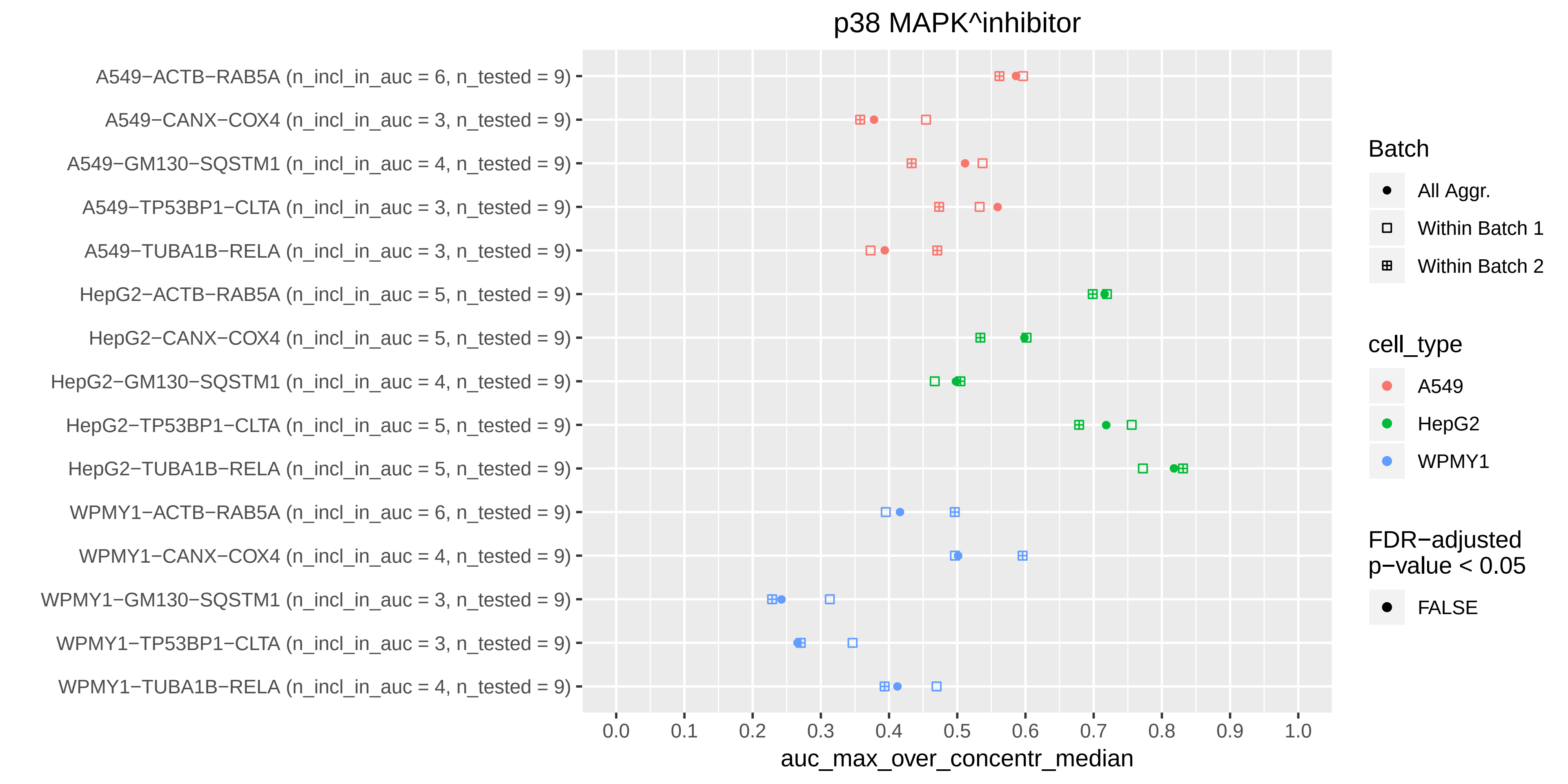
p38 MAPK^inhibitor
A549−ACTB−RAB5A (n_incl_in_auc = 6, n_tested = 9)
A549−CANX−COX4 (n_incl_in_auc = 3, n_tested = 9)
A549−GM130−SQSTM1 (n_incl_in_auc = 4, n_tested = 9)
A549−TP53BP1−CLTA (n_incl_in_auc = 3, n_tested = 9)
A549−TUBA1B−RELA (n_incl_in_auc = 3, n_tested = 9)
HepG2−ACTB−RAB5A (n_incl_in_auc = 5, n_tested = 9)
HepG2−CANX−COX4 (n_incl_in_auc = 5, n_tested = 9)
HepG2−GM130−SQSTM1 (n_incl_in_auc = 4, n_tested = 9)
HepG2−TP53BP1−CLTA (n_incl_in_auc = 5, n_tested = 9)
HepG2−TUBA1B−RELA (n_incl_in_auc = 5, n_tested = 9)
WPMY1−ACTB−RAB5A (n_incl_in_auc = 6, n_tested = 9)
WPMY1−CANX−COX4 (n_incl_in_auc = 4, n_tested = 9)
WPMY1−GM130−SQSTM1 (n_incl_in_auc = 3, n_tested = 9)
WPMY1−TP53BP1−CLTA (n_incl_in_auc = 3, n_tested = 9)
WPMY1−TUBA1B−RELA (n_incl_in_auc = 4, n_tested = 9)
0.0
0.1
0.2
0.3
0.4
0.5
0.6
0.7
0.8
0.9
1.0
auc_max_over_concentr_median
Batch
All Aggr.
Within Batch 1
Within Batch 2
cell_type
A549
HepG2
WPMY1
FDR−adjusted p−value < 0.05
FALSE

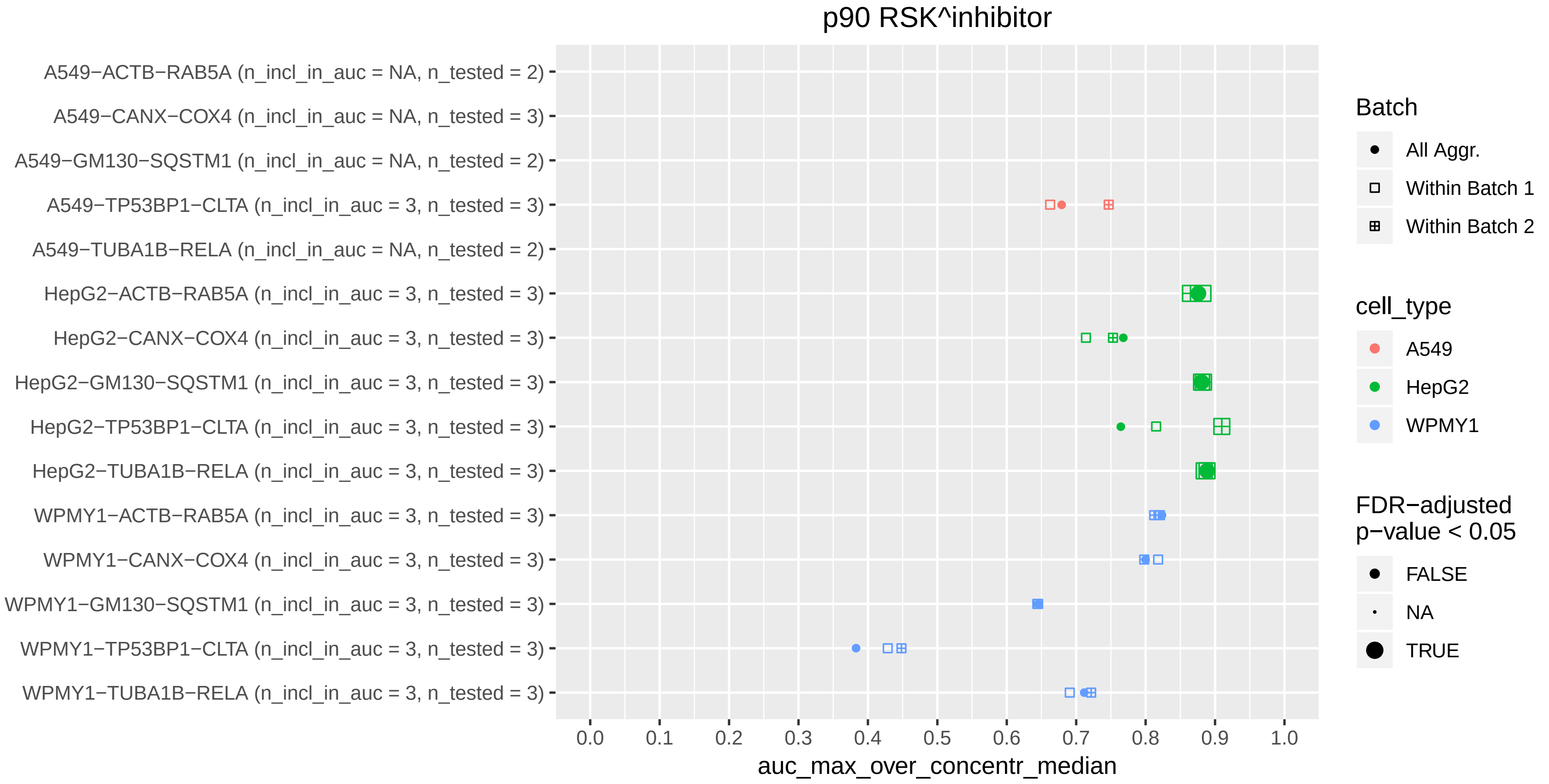

p90 RSK^inhibitor
A549−ACTB−RAB5A (n_incl_in_auc = NA, n_tested = 2)
A549−CANX−COX4 (n_incl_in_auc = NA, n_tested = 3)
A549−GM130−SQSTM1 (n_incl_in_auc = NA, n_tested = 2)
A549−TP53BP1−CLTA (n_incl_in_auc = 3, n_tested = 3)
A549−TUBA1B−RELA (n_incl_in_auc = NA, n_tested = 2)
HepG2−ACTB−RAB5A (n_incl_in_auc = 3, n_tested = 3)
HepG2−CANX−COX4 (n_incl_in_auc = 3, n_tested = 3)
HepG2−GM130−SQSTM1 (n_incl_in_auc = 3, n_tested = 3)
HepG2−TP53BP1−CLTA (n_incl_in_auc = 3, n_tested = 3)
HepG2−TUBA1B−RELA (n_incl_in_auc = 3, n_tested = 3)
WPMY1−ACTB−RAB5A (n_incl_in_auc = 3, n_tested = 3)
WPMY1−CANX−COX4 (n_incl_in_auc = 3, n_tested = 3)
WPMY1−GM130−SQSTM1 (n_incl_in_auc = 3, n_tested = 3)
WPMY1−TP53BP1−CLTA (n_incl_in_auc = 3, n_tested = 3)
WPMY1−TUBA1B−RELA (n_incl_in_auc = 3, n_tested = 3)
0.0
0.1
0.2
0.3
0.4
0.5
0.6
0.7
0.8
0.9
1.0
auc_max_over_concentr_median
Batch
All Aggr.
Within Batch 1
Within Batch 2
cell_type
A549
HepG2
WPMY1
FDR−adjusted p−value < 0.05
FALSE
NA
TRUE
